# RBM25 selectively stabilizes *XBP1s* mRNA independently of splicing factor activity

**DOI:** 10.64898/2026.11.04.764219

**Authors:** Anna Więch-Walów, Anna Barton, Michał Tracz, Sylwia Kozioł, Joanna Grzyb, Paulina Czechowicz, James F Collawn, Eric Chevet, Rafal Bartoszewski

## Abstract

The unfolded protein response (UPR) preserves endoplasmic reticulum (ER) proteostasis through coordinated transcriptional and translational programs that promote adaptation and prevents apoptosis. A central component of this response is the IRE1-XBP1 signaling axis, in which IRE1-mediated unconventional splicing of *XBP1* mRNA generates the active transcription factor XBP1s. While the mechanisms governing IRE1 activation and *XBP1* splicing have been extensively characterized, the post-transcriptional processes that sustain XBP1s expression remain poorly understood. Here, we identify RNA-binding motif protein 25 (RBM25) as a key post-transcriptional regulator of the UPR that selectively stabilizes spliced *XBP1* mRNA during ER stress. RBM25 binds the *XBP1s* 3′ untranslated region and prolongs *XBP1s* mRNA half-life without affecting unspliced *XBP1u* transcripts or IRE1-dependent regulated decay (RIDD) targets. Consequently, RBM25 depletion diminishes XBP1 transcriptional output, compromises adaptive UPR signaling, and sensitizes cells to ER stress-induced death across multiple stress paradigms. Together, our findings uncover a previously unrecognized RNA-based regulatory checkpoint that amplifies and sustains XBP1 signaling, thereby tuning the magnitude of the adaptive UPR and cellular survival under proteotoxic stress.

## INTRODUCTION

The unfolded protein response (UPR) is a conserved adaptive signaling pathway that maintains endoplasmic reticulum (ER) homeostasis when the protein folding demand exceeds ER capacity. In mammalian cells, the UPR is mediated by three ER resident transmembrane sensors—inositol requiring enzyme 1α (IRE1α), protein kinase RNA like ER kinase (PERK), and activating transcription factor 6 (ATF6). These proteins coordinate reductions in the ER protein load, the expansion of folding capacity, and the transcriptional reprogramming to restore ER homeostasis (1,2). When adaptive programs fail, prolonged UPR signaling induces cell death (1,2). Central to this response is IRE1α, the most evolutionary conserved ER stress sensor, whose endoribonuclease activity, together with the tRNA ligase RtcB catalyzes an unconventional splicing reaction that converts X box binding protein 1 (*XBP1*) mRNA into a translationally frameshifted transcript that encodes the potent transcription factor *XBP1s* (*3*). XBP1s drives adaptive programs to address ER homeostasis imbalance through the expression of ER chaperones, ER associated degradation components, and genes required for ER expansion, which are essential for mammalian cell viability (4,5).

In parallel, activated IRE1α degrades a subset of mRNAs and microRNAs through regulated IRE1 dependent decay (RIDD), a mechanism that can exert both adaptive and pro apoptotic effects (6–9). However, even though mammalian *XBP1s* mRNA is not a RIDD substrate (6–8), productive XBP1s output must be ensured in a stress-constrained environment characterized by elevated RNA turnover and attenuated translation (6). Sustained IRE1α signaling therefore presents the quantitative challenge of maintaining sufficient levels of the spliced *XBP1* mRNA transcript to produce enough XBP1s and drive adaptive transcriptional programs. How mammalian cells achieve robust XBP1s accumulation under these conditions has remained poorly characterized.

We reasoned that a mechanism to maintain *XBP1s* mRNA expression following splicing might involve RNA binding proteins (RBPs). Indeed, the 3′ untranslated region (3′UTR) of *XBP1* mRNA, although dispensable for splicing in mammals, is a critical determinant of *XBP1s* mRNA stability (10–12). In the same line in *Drosophila*, the RNA binding protein Pumilio binds *Xbp1* mRNA to potentiate *Xbp1s* expression by modulating RNA stability, highlighting an evolutionarily conserved role for RNA binding proteins (RBPs) in regulating UPR output (13). These observations raise the possibility that, in mammalian cells, RBPs could provide a distinct post transcriptional layer of control that ensures sufficient XBP1s accumulation.

To test this hypothesis, we used an unbiased, crosslinking-independent RNA affinity-mass spectrometry approach to identify human proteins associating with the 3′UTRs of spliced and unspliced *XBP1* transcripts under basal and ER stress conditions in HeLa cells. This approach identified RNA-binding motif protein 25 (RBM25) as a stress-enhanced interactor selectively recognizing *XBP1s* 3′UTR. Mechanistically, RBM25 functions as a constitutive RNA-binding factor that stabilizes *XBP1s* mRNA, thereby maintaining sufficient transcript levels to support robust IRE1-dependent UPR signaling and promote cell survival upon ER stress.

## RESULTS

### RBM25 selectively regulates XBP1s expression during ER stress

To identify RNA-binding proteins (RBPs) that interact with XBP1s 3′ untranslated regions (3′UTRs), we performed an unbiased RNA affinity pulldown–mass spectrometry screen using the 5′-biotinylated 3′ untranslated regions (3′UTRs) of both unspliced and spliced *XBP1* transcripts (*XBP1u* and *XBP1s*) as baits (14,15). In vitro–transcribed RNA baits were incubated, without crosslinking, at room temperature with clarified cytoplasmic lysates from HeLa S3 cells maintained under basal conditions or subjected to tunicamycin-induced ER stress. RNA–protein complexes were captured using streptavidin-coated magnetic beads and analyzed by bottom-up mass spectrometry–based proteomic profiling (**Figure 1A**).

**Figure 1.**
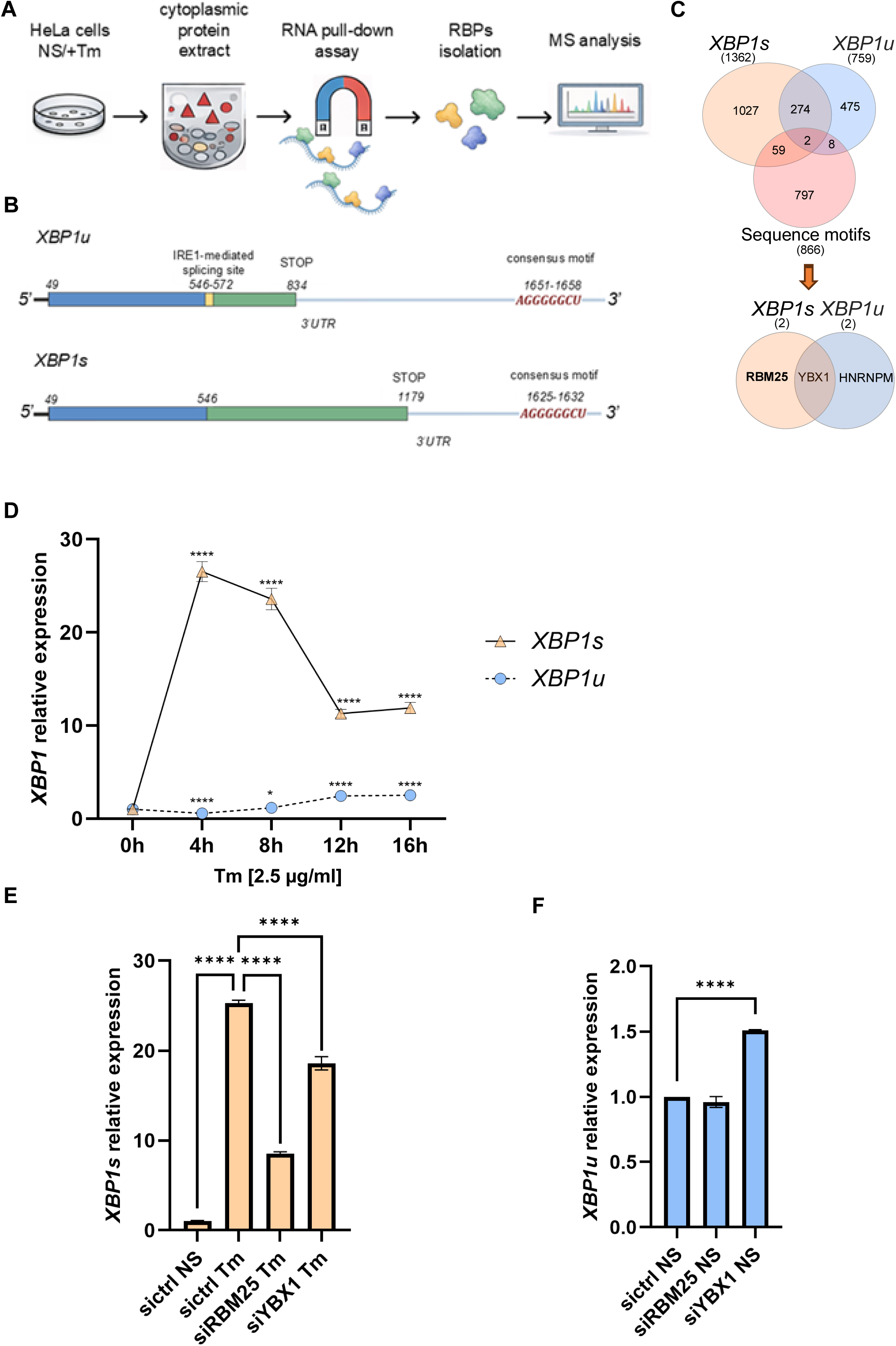
Proteomic mapping of XBP1 3′UTR-interacting RNA⍰binding proteins. **A.** Workflow illustrating the identification of proteins associating with *XBP1* mRNA during ER stress. Cytoplasmic extracts from HeLa S3 cells cultured in the presence or absence of tunicamycin (Tm) were subjected to RNA affinity purification using biotinylated transcripts corresponding to the 3′UTRs of *XBP1u* or *XBP1s*. RNA-protein complexes were recovered with streptavidin⍰coated magnetic beads, and bound RNA⍰binding proteins were subsequently purified and identified by mass spectrometry. **B.** Schematic representation of *XBP1u* and *XBP1s* mRNAs. *XBP1u* contains a 5′ coding region, the IRE1 cleavage site (nt 546-572), a stop codon at nt 834, and a 3′UTR harboring a conserved G⍰rich element (nt 1651-1658). *XBP1s* is generated by unconventional splicing beginning at nt 49 and includes the spliced junction, a stop codon at nt 1179, and a corresponding G⍰rich element within the 3′UTR (nt 1625-1632). These regions contain G⍰rich sequences identified in this study that are consistent with the reported binding preference of RBM25. **C.** Venn diagram illustrating the selection of proteins identified by the MS⍰based approach. Following the initial identification of proteins associated with the *XBP1s* and *XBP1u* 3′UTRs, candidates were filtered based on their enrichment profiles under condition⍰specific contexts (i.e., enrichment with XBP1u in non⍰stressed samples and with XBP1s under ER stress), UniProt annotation score, and the presence of predicted RNA binding motifs specifically located within the *XBP1* 3′UTRs, as identified using the RBPmap algorithm (see Supplementary Dataset 1). **D.** HeLa S3 cells transfected with control siRNA were treated with Tm (0.5 μg/ml) for the indicated times, and total RNA was isolated. *XBP1s* and *XBP1u* transcript levels were quantified by RT-qPCR, normalized to *RPLP0*, and expressed relative to unstressed controls. Data represent mean ± s.d. from three independent experiments. Statistical significance was determined using two⍰tailed tests (*P < 0.05, **P < 0.01, ***P < 0.001, n=6). **E** Quantitative RT-PCR analysis of *XBP1* transcripts in HeLa S3 cells transfected with control siRNA (sictrl) or *RBM25*⍰targeting siRNA (siRBM25) or *YBX1*-targeting siRNA (siYBX1) and treated with tunicamycin (Tm). Tm markedly induced *XBP1s* mRNA levels in sictrl cells, whereas *RBM25* depletion significantly attenuated this response (****P < 0.0001, n=6). **F**. Basal *XBP1u* mRNA levels were not affected by RBM25 depletion, and no differences were observed between untreated conditions (****P < 0.0001, n=6).

To prioritize candidates identified by proteomic profiling that are likely to directly interact with the *XBP1* 3′UTRs, proteins detected by mass spectrometry were further filtered based on their enrichment profiles under condition-specific contexts (i.e., enrichment with *XBP1u* in non-stressed samples and with *XBP1s* under ER stress), as well as the presence of predicted RNA-binding motifs specifically located within the *XBP1* 3′UTRs, using the RBPmap algorithm (16). This integrative analysis yielded two candidate RBPs: Y-box binding protein 1 (YBX1; UniProt P67809) and RNA-binding motif protein 25 (RBM25; UniProt P49756) (**Figure 1C)**.

To assess the potential roles of both YBX1 and RBM25 on XBP1s mRNA regulation, we first characterized the kinetics of *XBP1* mRNA isoforms (unspliced and spliced) expression during tunicamycin (Tm) induced ER stress in HeLa S3 cells (in the presence of control siRNA) (**Figure 1D**). In HeLa S3 cells, *XBP1s* accumulation peaked between 4 and 8 hours of ER stress, whereas *XBP1u* mRNA levels increased progressively up to 16 hours, consistent with previous reports [4].

We next performed siRNA-mediated knockdown of YBX1 or RBM25 and monitored *XBP1u* and *XBP1*s mRNA and protein abundance following 8 hours of ER stress induction with tunicamycin (**Figure 1E**). Strikingly, *RBM25* depletion selectively and robustly suppressed *XBP1s* expression at both the mRNA and protein levels (**Figure 1F**).

The magnitude of XBP1s protein reduction upon *RBM25* knockdown was comparable to that observed following pharmacological inhibition of IRE1 RNase activity (4µ8C). In contrast, silencing of *YBX1* resulted in only a modest increase in *XBP1u* and reduction in *XBP1s* transcript abundance, which was insufficient to significantly impact protein expression (**Figures 2** and **S Supplementary Fig. 1**). Importantly, *RBM25* knockdown had no effect on *XBP1u* mRNA levels, indicating isoform-specific regulation. To assess whether the effects of RBM25 on XBP1s mRNA might be indirectly mediated through altered IRE1 activity, we examined the expression of a canonical regulated IRE1 dependent decay (RIDD) substrate, diacylglycerol O acyltransferase 2 (*DGAT2*). As expected, *DGAT2* mRNA levels were reduced upon tunicamycin treatment and restored by pharmacological inhibition of IRE1 RNase. Notably, *RBM25* silencing did not affect *DGAT2* mRNA expression in ER stressed cells (**Figure 2C**), indicating that RBM25 regulates XBP1s independently of RIDD. Together, these findings identify RBM25 as a selective post transcriptional regulator of XBP1s mRNA expression which might operate through direct interaction between RBM25 and *XBP1s* 3’ UTR.

**Figure 2.**
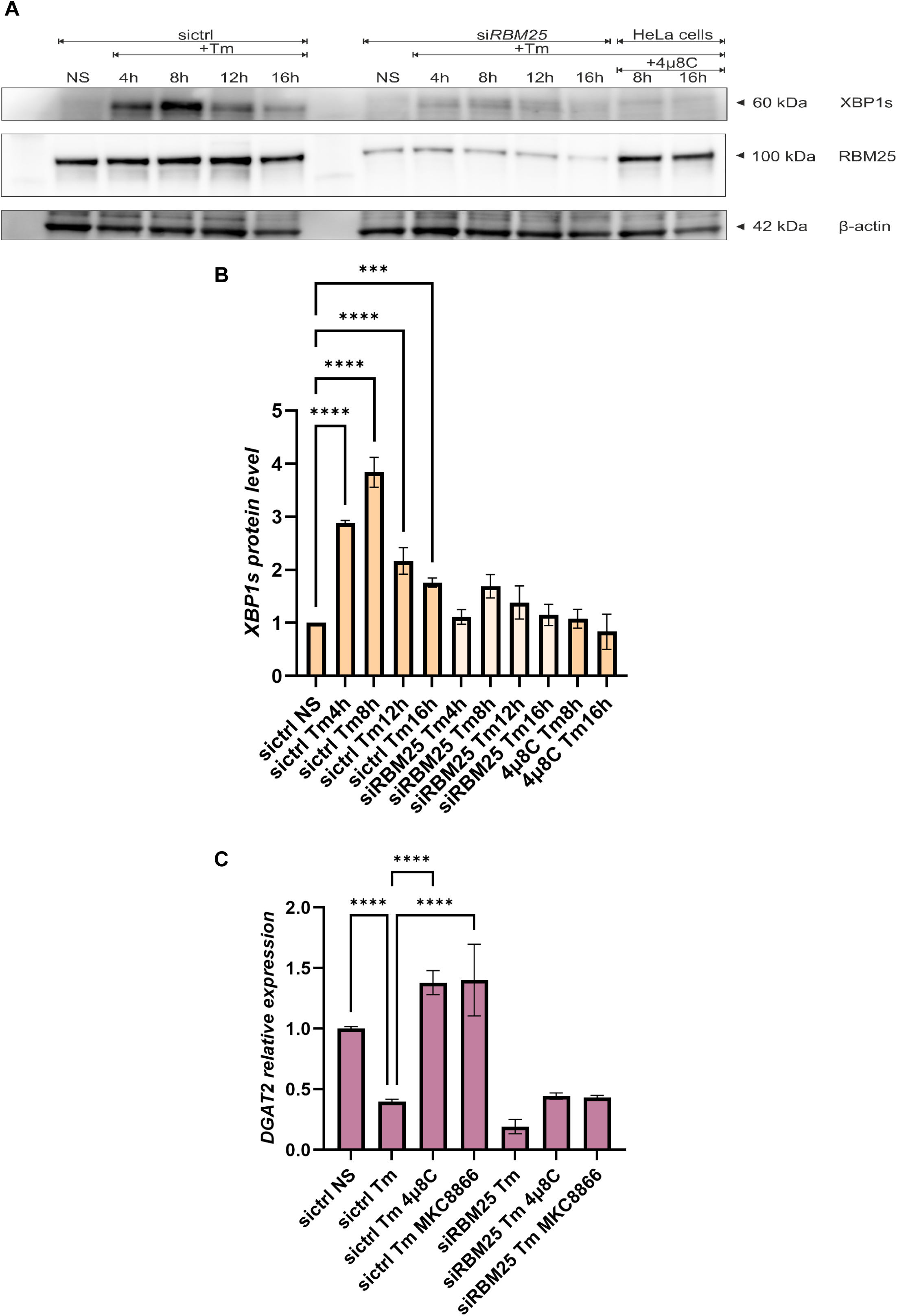
RBM25 depletion selectively impairs XBP1s expression during ER stress. **A.B.** Immunoblot analysis showing time⍰dependent accumulation of XBP1s protein following Tm treatment (4-16 h) in sictrl cells, which was markedly reduced upon RBM25 knockdown. β actin or total protein levels served as a loading and normalization control (**Supplementary** Figure 2). RBM25 protein levels remained stable upon Tm treatment in control cells, and efficient knockdown was confirmed by loss of the ∼100⍰kDa RBM25 band. Treatment with the IRE1 RNase inhibitor 4μ8C suppressed XBP1s accumulation, validating pathway specificity. **C**. Quantitative RT-PCR analysis of the RIDD target *DGAT2*. Under basal conditions, *DGAT2* mRNA levels were reduced in siRBM25 cells relative to sictrl (****P < 0.0001). Tm treatment significantly decreased *DGAT2* expression in sictrl cells (****P < 0.0001), with a comparable reduction observed upon *RBM25* depletion (**P < 0.01), indicating preserved RIDD activity. Transcript levels were normalized to *RPLP0* and expressed relative to unstressed controls. Data represent mean ± s.e.m. from three independent experiments. Statistical significance was determined using two tailed tests (*P < 0.05, **P < 0.01, ***P < 0.001, ****P < 0.0001, n=3).

### RBM25 directly associates with *XBP1* mRNA via its 3′UTR

To evaluate the direct interaction between RBM25 and *XBP1* mRNA that was suggested by the proteomic screen, we assessed RBM25 binding to either unspliced or spliced *XBP1* transcripts using RNA immunoprecipitation (RIP). Both *XBP1u* and *XBP1s* 3′UTRs contain conserved G rich RNA elements, including the sequence AGGGGGCU identified in our analysis (**Figure 1B**), which is consistent with the known preference of RBM25 for G rich binding motifs, suggesting that either isoform could serve as an RNA ligand (17,18). RIP assays using an anti RBM25 antibody were performed in HeLa S3 cells under basal conditions and following tunicamycin induced ER stress. Immunoprecipitation efficiency and specificity were confirmed by the selective enrichment of a ∼100 kDa RBM25 band in anti RBM25 samples by immunoblotting, while total protein staining verified comparable loading in the different experimental conditions (**Figure 3A)**.

**Figure 3.**
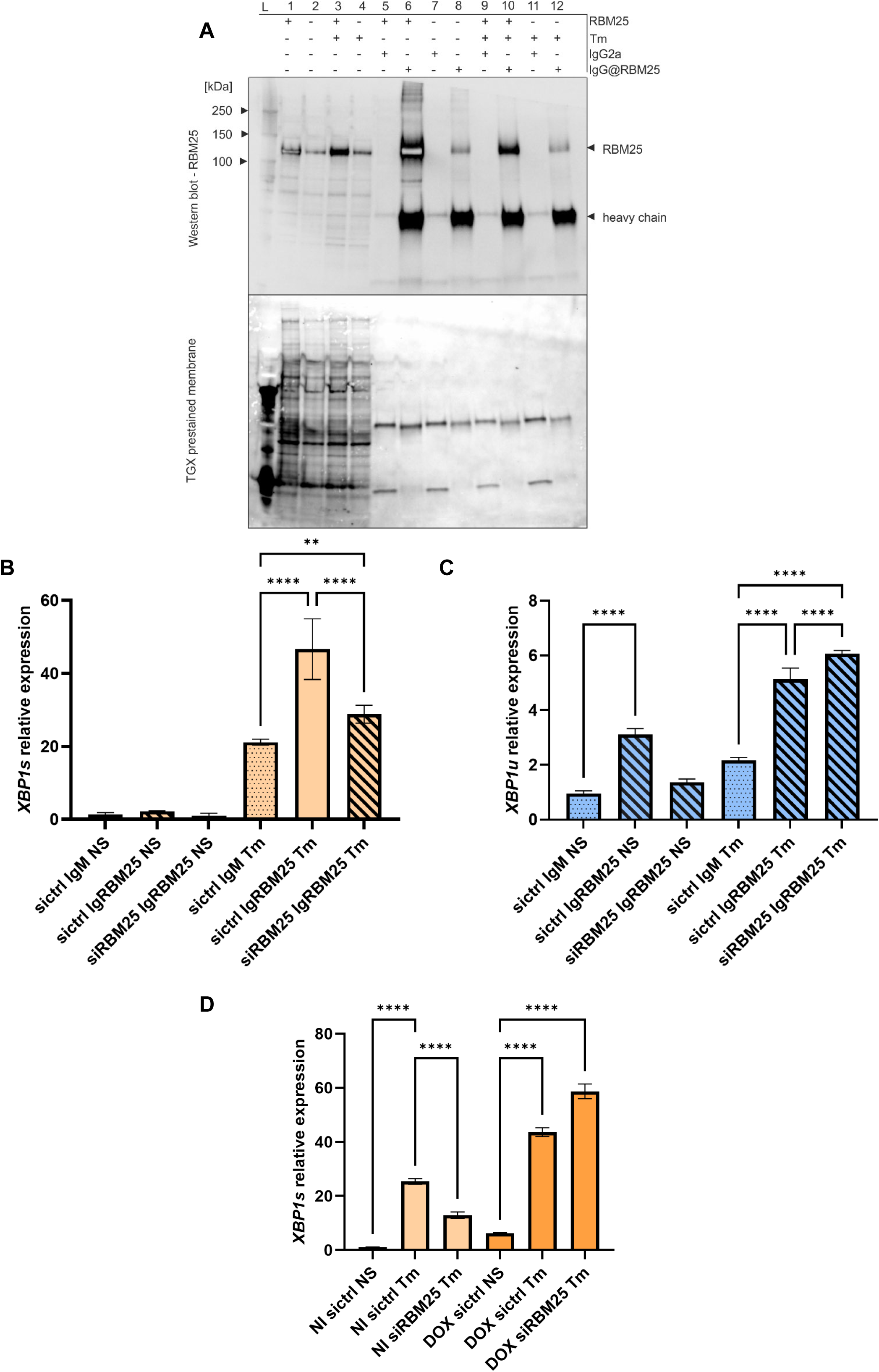
RBM25 associates with *XBP1s* mRNA under basal and ER stress conditions. **A.**RNA immunoprecipitation (RIP) analysis of RBM25 in HeLa S3 cells under non⍰stressed (NS) or tunicamycin (Tm)⍰treated conditions. Comparable RBM25 enrichment was observed in both conditions. Immunoblotting revealed a distinct ∼100⍰kDa RBM25 band in RBM25 immunoprecipitates, which was absent in IgG2a controls. Total protein staining confirmed equivalent input across samples. **B.** Quantitative RT-PCR analysis of *XBP1s* mRNA following RIP. Enrichment of *XBP1s* transcripts was observed in RBM25 immunoprecipitates relative to IgG2a controls, indicating a direct association between RBM25 and *XBP1s* mRNA in both sictrl (white) and siRBM25 (blue) cells. **C.** Quantitative RT-PCR analysis of *XBP1u* RNA following RIP. Although enrichment of *XBP1u* transcripts was detectable in RBM25 immunoprecipitates, the magnitude of enrichment was substantially lower than that observed for *XBP1s*, particularly under Tm treatment. **D**. Quantitative RT-PCR analysis of *XBP1s* expression in parental and inducible HeLa S3 XBP1s cell lines. In the absence of DOX, *XBP1s* levels under NS and Tm conditions were comparable between parental and inducible cells. Tm treatment significantly induced XBP1s expression in sictrl cells (****P < 0.0001), whereas *RBM25* knockdown markedly attenuated this response (****P < 0.0001). DOX treatment increased basal *XBP1s* levels and further enhanced Tm induced expression, as expected. Data represent mean ± s.e.m. from three independent experiments. Transcript levels were normalized to *RPLP0* and expressed relative to unstressed controls. Statistical significance was determined using two tailed tests (*P < 0.05, **P < 0.01, ***P < 0.001, ****P < 0.0001, n=3).

Quantification of co immunoprecipitated RNAs using isoform specific TaqMan probes revealed robust and selective enrichment of *XBP1s* mRNA in RBM25 immunoprecipitates under ER stress conditions (**Figure 3B**). In contrast, *XBP1u* mRNA was enriched to a lesser extent and exhibited detectable association with RBM25 under both basal and stressed conditions (**Figure 3C**). Importantly, enrichment of both transcripts was strongly reduced upon RBM25 knockdown, confirming the specificity of the interaction. These data demonstrate that RBM25 directly associates with *XBP1* mRNA, with preferential binding to the spliced isoform during ER stress.

To determine whether RBM25 binding and regulation of XBP1s depends on the *XBP1* 3′ UTR, we next exploited HeLa S3 cells engineered to express exogenous *XBP1s* lacking the native 3′UTR in an inducible manner (19). In these cells, endogenous *XBP1s* was robustly induced by ER stress and was markedly reduced upon *RBM25* depletion. In contrast, expression of the exogenous, 3′UTR deficient *XBP1s* transcript was not reduced by *RBM25* knockdown and showed a slight increase under ER stress conditions (**Figure 3D**). Together with the RIP data, these results demonstrate that RBM25 directly binds to the *XBP1* 3′UTR and selectively regulates XBP1s mRNA expression in a 3′UTR dependent fashion.

### RBM25 selectively stabilizes spliced *XBP1* mRNA during ER stress

To determine whether RBM25 binding to *XBP1s* mRNA affects transcript stability during ER stress, we examined the decay kinetics of *XBP1s* and *XBP1u* mRNAs in the presence or absence of RBM25. ER stress was induced with tunicamycin, after which global transcription was blocked and IRE1 RNase activity was pharmacologically inhibited to enable accurate assessment of *XBP1u* stability. In parallel, glyceraldehyde 3-phosphate dehydrogenase (*GAPDH*) mRNA decay was measured as a control for assay performance and global RNA turnover. Both XBP1 isoforms (u and s) exhibited relatively rapid turnover under ER stress conditions, with a half life of about 3.5 h for *XBP1u* and 2.9 h for *XBP1s*, whereas *GAPDH* mRNA was substantially more stable, with a half life of about 8.5 h, consistent with previous reports (11,20). Strikingly, RBM25 depletion selectively reduced the half-life of *XBP1s* mRNA (from 2.9 h to 1.8 h), while exerting no detectable effect on *XBP1u* or *GAPDH* mRNA stability (**Figure 4**). These results demonstrate that RBM25 selectively stabilizes *XBP1s* mRNA during ER stress, providing a mechanistic basis for the reduced XBP1s accumulation observed upon RBM25 silencing.

**Figure 4.**
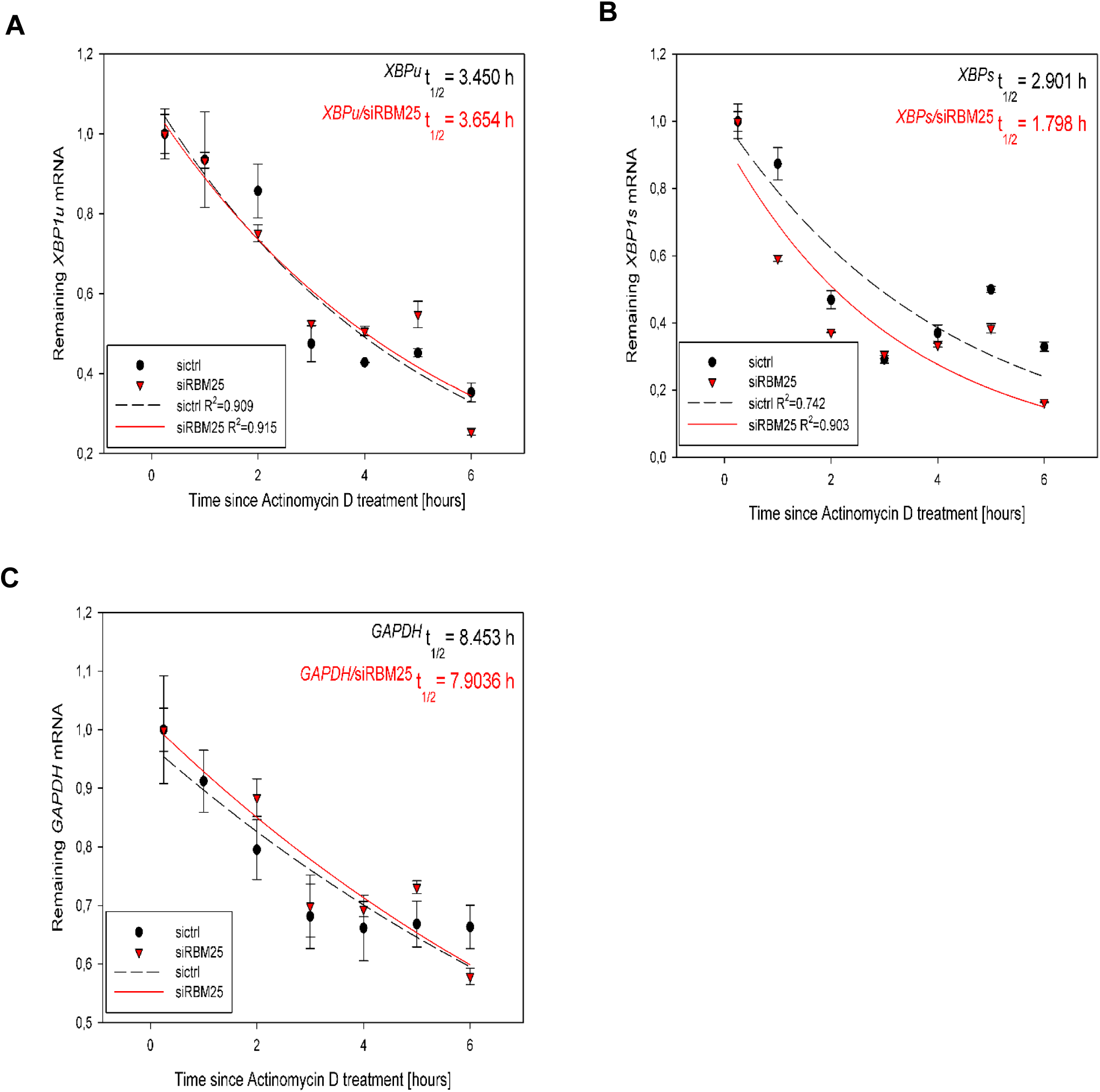
RBM25 selectively regulates the stability of *XBP1s* mRNA. HeLa S3 cells were transfected with control siRNA (sictrl) or *RBM25*⍰targeting siRNA (siRBM25), followed by treatment with tunicamycin (Tm), transcriptional blockade with actinomycin D, or inhibition of IRE1. mRNA abundance was measured over time by RT-qPCR. Transcript levels are expressed as the fraction remaining relative to time zero (C/C₀), and exponential decay curves were fitted for each condition. **A.** Decay analysis of *XBP1u* mRNA revealed comparable stability between sictrl and siRBM25 cells. **B.** Knockdown of *RBM25* markedly accelerated the decay of *XBP1s* mRNA, indicating a role in stabilizing the spliced transcript. **C.** *GAPDH* mRNA stability was unaffected by *RBM25* depletion, with overlapping decay profiles observed under both conditions. Transcripts levels were normalized to *RPLP0* RNA. Data represent mean ± s.e.m. from three independent experiments (n=3). Parameters of exponential decay fitting and associated statistical analyses are provided in **Supplementary Dataset 2.**

### RBM25 stabilizes *XBP1s* mRNA upon treatment with various ER stressors and in different cell types

To determine whether RBM25 dependent regulation of XBP1s mRNA represents a general feature of the UPR rather than a tunicamycin specific effect, we examined XBP1s mRNA induction in response to pharmacologically distinct ER stressors. HeLa S3 cells were exposed to thapsigargin (Tg), an inhibitor of the sarco/endoplasmic reticulum Ca²⁺ ATPase (SERCA); dithiothreitol (DTT), which disrupts disulfide bond formation; or brefeldin A (BFA), an inhibitor of ER–Golgi transport. In all cases, depletion of RBM25 resulted in a marked reduction of *XBP1s mRNA* expression compared with control cells (**Figure 5A**), independently of the cause of ER stress.

**Figure 5.**
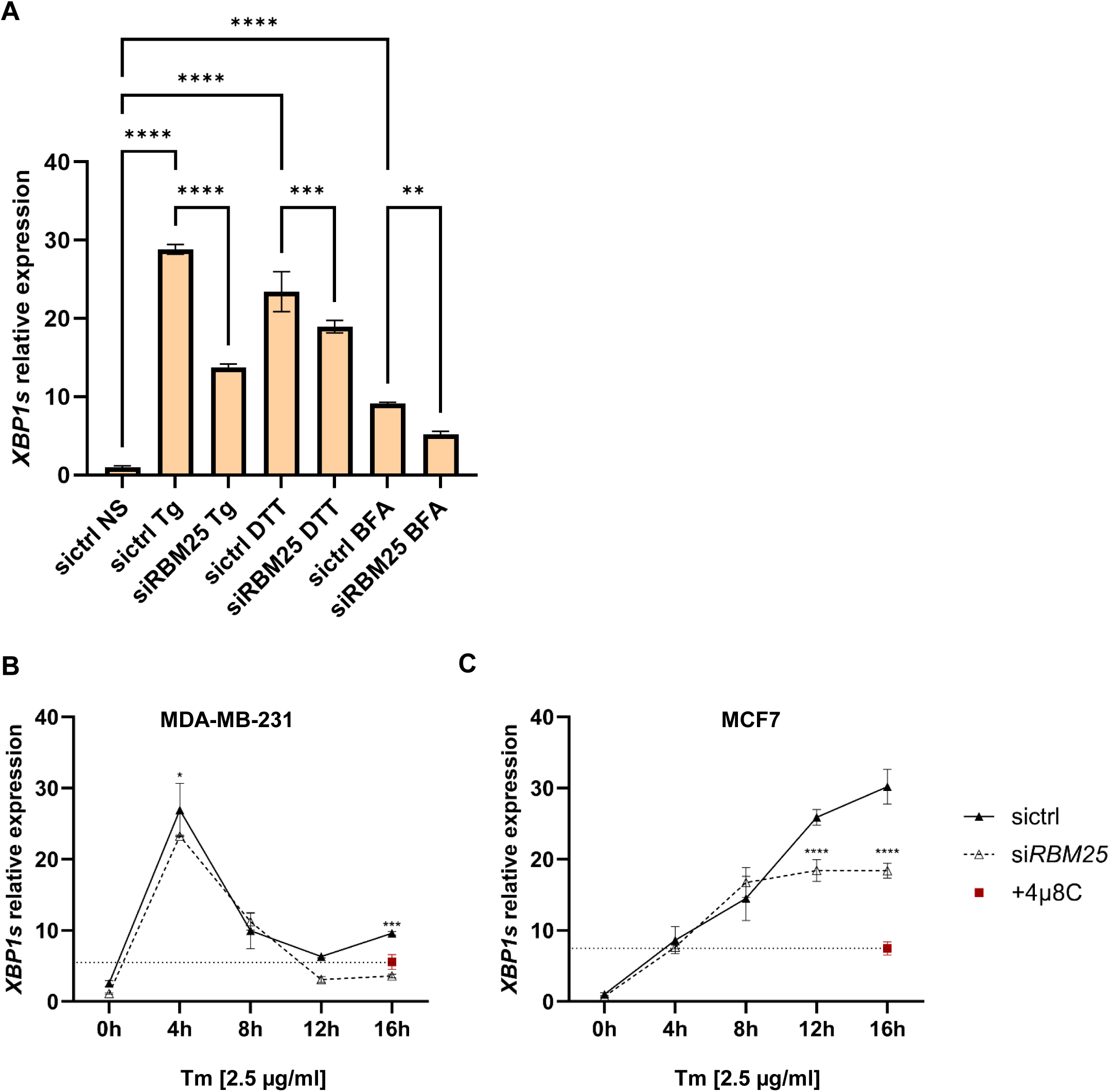
RBM25 is required for robust *XBP1s* induction across ER stress modalities and cell types. **A.** Quantitative RT-PCR analysis of *XBP1s* expression in HeLa S3 cells transfected with control siRNA (sictrl) or RBM25-targeting siRNA (siRBM25) following treatment with distinct ER stressors. Thapsigargin (Tg), dithiothreitol (DTT), and brefeldin A (BFA) each induced *XBP1s* expression in sictrl cells, whereas RBM25 depletion significantly attenuated this response across all conditions. **B**.**C**. Quantitative RT-PCR analysis of *XBP1s* expression in breast cancer cell lines with distinct basal IRE1-XBP1 activity. Under non-stressed (NS) conditions, basal XBP1s levels were low and unaffected by RBM25 depletion in both MDA-MB-231 and MCF7 cells. Tunicamycin (Tm) treatment induced a time-dependent increase in XBP1s expression in sictrl cells,. RBM25 knockdown significantly reduced XBP1s induction in both cell lines, with a stronger effect in MDA-MB-231 cells. Concurrent treatment with the IRE1 RNase inhibitor 4μ8C suppressed Tm-induced XBP1s expression to near-basal levels in both models. Transcript levels were normalized to *RPLP0* and expressed relative to unstressed controls. Data represent mean ± s.e.m. from three independent experiments. Statistical significance was determined using two-tailed tests (*P < 0.05, **P < 0.01, ***P < 0.001, ****P < 0.0001, n=3).

We next asked whether this regulatory requirement is preserved in cancer cells with distinct basal levels of IRE1-XBP1 signaling. In triple negative breast cancer MDA MB 231 cells, which exhibit constitutively elevated IRE1 RNase activity (21,22), *RBM25* silencing strongly attenuated *XBP1s* expression after 8 h of tunicamycin treatments, to an extent comparable to that observed upon pharmacological inhibition of IRE1. In contrast, in MCF7 breast cancer cells characterized by lower basal IRE1 activity (22,23), *RBM25* depletion during ER stress produced a significant but less pronounced reduction in *XBP1s* levels (**Figure 5BC**). Notably, in both cell lines, *RBM25* silencing impaired sustaine*d XBP1s* accumulation during prolonged UPR, limiting the attainment of maximal *XBP1s* levels following the initial response to ER stress. These findings support a role for RBM25 in stabilizing *XBP1s* mRNA rather than in regulating its initial induction. Collectively, these results indicate that RBM25 function scales with IRE1-XBP1 pathway output and is particularly critical in cellular contexts with high *XBP1s* demand. Furthermore, our data demonstrate that RBM25 acts as a broadly operating determinant of *XBP1s* mRNA accumulation, required for efficient activation of the IRE1 branch of the UPR across diverse ER stress conditions and cellular backgrounds.

### RBM25 is required for full activation of XBP1s-dependent UPR signaling

To assess the downstream consequences of RBM25 dependent regulation of *XBP1s* mRNA expression upon ER stress, we examined the expression of canonical XBP1s transcriptional targets, including the ER associated degradation component *EDEM1* and the chaperone *DNAJB9* (19,24,25). Tunicamycin treatment robustly induced expression of both mRNA in HeLa S3 cells, whereas this induction was significantly attenuated upon RBM25 silencing (**Figure 6AB**). Notably, silencing of *RBM25* did not impact *EDEM1* or *DNAJB9* expression under basal conditions, indicating that RBM25 is specifically required for stress induced UPR transcriptional output.

**Figure 6.**
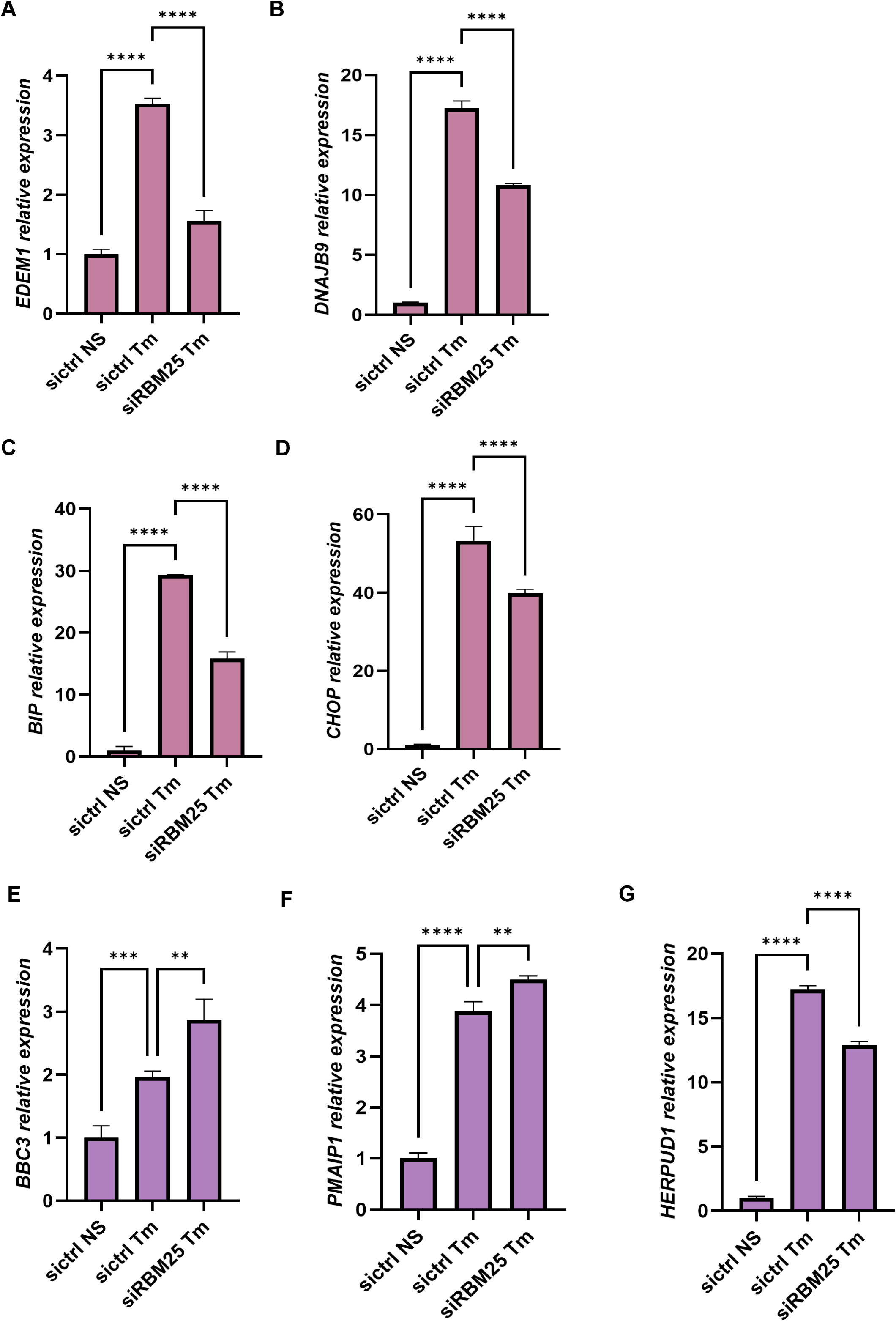
RBM25 depletion attenuates canonical UPR transcriptional output downstream of *XBP1s.* Quantitative RT-PCR analysis of XBP1s target genes in HeLa S3 cells transfected with control siRNA (sictrl) or *RBM25*⍰targeting siRNA (siRBM25). Under non⍰stressed (NS) conditions, basal expression of *EDEM1* (**A**) and *DNAJB9* (**B**) was low and unaffected by *RBM25* depletion. Tunicamycin (Tm) treatment robustly induced both transcripts in sictrl cells, whereas this induction was significantly attenuated upon *RBM25* knockdown indicating impaired activation of XBP1s dependent transcriptional programs. Tm treatment induced *BiP* (**C**) and *CHOP* (**D**) expression in control cells, consistent with activation of UPR signaling. RBM25 depletion reduced *BiP* induction by approximately 50% and resulted in a more modest decrease in *CHOP* expression, reflecting partial attenuation of downstream UPR signaling. To assess pathway specificity, expression of representative ATF4⍰and ATF6⍰dependent target genes was evaluated. RBM25 depletion led to a modest increase in the ATF4 targets *BBC3* (*PUMA*) (**E**) and *PMAIP1* (*NOXA*) (**F**), whereas the ATF6 target *HERPUD1* (**G**) was only slightly reduced. These results indicate that RBM25 depletion does not impair ATF4⍰or ATF6⍰dependent transcriptional programs but selectively attenuates the IRE1-XBP1 branch of the UPR. Transcript levels were normalized to *RPLP0* and expressed relative to unstressed controls. Data represent mean ± s.e.m. from three independent experiments. Statistical significance was determined using two⍰tailed tests (*P < 0.05, **P < 0.01, ***P < 0.001, ****P < 0.0001, n=3).

We next examined the expression of *BiP* and *CHOP*, key UPR effectors that are regulated by multiple signaling branches, including ATF6 and ATF4 in addition to *XBP1s* (26–28). Upon ER stress, *BiP* induction was reduced by approximately 50% following *RBM25* knockdown, whereas *CHOP* mRNA levels were also significantly decreased, albeit to a lesser extent (∼20%; **Figure 6CD**). Given that *BiP* and *CHOP* are not exclusive XBP1s targets, their partial sensitivity to *RBM25* depletion likely reflects compensatory regulation by parallel ATF6- and ATF4-dependent pathways.

To directly assess the specificity of RBM25 within the UPR network, we evaluated the expression of representative ATF4- and ATF6-dependent target genes. Expression of the ATF4 targets *PUMA* (*BBC3*) and *NOXA* (*PMAIP1*) (29) was not reduced but modestly increased in RBM25-depleted cells exposed to ER stress, whereas the ATF6 target *HERPUD1* (*30*) was only slightly decreased (**Figure 6E–G**). The modest upregulation of the pro-apoptotic factors *PUMA* and *NOXA* (31) is consistent with impaired adaptive capacity under conditions of reduced XBP1s levels. In contrast, the limited effect on *HERPUD1*, which is primarily regulated by ATF6 but also responsive to XBP1s (30), likely reflects partial dependence on XBP1s activity.

These results demonstrate that RBM25 silencing does not impair ATF4- or ATF6-dependent transcriptional programs. Collectively, these data indicate that RBM25 selectively supports the IRE1–XBP1 branch of the UPR by sustaining XBP1s-dependent transcriptional activity, while leaving parallel ATF4- and ATF6-mediated responses largely intact. Thus, RBM25 is required for maximal XBP1s-dependent activation of adaptive UPR signaling. Accordingly, loss of RBM25 compromises the adaptive UPR.

## DISCUSSION

The unfolded protein response requires finely tuned signaling outputs to restore endoplasmic reticulum (ER) proteostasis while avoiding premature engagement of apoptotic programs. Central to this balance is the IRE1–XBP1 pathway, which orchestrates adaptive transcriptional remodeling through generation of the spliced transcription factor XBP1s. While regulatory mechanisms controlling IRE1 activation, RNase activity, and *XBP1* mRNA splicing have been extensively studied, far less is known about how XBP1s abundance is quantitatively maintained once splicing is achieved. Our study identifies RBM25 as a previously unrecognized post-transcriptional determinant of *XBP1s* mRNA levels and reveals an additional RNA-centric checkpoint that shapes UPR signal strength downstream of IRE1. Our study identifies RBM25 as a previously unrecognized stabilizer of *XBP1s* mRNA, thereby sustaining XBP1s protein expression and transcriptional activity, and reveals an RNA-centric checkpoint that defines UPR signal strength downstream of IRE1.

We show that RBM25 selectively stabilizes the spliced form of *XBP1* mRNA through an interaction with its 3’ untranslated region, thereby sustaining XBP1s accumulation upon ER stress. This regulation operates independently of IRE1 RNase activity and does not impact RIDD activity, positioning RBM25 as an UPR modulator that acts in parallel to canonical enzymatic control by IRE1. Importantly, RBM25 does not affect the unspliced *XBP1u* transcript, highlighting that post-splicing mRNA fate, rather than splicing per se, is a critical determinant of the adaptive UPR output. These findings expand current models of UPR regulation by demonstrating that mRNA stability constitutes a key checkpoint defining the magnitude of XBP1-dependent transcriptional responses.

The role of RBM25 is observed under treatments with mechanistically distinct ER stressors and scales with pathway demand, being most pronounced in cellular contexts with high basal IRE1–XBP1 activity. This suggests that RBM25 functions as a quantitative amplifier that ensures sufficient availability of XBP1s under conditions of sustained ER stress, when transcription, translation, and RNA metabolism are globally constrained. By stabilizing *XBP1s* mRNA, RBM25 enables robust induction of downstream adaptive targets, including ER-associated degradation components and molecular chaperones, thereby supporting the proteostatic remodeling required for stress resolution.

Importantly, our data further reveal that RBM25 does not broadly modulate UPR signaling but instead exerts branch-specific effects. While *RBM25* depletion significantly attenuated the expression of canonical XBP1s target genes, transcriptional outputs associated with the ATF4 and ATF6 pathways remained largely preserved. The limited impact on representative ATF4 targets (*PUMA*, *NOXA*) and the modest effect on the ATF6-responsive gene *HERPUD1* indicate that RBM25 selectively reinforces the IRE1–XBP1 transcriptional axis rather than globally amplifying UPR signaling. This branch specificity distinguishes RBM25 from upstream regulators of ER stress responses and suggests that modulation of mRNA stability can provide pathway-selective control of UPR output. RBM25-dependent control of XBP1s also influences transcription of UPR cell fate decisions-related genes during ER stress, as attenuation of *XBP1s* stabilization upon RBM25 depletion blunts induction of adaptive UPR genes. This effect is most evident under conditions of acute ER stress that rely heavily on XBP1s signaling for survival, consistent with a model in which RBM25 primarily supports the early, adaptive phase of the UPR. The more modest impact on *CHOP* expression at early time points further suggests that RBM25 does not directly promote pro-apoptotic signaling but instead contributes to establishing an adaptive threshold that delays or prevents transition to terminal UPR outcomes.

RBM25 has previously been characterized as a nuclear RNA-binding protein regulating alternative pre-mRNA splicing at early stages of spliceosome assembly (17,32), with well-documented roles in controlling isoform decisions in apoptosis-related transcripts such as *BCL2L1* (BCL-x) (18) and in pathological splicing of *SCN5A* in heart failure (33). Our findings extend this functional repertoire by revealing a distinct role for RBM25 in the regulation of a fully processed mRNA. Unlike its canonical co-transcriptional splicing function, RBM25 directly binds the 3′UTR of spliced *XBP1* mRNA and selectively stabilizes the XBP1s transcript during ER stress. This mechanism identifies mRNA stability as an additional post-transcriptional checkpoint shaping IRE1–XBP1 signaling output.

Conceptually, this role parallels observations in *Drosophila*, where the RBP Pumilio promotes Xbp1 function by limiting RIDD-mediated mRNA decay (13). In mammalian cells, however, *XBP1s* is not itself a RIDD substrate (6–8). Instead, our data indicate that RBM25 ensures adequate accumulation of XBP1s in a cellular environment characterized by heightened RNA turnover and limited translational capacity during ER stress. Thus, rather than shielding XBP1s from IRE1-mediated degradation, RBM25 functions as a quantitative stabilizer that prolongs *XBP1s* mRNA persistence and enables effective transcriptional activation of adaptive UPR programs.

At first glance, the pro survival role of RBM25 identified here may appear inconsistent with prior reports linking elevated RBM25 levels to apoptosis in cardiac stress, where RBM25 has been implicated in CHOP dependent cell death (34). We propose that these seemingly divergent observations reflect context and time dependent roles of RBM25 within the UPR. Whereas our experiments capture early ER stress responses dominated by XBP1s mediated adaptation, prolonged or unresolved ER stress is known to shift UPR output toward CHOP driven apoptosis. Within this framework, RBM25 may support adaptive UPR signaling during early stress phases, while contributing to apoptotic outcomes under sustained stress or in specialized cell types such as cardiomyocytes. Consistent with this interpretation, elevated *RBM25* expression across multiple tumor types **(Supplementary Fig. 5**) suggests that RBM25 may support persistent XBP1s activity in malignancies characterized by chronic ER stress. Notably, RBM25 abundance, phosphorylation, and subcellular localization remained largely unchanged during ER stress **(Supplementary Fig. 2, 3 and 4)**, indicating that its regulatory function is mediated primarily through RNA interaction rather than stress dependent protein remodeling.

These findings also refine our understanding of how UPR signal strength and specificity are achieved at the post-transcriptional level. In contrast to mechanisms that regulate IRE1 activation or RNase activity, RBM25 operates downstream of splicing to control the persistence of the *XBP1s* transcript, thereby determining the magnitude and duration of its transcriptional output. By selectively stabilizing *XBP1s* mRNA without affecting parallel ATF4- or ATF6-driven programs, RBM25 acts as a quantitative amplifier that biases UPR signaling toward the adaptive IRE1-XBP1 branch. This selective reinforcement likely contributes to maintaining proteostatic capacity during early or moderate ER stress, while allowing other branches to remain responsive to prolonged or unresolved stress conditions. Such branch-selective tuning of UPR output may be particularly relevant in pathological contexts, such as cancer, where sustained activation of XBP1s supports proliferation and stress adaptation, and where uncoupling IRE1-XBP1 signaling from pro-apoptotic pathways could confer a selective survival advantage.

Collectively, our data support a revised model of the IRE1-XBP1 branch of the UPR (**Figure 7**), in which IRE1 mediated splicing generates XBP1s, while RBM25 ensures stability and persistence of the spliced transcript via its 3′UTR, together defining the amplitude and duration of adaptive UPR signaling. This RNA centric regulatory layer adds flexibility and tunability to UPR output and may be particularly relevant in pathological contexts such as cancer, where sustained XBP1s activity promotes survival, metabolic adaptation and therapy resistance.

**Figure 7.**
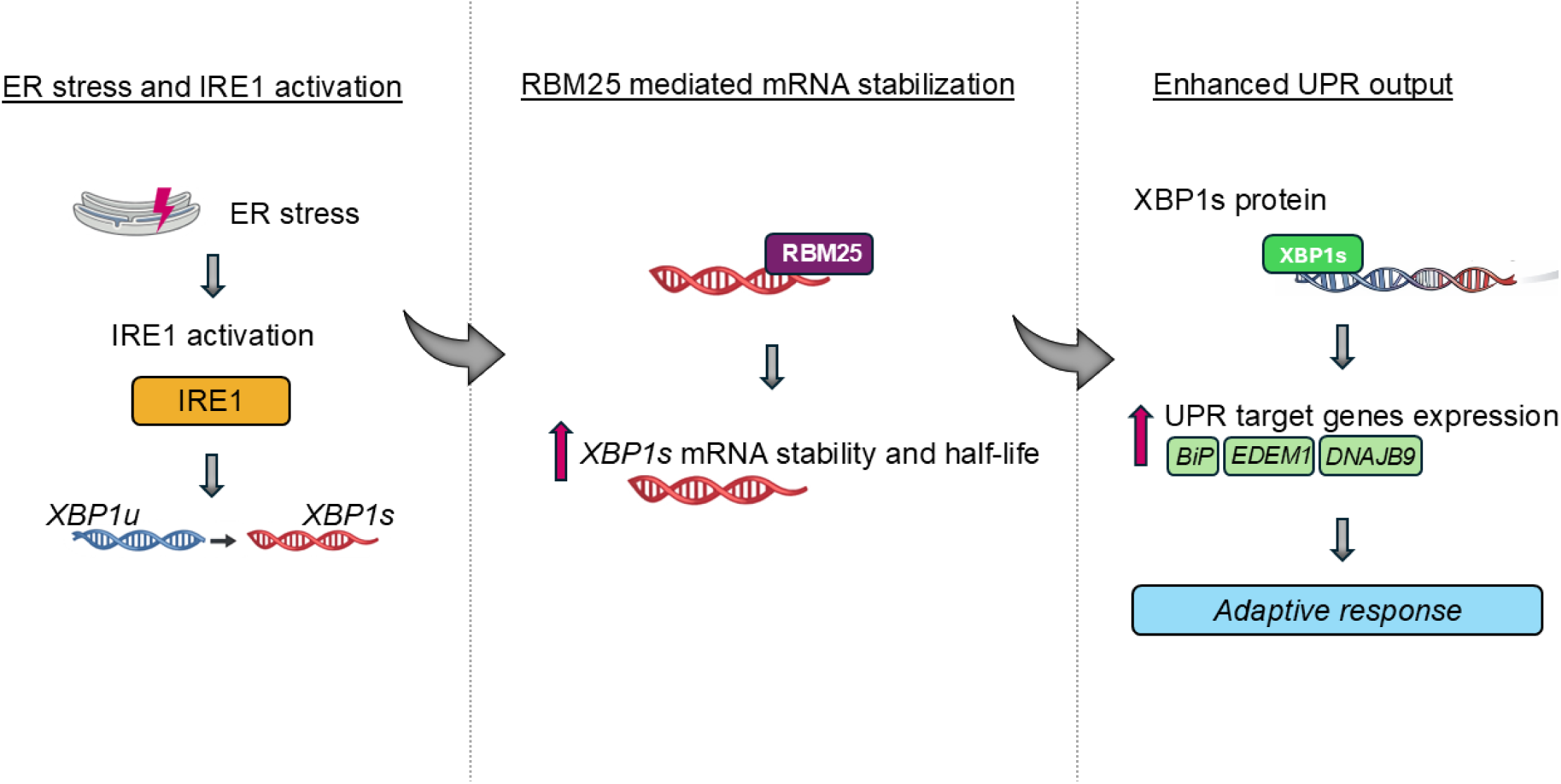
RBM25 stabilizes XBP1s mRNA to control IRE1-XBP1 pathway output. RBM25 binds the *XBP1s* 3′UTR and increases mRNA stability, enhancing XBP1s accumulation and downstream UPR gene expression, thereby acting as a post⍰transcriptional amplifier of adaptive ER stress responses.

Despite identifying RBM25 as a key post transcriptional regulator of XBP1s, the precise molecular mechanism underlying this stabilization remains to be defined. RBM25 may protect XBP1s from exonucleolytic decay, alter the composition or dynamics of XBP1 containing ribonucleoprotein complexes, or influence transcript localization during ER stress. Whether RBM25 acts autonomously or cooperates with additional RNA binding proteins, RNA modifications or stress induced remodeling of RNA-protein assemblies also remains unknown. Addressing these questions will be essential for understanding how RNA centric regulatory layers integrate with canonical UPR signaling and for evaluating whether RBM25 mediated control of XBP1s stability can be selectively manipulated to recalibrate maladaptive UPR signaling in disease settings such as cancer or cardiovascular disorders, without directly inhibiting IRE1 activity.

## METHODS

### Templates preparation

In order to create biotinylated RNA baits, based on (14) vectors encoding 3’UTR sequence of *XBP1s* (nucleotides 1180 -1776) or *XBP1u* (nucleotides 835 -1802), without poly(A) signal sequence, with the promoter sequence for polymerase T7 Φ2.5 at 5’ end were prepared. The templates for cloning were as follows: HmiT018550-MT06 for XBP1s and HmiT018551-MT06 for XBP1u (GeneCopoeia, Rockville, MD, USA). 20 ng of 3’UTR sequences templates underwent PCR reactions in the presence of 0.02 U/μL Phusion™ High -Fidelity DNA Polymerase (added after initial denaturation step; Thermo Scientific, F530S), 200 µM of each dNTP (Thermo Scientific, R0242), 1X Phusion™ HF Buffer and 0,5 µM of the appropriate forward and reverse primer (Table 1.). PCR reaction programs are presented in Table 2.

**Table 1.**
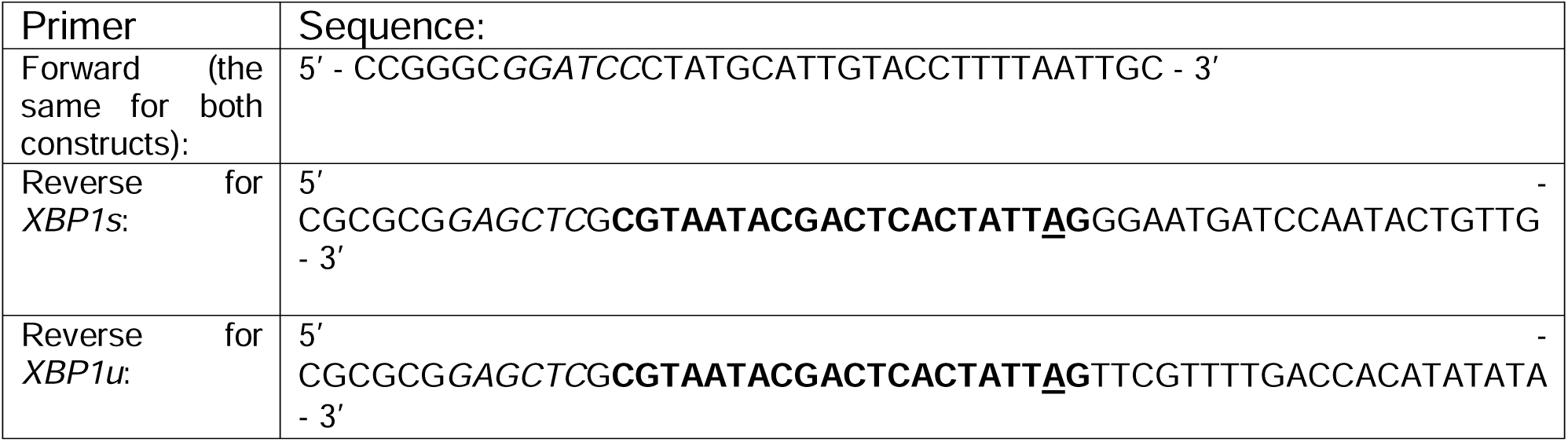
Primers used in the PCR reaction to amplify DNA sequences of 3’UTR of *XBP1s* and *XBP1u*. In bold: promoter sequence for polymerase T7 Φ2.5, **<u>A</u>** -is the nucleotide that during *in vitro* transcription biotin-AMP is inserted, in italics: restriction site for *BamHI* in forward primer or for *SacI* in reverse primers, that enabled cloning into pUC18 plasmid vector. All primers were synthesized by Genomed, Warsaw, Poland.

**Table 2.** PCR reaction programs for 3’UTRs amplification.

|  | **3’UTR of XBP1s** | **3’UTR of XBP1u** | **Step** | **No. of cycles** |
|---|---|---|---|---|
| temperature [°C] | 98 | 98 | initial denaturation | - |
| time [s] | 180 | 180 |  |  |
| pause |  |  | -polymerase addition |  |
| temperature [°C] | 98 | 98 | denaturation | X5 |
| time [s] | 10 | 10 |  |  |
| temperature [°C] | 57 | 55 | annealing |  |
| time [s] | 30 | 30 |  |  |
| temperature [°C] | 72 | 72 | extension |  |
| time [s] | 10 | 15 |  |  |
| temperature [°C] | 98 | 98 | denaturation | X20 |
| time [s] | 10 | 10 |  |  |
| temperature [°C] | 72 | 72 | annealing+ |  |
| time [s] | 40 | 45 | elongation |  |
| temperature [°C] | 72 | 72 | final extension | - |
| time [s] | 300 | 300 |  |  |
| temperature [°C] | 4 | 4 | hold |  |
| time [s] | ∞ | ∞ |  |  |

PCR products were purified with NucleoSpin Columns (MACHEREY-NAGEL GmbH & Co. KG, 740609.50S), digested with *BamHI* (Thermo Scientific, FD0054) and *SacI* (Thermo Scientific, FD1134) restriction enzymes, according to enzyme manufacturer’s recommendations, purified again with NucleoSpin Columns and ligated to of linearized with *BamHI/SacI* pUC18 vector in the molar ratio of 1.5:1. Ligation with T4 DNA Ligase (Thermo Scientific, EL0014) was conducted according to manufacturer’s instructions and *E. coli* DH5α competent cells were transformed according to protocol described in (35). Cells were plated on Luria Broth (LB; Invitrogen, 11548866) -agar plates with ampicillin (BioShop Canada Inc., AMP201.100) at a concentration of 100 μg/ml. The plates were incubated overnight at 37°C.

Bacterial minicultures were established for each construct. Colonies were placed in 3 ml of LB medium with 100 μg/ml ampicillin and incubated overnight at 37°C and 182 rpm. Plasmids were isolated with GeneMATRIX Basic DNA Purification Kit (*EURx* Sp. z o.o., E3545). The sequencing results performed at Genomed (Warsaw, Poland) confirmed the correctness of performed cloning.

| **Molecular cloning** |  |  |
|---|---|---|
| ampicillin | BioShop Canada Inc. | AMP201.100 |
| *BamHI* | Thermo Scientific | FD0054 |
| dNTP | Thermo Scientific | R0242 |
| GeneMATRIX Basic DNA Purification Kit | *EURx* Sp. z o.o. | E3545 |
| Luria Broth (LB) | Invitrogen | 11548866 |
| NucleoSpin Columns | MACHEREY-NAGEL GmbH & Co. KG | 740609.50S |
| Phusion™ High - Fidelity DNA Polymerase | Thermo Scientific | F530S |
| *SacI* | Thermo Scientific | FD1134 |
| T4 DNA Ligase | Thermo Scientific | EL0014 |
| XBP1s | GeneCopoeia | HmiT018550-MT06 |
| XBP1u | GeneCopoeia | HmiT018551-MT06 |

### *in vitro* Transcription

*XBP1s and XBP1u* transcripts were synthesized from *SacI* and *BamHI*-cut templates. Inserts encoding 3’UTRs of *XBP1s* or *XBP1u* were extracted from gel and purified with NucleoSpin Columns. *In vitro* transcription conditions were based on (14) with 50 U of RNA polymerase T7 Φ2.5 (Thermo Scientific, EP0112) for each 1 µg of template; choosing a fourfold excess of biotin -AMP (Jena Bioscience, NU-894-BIO-S) over ATP (Thermo Scientific, R0441) 2 mM vs 0.5 mM. Remaining nucleotides GTP, UTP, CTP (Thermo Scientific, R0461, R0471, R0451, respectively) were used in the final concentration of 1 mM. The reaction was carried at 37 °C, for 4 hours in the presence of 20 U RiboLock RNase Inhibitor (Thermo Scientific, EO0381) and 5 mM DTT (Sigma, 43815), followed by 10 min of enzyme deactivation at 70 °C. The reaction products were purified using RNA Clean & Concentrator-25 (Zymo Research, R1017) according to the manufacturer’s recommendations. The concentration of the RNA were assessed using a Multiskan SkyHigh Microplate Spectrophotometer (Life Technologies, Warsaw, Poland) by absorbance at 260 nm (A_260nm_).

| ***in vitro* transcription assay** |  |  |
|---|---|---|
| ATP | Thermo Scientific | R0441 |
| Biotin-AMP | Jena Bioscience | NU-894-BIO-S |
| CTP | Thermo Scientific | R0451 |
| DTT | Sigma | 43815 |
| GTP | Thermo Scientific | R0461 |
| NucleoSpin Columns | MACHEREY-NAGEL GmbH & Co. KG | 740609.50S |
| RNA polymerase T7 Φ2.5 | Thermo Scientific | EP0112 |
| UTP | Thermo Scientific | R0471 |
| RiboLock RNase Inhibitor | Thermo Scientific | EO0381 |
| RNA Clean & Concentrator-25 columns | Zymo Research | R1017 |
| UTP | Thermo Scientific | R0471 |

### Cell lines

All cells were obtained from ATCC (CCL-2.2™), MDA-MB-231 (HTB-26™) and MCF7 (HTB-22™). Cells were cultured in Minimum Essential Modified Eagle’s Medium with 2mM l-glutamine (MEM; Biowest, L0416), antibiotics (100 units/mL of penicillin, 100 µg/mL of streptomycin (Biowest, L0018)) and 10% fetal bovine serum (FBS, Sigma-Aldrich, F7524). The inducible stable HeLa S3 XBP1s cell line containing the cDNA sequence of XBP1s (NM_001079539.1) was obtained as described previously [3]. This cell line was cultured in MEM with 2 mM glutamine, hygromycin B (300 µg/ml) and 10% tetracycline free FBS (Takara Bio, USA, 631101). For the XBP1s 24 h induction, doxycycline (DOX; 400 µg/ml, Millipore-Sigma, D3072) was added to culture media for 24 h. All cell lines were cultured in a humidified incubator at 37°C in 5% CO2 in 100 mm dishes or 6-well plates and allowed to grow to 70 -80% confluence prior to the start of the experiments.

| **Cell culture** |  |  |
|---|---|---|
| doxycycline | Millipore-Sigma | D3072 |
| fetal bovine serum, FBS | Sigma-Aldrich | F7524 |
| hygromycin B | Invitrogen | 10687010 |
| Minimum Essential Modified Eagle’s | Biowest | L0416 |
| penicillin and streptomycin solution | Biowest | L0018 |
| tetracycline free FBS | Takara Bio | 631101 |

### Pharmacological induction of ER stress

Pharmacological induction of ER stress and activation of the UPR were performed according to previously described methods(36). Briefly, cells were treated with the compounds for the time periods specified: Tm (2.5 µg/ml; Sigma, T7765), Tg (50 nM; Sigma, T9033), BFA (0.4 µg/ml, Sigma, B6542), or DTT (4 mM; Sigma, 43815) for the time periods specified. Ctrl cells were treated with vehicle control (ctrl), DMSO (Sigma, D2650). To verify IRE1 activity, cells were treated with 10 µM 4µ8C (Sigma-Aldrich, SML0949) dissolved in DMSO (5,15,37).

| **Pharmacological ER stress inducers and IRE1 inhibitor** |  |  |
|---|---|---|
| 4µ8C | Sigma-Aldrich | SML0949 |
| brefeldin A, BFA | Sigma | B6542 |
| dimethyl sulfoxide, DMSO | Sigma | D2650 |
| dithiothreitol, DTT | Sigma | 43815 |
| thapsigargin, Tg | Sigma | T9033 |
| tunicamycin, Tm | Sigma | T7765 |

### siRNA transfections

siRNA for RBM25 (Ambion assay id s33912) and YBX1 (Ambion assay id s224139) were purchased from Thermo Scientific. Cells were transfected using the Lipofectamine RNAiMax (Invitrogen, 13778030) according to manufacturer’s protocol. The siRNAs were used at a final concentration of 20 nM for single knockdown. The transfected cells were cultured for 2 days prior to further analysis. Silencer Select Negative Control Ambion No. 1 (Invitrogen, 4404021) was used as a control.

| **siRNA transfections** |  |  |
|---|---|---|
| Lipofectamine RNAiMax | Invitrogen | 13778030 |
| sictrl | Invitrogen | 4404021 |
| siRBM25 | Ambion | s33912 |
| siYBX1 | Ambion | s224139 |

### Isolation of cytoplasmic proteins fraction and RBPs

HeLa S3 cells were cultured in 100-mm dishes (10 dishes per condition; no stress (NS) or Tm-treated). Cells were washed twice with ice-cold PBS (Biowest, L0616), scraped into 1□ml PBS per dish, and collected by centrifugation (5□min, 300□×□*g*, 4°C). After removing PBS, the packed cell volume (PCV) was determined. Cell pellets were resuspended in five volumes of lysis buffer (10□mM HEPES, 10□mM KCl, 1□mM MgCl₂, 1□mM DTT, and 1x Protease Inhibitor cOmplete Mini (Roche, 11836170001), pH 7.4) and incubated on ice for 20□min. Cells were lysed by passing the suspension 20 times through a syringe needle (G27). Lysates were clarified by ultracentrifugation (1□h, 100,000□×□*g*, 4°C), and protein concentration of the resulting cytoplasmic fraction was determined by A₂₈₀ absorbance.

For RNA pulldown assays, 1□µg of biotinylated RNA bait was incubated with 500□µg of cytoplasmic extract for 1□h at room temperature in the presence of 20□U RiboLock. Streptavidin magnetic beads (50□µl; Pierce, 88816) were added and incubated for an additional 1 h at room temperature with gentle agitation. Beads were collected using a magnetic rack and washed twice with wash buffer (0.6□M NaCl, 20□mM Tris, 10□mM EDTA, 1□mM DTT, 1× **protease inhibitor cocktail** (PIC), pH□7.4).

| **Proteins isolation, handling and Western blot utilities** |  |  |
|---|---|---|
| λ phosphatase | Biolabs | P0753S |
| 7.5% TGX<sup>TM</sup> FastCast<sup>TM</sup> polyacrylamide gel | Bio-Rad | 161-0171 |
| 4× Laemmli buffer | Bio-Rad | 1610747 |
| anti-β-actin | Cell Signaling Technology | #4967 |
| anti-rabbit IgG secondary antibody | Bio-Rad | 1706515 |
| anti-RBM25 antibody | Proteintech | 25297-1-AP |
| anti-XBP1s | Cell Signaling Technology | #12782 |
| Criterion TGX Stain-Free 4 - 15% gradient gels | Bio-Rad | 5678084 |
| DC Protein Assay | Bio-Rad | 5000111 |
| Dulbecco’s Phosphate Buffered Saline, PBS | Biowest | L0616 |
| Mini-PROTEAN TGX Stain-Free gels | Bio-Rad | 4568084 |
| IgG2a isotype control | MBL | M076-3 |
| ethylenediaminetetraacetic acid, EDTA | Sigma-Aldrich | E6758 |
| HEPES | Bioshop | HEP001.1 |
| magnesium chloride, MgCl<sub>2</sub> | Sigma | M8266 |
| Potassium chloride, KCl | Roth | P017.1 |
| Protease Inhibitor cOmplete Mini | Roche | 11836170001 |
| PVDF membrane | GE Healthcare | 10600023 |
| sodium chloride; NaCl | Sigma-Aldrich | S9888 |
| Streptavidin magnetic beads | Pierce | 88816 |
| SuperSep Phos-tag gels | FujiFilm | 192-17401 |
| SuperSignal West Femto substrate | Thermo Fisher Scientific | 34096 |
| Tris base | Millipore | 1.08387 |
| Tween-20 | IBI Scientific | IB07200 |

### Mass spectrometry analysis

Resin-bound proteins were denatured for 10 mins. in 65°C in presence of 0.5% sodium deoxycholate (DOC:Na, Merck). Next, 100 ng of Trypsin (Merck, EMS0006) was added to the samples for an overnight digestion in 37°C. On the following day, the peptide solution was separated from the resin, DOC:Na was removed via acidification and centrifugation, and the supernatants underwent desalting with the STAGE tip protocol (38). The obtained peptide pellet was resuspended in a 0.1% formic acid (FA) + 3% acetonitrile (ACN) solution.

Liquid Chromatography-Mass Spectrometry (LC-MS) was carried out on an M-Class Acquity nanoUPLC system coupled to a Synapt XS HDMS equipped with a nanoESI ion source interface. Mobile phase A consisted of H2O + 0.1% FA, while mobile phase B of ACN + 0.1% FA. Samples were separated on an analytical column (nanoEase HSS T3 C18 100Å, 1.8μm, 75μm x 150mm) kept at 40°C with a 40min linear gradient of 5-35%B at a 300nL/min flow rate. A 3min trapping step was performed prior to gradient separation. Injected volume was 3µL. MS data were collected in DIA mode, scanning at 2Hz through a 50-2000m/z range in ESI+ and analyzer mode set to Resolution (25000 FWHM at 785.84m/z). For MS2 of all ions, a collision energy ramp of 25-45V was set on instrument’s trap cell. Source conditions were fine-tuned and kept constant during acquisitions. A Glu-Fib (Waters) solution was acquired in the mass reference function, and the correction was applied post-acquisition.

Raw processing was performed using Progenesis QI for Proteomics v4.2.7 (Non-linear Dynamics). QI’’s autovalues were unchanged for the deconvolution, and the set low and high energy ion thresholds were 200 and 20, respectively. Deconvoluted data were searched via Ion Accounting against a redundant human protein sequence databank (UP000005640, 83526 sequences) appended with the CCP version of cRAP contaminant sequences (125 sequences, [2] 2] T. G. P. Machine. [Online]. Available: https://www.thegpm.org/crap/. [Accessed 2026].). The search parameters were as follows; peptide/fragment mass tolerance: 10/20 ppm; min. fragments/peptide: 3; min. fragments/protein: 7; min. peptides/protein: 1; max. protein mass: 1MDa; enzyme: trypsin; max. missed cleavages: 4; variable modification: oxidation of methionine; FDR: 4% (protein-level). Protein grouping was enabled, and relative quantitation was set according to non-conflicted peptides. Lists containing uniquely identified and quantified proteins from two independent experiments (SFxx1, SFxx2) were exported for further analysis.

### RNA Immunoprecipitation

RIP assays were performed using a miRNA RIP Kit (MBL, RN1005), adapted to assess RBM25 dependent RNA interactions in HeLa S3 cells under NS or Tm treatment (Tm, 8 h). For each condition, cells at ∼80% confluence were collected from four 100 mm dishes. Cells were harvested by scraping in ice cold PBS, pelleted, and lysed in mi Lysis Buffer (+) according to the manufacturer’s instructions.

For immunoprecipitation, 3 µg of anti RBM25 antibody (Proteintech, 25297 1 AP) or mouse IgG2a isotype control (MBL, M0763) were incubated with agarose beads (Thermo Scientific, 20421). Following immunoprecipitation and washing steps, RNA bound to antibody -bead complexes was extracted using the RNeasy Mini Kit (Qiagen, 74104). Briefly, 200 µl of RLT Lysis Buffer was added directly to washed beads, and RNA was eluted in 40 µl of nuclease free water. Input and post RIP fractions were evaluated by Western blot (WB) and qPCR as described in the manufacturer’s protocol.

| **RNA** |  |  |
|---|---|---|
| actinomycin D | Merck | A9415 |
| agarose beads | Thermo Scientific | 20421 |
| miRNA RIP Kit | MBL | RN1005 |
| RNeasy Mini Kit | Qiagen | 74104 |
| TaqMan™ RNA-to-CT™ 1-Step Kit | Applied Biosystems™ | 4392938 |

### Isolation of cellular RNA

Total cellular RNA was isolated using RNeasy Mini Kit (Qiagen, 74104), according to the manufacturer’s recommendations All RNA samples were depleted of genomic DNA according to manufacture instructions. RNA concentration was calculated based on the absorbance at 260 nm. RNA samples were stored at −20°C until use.

### Quantitative Real-Time PCR

The TaqMan™ RNA-to-CT™ 1-Step Kit (Applied Biosystems™, 4392938) was used as previously described [4], following the manufacturer’s protocol, including a 20-minute reverse-transcription step at 50°C. Relative mRNA levels were calculated using the 2⁻ΔΔCt method [7]. Superoxide dismutase 1 (SOD1; Hs00533490_m1) or TATA box binding protein (TBP; Hs00427620_m1) mRNAs served as endogenous controls. The stability of these reference genes in mRNA half-life experiments was validated using **ribosomal protein lateral**

### stalk subunit P0

(RPLP0; Hs00420895_g1), and GAPDH (Hs03929097_g1). Gene expression levels were quantified and analyzed in technical triplicates.

TaqMan probe assays used in this study included BIP (HSPA5; Hs00607129_gH), CHOP (Hs00358796_g1), DGAT2 (Hs01015913_m1), DNAJB9 (Hs01052402_m1), EDEM1 (Hs00976004_m1), RBM25 (Hs00402198_m1), XBP1s (Hs03929085_g1), XBP1u (Hs02856596_m1), and YBX1 (Hs00358903_g1). All probes were obtained from Life Technologies (Warsaw, Poland).

| **Gene expression assay** |  |  |
|---|---|---|
| *BIP (HSPA5)* | Thermo Fisher | Hs00607129_gH |
| *CHOP* |  | Hs00358796_g1 |
| *DGAT2* |  | Hs01015913_m1 |
| *DNAJB9* |  | Hs01052402_m1 |
| *EDEM1* |  | Hs00976004_m1 |
| *GAPDH* |  | Hs03929097_g1 |
| *RBM25* |  | Hs00402198_m1 |
| *RPLP0* |  | Hs00420895_g1 |
| *SOD1* |  | Hs00533490_m1 |
| *TBP* |  | Hs00427620_m1 |
| *XBP1s* |  | Hs03929085_g1 |
| *XBP1u* |  | Hs02856596_m1 |
| *YBX1* |  | Hs00358903_g1 |

### Measurement of RNA half-life

RNA half-lives of XBP1s, XBP1u, and GAPDH were measured as previously described [8, 9] with minor modifications. HeLa S3 cells transfected with siRBM25 or control siRNA were seeded in 35 mm dishes and grown to ∼80% confluence. Cells were exposed to 2.5□µg/ml Tm for 3□h prior to initiation of the assay. To simultaneously inhibit IRE1 RNase activity and block transcription, 10□µM 4µ8C and 5□µg/ml actinomycin D (Merck, A9415) were added at the start of the time course and maintained throughout the experiment. Total RNA was isolated at the indicated time points using the miRNeasy kit (Qiagen). RNA levels were quantified by TaqMan based real time PCR and normalized to endogenous *RPLP0* mRNA (amplified using a standard primer set). For each time point, measurements were obtained from three biological replicates across at least two independent experiments. Relative RNA abundance was expressed as the fold change compared with the initial time point (t□=□0). RNA decay kinetics were calculated from exponential regression using the equation C/C₀ = e⁻□□□, where C and C₀ represent RNA levels at times t and t₀, and kd is the decay constant, as described previously (39).

### Western Blots

Cells were lysed on ice for 15□min in RIPA buffer (Sigma-Aldrich, R0278) supplemented with cOmplete Mini protease inhibitor. Lysates were homogenized by passing the suspension 20 times through a 27 gauge needle. Insoluble debris was removed by centrifugation (15,000□×□g, 15□min, 4□°C). Protein concentrations were measured using the DC Protein Assay (Bio Rad, 5000111) with bovine serum albumin (BSA; Biowest, P6155-KG) as a standard.

Equal amounts of protein were mixed with 4× Laemmli buffer and heated for 5□min at 95□°C before separation on Criterion TGX Stain Free 4 -15% gradient gels (Bio Rad, 5678084) or Mini PROTEAN TGX Stain Free gels (Bio Rad, 4568084). Proteins were transferred to **polyvinylidene difluoride** (PVDF) membranes at 300□mA for 90□min at 4°C. Membranes were blocked overnight in TBS (20□mM Tris, 150□mM NaCl, pH□7.6) containing 5% BSA and 0.5% Tween 20 (IBI Scientific, IB07200). Primary antibodies were applied in TBST containing 3% BSA: XBP1s (Cell Signaling Technology, #12782, 1:1000), RBM25 (Proteintech, 25297 1 AP, 1:1000), and β actin (Cell Signaling Technology, #4967, 1:1000). Following washes in TBST, membranes were incubated with horseradish peroxidase -conjugated goat anti rabbit IgG secondary antibody (Bio Rad, 1706515). Signal detection was performed using SuperSignal West Femto substrate (Thermo Fisher Scientific, 34096), and chemiluminescence was recorded with a ChemiDoc XRS imaging system (Bio Rad). Band intensities were quantified using Image Lab software v4.1 (Bio Rad).

Samples were harvested in 1X NEBuffer for PMP (50 mM HEPES, 100 mM NaCl, 2 mM DTT, 0.01% Brij-35, pH 7.5; Biolabs, P0753S) with 1 mM MnCl₂ after 8 hours of Tm treatment or NS exposure, for either λ phosphatase treatment (+λ) or not (−λ). The soluble fractions underwent λ phosphatase treatment according to the manufacturer’s protocol. Samples for WB analysis were prepared as previously described, and equal amounts of protein were mixed with 2× Laemmli buffer. The mixture was then heated for 5 min at 95°C before separation on SuperSep Phos-tag gels (FujiFilm, 192-17401). After transferring to a PVDF membrane (GE Healthcare, 10600023) and blocking overnight in 5% BSA in TBST, anti-RBM25 antibody (Proteintech, 25297-1-AP) and anti-β-actin antibody (Cell Signaling Technology, #4967) were applied in 3% BSA in TBST (dilution 1:1000). WBs were developed as described above.

### Immunofluorescence microscopy

Immunofluorescence microscopy was performed as described previously. Briefly, cells were seeded onto 25-mm glass coverslips and cultured at 37°C for 48 h after siRNA transfection. After 48 h of culturing, cells were treated with ER stressors (Tm or Tg or vehicle (DMSO)) for 8 h before fixation with 4% paraformaldehyde (15 min at RT) in PBS and permeabilized with 0.1% Triton X-100 in PBS for 15 min at RT, washed 3 times for 2 min each with PBS, and then blocked with 2% BSA in PBS for 1h at RT. Cells were incubated with primary antibodies diluted (1:200) in blocking solution overnight at 4°C. Following 3 washing steps with PBS, the Alexa Fluor^TM^ 594 goat anti-rabbit IgG secondary antibody (A11037, Invitrogen) (1:1000) in blocking solution was applied and incubated 1h in the dark. The coverslips were stained in Hoechst solution in PBS (1:1000) for 20 min at RT and mounted directly on a supporting glass slide Imaging was performed using the Stellaris (Leica) platform with HC PL APO CS2 63x/1.40 oil objective. For excitation, a 405 nm diode or a white pulse laser were used, depending on the dye. Precise excitation wavelength was given in the image description. The collected data were processed and analyzed with LasX software (Leica, Wetzlar, Germany) (40) and Fiji software (41).

### Quantification and statistical analysis

All data are expressed as means ± SEM. For comparisons involving more than two groups, a one-way analysis of variance (ANOVA) was conducted, followed by Bonferroni’s post hoc test for multiple comparisons. Differences were considered significant at *p* < 0.05. All analyses were performed using GraphPad Prism version 10,6,0 (GraphPad Software, San Diego, CA). If representative results are presented, remaining results are available as datasets.

## Supporting information

Suppmental Materials

Data set 1

Data set 2

## DECLARATIONS

### Materials availability

Further information and requests for resources and reagents may be directed to and will be fulfilled by the corresponding authors.

### Data and code availability

All data reported in this paper will be shared by the lead contact upon request. This paper does not report original code. Any additional information required to reanalyze the data reported in this paper is available from the lead contact upon request. Mass spectrometry data were deposited to Zenodo: DOI 10.5281/zenodo.20667311

## Acknowledgments

This research was funded by This work was funded by National Science Center under OPUS 2022/47/B/NZ7/00051 contract to R.B. The purchase of M-Class Acquity-Synapt XS LC-MS system was financially supported by the “Excellence Initiative -Research University” program for the University of Wroclaw.

## Author contributions

Conceived and designed the experiments: R.B., A.W.W. Performed the experiments: A.B., A.W.W. Analyzed the data: R.B., A.W.W., P.C., S.K., A.B.; Participation in CLSM imaging, preparation of Supplementary Figure 1: J.G. Performed the mass spectrometry experiments: M.T.; Contributed reagents/materials/analysis tools: R.B. Wrote the manuscript: R.B. A.W.W. Revised the manuscript: J.F.C. and E.C.

## Declaration of generative AI and AI-assisted technologies

Generative AI language tools (Microsoft Copilot) were used to assist in correcting language and grammar in this document. No content was generated beyond linguistic refinement.

**Supplementary Figure 1.** R**B**M25**, but not YBX1, is required for efficient XBP1s stabilization during ER stress.** Immunoblot analysis of XBP1s protein following ER stress. Under NS conditions, XBP1s protein was minimally detectable across all siRNA treatments. Tm exposure induced marked accumulation of XBP1s protein at 4 and 8 h in sictrl and siYBX1 cells, whereas RBM25 depletion substantially impaired XBP1s protein accumulation despite comparable β actin loading. These results indicate that RBM25, but not YBX1, is required for efficient stabilization and accumulation of XBP1s during ER stress, without affecting XBP1u expression.

**Supplementary Figure 2. RBM25 protein abundance is stable during ER stress and efficiently reduced upon siRNA-mediated depletion. I**mmunoblot analysis of HeLa S3 cells transfected with control siRNA (sictrl) or RBM25-targeting siRNA (siRBM25) under non-stressed (NS) and tunicamycin (Tm)-treated conditions. RBM25 protein levels remained stable upon Tm treatment in sictrl cells, indicating that ER stress does not alter RBM25 abundance. In contrast, RBM25 levels were markedly reduced in siRBM25 cells under both basal and stressed conditions, confirming efficient and sustained knockdown. β-actin serves as a loading control. Total protein staining of SDS-PAGE gels and membranes confirmed equal sample loading and transfer across all conditions.

**Supplementary Figure 3. RBM25 phosphorylation is maintained during ER stress independently of IRE1 activity.** RBM25 mobility was assessed using Phos⍰tag electrophoresis in HeLa S3 cells under non⍰stressed (NS) and tunicamycin (Tm)⍰treated conditions. RBM25 resolved as two species: a faster⍰migrating non phosphorylated form (∼100 kDa) and a slower⍰migrating phosphorylated form. Treatment with λ phosphatase (+λ) shifted RBM25 mobility toward the faster⍰migrating species, confirming basal phosphorylation. Tm treatment did not markedly alter the relative abundance of the phosphorylated RBM25 species, indicating that RBM25 phosphorylation is preserved during ER stress. Consistent with conventional SDS-PAGE analysis, total RBM25 protein levels remained stable under both NS and Tm conditions. RBM25 knockdown (siRBM25) substantially reduced both total and phosphorylated RBM25 species, confirming efficient depletion. Co treatment with the IRE1 RNase inhibitor 4μ8C during Tm exposure did not affect the abundance of the phosphorylated RBM25 species, indicating that RBM25 phosphorylation occurs independently of IRE1 activity. β⍰actin serves as a loading control and showed uniform expression across all samples.

**Supplementary Figure 4. RBM25 nuclear localization is maintained during ER stress.** Confocal laser scanning microscopy (CLSM) images of HeLa S3 cells transfected with control siRNA (sictrl) or RBM25**⍰**targeting siRNA (siRBM25) under non⍰stressed (NS) conditions or following treatment with tunicamycin (Tm) or thapsigargin (Tg). RBM25 was detected by immunofluorescence (red), nuclei were visualized using Hoechst staining (blue), and bright field images are shown in grayscale. Merged images are displayed as overlays. Under NS conditions, sictrl cells exhibited clear nuclear localization of RBM25, whereas siRBM25 cells showed minimal signal, confirming efficient depletion. Induction of ER stress with Tm or Tg did not alter RBM25 localization in sictrl cells, with the protein remaining predominantly nuclear. In siRBM25 cells, weak diffuse staining was observed under stress conditions, consistent with residual signal. Bright⍰field images confirmed comparable cell morphology across all conditions. Images were adjusted for contrast only; no additional processing was applied. Scale bar, 50 μm. Imaging was performed using a HC PL APO CS2 63×/1.40 oil immersion objective.

**Supplementary Figure 5. R*B*M25 and *XBP1* expression patterns and correlation across human tumor types. A.** Differential expression analysis of *RBM25* mRNA in human tumor tissues (red) versus corresponding normal tissues (green), based on GEPIA database analysis (42). Only tumor types exhibiting statistically significant differences are shown (*P < 0.01). Sample numbers for each tissue type are indicated in parentheses. Expression levels are presented as transcripts per million (TPM). **B.** Differential expression analysis of *XBP1* mRNA across the same tumor types. *XBP1* expression is shown for tumor (red) and normal (green) tissues, limited to those cancer types displaying significant differential expression of *RBM25* (*P < 0.01). Sample sizes are indicated in parentheses. Expression values are presented as TPM. **C.** Pearson correlation analysis of *RBM25* and *XBP1* expression across multiple human tumor types. Green bars indicate tumor types with statistically significant correlations (*P < 0.05), whereas grey bars represent non⍰significant correlations.

Correlation analysis between *XBP1* and *RBM25* expression levels was performed using the GEPIA (Gene Expression Profiling Interactive Analysis) platform. Pearson correlation coefficients (*R*) and their corresponding *p*-values were calculated based on datasets from TCGA (The Cancer Genome Atlas). The results were visualized as a bar chart generated using Python 3 with the Matplotlib library (43).

**Supplementary Figure 6. Raw WB data.**

**Data Set 1. Mass Spectrometry analysis.**

**Data Set 2. mRNA half-life analysis.**

## Abbreviations

3’ UTR: 3’ untranslated region
4μ8C: IRE1 RNase inhibitor
ACN: acetonitrile
ATF4: activating transcription factor 4
ATF6: activating transcription factor 6
BFA: brefeldin A
BiP: Heat Shock Protein Family A (Hsp70) Member 5 (HSPA5)
BSA: bovine serum albumin
CHOP: DNA Damage Inducible Transcript 3
CLSM: confocal laser scanning microscopy
DGAT2: diacylglycerol O-acyltransferase 2
DNAJB9: DnaJ Heat Shock Protein Family (Hsp40) Member B9
DOX: doxycycline
DOC: sodium deoxycholate
DTT: dithiothreitol
EDEM1: ER Degradation Enhancing Alpha-Mannosidase Like Protein 1
ER: endoplasmic reticulum
FA: formic acid
FBS: fetal bovine serum
GAPDH: glyceraldehyde 3-phosphate dehydrogenase
HERPUD1: Homocysteine Inducible ER Protein With Ubiquitin Like Domain 1
IRE1α: inositol-requiring enzyme 1 α
LC-MS: liquid chromatography-mass spectrometry
LB: Luria broth
NS: non-stressed
PCV: packed cell volume
PERK: protein kinase RNA-like endoplasmic reticulum kinase
PIC: protease inhibitor cocktail
PMAIP1: Phorbol-12-Myristate-13-Acetate-Induced Protein 1 (NOXA)
PUMA: BCL2 Binding Component 3 (BBC3)
PVDF: polyvinylidene difluoride
RBP: RNA-binding protein
RBM25: RNA-binding motif protein 25
RIDD: regulated IRE1-dependent decay
RIP: RNA immunoprecipitation
RPLP0: ribosomal protein lateral stalk subunit P0
SERCA: sarco/endoplasmic reticulum Ca²⁺-ATPase
siRNA: small interfering RNA
SOD1: superoxide dismutase 1
TBP: TATA-box binding protein
Tg: thapsigargin
Tm: tunicamycin
UPR: unfolded protein response
WB: western blot
XBP1: X-box binding protein 1
XBP1s: spliced X-box binding protein 1
XBP1u: unspliced X-box binding protein 1
YBX1: Y-box binding protein 1

