## Supplementary material for "RBM25 selectively stabilizes *XBP1s* mRNA independently of splicing factor activity": Suppmental Materials

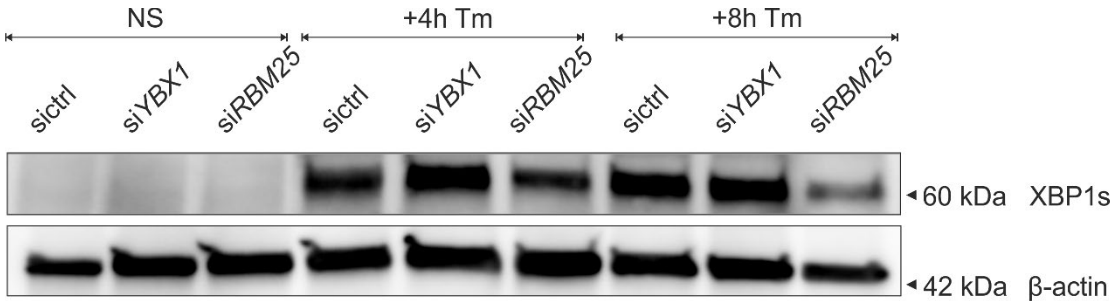


**Supplementary Figure 1. RBM25, but not YBX1, is required for efficient XBP1s stabilization during ER stress.**

Immunoblot analysis of XBP1s protein following ER stress. Under NS conditions, XBP1s protein was minimally detectable across all siRNA treatments. Tm exposure induced marked accumulation of XBP1s protein at 4 and 8 h in siCtrl and siYBX1 cells, whereas RBM25 depletion substantially impaired XBP1s protein accumulation despite comparable β-actin loading. These results indicate that RBM25, but not YBX1, is required for efficient stabilization and accumulation of XBP1s during ER stress, without affecting XBP1u expression.

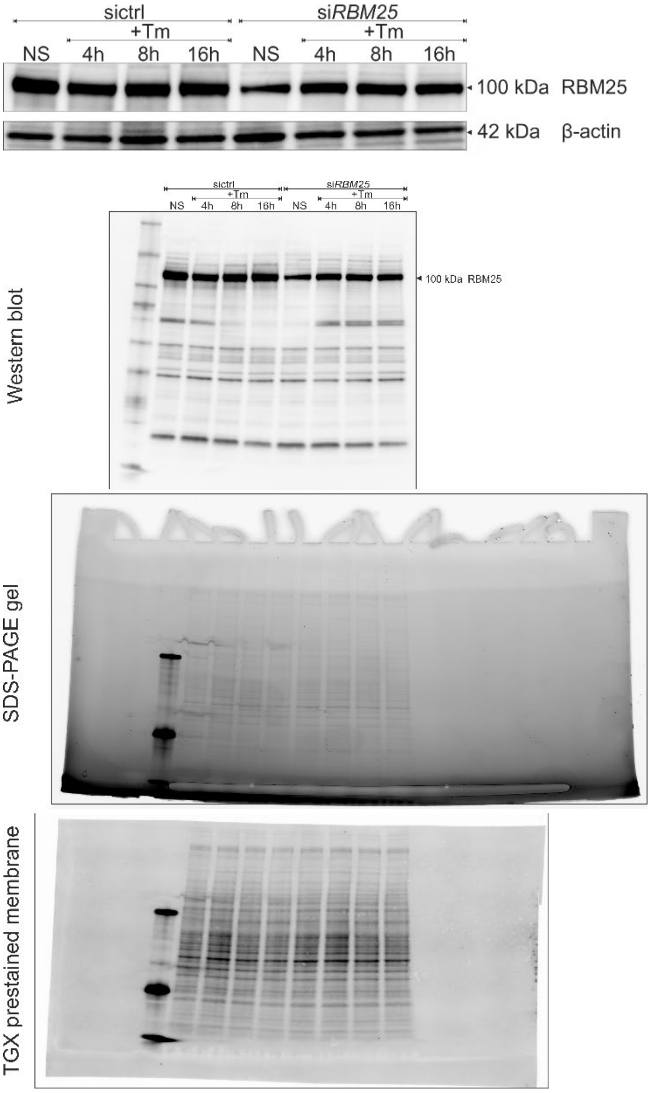


**Supplementary Figure 2. RBM25 protein abundance is stable during ER stress and efficiently reduced upon siRNA mediated depletion.**

Immunoblot analysis of HeLa S3 cells transfected with control siRNA (siCtrl) or RBM25 targeting siRNA (siRBM25) under non stressed (NS) and tunicamycin (Tm) treated conditions. RBM25 protein levels remained stable upon Tm treatment in siCtrl cells, indicating that ER stress does not alter RBM25 abundance. In contrast, RBM25 levels were markedly reduced in siRBM25 cells under both basal and stressed conditions, confirming efficient and sustained knockdown. β actin serves as a loading control. Total protein staining of SDS–PAGE gels and membranes confirmed equal sample loading and transfer across all conditions. These results demonstrate that RBM25 levels are not regulated by ER stress and that the reduction observed in siRBM25 samples reflects specific depletion rather than secondary stress-induced effects.

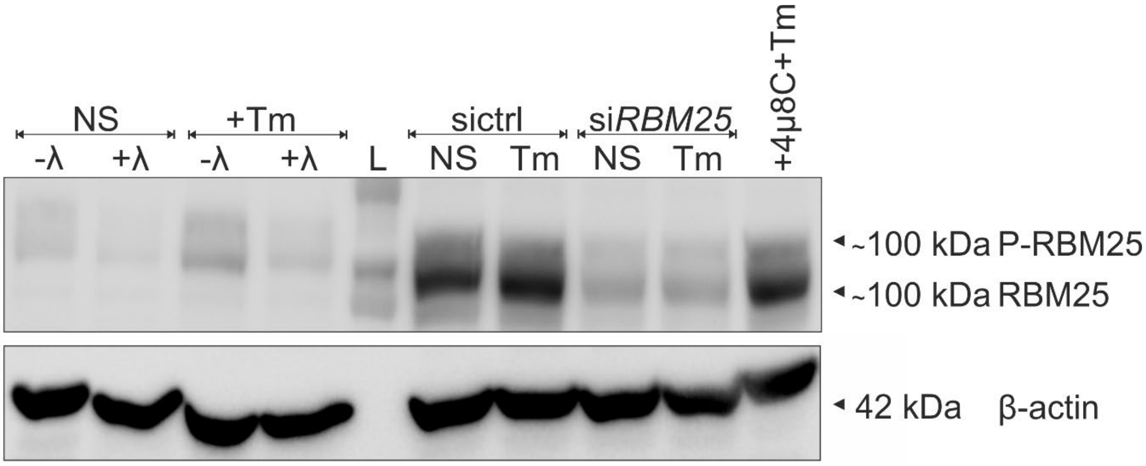


**Supplementary Figure 3. RBM25 phosphorylation is maintained during ER stress independently of IRE1 activity.**

RBM25 mobility was assessed using Phos-tag electrophoresis in HeLa S3 cells under non-stressed (NS) and tunicamycin (Tm)-treated conditions. RBM25 resolved as two species: a faster-migrating non-phosphorylated form (~100 kDa) and a slower-migrating phosphorylated form. Treatment with λ phosphatase (+λ) shifted RBM25 mobility toward the faster-migrating species, confirming basal phosphorylation. Tm treatment did not markedly alter the relative abundance of the phosphorylated RBM25 species, indicating that RBM25 phosphorylation is preserved during ER stress. Consistent with conventional SDS–PAGE analysis, total RBM25 protein levels remained stable under both NS and Tm conditions. *RBM25* knockdown (siRBM25) substantially reduced both total and phosphorylated RBM25 species, confirming efficient depletion. Co-treatment with the IRE1 RNase inhibitor 4μ8C during Tm exposure did not affect the abundance of the phosphorylated RBM25 species, indicating that RBM25 phosphorylation occurs independently of IRE1 activity. β-actin serves as a loading control and showed uniform expression across all samples.

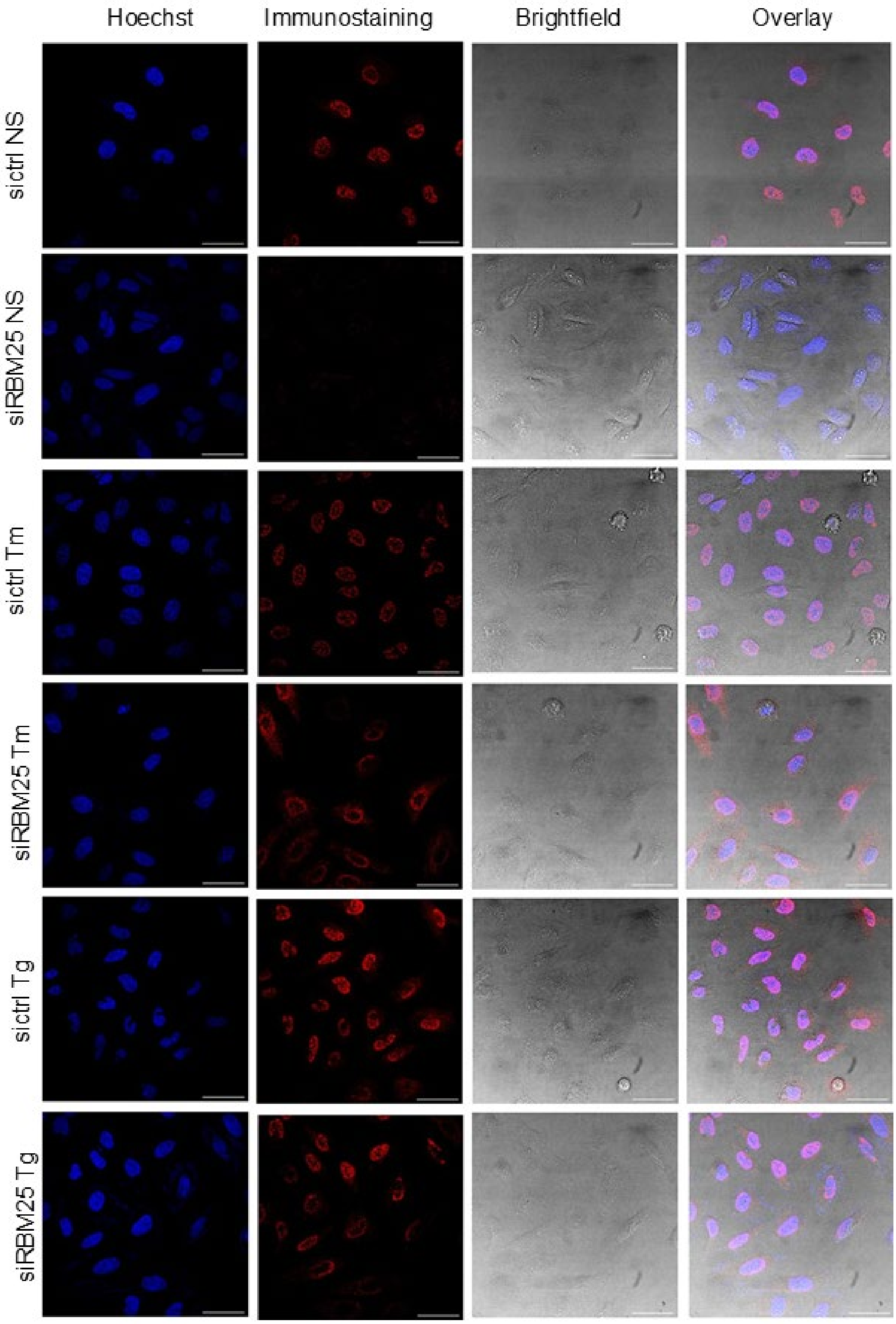


**Supplementary Figure 4. RBM25 nuclear localization is maintained during ER stress.**

Confocal laser scanning microscopy (CLSM) images of HeLa S3 cells transfected with control siRNA (siCtrl) or RBM25-targeting siRNA (siRBM25) under non-stressed (NS) conditions or following treatment with tunicamycin (Tm) or thapsigargin (Tg). RBM25 was detected by immunofluorescence (red), nuclei were visualized using Hoechst staining (blue), and bright-field images are shown in grayscale. Merged images are displayed as overlays. Under NS conditions, siCtrl cells exhibited clear nuclear localization of RBM25, whereas siRBM25 cells showed minimal signal, confirming efficient depletion. Induction of ER stress with Tm or Tg did not alter RBM25 localization in sictrl cells, with the protein remaining predominantly nuclear. In siRBM25 cells, weak diffuse staining was observed under stress conditions, consistent with residual signal. Bright-field images confirmed comparable cell morphology across all conditions. Images were adjusted for contrast only; no additional processing was applied. Scale bar, 50 µm. Imaging was performed using a HC PL APO CS2 63×/1.40 oil immersion objective. Hoechst staining of nucleus (blue, extinction 405 nm, emission 431-497 nm), immunostaining (red, exc. 589 nm, em. 599-677 nm), brighfield channel (gray) and overlay of all three channels.

**Supplementary Figure 5 A.**

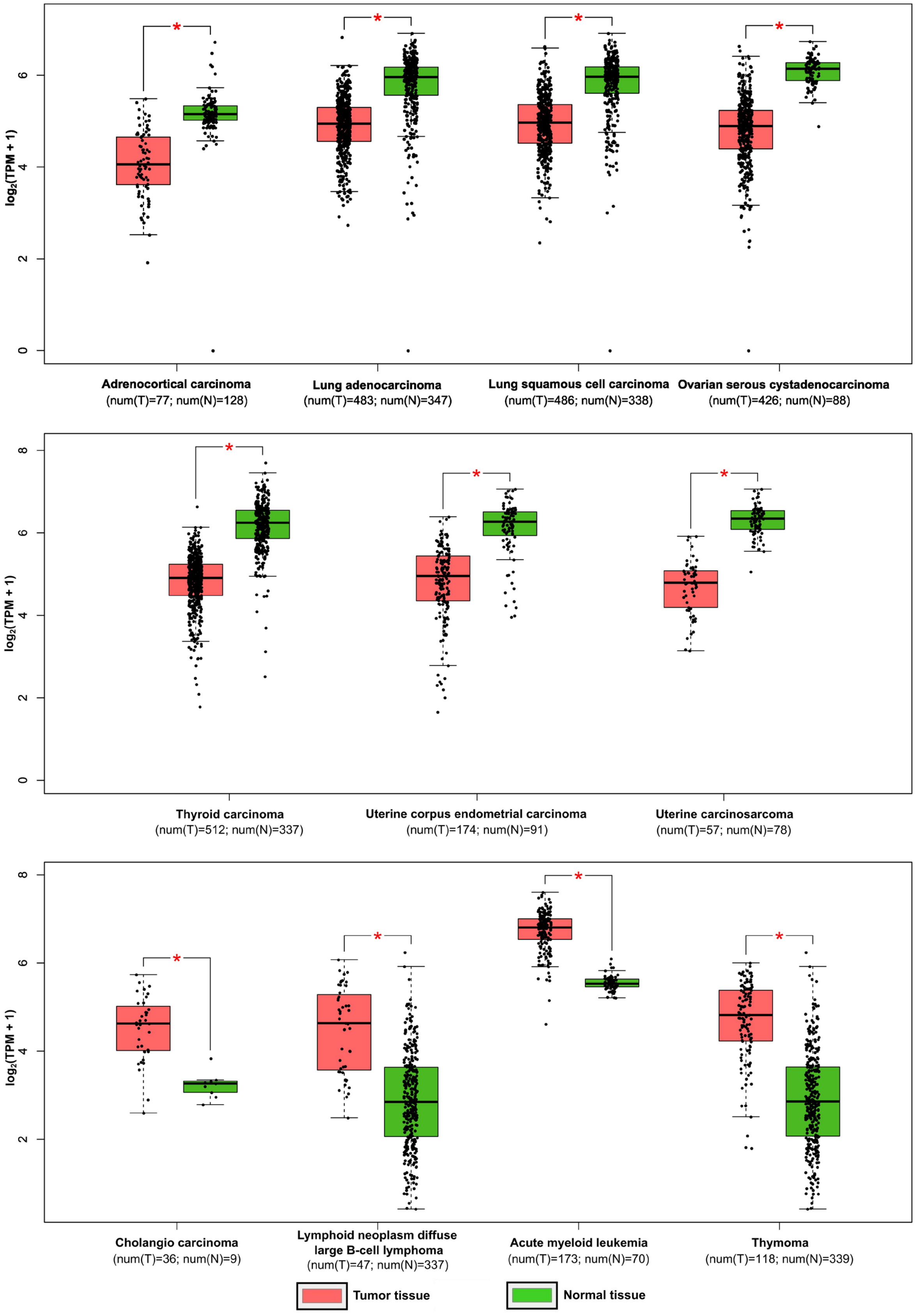


**Supplementary Figure 5 A.** *RBM25* and *XBP1* expression patterns and correlation across human tumor types. ***RBM25* expression in human tumor tissues (red) versus corresponding normal tissues (green)**

**Supplementary Figure 5 B.**

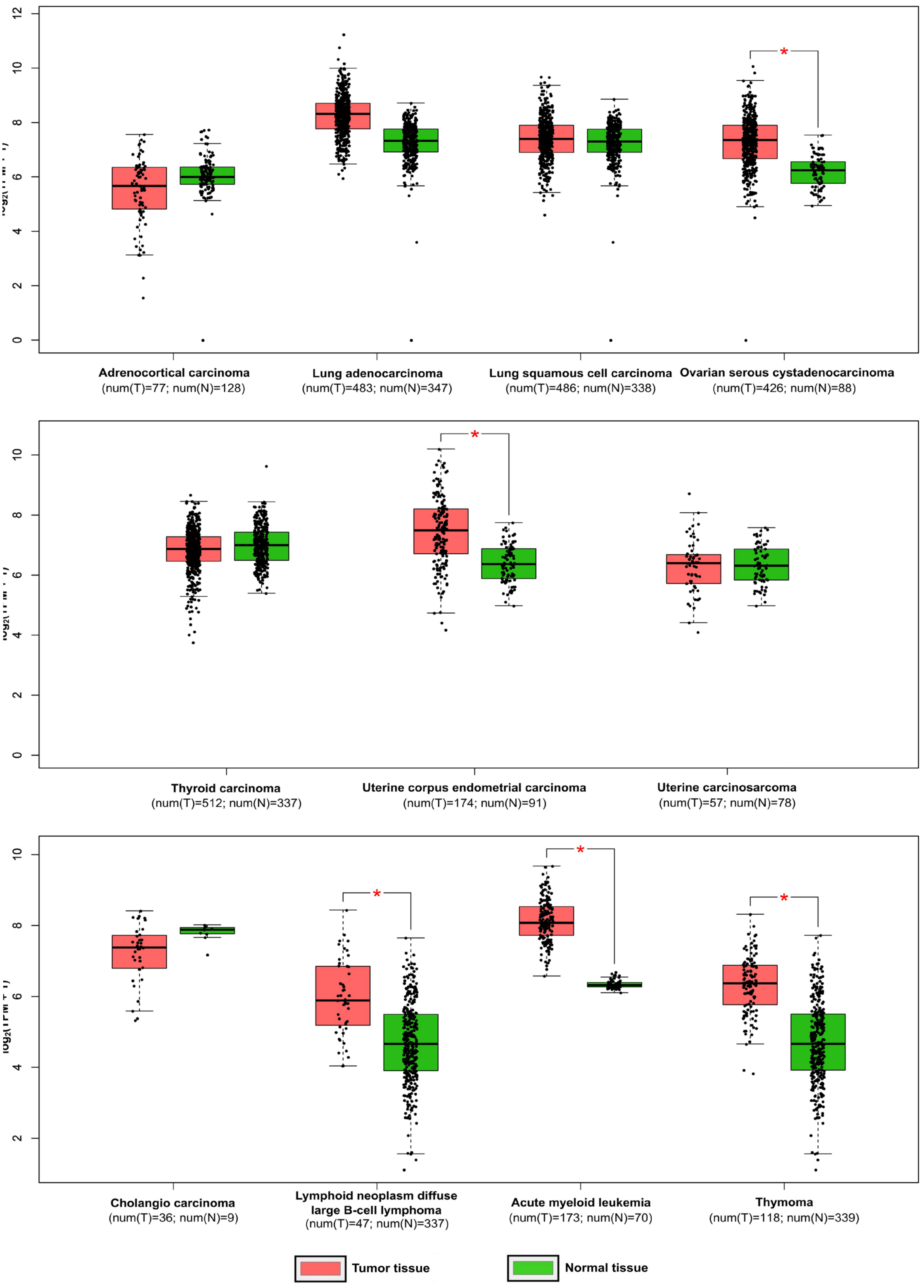


**Supplementary Figure 5B.** *RBM25* and *XBP1* expression patterns and correlation across human tumor types. ***XBP1s* across the same tumor types. *XBP1* expression is shown for tumor (red) and normal (green) tissues.**

**Supplementary Figure C.**

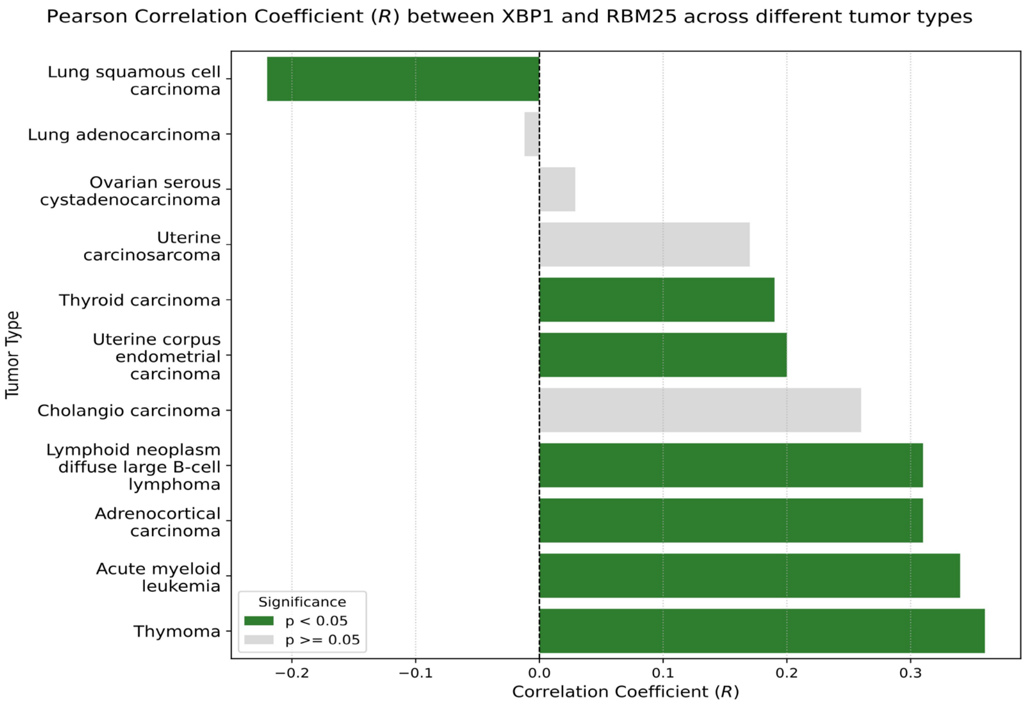


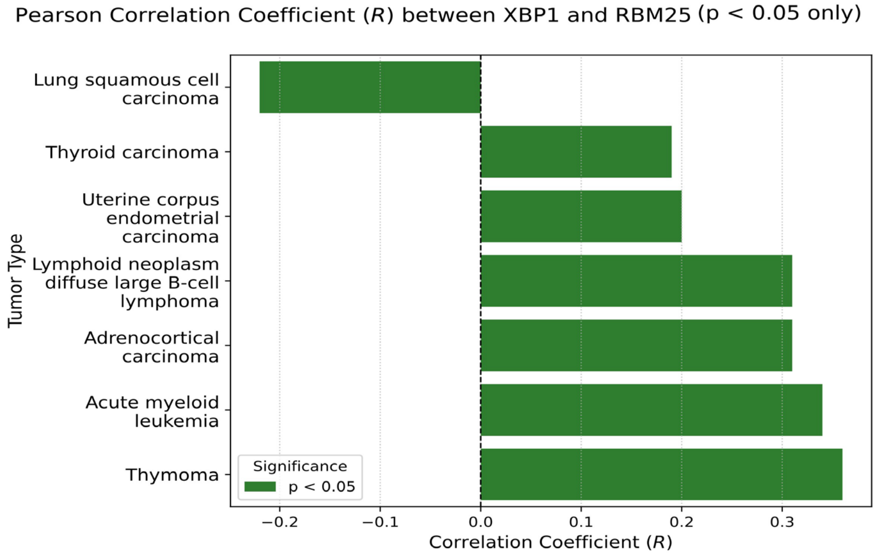


**Supplementary Figure C.** *RBM25* and *XBP1* expression patterns and correlation across human tumor types. **Pearson correlation analysis of *RBM25* and *XBP1* expression across multiple human tumor types.** Green bars indicate tumor types with statistically significant correlations (*P < 0.05), whereas grey bars represent non-significant correlations.

1. Source data: full-length uncropped western blots, TGX prestained SDS-PAGE gel and membrane for **Figure 2A (=Supplementary Figure 2).**

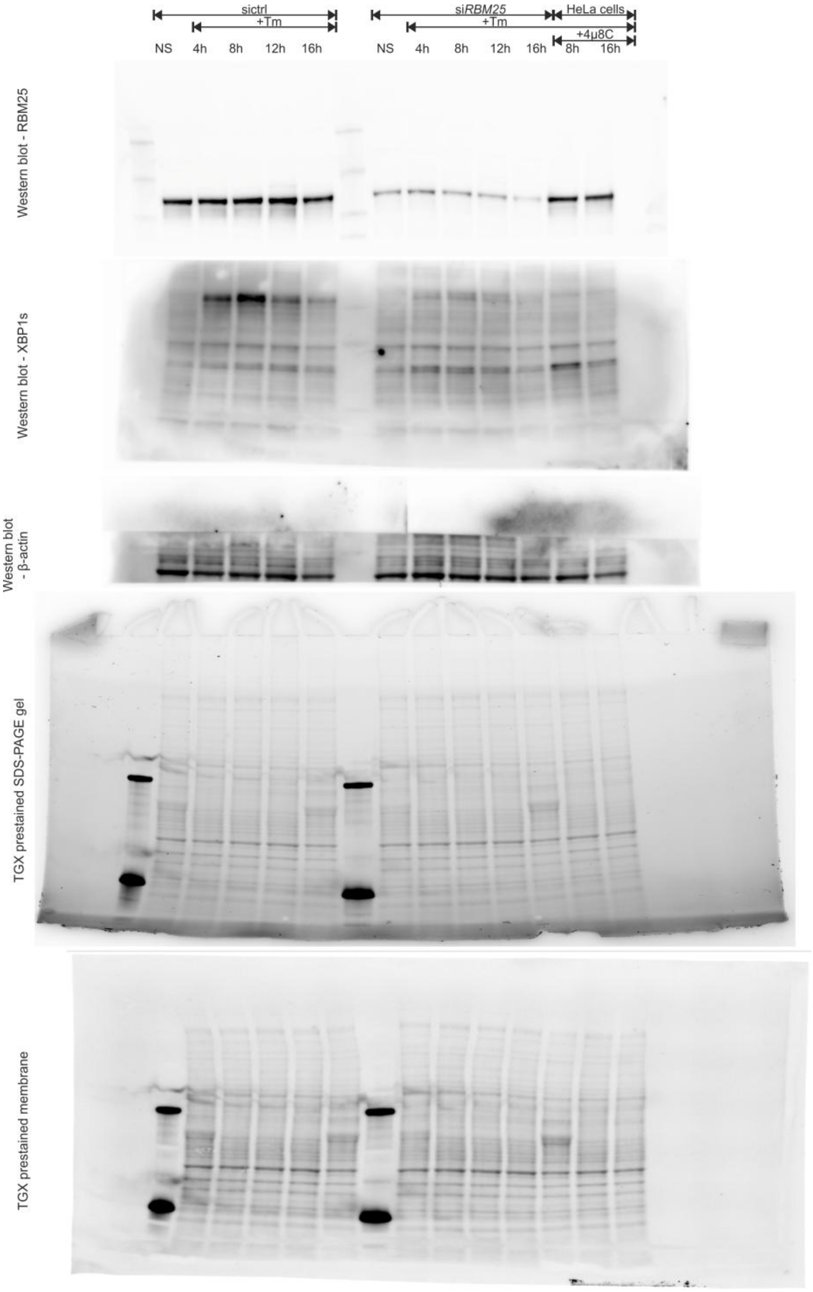

2. Source data: full-length uncropped western blot, TGX prestained SDS-PAGE gel and membrane for **Figure 3A.**

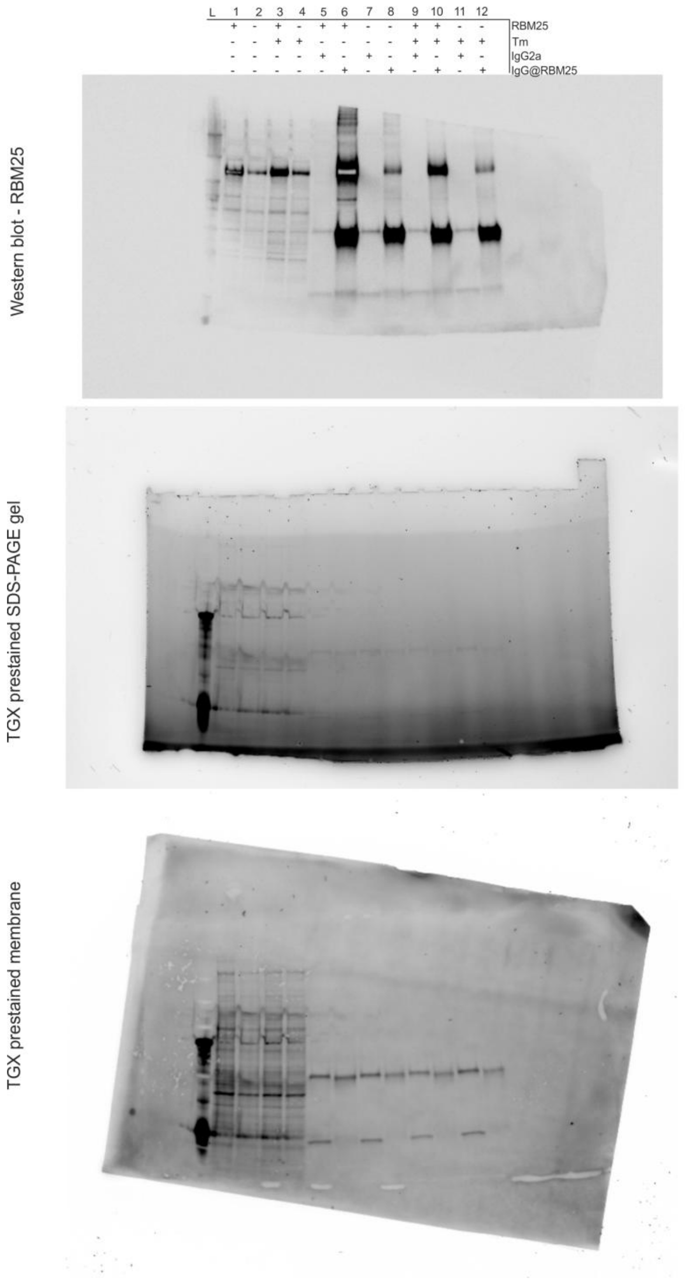

3. Source data: full-length uncropped western blots for **Supplementary Figure 1**.

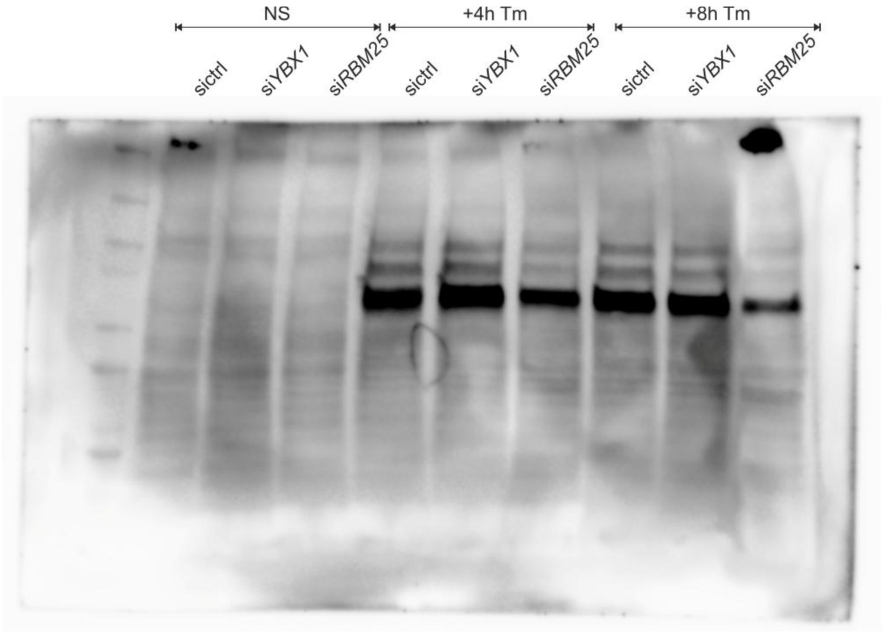


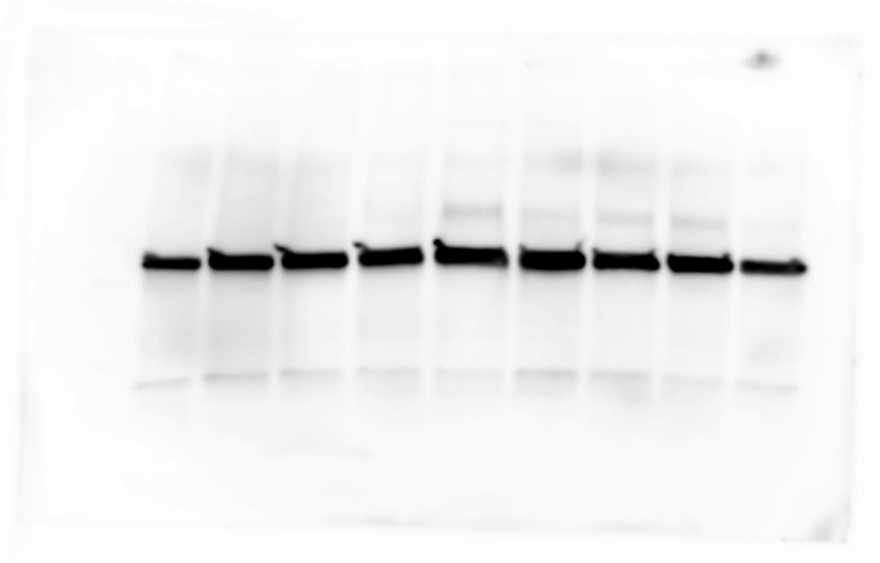

4. Source data: full-length uncropped western blot and TGX-prestained SDS-PAGE gel and membrane for **Figure 2A** for XBP1s (**replicate 2**).

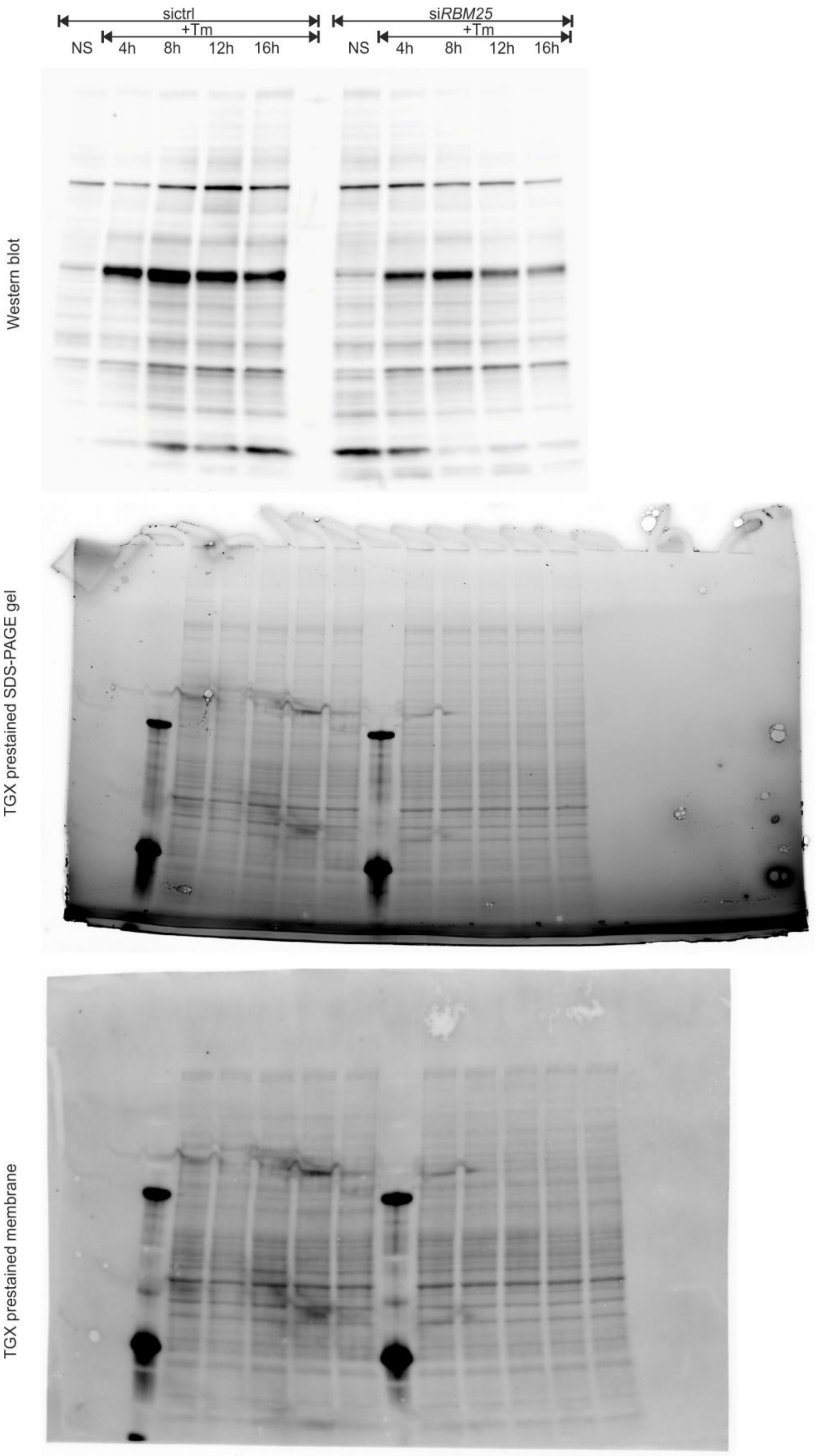

5. Source data: full-length uncropped western blots, TGX prestained SDS-PAGE gel and membrane for **Figure 2A** for RBM25 (**replicate 2**).

Western blot

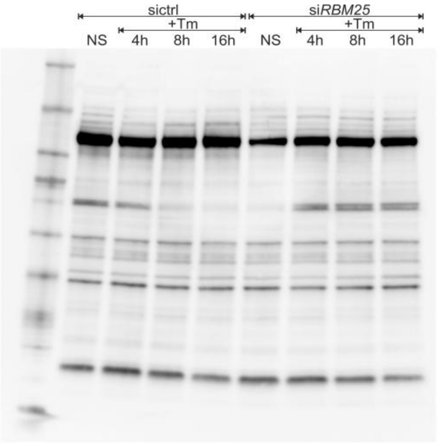


TGX prestained SDS-PAGE gel

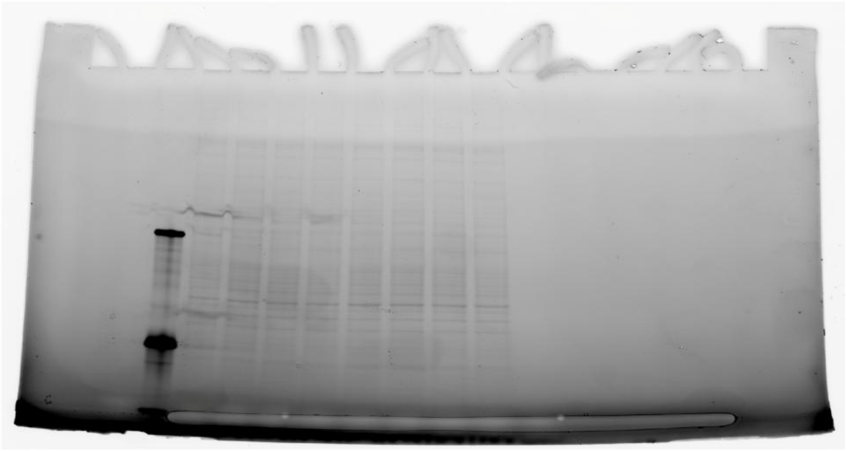

TGX prestained membrane

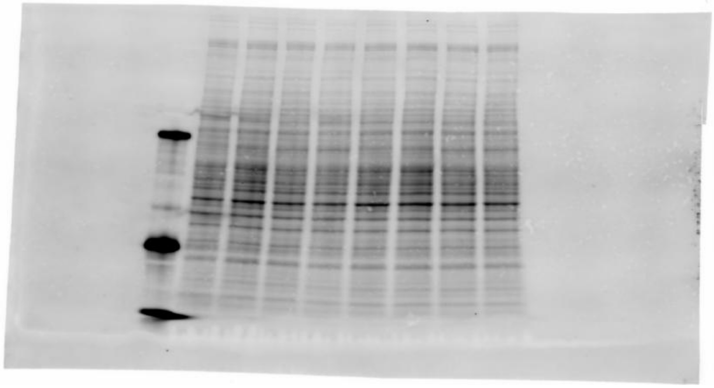

6. Source data: full-length uncropped western blots and TGX prestained SDS-PAGE gel for Figure 2A for XBP1s and RBM25 (**replicate 3**).

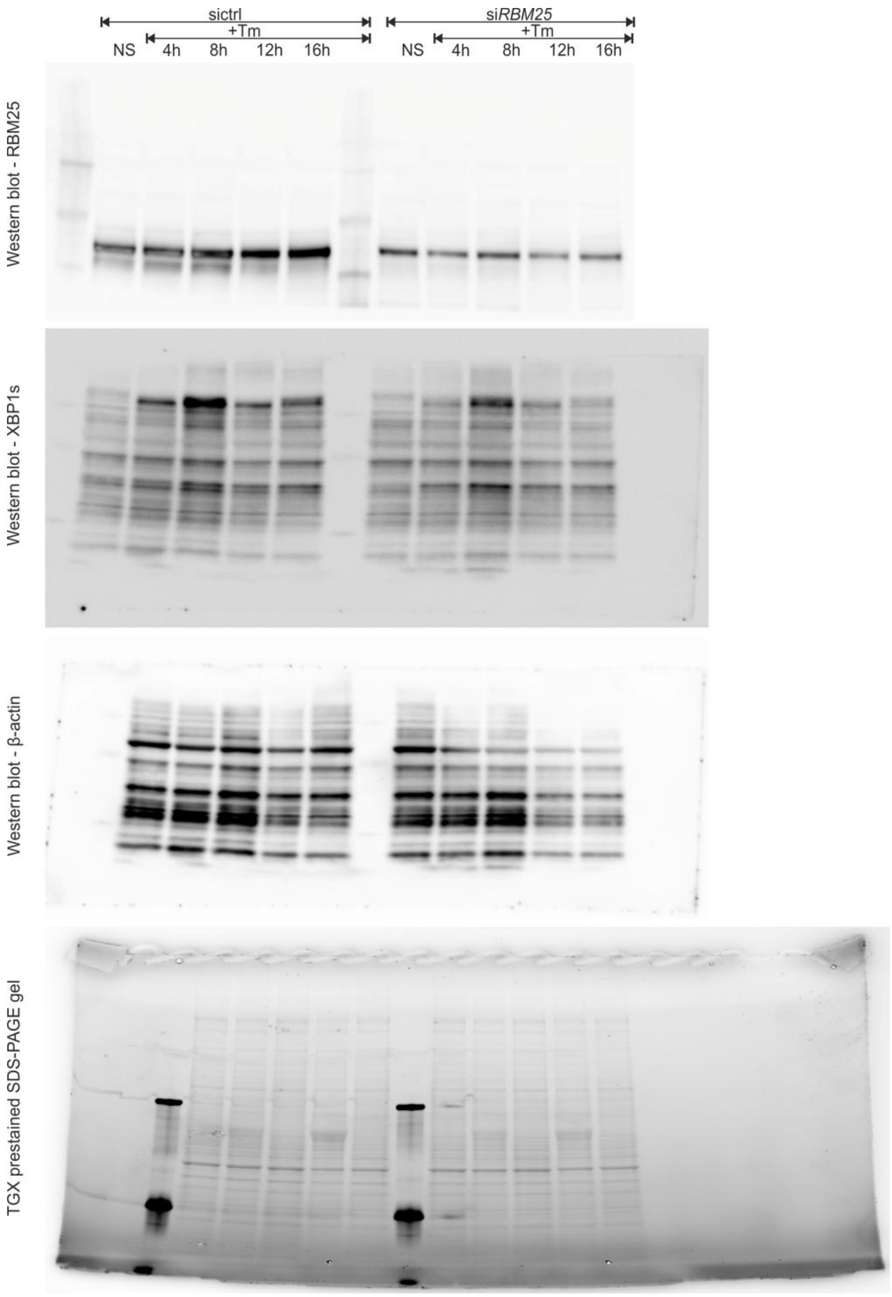

7. Source data: full-length uncropped western blots for **Supplementary Figure 3.**

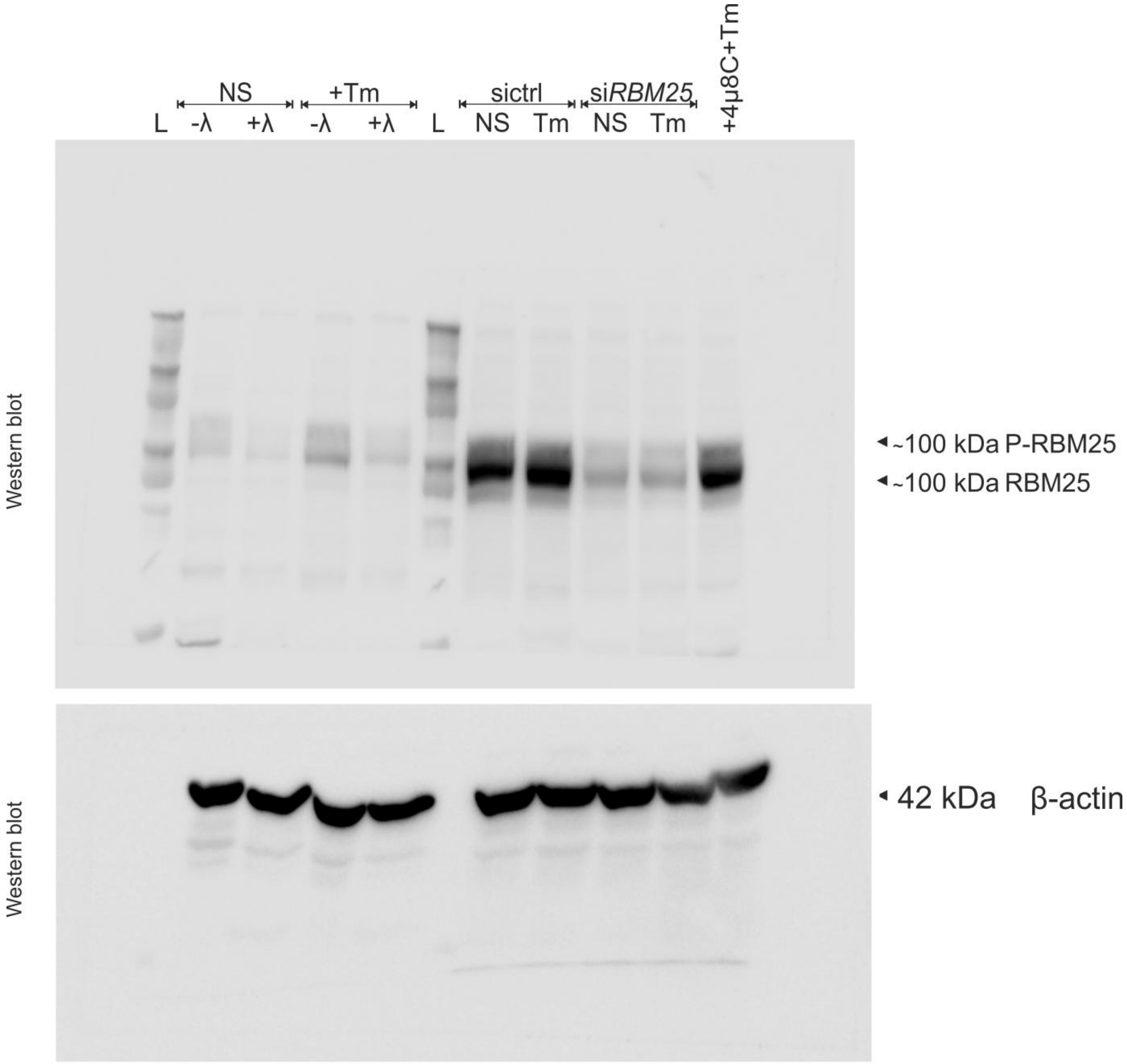
