## Supplementary material for "RBM25 selectively stabilizes *XBP1s* mRNA independently of splicing factor activity": Data set 2: Data Set 2.pdf

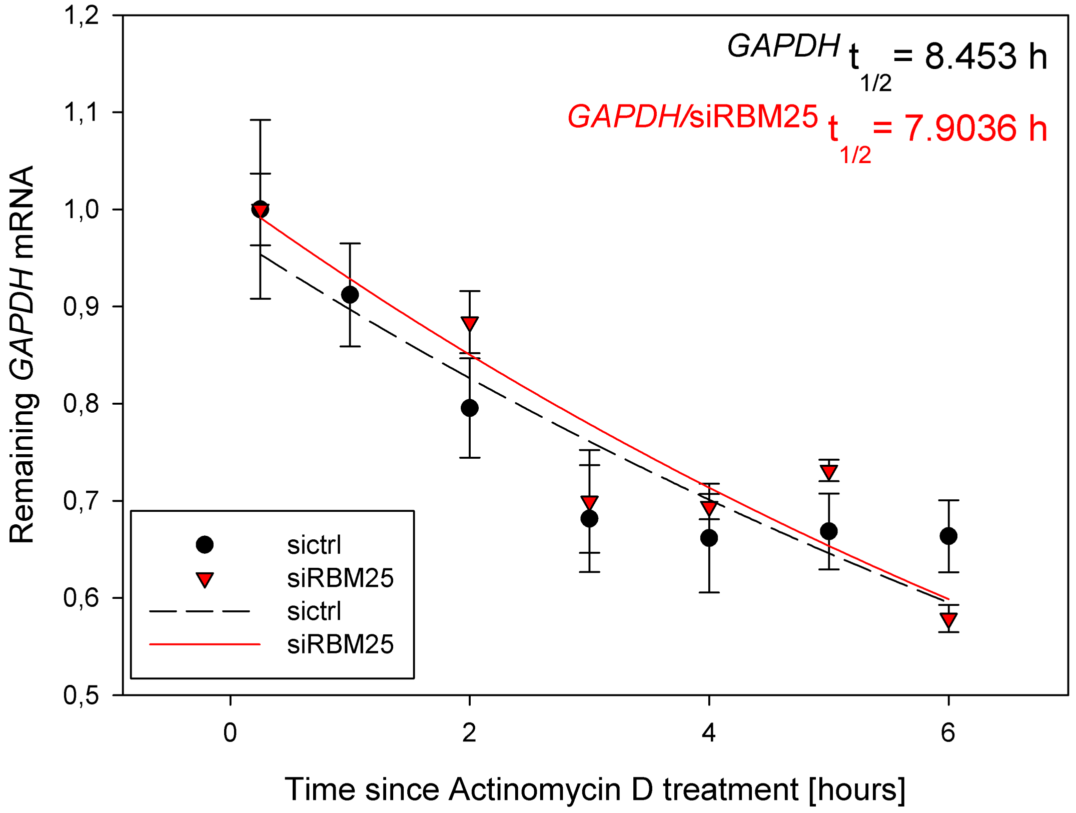

GAPDH t1/2 = 8.453 h
GAPDH/siRBM25 t1/2 = 7.9036 h
Remaining GAPDH mRNA
1,2
1,1
1,0
0,9
0,8
0,7
0,6
0,5
0
2
4
6
Time since Actinomycin D treatment [hours]
sictrl
siRBM25
sictrl
siRBM25

| | 1 | 2 | 3 | 4 | 5 | 6 |
|---|---|---|---|---|---|---|
| 1 | 0,2500 | 1,0000 | 0,0920 | 1,0000 | 0,0370 | 0,2500 |
| 2 | 1,0000 | 0,9120 | 0,0530 | | 0,0170 | 1,0000 |
| 3 | 2,0000 | 0,7956 | 0,0510 | 0,8840 | 0,0320 | 2,0000 |
| 4 | 3,0000 | 0,6817 | 0,0550 | 0,6995 | 0,0530 | 3,0000 |
| 5 | 4,0000 | 0,6616 | 0,0560 | 0,6942 | 0,0130 | 4,0000 |
| 6 | 5,0000 | 0,6686 | 0,0390 | 0,7313 | 0,0110 | 5,0000 |
| 7 | 6,0000 | 0,6636 | 0,0370 | 0,5789 | 0,0140 | 6,0000 |
| 8 | | | | | | |
| 9 | | | | | | |
| 10 | | | | | | |
| 11 | | | | | | |
| 12 | | | | | | |
| 13 | | | | | | |
| 14 | | | | | | |
| 15 | | | | | | |
| 16 | | | | | | |
| 17 | | | | | | |
| 18 | | | | | | |
| 19 | | | | | | |
| 20 | | | | | | |
| 21 | | | | | | |
| 22 | | | | | | |
| 23 | | | | | | |
| 24 | | | | | | |
| 25 | | | | | | |
| 26 | | | | | | |
| 27 | | | | | | |
| 28 | | | | | | |
| 29 | | | | | | |
| 30 | | | | | | |
| 31 | | | | | | |
| 32 | | | | | | |
| 33 | | | | | | |
| 34 | | | | | | |
| 35 | | | | | | |
| 36 | | | | | | |
| 37 | | | | | | |
| 38 | | | | | | |
| 39 | | | | | | |
| 40 | | | | | | |
| 41 | | | | | | |
| 42 | | | | | | |
| 43 | | | | | | |
| 44 | | | | | | |
| 45 | | | | | | |
| 46 | | | | | | |
| 47 | | | | | | |
| 48 | | | | | | |
| 49 | | | | | | |

| | 7-x column | 8-y column | 9 | 10-SumSq | 11-Iterations | 12-a-Final |
|---|---|---|---|---|---|---|
| 1 | 0,2500 | 1,0434 | | 0,0411 | 7,0000 | 1,0972 |
| 2 | 0,2725 | 1,0387 | | 0,0411 | 7,0000 | 1,0972 |
| 3 | 0,2949 | 1,0341 | | 0,0411 | 8,0000 | 1,0972 |
| 4 | 0,3174 | 1,0294 | | 0,0411 | 8,0000 | 1,0972 |
| 5 | 0,3398 | 1,0248 | | 0,0411 | 8,0000 | 1,0972 |
| 6 | 0,3623 | 1,0202 | | 0,0411 | 10,0000 | 1,0972 |
| 7 | 0,3848 | 1,0156 | | 0,0411 | 7,0000 | 1,0972 |
| 8 | 0,4072 | 1,0110 | | 0,0411 | 7,0000 | 1,0972 |
| 9 | 0,4297 | 1,0064 | | 0,0411 | 10,0000 | 1,0972 |
| 10 | 0,4521 | 1,0019 | | 0,0411 | 7,0000 | 1,0972 |
| 11 | 0,4746 | 0,9974 | | 0,0411 | 8,0000 | 1,0972 |
| 12 | 0,4971 | 0,9929 | | 0,0411 | 7,0000 | 1,0972 |
| 13 | 0,5195 | 0,9884 | | 0,0411 | 15,0000 | 1,0972 |
| 14 | 0,5420 | 0,9840 | | 0,0411 | 7,0000 | 1,0972 |
| 15 | 0,5645 | 0,9796 | | 0,0411 | 10,0000 | 1,0972 |
| 16 | 0,5869 | 0,9752 | | 0,0411 | 8,0000 | 1,0972 |
| 17 | 0,6094 | 0,9708 | | 0,0411 | 8,0000 | 1,0972 |
| 18 | 0,6318 | 0,9664 | | 0,0411 | 8,0000 | 1,0972 |
| 19 | 0,6543 | 0,9620 | | 0,0411 | 8,0000 | 1,0972 |
| 20 | 0,6768 | 0,9577 | | 0,0411 | 7,0000 | 1,0972 |
| 21 | 0,6992 | 0,9534 | | 0,0411 | 12,0000 | 1,0972 |
| 22 | 0,7217 | 0,9491 | | 0,0411 | 8,0000 | 1,0972 |
| 23 | 0,7441 | 0,9448 | | 0,0411 | 8,0000 | 1,0972 |
| 24 | 0,7666 | 0,9406 | | 0,0411 | 7,0000 | 1,0972 |
| 25 | 0,7891 | 0,9363 | | 0,0411 | 8,0000 | 1,0972 |
| 26 | 0,8115 | 0,9321 | | 0,0411 | 10,0000 | 1,0972 |
| 27 | 0,8340 | 0,9279 | | 0,0411 | 8,0000 | 1,0972 |
| 28 | 0,8564 | 0,9238 | | 0,0411 | 8,0000 | 1,0972 |
| 29 | 0,8789 | 0,9196 | | 0,0411 | 10,0000 | 1,0972 |
| 30 | 0,9014 | 0,9155 | | 0,0411 | 10,0000 | 1,0972 |
| 31 | 0,9238 | 0,9113 | | 0,0411 | 8,0000 | 1,0972 |
| 32 | 0,9463 | 0,9072 | | 0,0411 | 8,0000 | 1,0972 |
| 33 | 0,9688 | 0,9031 | | 0,0411 | 8,0000 | 1,0972 |
| 34 | 0,9912 | 0,8991 | | 0,0411 | 10,0000 | 1,0972 |
| 35 | 1,0137 | 0,8950 | | 0,0411 | 7,0000 | 1,0972 |
| 36 | 1,0361 | 0,8910 | | 0,0411 | 7,0000 | 1,0972 |
| 37 | 1,0586 | 0,8870 | | 0,0411 | 8,0000 | 1,0972 |
| 38 | 1,0811 | 0,8830 | | 0,0411 | 7,0000 | 1,0972 |
| 39 | 1,1035 | 0,8790 | | 0,0411 | 7,0000 | 1,0972 |
| 40 | 1,1260 | 0,8751 | | 0,0411 | 7,0000 | 1,0972 |
| 41 | 1,1484 | 0,8711 | | 0,0411 | 8,0000 | 1,0972 |
| 42 | 1,1709 | 0,8672 | | 0,0411 | 10,0000 | 1,0972 |
| 43 | 1,1934 | 0,8633 | | 0,0411 | 10,0000 | 1,0972 |
| 44 | 1,2158 | 0,8594 | | 0,0411 | 8,0000 | 1,0972 |
| 45 | 1,2383 | 0,8555 | | 0,0411 | 10,0000 | 1,0972 |
| 46 | 1,2607 | 0,8517 | | 0,0411 | 10,0000 | 1,0972 |
| 47 | 1,2832 | 0,8479 | | 0,0411 | 7,0000 | 1,0972 |
| 48 | 1,3057 | 0,8440 | | 0,0411 | 12,0000 | 1,0972 |
| 49 | 1,3281 | 0,8402 | | 0,0411 | 7,0000 | 1,0972 |

| | 13-b-Final | 14-x column 1 | 15-y column 1 | 16 | 17-SumSq 1 | 18-Iterations 1 |
|---|---|---|---|---|---|---|
| 1 | 0,2009 | 0,2500 | 1,0253 | | 0,0352 | 8,0000 |
| 2 | 0,2009 | 0,2725 | 1,0210 | | 0,0352 | 7,0000 |
| 3 | 0,2009 | 0,2949 | 1,0166 | | 0,0352 | 7,0000 |
| 4 | 0,2009 | 0,3174 | 1,0123 | | 0,0352 | 7,0000 |
| 5 | 0,2009 | 0,3398 | 1,0080 | | 0,0352 | 7,0000 |
| 6 | 0,2009 | 0,3623 | 1,0037 | | 0,0352 | 7,0000 |
| 7 | 0,2009 | 0,3848 | 0,9994 | | 0,0352 | 7,0000 |
| 8 | 0,2009 | 0,4072 | 0,9952 | | 0,0352 | 10,0000 |
| 9 | 0,2009 | 0,4297 | 0,9909 | | 0,0352 | 7,0000 |
| 10 | 0,2009 | 0,4521 | 0,9867 | | 0,0352 | 7,0000 |
| 11 | 0,2009 | 0,4746 | 0,9825 | | 0,0352 | 7,0000 |
| 12 | 0,2009 | 0,4971 | 0,9784 | | 0,0352 | 7,0000 |
| 13 | 0,2009 | 0,5195 | 0,9742 | | 0,0352 | 10,0000 |
| 14 | 0,2009 | 0,5420 | 0,9701 | | 0,0352 | 7,0000 |
| 15 | 0,2009 | 0,5645 | 0,9659 | | 0,0352 | 7,0000 |
| 16 | 0,2009 | 0,5869 | 0,9618 | | 0,0352 | 10,0000 |
| 17 | 0,2009 | 0,6094 | 0,9577 | | 0,0352 | 7,0000 |
| 18 | 0,2009 | 0,6318 | 0,9537 | | 0,0352 | 10,0000 |
| 19 | 0,2009 | 0,6543 | 0,9496 | | 0,0352 | 7,0000 |
| 20 | 0,2009 | 0,6768 | 0,9456 | | 0,0352 | 7,0000 |
| 21 | 0,2009 | 0,6992 | 0,9415 | | 0,0352 | 10,0000 |
| 22 | 0,2009 | 0,7217 | 0,9375 | | 0,0352 | 7,0000 |
| 23 | 0,2009 | 0,7441 | 0,9335 | | 0,0352 | 10,0000 |
| 24 | 0,2009 | 0,7666 | 0,9296 | | 0,0352 | 12,0000 |
| 25 | 0,2009 | 0,7891 | 0,9256 | | 0,0352 | 7,0000 |
| 26 | 0,2009 | 0,8115 | 0,9217 | | 0,0352 | 7,0000 |
| 27 | 0,2009 | 0,8340 | 0,9178 | | 0,0352 | 7,0000 |
| 28 | 0,2009 | 0,8564 | 0,9139 | | 0,0352 | 10,0000 |
| 29 | 0,2009 | 0,8789 | 0,9100 | | 0,0352 | 7,0000 |
| 30 | 0,2009 | 0,9014 | 0,9061 | | 0,0352 | 7,0000 |
| 31 | 0,2009 | 0,9238 | 0,9023 | | 0,0352 | 10,0000 |
| 32 | 0,2009 | 0,9463 | 0,8984 | | 0,0352 | 10,0000 |
| 33 | 0,2009 | 0,9688 | 0,8946 | | 0,0352 | 7,0000 |
| 34 | 0,2009 | 0,9912 | 0,8908 | | 0,0352 | 7,0000 |
| 35 | 0,2009 | 1,0137 | 0,8870 | | 0,0352 | 7,0000 |
| 36 | 0,2009 | 1,0361 | 0,8832 | | 0,0352 | 10,0000 |
| 37 | 0,2009 | 1,0586 | 0,8795 | | 0,0352 | 7,0000 |
| 38 | 0,2009 | 1,0811 | 0,8757 | | 0,0352 | 7,0000 |
| 39 | 0,2009 | 1,1035 | 0,8720 | | 0,0352 | 8,0000 |
| 40 | 0,2009 | 1,1260 | 0,8683 | | 0,0352 | 7,0000 |
| 41 | 0,2009 | 1,1484 | 0,8646 | | 0,0352 | 7,0000 |
| 42 | 0,2009 | 1,1709 | 0,8609 | | 0,0352 | 8,0000 |
| 43 | 0,2009 | 1,1934 | 0,8573 | | 0,0352 | 7,0000 |
| 44 | 0,2009 | 1,2158 | 0,8536 | | 0,0352 | 7,0000 |
| 45 | 0,2009 | 1,2383 | 0,8500 | | 0,0352 | 7,0000 |
| 46 | 0,2009 | 1,2607 | 0,8464 | | 0,0352 | 7,0000 |
| 47 | 0,2009 | 1,2832 | 0,8428 | | 0,0352 | 7,0000 |
| 48 | 0,2009 | 1,3057 | 0,8392 | | 0,0352 | 7,0000 |
| 49 | 0,2009 | 1,3281 | 0,8356 | | 0,0352 | 7,0000 |

| | 19-a-Final 1 | 20-b-Final 1 | 21-x column 2 | 22-y column 2 | 23 | 24-SumSq 2 |
|---|---|---|---|---|---|---|
| 1 | 1,0751 | 0,1897 | 0,2500 | 0,9536 | | 0,0164 |
| 2 | 1,0751 | 0,1897 | 0,2725 | 0,9518 | | 0,0164 |
| 3 | 1,0751 | 0,1897 | 0,2949 | 0,9501 | | 0,0164 |
| 4 | 1,0751 | 0,1897 | 0,3174 | 0,9483 | | 0,0164 |
| 5 | 1,0751 | 0,1897 | 0,3398 | 0,9466 | | 0,0164 |
| 6 | 1,0751 | 0,1897 | 0,3623 | 0,9448 | | 0,0164 |
| 7 | 1,0751 | 0,1897 | 0,3848 | 0,9431 | | 0,0164 |
| 8 | 1,0751 | 0,1897 | 0,4072 | 0,9414 | | 0,0164 |
| 9 | 1,0751 | 0,1897 | 0,4297 | 0,9396 | | 0,0164 |
| 10 | 1,0751 | 0,1897 | 0,4521 | 0,9379 | | 0,0164 |
| 11 | 1,0751 | 0,1897 | 0,4746 | 0,9362 | | 0,0164 |
| 12 | 1,0751 | 0,1897 | 0,4971 | 0,9345 | | 0,0164 |
| 13 | 1,0751 | 0,1897 | 0,5195 | 0,9327 | | 0,0164 |
| 14 | 1,0751 | 0,1897 | 0,5420 | 0,9310 | | 0,0164 |
| 15 | 1,0751 | 0,1897 | 0,5645 | 0,9293 | | 0,0164 |
| 16 | 1,0751 | 0,1897 | 0,5869 | 0,9276 | | 0,0164 |
| 17 | 1,0751 | 0,1897 | 0,6094 | 0,9259 | | 0,0164 |
| 18 | 1,0751 | 0,1897 | 0,6318 | 0,9242 | | 0,0164 |
| 19 | 1,0751 | 0,1897 | 0,6543 | 0,9225 | | 0,0164 |
| 20 | 1,0751 | 0,1897 | 0,6768 | 0,9208 | | 0,0164 |
| 21 | 1,0751 | 0,1897 | 0,6992 | 0,9191 | | 0,0164 |
| 22 | 1,0751 | 0,1897 | 0,7217 | 0,9174 | | 0,0164 |
| 23 | 1,0751 | 0,1897 | 0,7441 | 0,9157 | | 0,0164 |
| 24 | 1,0751 | 0,1897 | 0,7666 | 0,9140 | | 0,0164 |
| 25 | 1,0751 | 0,1897 | 0,7891 | 0,9124 | | 0,0164 |
| 26 | 1,0751 | 0,1897 | 0,8115 | 0,9107 | | 0,0164 |
| 27 | 1,0751 | 0,1897 | 0,8340 | 0,9090 | | 0,0164 |
| 28 | 1,0751 | 0,1897 | 0,8564 | 0,9073 | | 0,0164 |
| 29 | 1,0751 | 0,1897 | 0,8789 | 0,9057 | | 0,0164 |
| 30 | 1,0751 | 0,1897 | 0,9014 | 0,9040 | | 0,0164 |
| 31 | 1,0751 | 0,1897 | 0,9238 | 0,9023 | | 0,0164 |
| 32 | 1,0751 | 0,1897 | 0,9463 | 0,9007 | | 0,0164 |
| 33 | 1,0751 | 0,1897 | 0,9688 | 0,8990 | | 0,0164 |
| 34 | 1,0751 | 0,1897 | 0,9912 | 0,8974 | | 0,0164 |
| 35 | 1,0751 | 0,1897 | 1,0137 | 0,8957 | | 0,0164 |
| 36 | 1,0751 | 0,1897 | 1,0361 | 0,8941 | | 0,0164 |
| 37 | 1,0751 | 0,1897 | 1,0586 | 0,8924 | | 0,0164 |
| 38 | 1,0751 | 0,1897 | 1,0811 | 0,8908 | | 0,0164 |
| 39 | 1,0751 | 0,1897 | 1,1035 | 0,8891 | | 0,0164 |
| 40 | 1,0751 | 0,1897 | 1,1260 | 0,8875 | | 0,0164 |
| 41 | 1,0751 | 0,1897 | 1,1484 | 0,8859 | | 0,0164 |
| 42 | 1,0751 | 0,1897 | 1,1709 | 0,8842 | | 0,0164 |
| 43 | 1,0751 | 0,1897 | 1,1934 | 0,8826 | | 0,0164 |
| 44 | 1,0751 | 0,1897 | 1,2158 | 0,8810 | | 0,0164 |
| 45 | 1,0751 | 0,1897 | 1,2383 | 0,8794 | | 0,0164 |
| 46 | 1,0751 | 0,1897 | 1,2607 | 0,8777 | | 0,0164 |
| 47 | 1,0751 | 0,1897 | 1,2832 | 0,8761 | | 0,0164 |
| 48 | 1,0751 | 0,1897 | 1,3057 | 0,8745 | | 0,0164 |
| 49 | 1,0751 | 0,1897 | 1,3281 | 0,8729 | | 0,0164 |

| | 25-Iterations 2 | 26-a-Final 2 | 27-b-Final 2 | 28-x column 3 | 29-y column 3 | 30 |
|---|---|---|---|---|---|---|
| 1 | 9,0000 | 0,9733 | 0,0820 | 0,2500 | 1,0227 | |
| 2 | 9,0000 | 0,9733 | 0,0820 | 0,2725 | 1,0206 | |
| 3 | 9,0000 | 0,9733 | 0,0820 | 0,2949 | 1,0184 | |
| 4 | 9,0000 | 0,9733 | 0,0820 | 0,3174 | 1,0162 | |
| 5 | 9,0000 | 0,9733 | 0,0820 | 0,3398 | 1,0141 | |
| 6 | 9,0000 | 0,9733 | 0,0820 | 0,3623 | 1,0119 | |
| 7 | 9,0000 | 0,9733 | 0,0820 | 0,3848 | 1,0098 | |
| 8 | 9,0000 | 0,9733 | 0,0820 | 0,4072 | 1,0076 | |
| 9 | 9,0000 | 0,9733 | 0,0820 | 0,4297 | 1,0055 | |
| 10 | 9,0000 | 0,9733 | 0,0820 | 0,4521 | 1,0034 | |
| 11 | 9,0000 | 0,9733 | 0,0820 | 0,4746 | 1,0013 | |
| 12 | 9,0000 | 0,9733 | 0,0820 | 0,4971 | 0,9991 | |
| 13 | 9,0000 | 0,9733 | 0,0820 | 0,5195 | 0,9970 | |
| 14 | 9,0000 | 0,9733 | 0,0820 | 0,5420 | 0,9949 | |
| 15 | 9,0000 | 0,9733 | 0,0820 | 0,5645 | 0,9928 | |
| 16 | 9,0000 | 0,9733 | 0,0820 | 0,5869 | 0,9907 | |
| 17 | 9,0000 | 0,9733 | 0,0820 | 0,6094 | 0,9886 | |
| 18 | 9,0000 | 0,9733 | 0,0820 | 0,6318 | 0,9865 | |
| 19 | 9,0000 | 0,9733 | 0,0820 | 0,6543 | 0,9844 | |
| 20 | 9,0000 | 0,9733 | 0,0820 | 0,6768 | 0,9823 | |
| 21 | 9,0000 | 0,9733 | 0,0820 | 0,6992 | 0,9802 | |
| 22 | 9,0000 | 0,9733 | 0,0820 | 0,7217 | 0,9782 | |
| 23 | 9,0000 | 0,9733 | 0,0820 | 0,7441 | 0,9761 | |
| 24 | 9,0000 | 0,9733 | 0,0820 | 0,7666 | 0,9740 | |
| 25 | 9,0000 | 0,9733 | 0,0820 | 0,7891 | 0,9719 | |
| 26 | 9,0000 | 0,9733 | 0,0820 | 0,8115 | 0,9699 | |
| 27 | 9,0000 | 0,9733 | 0,0820 | 0,8340 | 0,9678 | |
| 28 | 9,0000 | 0,9733 | 0,0820 | 0,8564 | 0,9658 | |
| 29 | 9,0000 | 0,9733 | 0,0820 | 0,8789 | 0,9637 | |
| 30 | 9,0000 | 0,9733 | 0,0820 | 0,9014 | 0,9617 | |
| 31 | 9,0000 | 0,9733 | 0,0820 | 0,9238 | 0,9597 | |
| 32 | 9,0000 | 0,9733 | 0,0820 | 0,9463 | 0,9576 | |
| 33 | 9,0000 | 0,9733 | 0,0820 | 0,9688 | 0,9556 | |
| 34 | 9,0000 | 0,9733 | 0,0820 | 0,9912 | 0,9536 | |
| 35 | 9,0000 | 0,9733 | 0,0820 | 1,0137 | 0,9515 | |
| 36 | 9,0000 | 0,9733 | 0,0820 | 1,0361 | 0,9495 | |
| 37 | 8,0000 | 0,9733 | 0,0820 | 1,0586 | 0,9475 | |
| 38 | 9,0000 | 0,9733 | 0,0820 | 1,0811 | 0,9455 | |
| 39 | 9,0000 | 0,9733 | 0,0820 | 1,1035 | 0,9435 | |
| 40 | 9,0000 | 0,9733 | 0,0820 | 1,1260 | 0,9415 | |
| 41 | 9,0000 | 0,9733 | 0,0820 | 1,1484 | 0,9395 | |
| 42 | 9,0000 | 0,9733 | 0,0820 | 1,1709 | 0,9375 | |
| 43 | 9,0000 | 0,9733 | 0,0820 | 1,1934 | 0,9355 | |
| 44 | 9,0000 | 0,9733 | 0,0820 | 1,2158 | 0,9335 | |
| 45 | 9,0000 | 0,9733 | 0,0820 | 1,2383 | 0,9316 | |
| 46 | 9,0000 | 0,9733 | 0,0820 | 1,2607 | 0,9296 | |
| 47 | 9,0000 | 0,9733 | 0,0820 | 1,2832 | 0,9276 | |
| 48 | 9,0000 | 0,9733 | 0,0820 | 1,3057 | 0,9257 | |
| 49 | 9,0000 | 0,9733 | 0,0820 | 1,3281 | 0,9237 | |

| | 31-SumSq 3 | 32-Iterations 3 | 33-a-Final 3 | 34-b-Final 3 | 35-x column 4 | 36-y column 4 |
|---|---|---|---|---|---|---|
| 1 | 0,0189 | 8,0000 | 1,0472 | 0,0945 | 0,2500 | 0,9915 |
| 2 | 0,0189 | 8,0000 | 1,0472 | 0,0945 | 0,2725 | 0,9895 |
| 3 | 0,0189 | 8,0000 | 1,0472 | 0,0945 | 0,2949 | 0,9876 |
| 4 | 0,0189 | 8,0000 | 1,0472 | 0,0945 | 0,3174 | 0,9856 |
| 5 | 0,0189 | 8,0000 | 1,0472 | 0,0945 | 0,3398 | 0,9837 |
| 6 | 0,0189 | 8,0000 | 1,0472 | 0,0945 | 0,3623 | 0,9818 |
| 7 | 0,0189 | 7,0000 | 1,0472 | 0,0945 | 0,3848 | 0,9798 |
| 8 | 0,0189 | 8,0000 | 1,0472 | 0,0945 | 0,4072 | 0,9779 |
| 9 | 0,0189 | 8,0000 | 1,0472 | 0,0945 | 0,4297 | 0,9760 |
| 10 | 0,0189 | 8,0000 | 1,0472 | 0,0945 | 0,4521 | 0,9740 |
| 11 | 0,0189 | 8,0000 | 1,0472 | 0,0945 | 0,4746 | 0,9721 |
| 12 | 0,0189 | 8,0000 | 1,0472 | 0,0945 | 0,4971 | 0,9702 |
| 13 | 0,0189 | 8,0000 | 1,0472 | 0,0945 | 0,5195 | 0,9683 |
| 14 | 0,0189 | 8,0000 | 1,0472 | 0,0945 | 0,5420 | 0,9664 |
| 15 | 0,0189 | 8,0000 | 1,0472 | 0,0945 | 0,5645 | 0,9645 |
| 16 | 0,0189 | 8,0000 | 1,0472 | 0,0945 | 0,5869 | 0,9626 |
| 17 | 0,0189 | 8,0000 | 1,0472 | 0,0945 | 0,6094 | 0,9607 |
| 18 | 0,0189 | 8,0000 | 1,0472 | 0,0945 | 0,6318 | 0,9588 |
| 19 | 0,0189 | 8,0000 | 1,0472 | 0,0945 | 0,6543 | 0,9569 |
| 20 | 0,0189 | 8,0000 | 1,0472 | 0,0945 | 0,6768 | 0,9551 |
| 21 | 0,0189 | 8,0000 | 1,0472 | 0,0945 | 0,6992 | 0,9532 |
| 22 | 0,0189 | 8,0000 | 1,0472 | 0,0945 | 0,7217 | 0,9513 |
| 23 | 0,0189 | 8,0000 | 1,0472 | 0,0945 | 0,7441 | 0,9494 |
| 24 | 0,0189 | 8,0000 | 1,0472 | 0,0945 | 0,7666 | 0,9476 |
| 25 | 0,0189 | 7,0000 | 1,0472 | 0,0945 | 0,7891 | 0,9457 |
| 26 | 0,0189 | 8,0000 | 1,0472 | 0,0945 | 0,8115 | 0,9438 |
| 27 | 0,0189 | 8,0000 | 1,0472 | 0,0945 | 0,8340 | 0,9420 |
| 28 | 0,0189 | 8,0000 | 1,0472 | 0,0945 | 0,8564 | 0,9401 |
| 29 | 0,0189 | 8,0000 | 1,0472 | 0,0945 | 0,8789 | 0,9383 |
| 30 | 0,0189 | 8,0000 | 1,0472 | 0,0945 | 0,9014 | 0,9364 |
| 31 | 0,0189 | 8,0000 | 1,0472 | 0,0945 | 0,9238 | 0,9346 |
| 32 | 0,0189 | 8,0000 | 1,0472 | 0,0945 | 0,9463 | 0,9327 |
| 33 | 0,0189 | 8,0000 | 1,0472 | 0,0945 | 0,9688 | 0,9309 |
| 34 | 0,0189 | 8,0000 | 1,0472 | 0,0945 | 0,9912 | 0,9291 |
| 35 | 0,0189 | 8,0000 | 1,0472 | 0,0945 | 1,0137 | 0,9272 |
| 36 | 0,0189 | 8,0000 | 1,0472 | 0,0945 | 1,0361 | 0,9254 |
| 37 | 0,0189 | 8,0000 | 1,0472 | 0,0945 | 1,0586 | 0,9236 |
| 38 | 0,0189 | 8,0000 | 1,0472 | 0,0945 | 1,0811 | 0,9218 |
| 39 | 0,0189 | 7,0000 | 1,0472 | 0,0945 | 1,1035 | 0,9200 |
| 40 | 0,0189 | 8,0000 | 1,0472 | 0,0945 | 1,1260 | 0,9182 |
| 41 | 0,0189 | 8,0000 | 1,0472 | 0,0945 | 1,1484 | 0,9164 |
| 42 | 0,0189 | 8,0000 | 1,0472 | 0,0945 | 1,1709 | 0,9145 |
| 43 | 0,0189 | 8,0000 | 1,0472 | 0,0945 | 1,1934 | 0,9127 |
| 44 | 0,0189 | 8,0000 | 1,0472 | 0,0945 | 1,2158 | 0,9110 |
| 45 | 0,0189 | 8,0000 | 1,0472 | 0,0945 | 1,2383 | 0,9092 |
| 46 | 0,0189 | 8,0000 | 1,0472 | 0,0945 | 1,2607 | 0,9074 |
| 47 | 0,0189 | 8,0000 | 1,0472 | 0,0945 | 1,2832 | 0,9056 |
| 48 | 0,0189 | 7,0000 | 1,0472 | 0,0945 | 1,3057 | 0,9038 |
| 49 | 0,0189 | 8,0000 | 1,0472 | 0,0945 | 1,3281 | 0,9020 |

| | 37 | 38-SumSq 4 | 39-Iterations 4 | 40-a-Final 4 | 41-b-Final 4 |
|---|---|---|---|---|---|
| 1 | | 0,0143 | 8,0000 | 1,0134 | 0,0877 |
| 2 | | 0,0143 | 8,0000 | 1,0134 | 0,0877 |
| 3 | | 0,0143 | 8,0000 | 1,0134 | 0,0877 |
| 4 | | 0,0143 | 8,0000 | 1,0134 | 0,0877 |
| 5 | | 0,0143 | 8,0000 | 1,0134 | 0,0877 |
| 6 | | 0,0143 | 8,0000 | 1,0134 | 0,0877 |
| 7 | | 0,0143 | 8,0000 | 1,0134 | 0,0877 |
| 8 | | 0,0143 | 8,0000 | 1,0134 | 0,0877 |
| 9 | | 0,0143 | 8,0000 | 1,0134 | 0,0877 |
| 10 | | 0,0143 | 8,0000 | 1,0134 | 0,0877 |
| 11 | | 0,0143 | 8,0000 | 1,0134 | 0,0877 |
| 12 | | 0,0143 | 8,0000 | 1,0134 | 0,0877 |
| 13 | | 0,0143 | 8,0000 | 1,0134 | 0,0877 |
| 14 | | 0,0143 | 8,0000 | 1,0134 | 0,0877 |
| 15 | | 0,0143 | 8,0000 | 1,0134 | 0,0877 |
| 16 | | 0,0143 | 8,0000 | 1,0134 | 0,0877 |
| 17 | | 0,0143 | 8,0000 | 1,0134 | 0,0877 |
| 18 | | 0,0143 | 8,0000 | 1,0134 | 0,0877 |
| 19 | | 0,0143 | 8,0000 | 1,0134 | 0,0877 |
| 20 | | 0,0143 | 8,0000 | 1,0134 | 0,0877 |
| 21 | | 0,0143 | 8,0000 | 1,0134 | 0,0877 |
| 22 | | 0,0143 | 8,0000 | 1,0134 | 0,0877 |
| 23 | | 0,0143 | 8,0000 | 1,0134 | 0,0877 |
| 24 | | 0,0143 | 8,0000 | 1,0134 | 0,0877 |
| 25 | | 0,0143 | 8,0000 | 1,0134 | 0,0877 |
| 26 | | 0,0143 | 8,0000 | 1,0134 | 0,0877 |
| 27 | | 0,0143 | 8,0000 | 1,0134 | 0,0877 |
| 28 | | 0,0143 | 8,0000 | 1,0134 | 0,0877 |
| 29 | | 0,0143 | 8,0000 | 1,0134 | 0,0877 |
| 30 | | 0,0143 | 8,0000 | 1,0134 | 0,0877 |
| 31 | | 0,0143 | 8,0000 | 1,0134 | 0,0877 |
| 32 | | 0,0143 | 8,0000 | 1,0134 | 0,0877 |
| 33 | | 0,0143 | 7,0000 | 1,0134 | 0,0877 |
| 34 | | 0,0143 | 8,0000 | 1,0134 | 0,0877 |
| 35 | | 0,0143 | 8,0000 | 1,0134 | 0,0877 |
| 36 | | 0,0143 | 8,0000 | 1,0134 | 0,0877 |
| 37 | | 0,0143 | 8,0000 | 1,0134 | 0,0877 |
| 38 | | 0,0143 | 8,0000 | 1,0134 | 0,0877 |
| 39 | | 0,0143 | 8,0000 | 1,0134 | 0,0877 |
| 40 | | 0,0143 | 8,0000 | 1,0134 | 0,0877 |
| 41 | | 0,0143 | 8,0000 | 1,0134 | 0,0877 |
| 42 | | 0,0143 | 8,0000 | 1,0134 | 0,0877 |
| 43 | | 0,0143 | 8,0000 | 1,0134 | 0,0877 |
| 44 | | 0,0143 | 8,0000 | 1,0134 | 0,0877 |
| 45 | | 0,0143 | 8,0000 | 1,0134 | 0,0877 |
| 46 | | 0,0143 | 8,0000 | 1,0134 | 0,0877 |
| 47 | | 0,0143 | 8,0000 | 1,0134 | 0,0877 |
| 48 | | 0,0143 | 8,0000 | 1,0134 | 0,0877 |
| 49 | | 0,0143 | 8,0000 | 1,0134 | 0,0877 |

| | 1 | 2 | 3 | 4 | 5 | 6 |
|---|---|---|---|---|---|---|
| 50 | | | | | | |
| 51 | | | | | | |
| 52 | | | | | | |
| 53 | | | | | | |
| 54 | | | | | | |
| 55 | | | | | | |
| 56 | | | | | | |
| 57 | | | | | | |
| 58 | | | | | | |
| 59 | | | | | | |
| 60 | | | | | | |
| 61 | | | | | | |
| 62 | | | | | | |
| 63 | | | | | | |
| 64 | | | | | | |
| 65 | | | | | | |
| 66 | | | | | | |
| 67 | | | | | | |
| 68 | | | | | | |
| 69 | | | | | | |
| 70 | | | | | | |
| 71 | | | | | | |
| 72 | | | | | | |
| 73 | | | | | | |
| 74 | | | | | | |
| 75 | | | | | | |
| 76 | | | | | | |
| 77 | | | | | | |
| 78 | | | | | | |
| 79 | | | | | | |
| 80 | | | | | | |
| 81 | | | | | | |
| 82 | | | | | | |
| 83 | | | | | | |
| 84 | | | | | | |
| 85 | | | | | | |
| 86 | | | | | | |
| 87 | | | | | | |
| 88 | | | | | | |
| 89 | | | | | | |
| 90 | | | | | | |
| 91 | | | | | | |
| 92 | | | | | | |
| 93 | | | | | | |
| 94 | | | | | | |
| 95 | | | | | | |
| 96 | | | | | | |
| 97 | | | | | | |
| 98 | | | | | | |

| | 7-x column | 8-y column | 9 | 10-SumSq | 11-Iterations | 12-a-Final |
|---|---|---|---|---|---|---|
| 50 | 1,3506 | 0,8365 | | 0,0411 | 7,0000 | 1,0972 |
| 51 | 1,3730 | 0,8327 | | 0,0411 | 7,0000 | 1,0972 |
| 52 | 1,3955 | 0,8289 | | 0,0411 | 10,0000 | 1,0972 |
| 53 | 1,4180 | 0,8252 | | 0,0411 | 7,0000 | 1,0972 |
| 54 | 1,4404 | 0,8215 | | 0,0411 | 7,0000 | 1,0972 |
| 55 | 1,4629 | 0,8178 | | 0,0411 | 8,0000 | 1,0972 |
| 56 | 1,4854 | 0,8141 | | 0,0411 | 10,0000 | 1,0972 |
| 57 | 1,5078 | 0,8105 | | 0,0411 | 8,0000 | 1,0972 |
| 58 | 1,5303 | 0,8068 | | 0,0411 | 12,0000 | 1,0972 |
| 59 | 1,5527 | 0,8032 | | 0,0411 | 8,0000 | 1,0972 |
| 60 | 1,5752 | 0,7996 | | 0,0411 | 10,0000 | 1,0972 |
| 61 | 1,5977 | 0,7960 | | 0,0411 | 10,0000 | 1,0972 |
| 62 | 1,6201 | 0,7924 | | 0,0411 | 7,0000 | 1,0972 |
| 63 | 1,6426 | 0,7888 | | 0,0411 | 10,0000 | 1,0972 |
| 64 | 1,6650 | 0,7853 | | 0,0411 | 8,0000 | 1,0972 |
| 65 | 1,6875 | 0,7817 | | 0,0411 | 10,0000 | 1,0972 |
| 66 | 1,7100 | 0,7782 | | 0,0411 | 10,0000 | 1,0972 |
| 67 | 1,7324 | 0,7747 | | 0,0411 | 10,0000 | 1,0972 |
| 68 | 1,7549 | 0,7712 | | 0,0411 | 8,0000 | 1,0972 |
| 69 | 1,7773 | 0,7677 | | 0,0411 | 7,0000 | 1,0972 |
| 70 | 1,7998 | 0,7643 | | 0,0411 | 7,0000 | 1,0972 |
| 71 | 1,8223 | 0,7608 | | 0,0411 | 8,0000 | 1,0972 |
| 72 | 1,8447 | 0,7574 | | 0,0411 | 7,0000 | 1,0972 |
| 73 | 1,8672 | 0,7540 | | 0,0411 | 8,0000 | 1,0972 |
| 74 | 1,8896 | 0,7506 | | 0,0411 | 10,0000 | 1,0972 |
| 75 | 1,9121 | 0,7472 | | 0,0411 | 7,0000 | 1,0972 |
| 76 | 1,9346 | 0,7439 | | 0,0411 | 7,0000 | 1,0972 |
| 77 | 1,9570 | 0,7405 | | 0,0411 | 7,0000 | 1,0972 |
| 78 | 1,9795 | 0,7372 | | 0,0411 | 8,0000 | 1,0972 |
| 79 | 2,0020 | 0,7339 | | 0,0411 | 7,0000 | 1,0972 |
| 80 | 2,0244 | 0,7306 | | 0,0411 | 8,0000 | 1,0972 |
| 81 | 2,0469 | 0,7273 | | 0,0411 | 8,0000 | 1,0972 |
| 82 | 2,0693 | 0,7240 | | 0,0411 | 8,0000 | 1,0972 |
| 83 | 2,0918 | 0,7207 | | 0,0411 | 7,0000 | 1,0972 |
| 84 | 2,1143 | 0,7175 | | 0,0411 | 10,0000 | 1,0972 |
| 85 | 2,1367 | 0,7143 | | 0,0411 | 7,0000 | 1,0972 |
| 86 | 2,1592 | 0,7110 | | 0,0411 | 7,0000 | 1,0972 |
| 87 | 2,1816 | 0,7078 | | 0,0411 | 7,0000 | 1,0972 |
| 88 | 2,2041 | 0,7047 | | 0,0411 | 7,0000 | 1,0972 |
| 89 | 2,2266 | 0,7015 | | 0,0411 | 8,0000 | 1,0972 |
| 90 | 2,2490 | 0,6983 | | 0,0411 | 7,0000 | 1,0972 |
| 91 | 2,2715 | 0,6952 | | 0,0411 | 7,0000 | 1,0972 |
| 92 | 2,2939 | 0,6921 | | 0,0411 | 8,0000 | 1,0972 |
| 93 | 2,3164 | 0,6889 | | 0,0411 | 7,0000 | 1,0972 |
| 94 | 2,3389 | 0,6858 | | 0,0411 | 8,0000 | 1,0972 |
| 95 | 2,3613 | 0,6827 | | 0,0411 | 10,0000 | 1,0972 |
| 96 | 2,3838 | 0,6797 | | 0,0411 | 10,0000 | 1,0972 |
| 97 | 2,4063 | 0,6766 | | 0,0411 | 8,0000 | 1,0972 |
| 98 | 2,4287 | 0,6736 | | 0,0411 | 22,0000 | 1,0972 |

| | 13-b-Final | 14-x column 1 | 15-y column 1 | 16 | 17-SumSq 1 | 18-Iterations 1 |
|---|---|---|---|---|---|---|
| 50 | 0,2009 | 1,3506 | 0,8321 | | 0,0352 | 7,0000 |
| 51 | 0,2009 | 1,3730 | 0,8285 | | 0,0352 | 7,0000 |
| 52 | 0,2009 | 1,3955 | 0,8250 | | 0,0352 | 10,0000 |
| 53 | 0,2009 | 1,4180 | 0,8215 | | 0,0352 | 7,0000 |
| 54 | 0,2009 | 1,4404 | 0,8180 | | 0,0352 | 7,0000 |
| 55 | 0,2009 | 1,4629 | 0,8145 | | 0,0352 | 7,0000 |
| 56 | 0,2009 | 1,4854 | 0,8111 | | 0,0352 | 7,0000 |
| 57 | 0,2009 | 1,5078 | 0,8076 | | 0,0352 | 7,0000 |
| 58 | 0,2009 | 1,5303 | 0,8042 | | 0,0352 | 10,0000 |
| 59 | 0,2009 | 1,5527 | 0,8008 | | 0,0352 | 7,0000 |
| 60 | 0,2009 | 1,5752 | 0,7974 | | 0,0352 | 7,0000 |
| 61 | 0,2009 | 1,5977 | 0,7940 | | 0,0352 | 10,0000 |
| 62 | 0,2009 | 1,6201 | 0,7906 | | 0,0352 | 7,0000 |
| 63 | 0,2009 | 1,6426 | 0,7872 | | 0,0352 | 7,0000 |
| 64 | 0,2009 | 1,6650 | 0,7839 | | 0,0352 | 7,0000 |
| 65 | 0,2009 | 1,6875 | 0,7806 | | 0,0352 | 10,0000 |
| 66 | 0,2009 | 1,7100 | 0,7772 | | 0,0352 | 7,0000 |
| 67 | 0,2009 | 1,7324 | 0,7739 | | 0,0352 | 10,0000 |
| 68 | 0,2009 | 1,7549 | 0,7706 | | 0,0352 | 10,0000 |
| 69 | 0,2009 | 1,7773 | 0,7674 | | 0,0352 | 7,0000 |
| 70 | 0,2009 | 1,7998 | 0,7641 | | 0,0352 | 7,0000 |
| 71 | 0,2009 | 1,8223 | 0,7608 | | 0,0352 | 7,0000 |
| 72 | 0,2009 | 1,8447 | 0,7576 | | 0,0352 | 7,0000 |
| 73 | 0,2009 | 1,8672 | 0,7544 | | 0,0352 | 10,0000 |
| 74 | 0,2009 | 1,8896 | 0,7512 | | 0,0352 | 7,0000 |
| 75 | 0,2009 | 1,9121 | 0,7480 | | 0,0352 | 10,0000 |
| 76 | 0,2009 | 1,9346 | 0,7448 | | 0,0352 | 7,0000 |
| 77 | 0,2009 | 1,9570 | 0,7416 | | 0,0352 | 12,0000 |
| 78 | 0,2009 | 1,9795 | 0,7385 | | 0,0352 | 7,0000 |
| 79 | 0,2009 | 2,0020 | 0,7353 | | 0,0352 | 7,0000 |
| 80 | 0,2009 | 2,0244 | 0,7322 | | 0,0352 | 10,0000 |
| 81 | 0,2009 | 2,0469 | 0,7291 | | 0,0352 | 7,0000 |
| 82 | 0,2009 | 2,0693 | 0,7260 | | 0,0352 | 7,0000 |
| 83 | 0,2009 | 2,0918 | 0,7229 | | 0,0352 | 7,0000 |
| 84 | 0,2009 | 2,1143 | 0,7198 | | 0,0352 | 7,0000 |
| 85 | 0,2009 | 2,1367 | 0,7168 | | 0,0352 | 7,0000 |
| 86 | 0,2009 | 2,1592 | 0,7137 | | 0,0352 | 7,0000 |
| 87 | 0,2009 | 2,1816 | 0,7107 | | 0,0352 | 7,0000 |
| 88 | 0,2009 | 2,2041 | 0,7077 | | 0,0352 | 7,0000 |
| 89 | 0,2009 | 2,2266 | 0,7047 | | 0,0352 | 7,0000 |
| 90 | 0,2009 | 2,2490 | 0,7017 | | 0,0352 | 7,0000 |
| 91 | 0,2009 | 2,2715 | 0,6987 | | 0,0352 | 10,0000 |
| 92 | 0,2009 | 2,2939 | 0,6957 | | 0,0352 | 10,0000 |
| 93 | 0,2009 | 2,3164 | 0,6928 | | 0,0352 | 7,0000 |
| 94 | 0,2009 | 2,3389 | 0,6898 | | 0,0352 | 7,0000 |
| 95 | 0,2009 | 2,3613 | 0,6869 | | 0,0352 | 7,0000 |
| 96 | 0,2009 | 2,3838 | 0,6840 | | 0,0352 | 7,0000 |
| 97 | 0,2009 | 2,4063 | 0,6810 | | 0,0352 | 7,0000 |
| 98 | 0,2009 | 2,4287 | 0,6781 | | 0,0352 | 8,0000 |

| | 19-a-Final 1 | 20-b-Final 1 | 21-x column 2 | 22-y column 2 | 23 | 24-SumSq 2 |
|---|---|---|---|---|---|---|
| 50 | 1,0751 | 0,1897 | 1,3506 | 0,8713 | | 0,0164 |
| 51 | 1,0751 | 0,1897 | 1,3730 | 0,8697 | | 0,0164 |
| 52 | 1,0751 | 0,1897 | 1,3955 | 0,8681 | | 0,0164 |
| 53 | 1,0751 | 0,1897 | 1,4180 | 0,8665 | | 0,0164 |
| 54 | 1,0751 | 0,1897 | 1,4404 | 0,8649 | | 0,0164 |
| 55 | 1,0751 | 0,1897 | 1,4629 | 0,8633 | | 0,0164 |
| 56 | 1,0751 | 0,1897 | 1,4854 | 0,8617 | | 0,0164 |
| 57 | 1,0751 | 0,1897 | 1,5078 | 0,8601 | | 0,0164 |
| 58 | 1,0751 | 0,1897 | 1,5303 | 0,8586 | | 0,0164 |
| 59 | 1,0751 | 0,1897 | 1,5527 | 0,8570 | | 0,0164 |
| 60 | 1,0751 | 0,1897 | 1,5752 | 0,8554 | | 0,0164 |
| 61 | 1,0751 | 0,1897 | 1,5977 | 0,8538 | | 0,0164 |
| 62 | 1,0751 | 0,1897 | 1,6201 | 0,8523 | | 0,0164 |
| 63 | 1,0751 | 0,1897 | 1,6426 | 0,8507 | | 0,0164 |
| 64 | 1,0751 | 0,1897 | 1,6650 | 0,8491 | | 0,0164 |
| 65 | 1,0751 | 0,1897 | 1,6875 | 0,8476 | | 0,0164 |
| 66 | 1,0751 | 0,1897 | 1,7100 | 0,8460 | | 0,0164 |
| 67 | 1,0751 | 0,1897 | 1,7324 | 0,8444 | | 0,0164 |
| 68 | 1,0751 | 0,1897 | 1,7549 | 0,8429 | | 0,0164 |
| 69 | 1,0751 | 0,1897 | 1,7773 | 0,8413 | | 0,0164 |
| 70 | 1,0751 | 0,1897 | 1,7998 | 0,8398 | | 0,0164 |
| 71 | 1,0751 | 0,1897 | 1,8223 | 0,8382 | | 0,0164 |
| 72 | 1,0751 | 0,1897 | 1,8447 | 0,8367 | | 0,0164 |
| 73 | 1,0751 | 0,1897 | 1,8672 | 0,8352 | | 0,0164 |
| 74 | 1,0751 | 0,1897 | 1,8896 | 0,8336 | | 0,0164 |
| 75 | 1,0751 | 0,1897 | 1,9121 | 0,8321 | | 0,0164 |
| 76 | 1,0751 | 0,1897 | 1,9346 | 0,8306 | | 0,0164 |
| 77 | 1,0751 | 0,1897 | 1,9570 | 0,8290 | | 0,0164 |
| 78 | 1,0751 | 0,1897 | 1,9795 | 0,8275 | | 0,0164 |
| 79 | 1,0751 | 0,1897 | 2,0020 | 0,8260 | | 0,0164 |
| 80 | 1,0751 | 0,1897 | 2,0244 | 0,8245 | | 0,0164 |
| 81 | 1,0751 | 0,1897 | 2,0469 | 0,8229 | | 0,0164 |
| 82 | 1,0751 | 0,1897 | 2,0693 | 0,8214 | | 0,0164 |
| 83 | 1,0751 | 0,1897 | 2,0918 | 0,8199 | | 0,0164 |
| 84 | 1,0751 | 0,1897 | 2,1143 | 0,8184 | | 0,0164 |
| 85 | 1,0751 | 0,1897 | 2,1367 | 0,8169 | | 0,0164 |
| 86 | 1,0751 | 0,1897 | 2,1592 | 0,8154 | | 0,0164 |
| 87 | 1,0751 | 0,1897 | 2,1816 | 0,8139 | | 0,0164 |
| 88 | 1,0751 | 0,1897 | 2,2041 | 0,8124 | | 0,0164 |
| 89 | 1,0751 | 0,1897 | 2,2266 | 0,8109 | | 0,0164 |
| 90 | 1,0751 | 0,1897 | 2,2490 | 0,8094 | | 0,0164 |
| 91 | 1,0751 | 0,1897 | 2,2715 | 0,8079 | | 0,0164 |
| 92 | 1,0751 | 0,1897 | 2,2939 | 0,8064 | | 0,0164 |
| 93 | 1,0751 | 0,1897 | 2,3164 | 0,8050 | | 0,0164 |
| 94 | 1,0751 | 0,1897 | 2,3389 | 0,8035 | | 0,0164 |
| 95 | 1,0751 | 0,1897 | 2,3613 | 0,8020 | | 0,0164 |
| 96 | 1,0751 | 0,1897 | 2,3838 | 0,8005 | | 0,0164 |
| 97 | 1,0751 | 0,1897 | 2,4063 | 0,7990 | | 0,0164 |
| 98 | 1,0751 | 0,1897 | 2,4287 | 0,7976 | | 0,0164 |

| | 25-Iterations 2 | 26-a-Final 2 | 27-b-Final 2 | 28-x column 3 | 29-y column 3 | 30 |
|---|---|---|---|---|---|---|
| 50 | 9,0000 | 0,9733 | 0,0820 | 1,3506 | 0,9217 | |
| 51 | 9,0000 | 0,9733 | 0,0820 | 1,3730 | 0,9198 | |
| 52 | 9,0000 | 0,9733 | 0,0820 | 1,3955 | 0,9178 | |
| 53 | 9,0000 | 0,9733 | 0,0820 | 1,4180 | 0,9159 | |
| 54 | 9,0000 | 0,9733 | 0,0820 | 1,4404 | 0,9140 | |
| 55 | 9,0000 | 0,9733 | 0,0820 | 1,4629 | 0,9120 | |
| 56 | 9,0000 | 0,9733 | 0,0820 | 1,4854 | 0,9101 | |
| 57 | 9,0000 | 0,9733 | 0,0820 | 1,5078 | 0,9082 | |
| 58 | 9,0000 | 0,9733 | 0,0820 | 1,5303 | 0,9062 | |
| 59 | 9,0000 | 0,9733 | 0,0820 | 1,5527 | 0,9043 | |
| 60 | 9,0000 | 0,9733 | 0,0820 | 1,5752 | 0,9024 | |
| 61 | 9,0000 | 0,9733 | 0,0820 | 1,5977 | 0,9005 | |
| 62 | 9,0000 | 0,9733 | 0,0820 | 1,6201 | 0,8986 | |
| 63 | 9,0000 | 0,9733 | 0,0820 | 1,6426 | 0,8967 | |
| 64 | 9,0000 | 0,9733 | 0,0820 | 1,6650 | 0,8948 | |
| 65 | 9,0000 | 0,9733 | 0,0820 | 1,6875 | 0,8929 | |
| 66 | 9,0000 | 0,9733 | 0,0820 | 1,7100 | 0,8910 | |
| 67 | 9,0000 | 0,9733 | 0,0820 | 1,7324 | 0,8891 | |
| 68 | 9,0000 | 0,9733 | 0,0820 | 1,7549 | 0,8872 | |
| 69 | 9,0000 | 0,9733 | 0,0820 | 1,7773 | 0,8853 | |
| 70 | 9,0000 | 0,9733 | 0,0820 | 1,7998 | 0,8834 | |
| 71 | 9,0000 | 0,9733 | 0,0820 | 1,8223 | 0,8816 | |
| 72 | 9,0000 | 0,9733 | 0,0820 | 1,8447 | 0,8797 | |
| 73 | 9,0000 | 0,9733 | 0,0820 | 1,8672 | 0,8778 | |
| 74 | 9,0000 | 0,9733 | 0,0820 | 1,8896 | 0,8760 | |
| 75 | 9,0000 | 0,9733 | 0,0820 | 1,9121 | 0,8741 | |
| 76 | 9,0000 | 0,9733 | 0,0820 | 1,9346 | 0,8723 | |
| 77 | 9,0000 | 0,9733 | 0,0820 | 1,9570 | 0,8704 | |
| 78 | 9,0000 | 0,9733 | 0,0820 | 1,9795 | 0,8686 | |
| 79 | 9,0000 | 0,9733 | 0,0820 | 2,0020 | 0,8667 | |
| 80 | 9,0000 | 0,9733 | 0,0820 | 2,0244 | 0,8649 | |
| 81 | 9,0000 | 0,9733 | 0,0820 | 2,0469 | 0,8631 | |
| 82 | 9,0000 | 0,9733 | 0,0820 | 2,0693 | 0,8612 | |
| 83 | 9,0000 | 0,9733 | 0,0820 | 2,0918 | 0,8594 | |
| 84 | 9,0000 | 0,9733 | 0,0820 | 2,1143 | 0,8576 | |
| 85 | 9,0000 | 0,9733 | 0,0820 | 2,1367 | 0,8558 | |
| 86 | 9,0000 | 0,9733 | 0,0820 | 2,1592 | 0,8540 | |
| 87 | 9,0000 | 0,9733 | 0,0820 | 2,1816 | 0,8522 | |
| 88 | 8,0000 | 0,9733 | 0,0820 | 2,2041 | 0,8503 | |
| 89 | 8,0000 | 0,9733 | 0,0820 | 2,2266 | 0,8485 | |
| 90 | 9,0000 | 0,9733 | 0,0820 | 2,2490 | 0,8467 | |
| 91 | 9,0000 | 0,9733 | 0,0820 | 2,2715 | 0,8450 | |
| 92 | 9,0000 | 0,9733 | 0,0820 | 2,2939 | 0,8432 | |
| 93 | 8,0000 | 0,9733 | 0,0820 | 2,3164 | 0,8414 | |
| 94 | 9,0000 | 0,9733 | 0,0820 | 2,3389 | 0,8396 | |
| 95 | 9,0000 | 0,9733 | 0,0820 | 2,3613 | 0,8378 | |
| 96 | 9,0000 | 0,9733 | 0,0820 | 2,3838 | 0,8360 | |
| 97 | 8,0000 | 0,9733 | 0,0820 | 2,4063 | 0,8343 | |
| 98 | 9,0000 | 0,9733 | 0,0820 | 2,4287 | 0,8325 | |

| | 31-SumSq 3 | 32-Iterations 3 | 33-a-Final 3 | 34-b-Final 3 | 35-x column 4 | 36-y column 4 |
|---|---|---|---|---|---|---|
| 50 | 0,0189 | 8,0000 | 1,0472 | 0,0945 | 1,3506 | 0,9002 |
| 51 | 0,0189 | 8,0000 | 1,0472 | 0,0945 | 1,3730 | 0,8985 |
| 52 | 0,0189 | 8,0000 | 1,0472 | 0,0945 | 1,3955 | 0,8967 |
| 53 | 0,0189 | 8,0000 | 1,0472 | 0,0945 | 1,4180 | 0,8949 |
| 54 | 0,0189 | 8,0000 | 1,0472 | 0,0945 | 1,4404 | 0,8932 |
| 55 | 0,0189 | 8,0000 | 1,0472 | 0,0945 | 1,4629 | 0,8914 |
| 56 | 0,0189 | 8,0000 | 1,0472 | 0,0945 | 1,4854 | 0,8897 |
| 57 | 0,0189 | 8,0000 | 1,0472 | 0,0945 | 1,5078 | 0,8879 |
| 58 | 0,0189 | 8,0000 | 1,0472 | 0,0945 | 1,5303 | 0,8862 |
| 59 | 0,0189 | 8,0000 | 1,0472 | 0,0945 | 1,5527 | 0,8844 |
| 60 | 0,0189 | 8,0000 | 1,0472 | 0,0945 | 1,5752 | 0,8827 |
| 61 | 0,0189 | 8,0000 | 1,0472 | 0,0945 | 1,5977 | 0,8810 |
| 62 | 0,0189 | 8,0000 | 1,0472 | 0,0945 | 1,6201 | 0,8792 |
| 63 | 0,0189 | 8,0000 | 1,0472 | 0,0945 | 1,6426 | 0,8775 |
| 64 | 0,0189 | 8,0000 | 1,0472 | 0,0945 | 1,6650 | 0,8758 |
| 65 | 0,0189 | 8,0000 | 1,0472 | 0,0945 | 1,6875 | 0,8740 |
| 66 | 0,0189 | 7,0000 | 1,0472 | 0,0945 | 1,7100 | 0,8723 |
| 67 | 0,0189 | 8,0000 | 1,0472 | 0,0945 | 1,7324 | 0,8706 |
| 68 | 0,0189 | 8,0000 | 1,0472 | 0,0945 | 1,7549 | 0,8689 |
| 69 | 0,0189 | 8,0000 | 1,0472 | 0,0945 | 1,7773 | 0,8672 |
| 70 | 0,0189 | 8,0000 | 1,0472 | 0,0945 | 1,7998 | 0,8655 |
| 71 | 0,0189 | 8,0000 | 1,0472 | 0,0945 | 1,8223 | 0,8638 |
| 72 | 0,0189 | 8,0000 | 1,0472 | 0,0945 | 1,8447 | 0,8621 |
| 73 | 0,0189 | 8,0000 | 1,0472 | 0,0945 | 1,8672 | 0,8604 |
| 74 | 0,0189 | 8,0000 | 1,0472 | 0,0945 | 1,8896 | 0,8587 |
| 75 | 0,0189 | 8,0000 | 1,0472 | 0,0945 | 1,9121 | 0,8570 |
| 76 | 0,0189 | 8,0000 | 1,0472 | 0,0945 | 1,9346 | 0,8553 |
| 77 | 0,0189 | 8,0000 | 1,0472 | 0,0945 | 1,9570 | 0,8536 |
| 78 | 0,0189 | 8,0000 | 1,0472 | 0,0945 | 1,9795 | 0,8519 |
| 79 | 0,0189 | 8,0000 | 1,0472 | 0,0945 | 2,0020 | 0,8503 |
| 80 | 0,0189 | 8,0000 | 1,0472 | 0,0945 | 2,0244 | 0,8486 |
| 81 | 0,0189 | 7,0000 | 1,0472 | 0,0945 | 2,0469 | 0,8469 |
| 82 | 0,0189 | 8,0000 | 1,0472 | 0,0945 | 2,0693 | 0,8453 |
| 83 | 0,0189 | 8,0000 | 1,0472 | 0,0945 | 2,0918 | 0,8436 |
| 84 | 0,0189 | 8,0000 | 1,0472 | 0,0945 | 2,1143 | 0,8419 |
| 85 | 0,0189 | 8,0000 | 1,0472 | 0,0945 | 2,1367 | 0,8403 |
| 86 | 0,0189 | 8,0000 | 1,0472 | 0,0945 | 2,1592 | 0,8386 |
| 87 | 0,0189 | 8,0000 | 1,0472 | 0,0945 | 2,1816 | 0,8370 |
| 88 | 0,0189 | 8,0000 | 1,0472 | 0,0945 | 2,2041 | 0,8353 |
| 89 | 0,0189 | 8,0000 | 1,0472 | 0,0945 | 2,2266 | 0,8337 |
| 90 | 0,0189 | 8,0000 | 1,0472 | 0,0945 | 2,2490 | 0,8320 |
| 91 | 0,0189 | 8,0000 | 1,0472 | 0,0945 | 2,2715 | 0,8304 |
| 92 | 0,0189 | 8,0000 | 1,0472 | 0,0945 | 2,2939 | 0,8288 |
| 93 | 0,0189 | 8,0000 | 1,0472 | 0,0945 | 2,3164 | 0,8271 |
| 94 | 0,0189 | 8,0000 | 1,0472 | 0,0945 | 2,3389 | 0,8255 |
| 95 | 0,0189 | 8,0000 | 1,0472 | 0,0945 | 2,3613 | 0,8239 |
| 96 | 0,0189 | 8,0000 | 1,0472 | 0,0945 | 2,3838 | 0,8223 |
| 97 | 0,0189 | 8,0000 | 1,0472 | 0,0945 | 2,4063 | 0,8206 |
| 98 | 0,0189 | 8,0000 | 1,0472 | 0,0945 | 2,4287 | 0,8190 |

| | 37 | 38-SumSq 4 | 39-Iterations 4 | 40-a-Final 4 | 41-b-Final 4 |
|---|---|---|---|---|---|
| 50 | | 0,0143 | 8,0000 | 1,0134 | 0,0877 |
| 51 | | 0,0143 | 8,0000 | 1,0134 | 0,0877 |
| 52 | | 0,0143 | 8,0000 | 1,0134 | 0,0877 |
| 53 | | 0,0143 | 8,0000 | 1,0134 | 0,0877 |
| 54 | | 0,0143 | 8,0000 | 1,0134 | 0,0877 |
| 55 | | 0,0143 | 8,0000 | 1,0134 | 0,0877 |
| 56 | | 0,0143 | 8,0000 | 1,0134 | 0,0877 |
| 57 | | 0,0143 | 8,0000 | 1,0134 | 0,0877 |
| 58 | | 0,0143 | 8,0000 | 1,0134 | 0,0877 |
| 59 | | 0,0143 | 8,0000 | 1,0134 | 0,0877 |
| 60 | | 0,0143 | 8,0000 | 1,0134 | 0,0877 |
| 61 | | 0,0143 | 8,0000 | 1,0134 | 0,0877 |
| 62 | | 0,0143 | 8,0000 | 1,0134 | 0,0877 |
| 63 | | 0,0143 | 8,0000 | 1,0134 | 0,0877 |
| 64 | | 0,0143 | 8,0000 | 1,0134 | 0,0877 |
| 65 | | 0,0143 | 8,0000 | 1,0134 | 0,0877 |
| 66 | | 0,0143 | 8,0000 | 1,0134 | 0,0877 |
| 67 | | 0,0143 | 8,0000 | 1,0134 | 0,0877 |
| 68 | | 0,0143 | 8,0000 | 1,0134 | 0,0877 |
| 69 | | 0,0143 | 8,0000 | 1,0134 | 0,0877 |
| 70 | | 0,0143 | 8,0000 | 1,0134 | 0,0877 |
| 71 | | 0,0143 | 8,0000 | 1,0134 | 0,0877 |
| 72 | | 0,0143 | 8,0000 | 1,0134 | 0,0877 |
| 73 | | 0,0143 | 8,0000 | 1,0134 | 0,0877 |
| 74 | | 0,0143 | 8,0000 | 1,0134 | 0,0877 |
| 75 | | 0,0143 | 8,0000 | 1,0134 | 0,0877 |
| 76 | | 0,0143 | 8,0000 | 1,0134 | 0,0877 |
| 77 | | 0,0143 | 8,0000 | 1,0134 | 0,0877 |
| 78 | | 0,0143 | 8,0000 | 1,0134 | 0,0877 |
| 79 | | 0,0143 | 8,0000 | 1,0134 | 0,0877 |
| 80 | | 0,0143 | 8,0000 | 1,0134 | 0,0877 |
| 81 | | 0,0143 | 8,0000 | 1,0134 | 0,0877 |
| 82 | | 0,0143 | 8,0000 | 1,0134 | 0,0877 |
| 83 | | 0,0143 | 8,0000 | 1,0134 | 0,0877 |
| 84 | | 0,0143 | 8,0000 | 1,0134 | 0,0877 |
| 85 | | 0,0143 | 8,0000 | 1,0134 | 0,0877 |
| 86 | | 0,0143 | 8,0000 | 1,0134 | 0,0877 |
| 87 | | 0,0143 | 8,0000 | 1,0134 | 0,0877 |
| 88 | | 0,0143 | 8,0000 | 1,0134 | 0,0877 |
| 89 | | 0,0143 | 8,0000 | 1,0134 | 0,0877 |
| 90 | | 0,0143 | 8,0000 | 1,0134 | 0,0877 |
| 91 | | 0,0143 | 8,0000 | 1,0134 | 0,0877 |
| 92 | | 0,0143 | 8,0000 | 1,0134 | 0,0877 |
| 93 | | 0,0143 | 8,0000 | 1,0134 | 0,0877 |
| 94 | | 0,0143 | 8,0000 | 1,0134 | 0,0877 |
| 95 | | 0,0143 | 8,0000 | 1,0134 | 0,0877 |
| 96 | | 0,0143 | 8,0000 | 1,0134 | 0,0877 |
| 97 | | 0,0143 | 8,0000 | 1,0134 | 0,0877 |
| 98 | | 0,0143 | 8,0000 | 1,0134 | 0,0877 |

| | 1 | 2 | 3 | 4 | 5 | 6 |
|---|---|---|---|---|---|---|
| 99 | | | | | | |
| 100 | | | | | | |
| 101 | | | | | | |
| 102 | | | | | | |
| 103 | | | | | | |
| 104 | | | | | | |
| 105 | | | | | | |
| 106 | | | | | | |
| 107 | | | | | | |
| 108 | | | | | | |
| 109 | | | | | | |
| 110 | | | | | | |
| 111 | | | | | | |
| 112 | | | | | | |
| 113 | | | | | | |
| 114 | | | | | | |
| 115 | | | | | | |
| 116 | | | | | | |
| 117 | | | | | | |
| 118 | | | | | | |
| 119 | | | | | | |
| 120 | | | | | | |
| 121 | | | | | | |
| 122 | | | | | | |
| 123 | | | | | | |
| 124 | | | | | | |
| 125 | | | | | | |
| 126 | | | | | | |
| 127 | | | | | | |
| 128 | | | | | | |
| 129 | | | | | | |
| 130 | | | | | | |
| 131 | | | | | | |
| 132 | | | | | | |
| 133 | | | | | | |
| 134 | | | | | | |
| 135 | | | | | | |
| 136 | | | | | | |
| 137 | | | | | | |
| 138 | | | | | | |
| 139 | | | | | | |
| 140 | | | | | | |
| 141 | | | | | | |
| 142 | | | | | | |
| 143 | | | | | | |
| 144 | | | | | | |
| 145 | | | | | | |
| 146 | | | | | | |
| 147 | | | | | | |

| | 7-x column | 8-y column | 9 | 10-SumSq | 11-Iterations | 12-a-Final |
|---|---|---|---|---|---|---|
| 99 | 2,4512 | 0,6705 | | 0,0411 | 7,0000 | 1,0972 |
| 100 | 2,4736 | 0,6675 | | 0,0411 | 10,0000 | 1,0972 |
| 101 | 2,4961 | 0,6645 | | 0,0411 | 8,0000 | 1,0972 |
| 102 | 2,5186 | 0,6615 | | 0,0411 | 7,0000 | 1,0972 |
| 103 | 2,5410 | 0,6585 | | 0,0411 | 10,0000 | 1,0972 |
| 104 | 2,5635 | 0,6556 | | 0,0411 | 7,0000 | 1,0972 |
| 105 | 2,5859 | 0,6526 | | 0,0411 | 7,0000 | 1,0972 |
| 106 | 2,6084 | 0,6497 | | 0,0411 | 10,0000 | 1,0972 |
| 107 | 2,6309 | 0,6468 | | 0,0411 | 7,0000 | 1,0972 |
| 108 | 2,6533 | 0,6439 | | 0,0411 | 7,0000 | 1,0972 |
| 109 | 2,6758 | 0,6410 | | 0,0411 | 7,0000 | 1,0972 |
| 110 | 2,6982 | 0,6381 | | 0,0411 | 10,0000 | 1,0972 |
| 111 | 2,7207 | 0,6352 | | 0,0411 | 10,0000 | 1,0972 |
| 112 | 2,7432 | 0,6323 | | 0,0411 | 10,0000 | 1,0972 |
| 113 | 2,7656 | 0,6295 | | 0,0411 | 7,0000 | 1,0972 |
| 114 | 2,7881 | 0,6267 | | 0,0411 | 7,0000 | 1,0972 |
| 115 | 2,8105 | 0,6238 | | 0,0411 | 8,0000 | 1,0972 |
| 116 | 2,8330 | 0,6210 | | 0,0411 | 7,0000 | 1,0972 |
| 117 | 2,8555 | 0,6182 | | 0,0411 | 10,0000 | 1,0972 |
| 118 | 2,8779 | 0,6154 | | 0,0411 | 10,0000 | 1,0972 |
| 119 | 2,9004 | 0,6127 | | 0,0411 | 7,0000 | 1,0972 |
| 120 | 2,9229 | 0,6099 | | 0,0411 | 7,0000 | 1,0972 |
| 121 | 2,9453 | 0,6072 | | 0,0411 | 7,0000 | 1,0972 |
| 122 | 2,9678 | 0,6044 | | 0,0411 | 7,0000 | 1,0972 |
| 123 | 2,9902 | 0,6017 | | 0,0411 | 7,0000 | 1,0972 |
| 124 | 3,0127 | 0,5990 | | 0,0411 | 7,0000 | 1,0972 |
| 125 | 3,0352 | 0,5963 | | 0,0411 | 10,0000 | 1,0972 |
| 126 | 3,0576 | 0,5936 | | 0,0411 | 8,0000 | 1,0972 |
| 127 | 3,0801 | 0,5910 | | 0,0411 | 8,0000 | 1,0972 |
| 128 | 3,1025 | 0,5883 | | 0,0411 | 7,0000 | 1,0972 |
| 129 | 3,1250 | 0,5856 | | 0,0411 | 10,0000 | 1,0972 |
| 130 | 3,1475 | 0,5830 | | 0,0411 | 7,0000 | 1,0972 |
| 131 | 3,1699 | 0,5804 | | 0,0411 | 7,0000 | 1,0972 |
| 132 | 3,1924 | 0,5778 | | 0,0411 | 7,0000 | 1,0972 |
| 133 | 3,2148 | 0,5752 | | 0,0411 | 8,0000 | 1,0972 |
| 134 | 3,2373 | 0,5726 | | 0,0411 | 10,0000 | 1,0972 |
| 135 | 3,2598 | 0,5700 | | 0,0411 | 7,0000 | 1,0972 |
| 136 | 3,2822 | 0,5674 | | 0,0411 | 7,0000 | 1,0972 |
| 137 | 3,3047 | 0,5649 | | 0,0411 | 33,0000 | 1,0972 |
| 138 | 3,3271 | 0,5623 | | 0,0411 | 8,0000 | 1,0972 |
| 139 | 3,3496 | 0,5598 | | 0,0411 | 7,0000 | 1,0972 |
| 140 | 3,3721 | 0,5573 | | 0,0411 | 7,0000 | 1,0972 |
| 141 | 3,3945 | 0,5548 | | 0,0411 | 8,0000 | 1,0972 |
| 142 | 3,4170 | 0,5523 | | 0,0411 | 7,0000 | 1,0972 |
| 143 | 3,4395 | 0,5498 | | 0,0411 | 8,0000 | 1,0972 |
| 144 | 3,4619 | 0,5473 | | 0,0411 | 7,0000 | 1,0972 |
| 145 | 3,4844 | 0,5449 | | 0,0411 | 7,0000 | 1,0972 |
| 146 | 3,5068 | 0,5424 | | 0,0411 | 7,0000 | 1,0972 |
| 147 | 3,5293 | 0,5400 | | 0,0411 | 8,0000 | 1,0972 |

| | 13-b-Final | 14-x column 1 | 15-y column 1 | 16 | 17-SumSq 1 | 18-Iterations 1 |
|---|---|---|---|---|---|---|
| 99 | 0,2009 | 2,4512 | 0,6753 | | 0,0352 | 7,0000 |
| 100 | 0,2009 | 2,4736 | 0,6724 | | 0,0352 | 19,0000 |
| 101 | 0,2009 | 2,4961 | 0,6695 | | 0,0352 | 10,0000 |
| 102 | 0,2009 | 2,5186 | 0,6667 | | 0,0352 | 10,0000 |
| 103 | 0,2009 | 2,5410 | 0,6638 | | 0,0352 | 7,0000 |
| 104 | 0,2009 | 2,5635 | 0,6610 | | 0,0352 | 7,0000 |
| 105 | 0,2009 | 2,5859 | 0,6582 | | 0,0352 | 7,0000 |
| 106 | 0,2009 | 2,6084 | 0,6554 | | 0,0352 | 10,0000 |
| 107 | 0,2009 | 2,6309 | 0,6526 | | 0,0352 | 7,0000 |
| 108 | 0,2009 | 2,6533 | 0,6499 | | 0,0352 | 7,0000 |
| 109 | 0,2009 | 2,6758 | 0,6471 | | 0,0352 | 10,0000 |
| 110 | 0,2009 | 2,6982 | 0,6443 | | 0,0352 | 7,0000 |
| 111 | 0,2009 | 2,7207 | 0,6416 | | 0,0352 | 7,0000 |
| 112 | 0,2009 | 2,7432 | 0,6389 | | 0,0352 | 10,0000 |
| 113 | 0,2009 | 2,7656 | 0,6362 | | 0,0352 | 7,0000 |
| 114 | 0,2009 | 2,7881 | 0,6334 | | 0,0352 | 8,0000 |
| 115 | 0,2009 | 2,8105 | 0,6308 | | 0,0352 | 7,0000 |
| 116 | 0,2009 | 2,8330 | 0,6281 | | 0,0352 | 7,0000 |
| 117 | 0,2009 | 2,8555 | 0,6254 | | 0,0352 | 7,0000 |
| 118 | 0,2009 | 2,8779 | 0,6227 | | 0,0352 | 7,0000 |
| 119 | 0,2009 | 2,9004 | 0,6201 | | 0,0352 | 10,0000 |
| 120 | 0,2009 | 2,9229 | 0,6175 | | 0,0352 | 7,0000 |
| 121 | 0,2009 | 2,9453 | 0,6148 | | 0,0352 | 7,0000 |
| 122 | 0,2009 | 2,9678 | 0,6122 | | 0,0352 | 7,0000 |
| 123 | 0,2009 | 2,9902 | 0,6096 | | 0,0352 | 7,0000 |
| 124 | 0,2009 | 3,0127 | 0,6070 | | 0,0352 | 7,0000 |
| 125 | 0,2009 | 3,0352 | 0,6044 | | 0,0352 | 7,0000 |
| 126 | 0,2009 | 3,0576 | 0,6019 | | 0,0352 | 7,0000 |
| 127 | 0,2009 | 3,0801 | 0,5993 | | 0,0352 | 10,0000 |
| 128 | 0,2009 | 3,1025 | 0,5968 | | 0,0352 | 7,0000 |
| 129 | 0,2009 | 3,1250 | 0,5942 | | 0,0352 | 7,0000 |
| 130 | 0,2009 | 3,1475 | 0,5917 | | 0,0352 | 7,0000 |
| 131 | 0,2009 | 3,1699 | 0,5892 | | 0,0352 | 7,0000 |
| 132 | 0,2009 | 3,1924 | 0,5867 | | 0,0352 | 8,0000 |
| 133 | 0,2009 | 3,2148 | 0,5842 | | 0,0352 | 7,0000 |
| 134 | 0,2009 | 3,2373 | 0,5817 | | 0,0352 | 7,0000 |
| 135 | 0,2009 | 3,2598 | 0,5792 | | 0,0352 | 7,0000 |
| 136 | 0,2009 | 3,2822 | 0,5768 | | 0,0352 | 7,0000 |
| 137 | 0,2009 | 3,3047 | 0,5743 | | 0,0352 | 10,0000 |
| 138 | 0,2009 | 3,3271 | 0,5719 | | 0,0352 | 7,0000 |
| 139 | 0,2009 | 3,3496 | 0,5694 | | 0,0352 | 7,0000 |
| 140 | 0,2009 | 3,3721 | 0,5670 | | 0,0352 | 7,0000 |
| 141 | 0,2009 | 3,3945 | 0,5646 | | 0,0352 | 8,0000 |
| 142 | 0,2009 | 3,4170 | 0,5622 | | 0,0352 | 8,0000 |
| 143 | 0,2009 | 3,4395 | 0,5598 | | 0,0352 | 7,0000 |
| 144 | 0,2009 | 3,4619 | 0,5574 | | 0,0352 | 8,0000 |
| 145 | 0,2009 | 3,4844 | 0,5551 | | 0,0352 | 7,0000 |
| 146 | 0,2009 | 3,5068 | 0,5527 | | 0,0352 | 15,0000 |
| 147 | 0,2009 | 3,5293 | 0,5503 | | 0,0352 | 7,0000 |

| | 19-a-Final 1 | 20-b-Final 1 | 21-x column 2 | 22-y column 2 | 23 | 24-SumSq 2 |
|---|---|---|---|---|---|---|
| 99 | 1,0751 | 0,1897 | 2,4512 | 0,7961 | | 0,0164 |
| 100 | 1,0751 | 0,1897 | 2,4736 | 0,7946 | | 0,0164 |
| 101 | 1,0751 | 0,1897 | 2,4961 | 0,7932 | | 0,0164 |
| 102 | 1,0751 | 0,1897 | 2,5186 | 0,7917 | | 0,0164 |
| 103 | 1,0751 | 0,1897 | 2,5410 | 0,7903 | | 0,0164 |
| 104 | 1,0751 | 0,1897 | 2,5635 | 0,7888 | | 0,0164 |
| 105 | 1,0751 | 0,1897 | 2,5859 | 0,7874 | | 0,0164 |
| 106 | 1,0751 | 0,1897 | 2,6084 | 0,7859 | | 0,0164 |
| 107 | 1,0751 | 0,1897 | 2,6309 | 0,7845 | | 0,0164 |
| 108 | 1,0751 | 0,1897 | 2,6533 | 0,7830 | | 0,0164 |
| 109 | 1,0751 | 0,1897 | 2,6758 | 0,7816 | | 0,0164 |
| 110 | 1,0751 | 0,1897 | 2,6982 | 0,7801 | | 0,0164 |
| 111 | 1,0751 | 0,1897 | 2,7207 | 0,7787 | | 0,0164 |
| 112 | 1,0751 | 0,1897 | 2,7432 | 0,7773 | | 0,0164 |
| 113 | 1,0751 | 0,1897 | 2,7656 | 0,7758 | | 0,0164 |
| 114 | 1,0751 | 0,1897 | 2,7881 | 0,7744 | | 0,0164 |
| 115 | 1,0751 | 0,1897 | 2,8105 | 0,7730 | | 0,0164 |
| 116 | 1,0751 | 0,1897 | 2,8330 | 0,7716 | | 0,0164 |
| 117 | 1,0751 | 0,1897 | 2,8555 | 0,7702 | | 0,0164 |
| 118 | 1,0751 | 0,1897 | 2,8779 | 0,7687 | | 0,0164 |
| 119 | 1,0751 | 0,1897 | 2,9004 | 0,7673 | | 0,0164 |
| 120 | 1,0751 | 0,1897 | 2,9229 | 0,7659 | | 0,0164 |
| 121 | 1,0751 | 0,1897 | 2,9453 | 0,7645 | | 0,0164 |
| 122 | 1,0751 | 0,1897 | 2,9678 | 0,7631 | | 0,0164 |
| 123 | 1,0751 | 0,1897 | 2,9902 | 0,7617 | | 0,0164 |
| 124 | 1,0751 | 0,1897 | 3,0127 | 0,7603 | | 0,0164 |
| 125 | 1,0751 | 0,1897 | 3,0352 | 0,7589 | | 0,0164 |
| 126 | 1,0751 | 0,1897 | 3,0576 | 0,7575 | | 0,0164 |
| 127 | 1,0751 | 0,1897 | 3,0801 | 0,7561 | | 0,0164 |
| 128 | 1,0751 | 0,1897 | 3,1025 | 0,7547 | | 0,0164 |
| 129 | 1,0751 | 0,1897 | 3,1250 | 0,7533 | | 0,0164 |
| 130 | 1,0751 | 0,1897 | 3,1475 | 0,7519 | | 0,0164 |
| 131 | 1,0751 | 0,1897 | 3,1699 | 0,7505 | | 0,0164 |
| 132 | 1,0751 | 0,1897 | 3,1924 | 0,7492 | | 0,0164 |
| 133 | 1,0751 | 0,1897 | 3,2148 | 0,7478 | | 0,0164 |
| 134 | 1,0751 | 0,1897 | 3,2373 | 0,7464 | | 0,0164 |
| 135 | 1,0751 | 0,1897 | 3,2598 | 0,7450 | | 0,0164 |
| 136 | 1,0751 | 0,1897 | 3,2822 | 0,7437 | | 0,0164 |
| 137 | 1,0751 | 0,1897 | 3,3047 | 0,7423 | | 0,0164 |
| 138 | 1,0751 | 0,1897 | 3,3271 | 0,7409 | | 0,0164 |
| 139 | 1,0751 | 0,1897 | 3,3496 | 0,7396 | | 0,0164 |
| 140 | 1,0751 | 0,1897 | 3,3721 | 0,7382 | | 0,0164 |
| 141 | 1,0751 | 0,1897 | 3,3945 | 0,7369 | | 0,0164 |
| 142 | 1,0751 | 0,1897 | 3,4170 | 0,7355 | | 0,0164 |
| 143 | 1,0751 | 0,1897 | 3,4395 | 0,7341 | | 0,0164 |
| 144 | 1,0751 | 0,1897 | 3,4619 | 0,7328 | | 0,0164 |
| 145 | 1,0751 | 0,1897 | 3,4844 | 0,7314 | | 0,0164 |
| 146 | 1,0751 | 0,1897 | 3,5068 | 0,7301 | | 0,0164 |
| 147 | 1,0751 | 0,1897 | 3,5293 | 0,7288 | | 0,0164 |

| | 25-Iterations 2 | 26-a-Final 2 | 27-b-Final 2 | 28-x column 3 | 29-y column 3 | 30 |
|---|---|---|---|---|---|---|
| 99 | 8,0000 | 0,9733 | 0,0820 | 2,4512 | 0,8307 | |
| 100 | 9,0000 | 0,9733 | 0,0820 | 2,4736 | 0,8290 | |
| 101 | 9,0000 | 0,9733 | 0,0820 | 2,4961 | 0,8272 | |
| 102 | 9,0000 | 0,9733 | 0,0820 | 2,5186 | 0,8255 | |
| 103 | 9,0000 | 0,9733 | 0,0820 | 2,5410 | 0,8237 | |
| 104 | 9,0000 | 0,9733 | 0,0820 | 2,5635 | 0,8220 | |
| 105 | 9,0000 | 0,9733 | 0,0820 | 2,5859 | 0,8202 | |
| 106 | 9,0000 | 0,9733 | 0,0820 | 2,6084 | 0,8185 | |
| 107 | 9,0000 | 0,9733 | 0,0820 | 2,6309 | 0,8168 | |
| 108 | 9,0000 | 0,9733 | 0,0820 | 2,6533 | 0,8150 | |
| 109 | 9,0000 | 0,9733 | 0,0820 | 2,6758 | 0,8133 | |
| 110 | 8,0000 | 0,9733 | 0,0820 | 2,6982 | 0,8116 | |
| 111 | 8,0000 | 0,9733 | 0,0820 | 2,7207 | 0,8099 | |
| 112 | 9,0000 | 0,9733 | 0,0820 | 2,7432 | 0,8081 | |
| 113 | 9,0000 | 0,9733 | 0,0820 | 2,7656 | 0,8064 | |
| 114 | 9,0000 | 0,9733 | 0,0820 | 2,7881 | 0,8047 | |
| 115 | 9,0000 | 0,9733 | 0,0820 | 2,8105 | 0,8030 | |
| 116 | 9,0000 | 0,9733 | 0,0820 | 2,8330 | 0,8013 | |
| 117 | 9,0000 | 0,9733 | 0,0820 | 2,8555 | 0,7996 | |
| 118 | 9,0000 | 0,9733 | 0,0820 | 2,8779 | 0,7979 | |
| 119 | 8,0000 | 0,9733 | 0,0820 | 2,9004 | 0,7962 | |
| 120 | 9,0000 | 0,9733 | 0,0820 | 2,9229 | 0,7945 | |
| 121 | 8,0000 | 0,9733 | 0,0820 | 2,9453 | 0,7929 | |
| 122 | 8,0000 | 0,9733 | 0,0820 | 2,9678 | 0,7912 | |
| 123 | 8,0000 | 0,9733 | 0,0820 | 2,9902 | 0,7895 | |
| 124 | 8,0000 | 0,9733 | 0,0820 | 3,0127 | 0,7878 | |
| 125 | 8,0000 | 0,9733 | 0,0820 | 3,0352 | 0,7862 | |
| 126 | 8,0000 | 0,9733 | 0,0820 | 3,0576 | 0,7845 | |
| 127 | 8,0000 | 0,9733 | 0,0820 | 3,0801 | 0,7828 | |
| 128 | 8,0000 | 0,9733 | 0,0820 | 3,1025 | 0,7812 | |
| 129 | 8,0000 | 0,9733 | 0,0820 | 3,1250 | 0,7795 | |
| 130 | 8,0000 | 0,9733 | 0,0820 | 3,1475 | 0,7779 | |
| 131 | 8,0000 | 0,9733 | 0,0820 | 3,1699 | 0,7762 | |
| 132 | 8,0000 | 0,9733 | 0,0820 | 3,1924 | 0,7746 | |
| 133 | 8,0000 | 0,9733 | 0,0820 | 3,2148 | 0,7729 | |
| 134 | 8,0000 | 0,9733 | 0,0820 | 3,2373 | 0,7713 | |
| 135 | 8,0000 | 0,9733 | 0,0820 | 3,2598 | 0,7696 | |
| 136 | 8,0000 | 0,9733 | 0,0820 | 3,2822 | 0,7680 | |
| 137 | 8,0000 | 0,9733 | 0,0820 | 3,3047 | 0,7664 | |
| 138 | 8,0000 | 0,9733 | 0,0820 | 3,3271 | 0,7648 | |
| 139 | 8,0000 | 0,9733 | 0,0820 | 3,3496 | 0,7631 | |
| 140 | 8,0000 | 0,9733 | 0,0820 | 3,3721 | 0,7615 | |
| 141 | 8,0000 | 0,9733 | 0,0820 | 3,3945 | 0,7599 | |
| 142 | 8,0000 | 0,9733 | 0,0820 | 3,4170 | 0,7583 | |
| 143 | 7,0000 | 0,9733 | 0,0820 | 3,4395 | 0,7567 | |
| 144 | 8,0000 | 0,9733 | 0,0820 | 3,4619 | 0,7551 | |
| 145 | 8,0000 | 0,9733 | 0,0820 | 3,4844 | 0,7535 | |
| 146 | 8,0000 | 0,9733 | 0,0820 | 3,5068 | 0,7519 | |
| 147 | 8,0000 | 0,9733 | 0,0820 | 3,5293 | 0,7503 | |

| | 31-SumSq 3 | 32-Iterations 3 | 33-a-Final 3 | 34-b-Final 3 | 35-x column 4 | 36-y column 4 |
|---|---|---|---|---|---|---|
| 99 | 0,0189 | 8,0000 | 1,0472 | 0,0945 | 2,4512 | 0,8174 |
| 100 | 0,0189 | 8,0000 | 1,0472 | 0,0945 | 2,4736 | 0,8158 |
| 101 | 0,0189 | 8,0000 | 1,0472 | 0,0945 | 2,4961 | 0,8142 |
| 102 | 0,0189 | 8,0000 | 1,0472 | 0,0945 | 2,5186 | 0,8126 |
| 103 | 0,0189 | 8,0000 | 1,0472 | 0,0945 | 2,5410 | 0,8110 |
| 104 | 0,0189 | 8,0000 | 1,0472 | 0,0945 | 2,5635 | 0,8094 |
| 105 | 0,0189 | 8,0000 | 1,0472 | 0,0945 | 2,5859 | 0,8078 |
| 106 | 0,0189 | 8,0000 | 1,0472 | 0,0945 | 2,6084 | 0,8062 |
| 107 | 0,0189 | 8,0000 | 1,0472 | 0,0945 | 2,6309 | 0,8046 |
| 108 | 0,0189 | 8,0000 | 1,0472 | 0,0945 | 2,6533 | 0,8031 |
| 109 | 0,0189 | 8,0000 | 1,0472 | 0,0945 | 2,6758 | 0,8015 |
| 110 | 0,0189 | 8,0000 | 1,0472 | 0,0945 | 2,6982 | 0,7999 |
| 111 | 0,0189 | 8,0000 | 1,0472 | 0,0945 | 2,7207 | 0,7983 |
| 112 | 0,0189 | 8,0000 | 1,0472 | 0,0945 | 2,7432 | 0,7968 |
| 113 | 0,0189 | 8,0000 | 1,0472 | 0,0945 | 2,7656 | 0,7952 |
| 114 | 0,0189 | 8,0000 | 1,0472 | 0,0945 | 2,7881 | 0,7936 |
| 115 | 0,0189 | 8,0000 | 1,0472 | 0,0945 | 2,8105 | 0,7921 |
| 116 | 0,0189 | 8,0000 | 1,0472 | 0,0945 | 2,8330 | 0,7905 |
| 117 | 0,0189 | 8,0000 | 1,0472 | 0,0945 | 2,8555 | 0,7889 |
| 118 | 0,0189 | 8,0000 | 1,0472 | 0,0945 | 2,8779 | 0,7874 |
| 119 | 0,0189 | 8,0000 | 1,0472 | 0,0945 | 2,9004 | 0,7858 |
| 120 | 0,0189 | 8,0000 | 1,0472 | 0,0945 | 2,9229 | 0,7843 |
| 121 | 0,0189 | 8,0000 | 1,0472 | 0,0945 | 2,9453 | 0,7828 |
| 122 | 0,0189 | 8,0000 | 1,0472 | 0,0945 | 2,9678 | 0,7812 |
| 123 | 0,0189 | 8,0000 | 1,0472 | 0,0945 | 2,9902 | 0,7797 |
| 124 | 0,0189 | 8,0000 | 1,0472 | 0,0945 | 3,0127 | 0,7781 |
| 125 | 0,0189 | 8,0000 | 1,0472 | 0,0945 | 3,0352 | 0,7766 |
| 126 | 0,0189 | 8,0000 | 1,0472 | 0,0945 | 3,0576 | 0,7751 |
| 127 | 0,0189 | 8,0000 | 1,0472 | 0,0945 | 3,0801 | 0,7736 |
| 128 | 0,0189 | 6,0000 | 1,0472 | 0,0945 | 3,1025 | 0,7720 |
| 129 | 0,0189 | 7,0000 | 1,0472 | 0,0945 | 3,1250 | 0,7705 |
| 130 | 0,0189 | 7,0000 | 1,0472 | 0,0945 | 3,1475 | 0,7690 |
| 131 | 0,0189 | 8,0000 | 1,0472 | 0,0945 | 3,1699 | 0,7675 |
| 132 | 0,0189 | 8,0000 | 1,0472 | 0,0945 | 3,1924 | 0,7660 |
| 133 | 0,0189 | 8,0000 | 1,0472 | 0,0945 | 3,2148 | 0,7645 |
| 134 | 0,0189 | 8,0000 | 1,0472 | 0,0945 | 3,2373 | 0,7630 |
| 135 | 0,0189 | 8,0000 | 1,0472 | 0,0945 | 3,2598 | 0,7615 |
| 136 | 0,0189 | 8,0000 | 1,0472 | 0,0945 | 3,2822 | 0,7600 |
| 137 | 0,0189 | 7,0000 | 1,0472 | 0,0945 | 3,3047 | 0,7585 |
| 138 | 0,0189 | 8,0000 | 1,0472 | 0,0945 | 3,3271 | 0,7570 |
| 139 | 0,0189 | 8,0000 | 1,0472 | 0,0945 | 3,3496 | 0,7555 |
| 140 | 0,0189 | 8,0000 | 1,0472 | 0,0945 | 3,3721 | 0,7540 |
| 141 | 0,0189 | 8,0000 | 1,0472 | 0,0945 | 3,3945 | 0,7525 |
| 142 | 0,0189 | 8,0000 | 1,0472 | 0,0945 | 3,4170 | 0,7510 |
| 143 | 0,0189 | 8,0000 | 1,0472 | 0,0945 | 3,4395 | 0,7496 |
| 144 | 0,0189 | 8,0000 | 1,0472 | 0,0945 | 3,4619 | 0,7481 |
| 145 | 0,0189 | 8,0000 | 1,0472 | 0,0945 | 3,4844 | 0,7466 |
| 146 | 0,0189 | 8,0000 | 1,0472 | 0,0945 | 3,5068 | 0,7451 |
| 147 | 0,0189 | 7,0000 | 1,0472 | 0,0945 | 3,5293 | 0,7437 |

| | 37 | 38-SumSq 4 | 39-Iterations 4 | 40-a-Final 4 | 41-b-Final 4 |
|---|---|---|---|---|---|
| 99 | | 0,0143 | 8,0000 | 1,0134 | 0,0877 |
| 100 | | 0,0143 | 8,0000 | 1,0134 | 0,0877 |
| 101 | | 0,0143 | 8,0000 | 1,0134 | 0,0877 |
| 102 | | 0,0143 | 8,0000 | 1,0134 | 0,0877 |
| 103 | | 0,0143 | 8,0000 | 1,0134 | 0,0877 |
| 104 | | 0,0143 | 8,0000 | 1,0134 | 0,0877 |
| 105 | | 0,0143 | 8,0000 | 1,0134 | 0,0877 |
| 106 | | 0,0143 | 8,0000 | 1,0134 | 0,0877 |
| 107 | | 0,0143 | 8,0000 | 1,0134 | 0,0877 |
| 108 | | 0,0143 | 8,0000 | 1,0134 | 0,0877 |
| 109 | | 0,0143 | 8,0000 | 1,0134 | 0,0877 |
| 110 | | 0,0143 | 8,0000 | 1,0134 | 0,0877 |
| 111 | | 0,0143 | 8,0000 | 1,0134 | 0,0877 |
| 112 | | 0,0143 | 8,0000 | 1,0134 | 0,0877 |
| 113 | | 0,0143 | 8,0000 | 1,0134 | 0,0877 |
| 114 | | 0,0143 | 8,0000 | 1,0134 | 0,0877 |
| 115 | | 0,0143 | 8,0000 | 1,0134 | 0,0877 |
| 116 | | 0,0143 | 8,0000 | 1,0134 | 0,0877 |
| 117 | | 0,0143 | 8,0000 | 1,0134 | 0,0877 |
| 118 | | 0,0143 | 8,0000 | 1,0134 | 0,0877 |
| 119 | | 0,0143 | 8,0000 | 1,0134 | 0,0877 |
| 120 | | 0,0143 | 8,0000 | 1,0134 | 0,0877 |
| 121 | | 0,0143 | 8,0000 | 1,0134 | 0,0877 |
| 122 | | 0,0143 | 8,0000 | 1,0134 | 0,0877 |
| 123 | | 0,0143 | 8,0000 | 1,0134 | 0,0877 |
| 124 | | 0,0143 | 8,0000 | 1,0134 | 0,0877 |
| 125 | | 0,0143 | 8,0000 | 1,0134 | 0,0877 |
| 126 | | 0,0143 | 8,0000 | 1,0134 | 0,0877 |
| 127 | | 0,0143 | 8,0000 | 1,0134 | 0,0877 |
| 128 | | 0,0143 | 8,0000 | 1,0134 | 0,0877 |
| 129 | | 0,0143 | 7,0000 | 1,0134 | 0,0877 |
| 130 | | 0,0143 | 8,0000 | 1,0134 | 0,0877 |
| 131 | | 0,0143 | 8,0000 | 1,0134 | 0,0877 |
| 132 | | 0,0143 | 8,0000 | 1,0134 | 0,0877 |
| 133 | | 0,0143 | 8,0000 | 1,0134 | 0,0877 |
| 134 | | 0,0143 | 8,0000 | 1,0134 | 0,0877 |
| 135 | | 0,0143 | 8,0000 | 1,0134 | 0,0877 |
| 136 | | 0,0143 | 8,0000 | 1,0134 | 0,0877 |
| 137 | | 0,0143 | 8,0000 | 1,0134 | 0,0877 |
| 138 | | 0,0143 | 8,0000 | 1,0134 | 0,0877 |
| 139 | | 0,0143 | 8,0000 | 1,0134 | 0,0877 |
| 140 | | 0,0143 | 8,0000 | 1,0134 | 0,0877 |
| 141 | | 0,0143 | 8,0000 | 1,0134 | 0,0877 |
| 142 | | 0,0143 | 8,0000 | 1,0134 | 0,0877 |
| 143 | | 0,0143 | 8,0000 | 1,0134 | 0,0877 |
| 144 | | 0,0143 | 8,0000 | 1,0134 | 0,0877 |
| 145 | | 0,0143 | 8,0000 | 1,0134 | 0,0877 |
| 146 | | 0,0143 | 8,0000 | 1,0134 | 0,0877 |
| 147 | | 0,0143 | 8,0000 | 1,0134 | 0,0877 |

| | 1 | 2 | 3 | 4 | 5 | 6 |
|---|---|---|---|---|---|---|
| 148 | | | | | | |
| 149 | | | | | | |
| 150 | | | | | | |
| 151 | | | | | | |
| 152 | | | | | | |
| 153 | | | | | | |
| 154 | | | | | | |
| 155 | | | | | | |
| 156 | | | | | | |
| 157 | | | | | | |
| 158 | | | | | | |
| 159 | | | | | | |
| 160 | | | | | | |
| 161 | | | | | | |
| 162 | | | | | | |
| 163 | | | | | | |
| 164 | | | | | | |
| 165 | | | | | | |
| 166 | | | | | | |
| 167 | | | | | | |
| 168 | | | | | | |
| 169 | | | | | | |
| 170 | | | | | | |
| 171 | | | | | | |
| 172 | | | | | | |
| 173 | | | | | | |
| 174 | | | | | | |
| 175 | | | | | | |
| 176 | | | | | | |
| 177 | | | | | | |
| 178 | | | | | | |
| 179 | | | | | | |
| 180 | | | | | | |
| 181 | | | | | | |
| 182 | | | | | | |
| 183 | | | | | | |
| 184 | | | | | | |
| 185 | | | | | | |
| 186 | | | | | | |
| 187 | | | | | | |
| 188 | | | | | | |
| 189 | | | | | | |
| 190 | | | | | | |
| 191 | | | | | | |
| 192 | | | | | | |
| 193 | | | | | | |
| 194 | | | | | | |
| 195 | | | | | | |
| 196 | | | | | | |

| | 7-x column | 8-y column | 9 | 10-SumSq | 11-Iterations | 12-a-Final |
|---|---|---|---|---|---|---|
| 148 | 3,5518 | 0,5375 | | 0,0411 | 8,0000 | 1,0972 |
| 149 | 3,5742 | 0,5351 | | 0,0411 | 8,0000 | 1,0972 |
| 150 | 3,5967 | 0,5327 | | 0,0411 | 7,0000 | 1,0972 |
| 151 | 3,6191 | 0,5303 | | 0,0411 | 8,0000 | 1,0972 |
| 152 | 3,6416 | 0,5279 | | 0,0411 | 7,0000 | 1,0972 |
| 153 | 3,6641 | 0,5255 | | 0,0411 | 7,0000 | 1,0972 |
| 154 | 3,6865 | 0,5232 | | 0,0411 | 10,0000 | 1,0972 |
| 155 | 3,7090 | 0,5208 | | 0,0411 | 7,0000 | 1,0972 |
| 156 | 3,7314 | 0,5185 | | 0,0411 | 7,0000 | 1,0972 |
| 157 | 3,7539 | 0,5161 | | 0,0411 | 7,0000 | 1,0972 |
| 158 | 3,7764 | 0,5138 | | 0,0411 | 7,0000 | 1,0972 |
| 159 | 3,7988 | 0,5115 | | 0,0411 | 7,0000 | 1,0972 |
| 160 | 3,8213 | 0,5092 | | 0,0411 | 7,0000 | 1,0972 |
| 161 | 3,8438 | 0,5069 | | 0,0411 | 7,0000 | 1,0972 |
| 162 | 3,8662 | 0,5046 | | 0,0411 | 7,0000 | 1,0972 |
| 163 | 3,8887 | 0,5023 | | 0,0411 | 7,0000 | 1,0972 |
| 164 | 3,9111 | 0,5001 | | 0,0411 | 7,0000 | 1,0972 |
| 165 | 3,9336 | 0,4978 | | 0,0411 | 7,0000 | 1,0972 |
| 166 | 3,9561 | 0,4956 | | 0,0411 | 7,0000 | 1,0972 |
| 167 | 3,9785 | 0,4934 | | 0,0411 | 7,0000 | 1,0972 |
| 168 | 4,0010 | 0,4911 | | 0,0411 | 7,0000 | 1,0972 |
| 169 | 4,0234 | 0,4889 | | 0,0411 | 7,0000 | 1,0972 |
| 170 | 4,0459 | 0,4867 | | 0,0411 | 7,0000 | 1,0972 |
| 171 | 4,0684 | 0,4845 | | 0,0411 | 7,0000 | 1,0972 |
| 172 | 4,0908 | 0,4824 | | 0,0411 | 7,0000 | 1,0972 |
| 173 | 4,1133 | 0,4802 | | 0,0411 | 7,0000 | 1,0972 |
| 174 | 4,1357 | 0,4780 | | 0,0411 | 7,0000 | 1,0972 |
| 175 | 4,1582 | 0,4759 | | 0,0411 | 7,0000 | 1,0972 |
| 176 | 4,1807 | 0,4737 | | 0,0411 | 7,0000 | 1,0972 |
| 177 | 4,2031 | 0,4716 | | 0,0411 | 7,0000 | 1,0972 |
| 178 | 4,2256 | 0,4695 | | 0,0411 | 7,0000 | 1,0972 |
| 179 | 4,2480 | 0,4674 | | 0,0411 | 7,0000 | 1,0972 |
| 180 | 4,2705 | 0,4653 | | 0,0411 | 7,0000 | 1,0972 |
| 181 | 4,2930 | 0,4632 | | 0,0411 | 7,0000 | 1,0972 |
| 182 | 4,3154 | 0,4611 | | 0,0411 | 7,0000 | 1,0972 |
| 183 | 4,3379 | 0,4590 | | 0,0411 | 7,0000 | 1,0972 |
| 184 | 4,3604 | 0,4569 | | 0,0411 | 7,0000 | 1,0972 |
| 185 | 4,3828 | 0,4549 | | 0,0411 | 7,0000 | 1,0972 |
| 186 | 4,4053 | 0,4528 | | 0,0411 | 7,0000 | 1,0972 |
| 187 | 4,4277 | 0,4508 | | 0,0411 | 7,0000 | 1,0972 |
| 188 | 4,4502 | 0,4488 | | 0,0411 | 7,0000 | 1,0972 |
| 189 | 4,4727 | 0,4467 | | 0,0411 | 7,0000 | 1,0972 |
| 190 | 4,4951 | 0,4447 | | 0,0411 | 7,0000 | 1,0972 |
| 191 | 4,5176 | 0,4427 | | 0,0411 | 7,0000 | 1,0972 |
| 192 | 4,5400 | 0,4407 | | 0,0411 | 7,0000 | 1,0972 |
| 193 | 4,5625 | 0,4387 | | 0,0411 | 7,0000 | 1,0972 |
| 194 | 4,5850 | 0,4368 | | 0,0411 | 7,0000 | 1,0972 |
| 195 | 4,6074 | 0,4348 | | 0,0411 | 7,0000 | 1,0972 |
| 196 | 4,6299 | 0,4328 | | 0,0411 | 7,0000 | 1,0972 |

| | 13-b-Final | 14-x column 1 | 15-y column 1 | 16 | 17-SumSq 1 | 18-Iterations 1 |
|---|---|---|---|---|---|---|
| 148 | 0,2009 | 3,5518 | 0,5480 | | 0,0352 | 7,0000 |
| 149 | 0,2009 | 3,5742 | 0,5457 | | 0,0352 | 12,0000 |
| 150 | 0,2009 | 3,5967 | 0,5434 | | 0,0352 | 33,0000 |
| 151 | 0,2009 | 3,6191 | 0,5410 | | 0,0352 | 7,0000 |
| 152 | 0,2009 | 3,6416 | 0,5387 | | 0,0352 | 7,0000 |
| 153 | 0,2009 | 3,6641 | 0,5364 | | 0,0352 | 7,0000 |
| 154 | 0,2009 | 3,6865 | 0,5342 | | 0,0352 | 10,0000 |
| 155 | 0,2009 | 3,7090 | 0,5319 | | 0,0352 | 7,0000 |
| 156 | 0,2009 | 3,7314 | 0,5296 | | 0,0352 | 7,0000 |
| 157 | 0,2009 | 3,7539 | 0,5274 | | 0,0352 | 7,0000 |
| 158 | 0,2009 | 3,7764 | 0,5251 | | 0,0352 | 7,0000 |
| 159 | 0,2009 | 3,7988 | 0,5229 | | 0,0352 | 7,0000 |
| 160 | 0,2009 | 3,8213 | 0,5207 | | 0,0352 | 7,0000 |
| 161 | 0,2009 | 3,8438 | 0,5185 | | 0,0352 | 7,0000 |
| 162 | 0,2009 | 3,8662 | 0,5163 | | 0,0352 | 10,0000 |
| 163 | 0,2009 | 3,8887 | 0,5141 | | 0,0352 | 7,0000 |
| 164 | 0,2009 | 3,9111 | 0,5119 | | 0,0352 | 7,0000 |
| 165 | 0,2009 | 3,9336 | 0,5097 | | 0,0352 | 7,0000 |
| 166 | 0,2009 | 3,9561 | 0,5075 | | 0,0352 | 7,0000 |
| 167 | 0,2009 | 3,9785 | 0,5054 | | 0,0352 | 7,0000 |
| 168 | 0,2009 | 4,0010 | 0,5032 | | 0,0352 | 7,0000 |
| 169 | 0,2009 | 4,0234 | 0,5011 | | 0,0352 | 7,0000 |
| 170 | 0,2009 | 4,0459 | 0,4990 | | 0,0352 | 7,0000 |
| 171 | 0,2009 | 4,0684 | 0,4968 | | 0,0352 | 7,0000 |
| 172 | 0,2009 | 4,0908 | 0,4947 | | 0,0352 | 7,0000 |
| 173 | 0,2009 | 4,1133 | 0,4926 | | 0,0352 | 7,0000 |
| 174 | 0,2009 | 4,1357 | 0,4905 | | 0,0352 | 10,0000 |
| 175 | 0,2009 | 4,1582 | 0,4884 | | 0,0352 | 7,0000 |
| 176 | 0,2009 | 4,1807 | 0,4864 | | 0,0352 | 7,0000 |
| 177 | 0,2009 | 4,2031 | 0,4843 | | 0,0352 | 10,0000 |
| 178 | 0,2009 | 4,2256 | 0,4822 | | 0,0352 | 7,0000 |
| 179 | 0,2009 | 4,2480 | 0,4802 | | 0,0352 | 7,0000 |
| 180 | 0,2009 | 4,2705 | 0,4781 | | 0,0352 | 7,0000 |
| 181 | 0,2009 | 4,2930 | 0,4761 | | 0,0352 | 7,0000 |
| 182 | 0,2009 | 4,3154 | 0,4741 | | 0,0352 | 10,0000 |
| 183 | 0,2009 | 4,3379 | 0,4721 | | 0,0352 | 7,0000 |
| 184 | 0,2009 | 4,3604 | 0,4701 | | 0,0352 | 7,0000 |
| 185 | 0,2009 | 4,3828 | 0,4681 | | 0,0352 | 7,0000 |
| 186 | 0,2009 | 4,4053 | 0,4661 | | 0,0352 | 7,0000 |
| 187 | 0,2009 | 4,4277 | 0,4641 | | 0,0352 | 7,0000 |
| 188 | 0,2009 | 4,4502 | 0,4621 | | 0,0352 | 7,0000 |
| 189 | 0,2009 | 4,4727 | 0,4601 | | 0,0352 | 7,0000 |
| 190 | 0,2009 | 4,4951 | 0,4582 | | 0,0352 | 7,0000 |
| 191 | 0,2009 | 4,5176 | 0,4562 | | 0,0352 | 7,0000 |
| 192 | 0,2009 | 4,5400 | 0,4543 | | 0,0352 | 9,0000 |
| 193 | 0,2009 | 4,5625 | 0,4524 | | 0,0352 | 7,0000 |
| 194 | 0,2009 | 4,5850 | 0,4504 | | 0,0352 | 7,0000 |
| 195 | 0,2009 | 4,6074 | 0,4485 | | 0,0352 | 7,0000 |
| 196 | 0,2009 | 4,6299 | 0,4466 | | 0,0352 | 7,0000 |

| | 19-a-Final 1 | 20-b-Final 1 | 21-x column 2 | 22-y column 2 | 23 | 24-SumSq 2 |
|---|---|---|---|---|---|---|
| 148 | 1,0751 | 0,1897 | 3,5518 | 0,7274 | | 0,0164 |
| 149 | 1,0751 | 0,1897 | 3,5742 | 0,7261 | | 0,0164 |
| 150 | 1,0751 | 0,1897 | 3,5967 | 0,7247 | | 0,0164 |
| 151 | 1,0751 | 0,1897 | 3,6191 | 0,7234 | | 0,0164 |
| 152 | 1,0751 | 0,1897 | 3,6416 | 0,7221 | | 0,0164 |
| 153 | 1,0751 | 0,1897 | 3,6641 | 0,7207 | | 0,0164 |
| 154 | 1,0751 | 0,1897 | 3,6865 | 0,7194 | | 0,0164 |
| 155 | 1,0751 | 0,1897 | 3,7090 | 0,7181 | | 0,0164 |
| 156 | 1,0751 | 0,1897 | 3,7314 | 0,7168 | | 0,0164 |
| 157 | 1,0751 | 0,1897 | 3,7539 | 0,7155 | | 0,0164 |
| 158 | 1,0751 | 0,1897 | 3,7764 | 0,7141 | | 0,0164 |
| 159 | 1,0751 | 0,1897 | 3,7988 | 0,7128 | | 0,0164 |
| 160 | 1,0751 | 0,1897 | 3,8213 | 0,7115 | | 0,0164 |
| 161 | 1,0751 | 0,1897 | 3,8438 | 0,7102 | | 0,0164 |
| 162 | 1,0751 | 0,1897 | 3,8662 | 0,7089 | | 0,0164 |
| 163 | 1,0751 | 0,1897 | 3,8887 | 0,7076 | | 0,0164 |
| 164 | 1,0751 | 0,1897 | 3,9111 | 0,7063 | | 0,0164 |
| 165 | 1,0751 | 0,1897 | 3,9336 | 0,7050 | | 0,0164 |
| 166 | 1,0751 | 0,1897 | 3,9561 | 0,7037 | | 0,0164 |
| 167 | 1,0751 | 0,1897 | 3,9785 | 0,7024 | | 0,0164 |
| 168 | 1,0751 | 0,1897 | 4,0010 | 0,7011 | | 0,0164 |
| 169 | 1,0751 | 0,1897 | 4,0234 | 0,6998 | | 0,0164 |
| 170 | 1,0751 | 0,1897 | 4,0459 | 0,6985 | | 0,0164 |
| 171 | 1,0751 | 0,1897 | 4,0684 | 0,6972 | | 0,0164 |
| 172 | 1,0751 | 0,1897 | 4,0908 | 0,6960 | | 0,0164 |
| 173 | 1,0751 | 0,1897 | 4,1133 | 0,6947 | | 0,0164 |
| 174 | 1,0751 | 0,1897 | 4,1357 | 0,6934 | | 0,0164 |
| 175 | 1,0751 | 0,1897 | 4,1582 | 0,6921 | | 0,0164 |
| 176 | 1,0751 | 0,1897 | 4,1807 | 0,6909 | | 0,0164 |
| 177 | 1,0751 | 0,1897 | 4,2031 | 0,6896 | | 0,0164 |
| 178 | 1,0751 | 0,1897 | 4,2256 | 0,6883 | | 0,0164 |
| 179 | 1,0751 | 0,1897 | 4,2480 | 0,6870 | | 0,0164 |
| 180 | 1,0751 | 0,1897 | 4,2705 | 0,6858 | | 0,0164 |
| 181 | 1,0751 | 0,1897 | 4,2930 | 0,6845 | | 0,0164 |
| 182 | 1,0751 | 0,1897 | 4,3154 | 0,6833 | | 0,0164 |
| 183 | 1,0751 | 0,1897 | 4,3379 | 0,6820 | | 0,0164 |
| 184 | 1,0751 | 0,1897 | 4,3604 | 0,6807 | | 0,0164 |
| 185 | 1,0751 | 0,1897 | 4,3828 | 0,6795 | | 0,0164 |
| 186 | 1,0751 | 0,1897 | 4,4053 | 0,6782 | | 0,0164 |
| 187 | 1,0751 | 0,1897 | 4,4277 | 0,6770 | | 0,0164 |
| 188 | 1,0751 | 0,1897 | 4,4502 | 0,6758 | | 0,0164 |
| 189 | 1,0751 | 0,1897 | 4,4727 | 0,6745 | | 0,0164 |
| 190 | 1,0751 | 0,1897 | 4,4951 | 0,6733 | | 0,0164 |
| 191 | 1,0751 | 0,1897 | 4,5176 | 0,6720 | | 0,0164 |
| 192 | 1,0751 | 0,1897 | 4,5400 | 0,6708 | | 0,0164 |
| 193 | 1,0751 | 0,1897 | 4,5625 | 0,6696 | | 0,0164 |
| 194 | 1,0751 | 0,1897 | 4,5850 | 0,6683 | | 0,0164 |
| 195 | 1,0751 | 0,1897 | 4,6074 | 0,6671 | | 0,0164 |
| 196 | 1,0751 | 0,1897 | 4,6299 | 0,6659 | | 0,0164 |

| | 25-Iterations 2 | 26-a-Final 2 | 27-b-Final 2 | 28-x column 3 | 29-y column 3 | 30 |
|---|---|---|---|---|---|---|
| 148 | 8,0000 | 0,9733 | 0,0820 | 3,5518 | 0,7487 | |
| 149 | 8,0000 | 0,9733 | 0,0820 | 3,5742 | 0,7471 | |
| 150 | 8,0000 | 0,9733 | 0,0820 | 3,5967 | 0,7455 | |
| 151 | 8,0000 | 0,9733 | 0,0820 | 3,6191 | 0,7440 | |
| 152 | 8,0000 | 0,9733 | 0,0820 | 3,6416 | 0,7424 | |
| 153 | 8,0000 | 0,9733 | 0,0820 | 3,6641 | 0,7408 | |
| 154 | 8,0000 | 0,9733 | 0,0820 | 3,6865 | 0,7392 | |
| 155 | 8,0000 | 0,9733 | 0,0820 | 3,7090 | 0,7377 | |
| 156 | 8,0000 | 0,9733 | 0,0820 | 3,7314 | 0,7361 | |
| 157 | 7,0000 | 0,9733 | 0,0820 | 3,7539 | 0,7346 | |
| 158 | 7,0000 | 0,9733 | 0,0820 | 3,7764 | 0,7330 | |
| 159 | 8,0000 | 0,9733 | 0,0820 | 3,7988 | 0,7314 | |
| 160 | 8,0000 | 0,9733 | 0,0820 | 3,8213 | 0,7299 | |
| 161 | 8,0000 | 0,9733 | 0,0820 | 3,8438 | 0,7283 | |
| 162 | 8,0000 | 0,9733 | 0,0820 | 3,8662 | 0,7268 | |
| 163 | 8,0000 | 0,9733 | 0,0820 | 3,8887 | 0,7253 | |
| 164 | 8,0000 | 0,9733 | 0,0820 | 3,9111 | 0,7237 | |
| 165 | 8,0000 | 0,9733 | 0,0820 | 3,9336 | 0,7222 | |
| 166 | 8,0000 | 0,9733 | 0,0820 | 3,9561 | 0,7207 | |
| 167 | 8,0000 | 0,9733 | 0,0820 | 3,9785 | 0,7191 | |
| 168 | 8,0000 | 0,9733 | 0,0820 | 4,0010 | 0,7176 | |
| 169 | 8,0000 | 0,9733 | 0,0820 | 4,0234 | 0,7161 | |
| 170 | 8,0000 | 0,9733 | 0,0820 | 4,0459 | 0,7146 | |
| 171 | 8,0000 | 0,9733 | 0,0820 | 4,0684 | 0,7131 | |
| 172 | 8,0000 | 0,9733 | 0,0820 | 4,0908 | 0,7115 | |
| 173 | 8,0000 | 0,9733 | 0,0820 | 4,1133 | 0,7100 | |
| 174 | 8,0000 | 0,9733 | 0,0820 | 4,1357 | 0,7085 | |
| 175 | 8,0000 | 0,9733 | 0,0820 | 4,1582 | 0,7070 | |
| 176 | 8,0000 | 0,9733 | 0,0820 | 4,1807 | 0,7055 | |
| 177 | 8,0000 | 0,9733 | 0,0820 | 4,2031 | 0,7040 | |
| 178 | 8,0000 | 0,9733 | 0,0820 | 4,2256 | 0,7025 | |
| 179 | 8,0000 | 0,9733 | 0,0820 | 4,2480 | 0,7011 | |
| 180 | 8,0000 | 0,9733 | 0,0820 | 4,2705 | 0,6996 | |
| 181 | 8,0000 | 0,9733 | 0,0820 | 4,2930 | 0,6981 | |
| 182 | 8,0000 | 0,9733 | 0,0820 | 4,3154 | 0,6966 | |
| 183 | 8,0000 | 0,9733 | 0,0820 | 4,3379 | 0,6951 | |
| 184 | 8,0000 | 0,9733 | 0,0820 | 4,3604 | 0,6937 | |
| 185 | 8,0000 | 0,9733 | 0,0820 | 4,3828 | 0,6922 | |
| 186 | 8,0000 | 0,9733 | 0,0820 | 4,4053 | 0,6907 | |
| 187 | 8,0000 | 0,9733 | 0,0820 | 4,4277 | 0,6893 | |
| 188 | 8,0000 | 0,9733 | 0,0820 | 4,4502 | 0,6878 | |
| 189 | 8,0000 | 0,9733 | 0,0820 | 4,4727 | 0,6863 | |
| 190 | 8,0000 | 0,9733 | 0,0820 | 4,4951 | 0,6849 | |
| 191 | 8,0000 | 0,9733 | 0,0820 | 4,5176 | 0,6834 | |
| 192 | 8,0000 | 0,9733 | 0,0820 | 4,5400 | 0,6820 | |
| 193 | 8,0000 | 0,9733 | 0,0820 | 4,5625 | 0,6805 | |
| 194 | 8,0000 | 0,9733 | 0,0820 | 4,5850 | 0,6791 | |
| 195 | 7,0000 | 0,9733 | 0,0820 | 4,6074 | 0,6777 | |
| 196 | 8,0000 | 0,9733 | 0,0820 | 4,6299 | 0,6762 | |

| | 31-SumSq 3 | 32-Iterations 3 | 33-a-Final 3 | 34-b-Final 3 | 35-x column 4 | 36-y column 4 |
|---|---|---|---|---|---|---|
| 148 | 0,0189 | 7,0000 | 1,0472 | 0,0945 | 3,5518 | 0,7422 |
| 149 | 0,0189 | 8,0000 | 1,0472 | 0,0945 | 3,5742 | 0,7408 |
| 150 | 0,0189 | 7,0000 | 1,0472 | 0,0945 | 3,5967 | 0,7393 |
| 151 | 0,0189 | 8,0000 | 1,0472 | 0,0945 | 3,6191 | 0,7378 |
| 152 | 0,0189 | 7,0000 | 1,0472 | 0,0945 | 3,6416 | 0,7364 |
| 153 | 0,0189 | 7,0000 | 1,0472 | 0,0945 | 3,6641 | 0,7349 |
| 154 | 0,0189 | 7,0000 | 1,0472 | 0,0945 | 3,6865 | 0,7335 |
| 155 | 0,0189 | 7,0000 | 1,0472 | 0,0945 | 3,7090 | 0,7320 |
| 156 | 0,0189 | 7,0000 | 1,0472 | 0,0945 | 3,7314 | 0,7306 |
| 157 | 0,0189 | 7,0000 | 1,0472 | 0,0945 | 3,7539 | 0,7292 |
| 158 | 0,0189 | 7,0000 | 1,0472 | 0,0945 | 3,7764 | 0,7277 |
| 159 | 0,0189 | 7,0000 | 1,0472 | 0,0945 | 3,7988 | 0,7263 |
| 160 | 0,0189 | 7,0000 | 1,0472 | 0,0945 | 3,8213 | 0,7249 |
| 161 | 0,0189 | 7,0000 | 1,0472 | 0,0945 | 3,8438 | 0,7234 |
| 162 | 0,0189 | 7,0000 | 1,0472 | 0,0945 | 3,8662 | 0,7220 |
| 163 | 0,0189 | 7,0000 | 1,0472 | 0,0945 | 3,8887 | 0,7206 |
| 164 | 0,0189 | 7,0000 | 1,0472 | 0,0945 | 3,9111 | 0,7192 |
| 165 | 0,0189 | 7,0000 | 1,0472 | 0,0945 | 3,9336 | 0,7178 |
| 166 | 0,0189 | 7,0000 | 1,0472 | 0,0945 | 3,9561 | 0,7164 |
| 167 | 0,0189 | 7,0000 | 1,0472 | 0,0945 | 3,9785 | 0,7149 |
| 168 | 0,0189 | 7,0000 | 1,0472 | 0,0945 | 4,0010 | 0,7135 |
| 169 | 0,0189 | 7,0000 | 1,0472 | 0,0945 | 4,0234 | 0,7121 |
| 170 | 0,0189 | 7,0000 | 1,0472 | 0,0945 | 4,0459 | 0,7107 |
| 171 | 0,0189 | 7,0000 | 1,0472 | 0,0945 | 4,0684 | 0,7093 |
| 172 | 0,0189 | 7,0000 | 1,0472 | 0,0945 | 4,0908 | 0,7079 |
| 173 | 0,0189 | 7,0000 | 1,0472 | 0,0945 | 4,1133 | 0,7065 |
| 174 | 0,0189 | 7,0000 | 1,0472 | 0,0945 | 4,1357 | 0,7052 |
| 175 | 0,0189 | 7,0000 | 1,0472 | 0,0945 | 4,1582 | 0,7038 |
| 176 | 0,0189 | 7,0000 | 1,0472 | 0,0945 | 4,1807 | 0,7024 |
| 177 | 0,0189 | 7,0000 | 1,0472 | 0,0945 | 4,2031 | 0,7010 |
| 178 | 0,0189 | 7,0000 | 1,0472 | 0,0945 | 4,2256 | 0,6996 |
| 179 | 0,0189 | 7,0000 | 1,0472 | 0,0945 | 4,2480 | 0,6982 |
| 180 | 0,0189 | 7,0000 | 1,0472 | 0,0945 | 4,2705 | 0,6969 |
| 181 | 0,0189 | 7,0000 | 1,0472 | 0,0945 | 4,2930 | 0,6955 |
| 182 | 0,0189 | 7,0000 | 1,0472 | 0,0945 | 4,3154 | 0,6941 |
| 183 | 0,0189 | 7,0000 | 1,0472 | 0,0945 | 4,3379 | 0,6928 |
| 184 | 0,0189 | 7,0000 | 1,0472 | 0,0945 | 4,3604 | 0,6914 |
| 185 | 0,0189 | 7,0000 | 1,0472 | 0,0945 | 4,3828 | 0,6900 |
| 186 | 0,0189 | 7,0000 | 1,0472 | 0,0945 | 4,4053 | 0,6887 |
| 187 | 0,0189 | 7,0000 | 1,0472 | 0,0945 | 4,4277 | 0,6873 |
| 188 | 0,0189 | 7,0000 | 1,0472 | 0,0945 | 4,4502 | 0,6860 |
| 189 | 0,0189 | 7,0000 | 1,0472 | 0,0945 | 4,4727 | 0,6846 |
| 190 | 0,0189 | 7,0000 | 1,0472 | 0,0945 | 4,4951 | 0,6833 |
| 191 | 0,0189 | 7,0000 | 1,0472 | 0,0945 | 4,5176 | 0,6819 |
| 192 | 0,0189 | 7,0000 | 1,0472 | 0,0945 | 4,5400 | 0,6806 |
| 193 | 0,0189 | 7,0000 | 1,0472 | 0,0945 | 4,5625 | 0,6793 |
| 194 | 0,0189 | 6,0000 | 1,0472 | 0,0945 | 4,5850 | 0,6779 |
| 195 | 0,0189 | 7,0000 | 1,0472 | 0,0945 | 4,6074 | 0,6766 |
| 196 | 0,0189 | 7,0000 | 1,0472 | 0,0945 | 4,6299 | 0,6753 |

| | 37 | 38-SumSq 4 | 39-Iterations 4 | 40-a-Final 4 | 41-b-Final 4 |
|---|---|---|---|---|---|
| 148 | | 0,0143 | 8,0000 | 1,0134 | 0,0877 |
| 149 | | 0,0143 | 8,0000 | 1,0134 | 0,0877 |
| 150 | | 0,0143 | 8,0000 | 1,0134 | 0,0877 |
| 151 | | 0,0143 | 8,0000 | 1,0134 | 0,0877 |
| 152 | | 0,0143 | 8,0000 | 1,0134 | 0,0877 |
| 153 | | 0,0143 | 8,0000 | 1,0134 | 0,0877 |
| 154 | | 0,0143 | 8,0000 | 1,0134 | 0,0877 |
| 155 | | 0,0143 | 8,0000 | 1,0134 | 0,0877 |
| 156 | | 0,0143 | 8,0000 | 1,0134 | 0,0877 |
| 157 | | 0,0143 | 8,0000 | 1,0134 | 0,0877 |
| 158 | | 0,0143 | 8,0000 | 1,0134 | 0,0877 |
| 159 | | 0,0143 | 8,0000 | 1,0134 | 0,0877 |
| 160 | | 0,0143 | 8,0000 | 1,0134 | 0,0877 |
| 161 | | 0,0143 | 8,0000 | 1,0134 | 0,0877 |
| 162 | | 0,0143 | 8,0000 | 1,0134 | 0,0877 |
| 163 | | 0,0143 | 8,0000 | 1,0134 | 0,0877 |
| 164 | | 0,0143 | 8,0000 | 1,0134 | 0,0877 |
| 165 | | 0,0143 | 8,0000 | 1,0134 | 0,0877 |
| 166 | | 0,0143 | 7,0000 | 1,0134 | 0,0877 |
| 167 | | 0,0143 | 8,0000 | 1,0134 | 0,0877 |
| 168 | | 0,0143 | 8,0000 | 1,0134 | 0,0877 |
| 169 | | 0,0143 | 8,0000 | 1,0134 | 0,0877 |
| 170 | | 0,0143 | 8,0000 | 1,0134 | 0,0877 |
| 171 | | 0,0143 | 8,0000 | 1,0134 | 0,0877 |
| 172 | | 0,0143 | 8,0000 | 1,0134 | 0,0877 |
| 173 | | 0,0143 | 8,0000 | 1,0134 | 0,0877 |
| 174 | | 0,0143 | 8,0000 | 1,0134 | 0,0877 |
| 175 | | 0,0143 | 7,0000 | 1,0134 | 0,0877 |
| 176 | | 0,0143 | 8,0000 | 1,0134 | 0,0877 |
| 177 | | 0,0143 | 7,0000 | 1,0134 | 0,0877 |
| 178 | | 0,0143 | 7,0000 | 1,0134 | 0,0877 |
| 179 | | 0,0143 | 7,0000 | 1,0134 | 0,0877 |
| 180 | | 0,0143 | 7,0000 | 1,0134 | 0,0877 |
| 181 | | 0,0143 | 7,0000 | 1,0134 | 0,0877 |
| 182 | | 0,0143 | 7,0000 | 1,0134 | 0,0877 |
| 183 | | 0,0143 | 7,0000 | 1,0134 | 0,0877 |
| 184 | | 0,0143 | 7,0000 | 1,0134 | 0,0877 |
| 185 | | 0,0143 | 7,0000 | 1,0134 | 0,0877 |
| 186 | | 0,0143 | 7,0000 | 1,0134 | 0,0877 |
| 187 | | 0,0143 | 7,0000 | 1,0134 | 0,0877 |
| 188 | | 0,0143 | 7,0000 | 1,0134 | 0,0877 |
| 189 | | 0,0143 | 7,0000 | 1,0134 | 0,0877 |
| 190 | | 0,0143 | 6,0000 | 1,0134 | 0,0877 |
| 191 | | 0,0143 | 7,0000 | 1,0134 | 0,0877 |
| 192 | | 0,0143 | 7,0000 | 1,0134 | 0,0877 |
| 193 | | 0,0143 | 7,0000 | 1,0134 | 0,0877 |
| 194 | | 0,0143 | 7,0000 | 1,0134 | 0,0877 |
| 195 | | 0,0143 | 7,0000 | 1,0134 | 0,0877 |
| 196 | | 0,0143 | 7,0000 | 1,0134 | 0,0877 |

| | 1 | 2 | 3 | 4 | 5 | 6 |
|---|---|---|---|---|---|---|
| 197 | | | | | | |
| 198 | | | | | | |
| 199 | | | | | | |
| 200 | | | | | | |
| 201 | | | | | | |
| 202 | | | | | | |
| 203 | | | | | | |
| 204 | | | | | | |
| 205 | | | | | | |
| 206 | | | | | | |
| 207 | | | | | | |
| 208 | | | | | | |
| 209 | | | | | | |
| 210 | | | | | | |
| 211 | | | | | | |
| 212 | | | | | | |
| 213 | | | | | | |
| 214 | | | | | | |
| 215 | | | | | | |
| 216 | | | | | | |
| 217 | | | | | | |
| 218 | | | | | | |
| 219 | | | | | | |
| 220 | | | | | | |
| 221 | | | | | | |
| 222 | | | | | | |
| 223 | | | | | | |
| 224 | | | | | | |
| 225 | | | | | | |
| 226 | | | | | | |
| 227 | | | | | | |
| 228 | | | | | | |
| 229 | | | | | | |
| 230 | | | | | | |
| 231 | | | | | | |
| 232 | | | | | | |
| 233 | | | | | | |
| 234 | | | | | | |
| 235 | | | | | | |
| 236 | | | | | | |
| 237 | | | | | | |
| 238 | | | | | | |
| 239 | | | | | | |
| 240 | | | | | | |
| 241 | | | | | | |
| 242 | | | | | | |
| 243 | | | | | | |
| 244 | | | | | | |
| 245 | | | | | | |

| | 7-x column | 8-y column | 9 | 10-SumSq | 11-Iterations | 12-a-Final |
|---|---|---|---|---|---|---|
| 197 | 4,6523 | 0,4309 | | 0,0411 | 7,0000 | 1,0972 |
| 198 | 4,6748 | 0,4290 | | 0,0411 | 7,0000 | 1,0972 |
| 199 | 4,6973 | 0,4270 | | 0,0411 | 7,0000 | 1,0972 |
| 200 | 4,7197 | 0,4251 | | 0,0411 | 6,0000 | 1,0972 |
| 201 | 4,7422 | 0,4232 | | | | |
| 202 | 4,7646 | 0,4213 | | | | |
| 203 | 4,7871 | 0,4194 | | | | |
| 204 | 4,8096 | 0,4175 | | | | |
| 205 | 4,8320 | 0,4156 | | | | |
| 206 | 4,8545 | 0,4138 | | | | |
| 207 | 4,8770 | 0,4119 | | | | |
| 208 | 4,8994 | 0,4100 | | | | |
| 209 | 4,9219 | 0,4082 | | | | |
| 210 | 4,9443 | 0,4064 | | | | |
| 211 | 4,9668 | 0,4045 | | | | |
| 212 | 4,9893 | 0,4027 | | | | |
| 213 | 5,0117 | 0,4009 | | | | |
| 214 | 5,0342 | 0,3991 | | | | |
| 215 | 5,0566 | 0,3973 | | | | |
| 216 | 5,0791 | 0,3955 | | | | |
| 217 | 5,1016 | 0,3937 | | | | |
| 218 | 5,1240 | 0,3919 | | | | |
| 219 | 5,1465 | 0,3902 | | | | |
| 220 | 5,1689 | 0,3884 | | | | |
| 221 | 5,1914 | 0,3867 | | | | |
| 222 | 5,2139 | 0,3849 | | | | |
| 223 | 5,2363 | 0,3832 | | | | |
| 224 | 5,2588 | 0,3815 | | | | |
| 225 | 5,2813 | 0,3798 | | | | |
| 226 | 5,3037 | 0,3780 | | | | |
| 227 | 5,3262 | 0,3763 | | | | |
| 228 | 5,3486 | 0,3747 | | | | |
| 229 | 5,3711 | 0,3730 | | | | |
| 230 | 5,3936 | 0,3713 | | | | |
| 231 | 5,4160 | 0,3696 | | | | |
| 232 | 5,4385 | 0,3680 | | | | |
| 233 | 5,4609 | 0,3663 | | | | |
| 234 | 5,4834 | 0,3646 | | | | |
| 235 | 5,5059 | 0,3630 | | | | |
| 236 | 5,5283 | 0,3614 | | | | |
| 237 | 5,5508 | 0,3597 | | | | |
| 238 | 5,5732 | 0,3581 | | | | |
| 239 | 5,5957 | 0,3565 | | | | |
| 240 | 5,6182 | 0,3549 | | | | |
| 241 | 5,6406 | 0,3533 | | | | |
| 242 | 5,6631 | 0,3517 | | | | |
| 243 | 5,6855 | 0,3501 | | | | |
| 244 | 5,7080 | 0,3486 | | | | |
| 245 | 5,7305 | 0,3470 | | | | |

| | 13-b-Final | 14-x column 1 | 15-y column 1 | 16 | 17-SumSq 1 | 18-Iterations 1 |
|---|---|---|---|---|---|---|
| 197 | 0,2009 | 4,6523 | 0,4447 | | 0,0352 | 7,0000 |
| 198 | 0,2009 | 4,6748 | 0,4428 | | 0,0352 | 7,0000 |
| 199 | 0,2009 | 4,6973 | 0,4409 | | 0,0352 | 7,0000 |
| 200 | 0,2009 | 4,7197 | 0,4391 | | 0,0352 | 6,0000 |
| 201 | | 4,7422 | 0,4372 | | | |
| 202 | | 4,7646 | 0,4353 | | | |
| 203 | | 4,7871 | 0,4335 | | | |
| 204 | | 4,8096 | 0,4317 | | | |
| 205 | | 4,8320 | 0,4298 | | | |
| 206 | | 4,8545 | 0,4280 | | | |
| 207 | | 4,8770 | 0,4262 | | | |
| 208 | | 4,8994 | 0,4244 | | | |
| 209 | | 4,9219 | 0,4226 | | | |
| 210 | | 4,9443 | 0,4208 | | | |
| 211 | | 4,9668 | 0,4190 | | | |
| 212 | | 4,9893 | 0,4172 | | | |
| 213 | | 5,0117 | 0,4154 | | | |
| 214 | | 5,0342 | 0,4136 | | | |
| 215 | | 5,0566 | 0,4119 | | | |
| 216 | | 5,0791 | 0,4101 | | | |
| 217 | | 5,1016 | 0,4084 | | | |
| 218 | | 5,1240 | 0,4067 | | | |
| 219 | | 5,1465 | 0,4049 | | | |
| 220 | | 5,1689 | 0,4032 | | | |
| 221 | | 5,1914 | 0,4015 | | | |
| 222 | | 5,2139 | 0,3998 | | | |
| 223 | | 5,2363 | 0,3981 | | | |
| 224 | | 5,2588 | 0,3964 | | | |
| 225 | | 5,2813 | 0,3947 | | | |
| 226 | | 5,3037 | 0,3930 | | | |
| 227 | | 5,3262 | 0,3914 | | | |
| 228 | | 5,3486 | 0,3897 | | | |
| 229 | | 5,3711 | 0,3880 | | | |
| 230 | | 5,3936 | 0,3864 | | | |
| 231 | | 5,4160 | 0,3847 | | | |
| 232 | | 5,4385 | 0,3831 | | | |
| 233 | | 5,4609 | 0,3815 | | | |
| 234 | | 5,4834 | 0,3798 | | | |
| 235 | | 5,5059 | 0,3782 | | | |
| 236 | | 5,5283 | 0,3766 | | | |
| 237 | | 5,5508 | 0,3750 | | | |
| 238 | | 5,5732 | 0,3734 | | | |
| 239 | | 5,5957 | 0,3718 | | | |
| 240 | | 5,6182 | 0,3703 | | | |
| 241 | | 5,6406 | 0,3687 | | | |
| 242 | | 5,6631 | 0,3671 | | | |
| 243 | | 5,6855 | 0,3656 | | | |
| 244 | | 5,7080 | 0,3640 | | | |
| 245 | | 5,7305 | 0,3625 | | | |

| | 19-a-Final 1 | 20-b-Final 1 | 21-x column 2 | 22-y column 2 | 23 | 24-SumSq 2 |
|---|---|---|---|---|---|---|
| 197 | 1,0751 | 0,1897 | 4,6523 | 0,6646 | | 0,0164 |
| 198 | 1,0751 | 0,1897 | 4,6748 | 0,6634 | | 0,0164 |
| 199 | 1,0751 | 0,1897 | 4,6973 | 0,6622 | | 0,0164 |
| 200 | 1,0751 | 0,1897 | 4,7197 | 0,6610 | | 0,0164 |
| 201 | | | 4,7422 | 0,6598 | | |
| 202 | | | 4,7646 | 0,6586 | | |
| 203 | | | 4,7871 | 0,6573 | | |
| 204 | | | 4,8096 | 0,6561 | | |
| 205 | | | 4,8320 | 0,6549 | | |
| 206 | | | 4,8545 | 0,6537 | | |
| 207 | | | 4,8770 | 0,6525 | | |
| 208 | | | 4,8994 | 0,6513 | | |
| 209 | | | 4,9219 | 0,6501 | | |
| 210 | | | 4,9443 | 0,6489 | | |
| 211 | | | 4,9668 | 0,6477 | | |
| 212 | | | 4,9893 | 0,6465 | | |
| 213 | | | 5,0117 | 0,6453 | | |
| 214 | | | 5,0342 | 0,6442 | | |
| 215 | | | 5,0566 | 0,6430 | | |
| 216 | | | 5,0791 | 0,6418 | | |
| 217 | | | 5,1016 | 0,6406 | | |
| 218 | | | 5,1240 | 0,6394 | | |
| 219 | | | 5,1465 | 0,6383 | | |
| 220 | | | 5,1689 | 0,6371 | | |
| 221 | | | 5,1914 | 0,6359 | | |
| 222 | | | 5,2139 | 0,6347 | | |
| 223 | | | 5,2363 | 0,6336 | | |
| 224 | | | 5,2588 | 0,6324 | | |
| 225 | | | 5,2813 | 0,6312 | | |
| 226 | | | 5,3037 | 0,6301 | | |
| 227 | | | 5,3262 | 0,6289 | | |
| 228 | | | 5,3486 | 0,6278 | | |
| 229 | | | 5,3711 | 0,6266 | | |
| 230 | | | 5,3936 | 0,6255 | | |
| 231 | | | 5,4160 | 0,6243 | | |
| 232 | | | 5,4385 | 0,6232 | | |
| 233 | | | 5,4609 | 0,6220 | | |
| 234 | | | 5,4834 | 0,6209 | | |
| 235 | | | 5,5059 | 0,6197 | | |
| 236 | | | 5,5283 | 0,6186 | | |
| 237 | | | 5,5508 | 0,6174 | | |
| 238 | | | 5,5732 | 0,6163 | | |
| 239 | | | 5,5957 | 0,6152 | | |
| 240 | | | 5,6182 | 0,6140 | | |
| 241 | | | 5,6406 | 0,6129 | | |
| 242 | | | 5,6631 | 0,6118 | | |
| 243 | | | 5,6855 | 0,6107 | | |
| 244 | | | 5,7080 | 0,6095 | | |
| 245 | | | 5,7305 | 0,6084 | | |

| | 25-Iterations 2 | 26-a-Final 2 | 27-b-Final 2 | 28-x column 3 | 29-y column 3 | 30 |
|---|---|---|---|---|---|---|
| 197 | 8,0000 | 0,9733 | 0,0820 | 4,6523 | 0,6748 | |
| 198 | 8,0000 | 0,9733 | 0,0820 | 4,6748 | 0,6734 | |
| 199 | 8,0000 | 0,9733 | 0,0820 | 4,6973 | 0,6719 | |
| 200 | 8,0000 | 0,9733 | 0,0820 | 4,7197 | 0,6705 | |
| 201 | | | | 4,7422 | 0,6691 | |
| 202 | | | | 4,7646 | 0,6677 | |
| 203 | | | | 4,7871 | 0,6663 | |
| 204 | | | | 4,8096 | 0,6648 | |
| 205 | | | | 4,8320 | 0,6634 | |
| 206 | | | | 4,8545 | 0,6620 | |
| 207 | | | | 4,8770 | 0,6606 | |
| 208 | | | | 4,8994 | 0,6592 | |
| 209 | | | | 4,9219 | 0,6578 | |
| 210 | | | | 4,9443 | 0,6564 | |
| 211 | | | | 4,9668 | 0,6550 | |
| 212 | | | | 4,9893 | 0,6537 | |
| 213 | | | | 5,0117 | 0,6523 | |
| 214 | | | | 5,0342 | 0,6509 | |
| 215 | | | | 5,0566 | 0,6495 | |
| 216 | | | | 5,0791 | 0,6481 | |
| 217 | | | | 5,1016 | 0,6468 | |
| 218 | | | | 5,1240 | 0,6454 | |
| 219 | | | | 5,1465 | 0,6440 | |
| 220 | | | | 5,1689 | 0,6427 | |
| 221 | | | | 5,1914 | 0,6413 | |
| 222 | | | | 5,2139 | 0,6399 | |
| 223 | | | | 5,2363 | 0,6386 | |
| 224 | | | | 5,2588 | 0,6372 | |
| 225 | | | | 5,2813 | 0,6359 | |
| 226 | | | | 5,3037 | 0,6345 | |
| 227 | | | | 5,3262 | 0,6332 | |
| 228 | | | | 5,3486 | 0,6318 | |
| 229 | | | | 5,3711 | 0,6305 | |
| 230 | | | | 5,3936 | 0,6292 | |
| 231 | | | | 5,4160 | 0,6278 | |
| 232 | | | | 5,4385 | 0,6265 | |
| 233 | | | | 5,4609 | 0,6252 | |
| 234 | | | | 5,4834 | 0,6238 | |
| 235 | | | | 5,5059 | 0,6225 | |
| 236 | | | | 5,5283 | 0,6212 | |
| 237 | | | | 5,5508 | 0,6199 | |
| 238 | | | | 5,5732 | 0,6186 | |
| 239 | | | | 5,5957 | 0,6173 | |
| 240 | | | | 5,6182 | 0,6160 | |
| 241 | | | | 5,6406 | 0,6146 | |
| 242 | | | | 5,6631 | 0,6133 | |
| 243 | | | | 5,6855 | 0,6120 | |
| 244 | | | | 5,7080 | 0,6108 | |
| 245 | | | | 5,7305 | 0,6095 | |

| | 31-SumSq 3 | 32-Iterations 3 | 33-a-Final 3 | 34-b-Final 3 | 35-x column 4 | 36-y column 4 |
|---|---|---|---|---|---|---|
| 197 | 0,0189 | 7,0000 | 1,0472 | 0,0945 | 4,6523 | 0,6739 |
| 198 | 0,0189 | 7,0000 | 1,0472 | 0,0945 | 4,6748 | 0,6726 |
| 199 | 0,0189 | 7,0000 | 1,0472 | 0,0945 | 4,6973 | 0,6713 |
| 200 | 0,0189 | 7,0000 | 1,0472 | 0,0945 | 4,7197 | 0,6700 |
| 201 | | | | | 4,7422 | 0,6686 |
| 202 | | | | | 4,7646 | 0,6673 |
| 203 | | | | | 4,7871 | 0,6660 |
| 204 | | | | | 4,8096 | 0,6647 |
| 205 | | | | | 4,8320 | 0,6634 |
| 206 | | | | | 4,8545 | 0,6621 |
| 207 | | | | | 4,8770 | 0,6608 |
| 208 | | | | | 4,8994 | 0,6595 |
| 209 | | | | | 4,9219 | 0,6582 |
| 210 | | | | | 4,9443 | 0,6569 |
| 211 | | | | | 4,9668 | 0,6556 |
| 212 | | | | | 4,9893 | 0,6543 |
| 213 | | | | | 5,0117 | 0,6530 |
| 214 | | | | | 5,0342 | 0,6517 |
| 215 | | | | | 5,0566 | 0,6504 |
| 216 | | | | | 5,0791 | 0,6492 |
| 217 | | | | | 5,1016 | 0,6479 |
| 218 | | | | | 5,1240 | 0,6466 |
| 219 | | | | | 5,1465 | 0,6453 |
| 220 | | | | | 5,1689 | 0,6441 |
| 221 | | | | | 5,1914 | 0,6428 |
| 222 | | | | | 5,2139 | 0,6415 |
| 223 | | | | | 5,2363 | 0,6403 |
| 224 | | | | | 5,2588 | 0,6390 |
| 225 | | | | | 5,2813 | 0,6378 |
| 226 | | | | | 5,3037 | 0,6365 |
| 227 | | | | | 5,3262 | 0,6353 |
| 228 | | | | | 5,3486 | 0,6340 |
| 229 | | | | | 5,3711 | 0,6328 |
| 230 | | | | | 5,3936 | 0,6315 |
| 231 | | | | | 5,4160 | 0,6303 |
| 232 | | | | | 5,4385 | 0,6290 |
| 233 | | | | | 5,4609 | 0,6278 |
| 234 | | | | | 5,4834 | 0,6266 |
| 235 | | | | | 5,5059 | 0,6253 |
| 236 | | | | | 5,5283 | 0,6241 |
| 237 | | | | | 5,5508 | 0,6229 |
| 238 | | | | | 5,5732 | 0,6216 |
| 239 | | | | | 5,5957 | 0,6204 |
| 240 | | | | | 5,6182 | 0,6192 |
| 241 | | | | | 5,6406 | 0,6180 |
| 242 | | | | | 5,6631 | 0,6168 |
| 243 | | | | | 5,6855 | 0,6155 |
| 244 | | | | | 5,7080 | 0,6143 |
| 245 | | | | | 5,7305 | 0,6131 |

| | 37 | 38-SumSq 4 | 39-Iterations 4 | 40-a-Final 4 | 41-b-Final 4 |
|---|---|---|---|---|---|
| 197 | | 0,0143 | 7,0000 | 1,0134 | 0,0877 |
| 198 | | 0,0143 | 7,0000 | 1,0134 | 0,0877 |
| 199 | | 0,0143 | 7,0000 | 1,0134 | 0,0877 |
| 200 | | 0,0143 | 7,0000 | 1,0134 | 0,0877 |
| 201 | | | | | |
| 202 | | | | | |
| 203 | | | | | |
| 204 | | | | | |
| 205 | | | | | |
| 206 | | | | | |
| 207 | | | | | |
| 208 | | | | | |
| 209 | | | | | |
| 210 | | | | | |
| 211 | | | | | |
| 212 | | | | | |
| 213 | | | | | |
| 214 | | | | | |
| 215 | | | | | |
| 216 | | | | | |
| 217 | | | | | |
| 218 | | | | | |
| 219 | | | | | |
| 220 | | | | | |
| 221 | | | | | |
| 222 | | | | | |
| 223 | | | | | |
| 224 | | | | | |
| 225 | | | | | |
| 226 | | | | | |
| 227 | | | | | |
| 228 | | | | | |
| 229 | | | | | |
| 230 | | | | | |
| 231 | | | | | |
| 232 | | | | | |
| 233 | | | | | |
| 234 | | | | | |
| 235 | | | | | |
| 236 | | | | | |
| 237 | | | | | |
| 238 | | | | | |
| 239 | | | | | |
| 240 | | | | | |
| 241 | | | | | |
| 242 | | | | | |
| 243 | | | | | |
| 244 | | | | | |
| 245 | | | | | |

| | 1 | 2 | 3 | 4 | 5 | 6 |
|---|---|---|---|---|---|---|
| 246 | | | | | | |
| 247 | | | | | | |
| 248 | | | | | | |
| 249 | | | | | | |
| 250 | | | | | | |
| 251 | | | | | | |
| 252 | | | | | | |
| 253 | | | | | | |
| 254 | | | | | | |
| 255 | | | | | | |
| 256 | | | | | | |
| 257 | | | | | | |

| | 7-x column | 8-y column | 9 | 10-SumSq | 11-Iterations | 12-a-Final |
|---|---|---|---|---|---|---|
| 246 | 5,7529 | 0,3454 | | | | |
| 247 | 5,7754 | 0,3439 | | | | |
| 248 | 5,7979 | 0,3423 | | | | |
| 249 | 5,8203 | 0,3408 | | | | |
| 250 | 5,8428 | 0,3392 | | | | |
| 251 | 5,8652 | 0,3377 | | | | |
| 252 | 5,8877 | 0,3362 | | | | |
| 253 | 5,9102 | 0,3347 | | | | |
| 254 | 5,9326 | 0,3332 | | | | |
| 255 | 5,9551 | 0,3317 | | | | |
| 256 | 5,9775 | 0,3302 | | | | |
| 257 | 6,0000 | 0,3287 | | | | |

| | 13-b-Final | 14-x column 1 | 15-y column 1 | 16 | 17-SumSq 1 | 18-Iterations 1 |
|---|---|---|---|---|---|---|
| 246 | | 5,7529 | 0,3609 | | | |
| 247 | | 5,7754 | 0,3594 | | | |
| 248 | | 5,7979 | 0,3578 | | | |
| 249 | | 5,8203 | 0,3563 | | | |
| 250 | | 5,8428 | 0,3548 | | | |
| 251 | | 5,8652 | 0,3533 | | | |
| 252 | | 5,8877 | 0,3518 | | | |
| 253 | | 5,9102 | 0,3503 | | | |
| 254 | | 5,9326 | 0,3488 | | | |
| 255 | | 5,9551 | 0,3473 | | | |
| 256 | | 5,9775 | 0,3459 | | | |
| 257 | | 6,0000 | 0,3444 | | | |

| | 19-a-Final 1 | 20-b-Final 1 | 21-x column 2 | 22-y column 2 | 23 | 24-SumSq 2 |
|---|---|---|---|---|---|---|
| 246 | | | 5,7529 | 0,6073 | | |
| 247 | | | 5,7754 | 0,6062 | | |
| 248 | | | 5,7979 | 0,6051 | | |
| 249 | | | 5,8203 | 0,6039 | | |
| 250 | | | 5,8428 | 0,6028 | | |
| 251 | | | 5,8652 | 0,6017 | | |
| 252 | | | 5,8877 | 0,6006 | | |
| 253 | | | 5,9102 | 0,5995 | | |
| 254 | | | 5,9326 | 0,5984 | | |
| 255 | | | 5,9551 | 0,5973 | | |
| 256 | | | 5,9775 | 0,5962 | | |
| 257 | | | 6,0000 | 0,5951 | | |

| | 25-Iterations 2 | 26-a-Final 2 | 27-b-Final 2 | 28-x column 3 | 29-y column 3 | 30 |
|---|---|---|---|---|---|---|
| 246 | | | | 5,7529 | 0,6082 | |
| 247 | | | | 5,7754 | 0,6069 | |
| 248 | | | | 5,7979 | 0,6056 | |
| 249 | | | | 5,8203 | 0,6043 | |
| 250 | | | | 5,8428 | 0,6030 | |
| 251 | | | | 5,8652 | 0,6017 | |
| 252 | | | | 5,8877 | 0,6005 | |
| 253 | | | | 5,9102 | 0,5992 | |
| 254 | | | | 5,9326 | 0,5979 | |
| 255 | | | | 5,9551 | 0,5967 | |
| 256 | | | | 5,9775 | 0,5954 | |
| 257 | | | | 6,0000 | 0,5941 | |

| | 31-SumSq 3 | 32-Iterations 3 | 33-a-Final 3 | 34-b-Final 3 | 35-x column 4 | 36-y column 4 |
|---|---|---|---|---|---|---|
| 246 | | | | | 5,7529 | 0,6119 |
| 247 | | | | | 5,7754 | 0,6107 |
| 248 | | | | | 5,7979 | 0,6095 |
| 249 | | | | | 5,8203 | 0,6083 |
| 250 | | | | | 5,8428 | 0,6071 |
| 251 | | | | | 5,8652 | 0,6059 |
| 252 | | | | | 5,8877 | 0,6047 |
| 253 | | | | | 5,9102 | 0,6035 |
| 254 | | | | | 5,9326 | 0,6024 |
| 255 | | | | | 5,9551 | 0,6012 |
| 256 | | | | | 5,9775 | 0,6000 |
| 257 | | | | | 6,0000 | 0,5988 |

| | 37 | 38-SumSq 4 | 39-Iterations 4 | 40-a-Final 4 | 41-b-Final 4 |
|---|---|---|---|---|---|
| 246 | | | | | |
| 247 | | | | | |
| 248 | | | | | |
| 249 | | | | | |
| 250 | | | | | |
| 251 | | | | | |
| 252 | | | | | |
| 253 | | | | | |
| 254 | | | | | |
| 255 | | | | | |
| 256 | | | | | |
| 257 | | | | | |

**Nonlinear Regression - Dynamic Fitting** **czwartek, maj 07, 2026, 16:03:32**

**Data Source: Copy (2) of Data 1 in Notebook1 half life calculations.JNB**
**Equation: Exponential Decay; Single, 2 Parameter**
f = a*exp(-b*x)

**Dynamic Fit Options:**

| | |
|---|---|
| Total Number of Fits | 200 |
| Maximum Number of Iterations | 200 |

**Parameter Ranges for Initial Estimates:**

| | **Minimum** | **Maximum** |
|---|---|---|
| a | 0,9900 | 1,0000 |
| b | 0,0000 | 1,0000 |

**Summary of Fit Results:**
Converged 100,0%

**Results for the Overall Best-Fit Solution:**

| **R** | **Rsqr** | **Adj Rsqr** | **Standard Error of Estimate** |
|---|---|---|---|
| 0,9259 | 0,8573 | 0,8288 | 0,0572 |

| | **Coefficient** | **Std. Error** | **t** | **P** |
|---|---|---|---|---|
| a | 0,9733 | 0,0445 | 21,8876 | <0,0001 |
| b | 0,0820 | 0,0149 | 5,5033 | 0,0027 |

**Analysis of Variance:**

Analysis of Variance:

| | **DF** | **SS** | **MS** |
|---|---|---|---|
| Regression | 2 | 4,2382 | 2,1191 |
| Residual | 5 | 0,0164 | 0,0033 |
| Total | 7 | 4,2546 | 0,6078 |

Corrected for the mean of the observations:

| | **DF** | **SS** | **MS** | **F** | **P** |
|---|---|---|---|---|---|
| Regression | 1 | 0,0984 | 0,0984 | 30,0423 | 0,0028 |
| Residual | 5 | 0,0164 | 0,0033 | | |
| Total | 6 | 0,1148 | 0,0191 | | |

**Statistical Tests:**

**Normality Test (Shapiro-Wilk)** Passed (P = 0,8532)

W Statistic= 0,9641 Significance Level = <0,0001

**Constant Variance Test** Passed (P = 0,6019)

**Fit Equation Description:**

```
[Variables]
x = col(1)
y = col(2)
reciprocal_y = 1/abs(y)
reciprocal_ysquare = 1/y^2
'Automatic Initial Parameter Estimate Functions
F(q)=if(size(x)>1; if(total(abs(y))>0; ape(x;log(abs(y));1;0;1); -306); 0)
asign(q)=if(mean(q)>=0;1;-1)
[Parameters]
a = if(F(0)[1]< 307; if(F(0)[1]>-307; asign(y)*10^F(0)[1]; asign(y)*10^(-307)); asign(y)*10^307) "Auto
{{previous: 0,973333}} {{MinRange: 0,99}} {{MaxRange: 1}}
b = if(x50(x;y)-min(x)=0; 1;  -ln(,5)/(x50(x;y)-min(x)))  "Auto {{previous: 0,0819964}} {{MinRange: 0}}
{{MaxRange: 1}}
[Equation]
f = a*exp(-b*x)
fit f to y
"fit f to y with weight reciprocal_y
"fit f to y with weight reciprocal_ysquare
[Constraints]
b>0
[Options]
tolerance=1e-10
stepsize=1
iterations=200
```

Number of Iterations Performed = 9

7**Nonlinear Regression - Dynamic Fitting** **czwartek, maj 07, 2026, 16:09:09**

**Data Source: Copy (2) of Data 1 in Notebook1 half life calculations.JNB**
**Equation: Exponential Decay; Single, 2 Parameter**
f = a*exp(-b*x)

**Dynamic Fit Options:**

| | |
|---|---|
| Total Number of Fits | 200 |
| Maximum Number of Iterations | 200 |

**Parameter Ranges for Initial Estimates:**

| | Minimum | Maximum |
|---|---|---|
| a | 0,9900 | 1,0000 |
| b | 0,0000 | 1,0000 |

**Summary of Fit Results:**

| | |
|---|---|
| Converged | 100,0% |

**Results for the Overall Best-Fit Solution:**

| R | Rsqr | Adj Rsqr | Standard Error of Estimate |
|---|---|---|---|
| 0,9353 | 0,8749 | 0,8436 | 0,0598 |

| | Coefficient | Std. Error | t | P |
|---|---|---|---|---|
| a | 1,0134 | 0,0554 | 18,2908 | <0,0001 |
| b | 0,0877 | 0,0164 | 5,3511 | 0,0059 |

**Analysis of Variance:**

Analysis of Variance:

| | DF | SS | MS |
|---|---|---|---|
| Regression | 2 | 3,6082 | 1,8041 |
| Residual | 4 | 0,0143 | 0,0036 |
| Total | 6 | 3,6225 | 0,6037 |

Corrected for the mean of the observations:

| | DF | SS | MS | F | P |
|---|---|---|---|---|---|
| Regression | 1 | 0,1001 | 0,1001 | 27,9656 | 0,0061 |
| Residual | 4 | 0,0143 | 0,0036 | | |
| Total | 5 | 0,1145 | 0,0229 | | |

**Statistical Tests:**

**Normality Test (Shapiro-Wilk)** Passed (P = 0,9550)

W Statistic= 0,9807 Significance Level = <0,0001

**Constant Variance Test** Passed (P = 0,0600)

**Fit Equation Description:**

```
[Variables]
x = col(6)
y = col(4)
reciprocal_y = 1/abs(y)
reciprocal_ysquare = 1/y^2
'Automatic Initial Parameter Estimate Functions
F(q)=if(size(x)>1; if(total(abs(y))>0; ape(x;log(abs(y));1;0;1); -306); 0)
asign(q)=if(mean(q)>=0;1;-1)
[Parameters]
a = if(F(0)[1]< 307; if(F(0)[1]>-307; asign(y)*10^F(0)[1]; asign(y)*10^(-307)); asign(y)*10^307) "Auto {{previous: 1,01345}} {{MinRange: 0,99}} {{MaxRange: 1}}
b = if(x50(x;y)-min(x)=0; 1;  -ln(,5)/(x50(x;y)-min(x)))  "Auto {{previous: 0,0876973}} {{MinRange: 0}} {{MaxRange: 1}}
[Equation]
f = a*exp(-b*x)
fit f to y
"fit f to y with weight reciprocal_y
"fit f to y with weight reciprocal_ysquare
[Constraints]
b>0
[Options]
tolerance=1e-10
stepsize=1
iterations=200
```

Number of Iterations Performed = 8

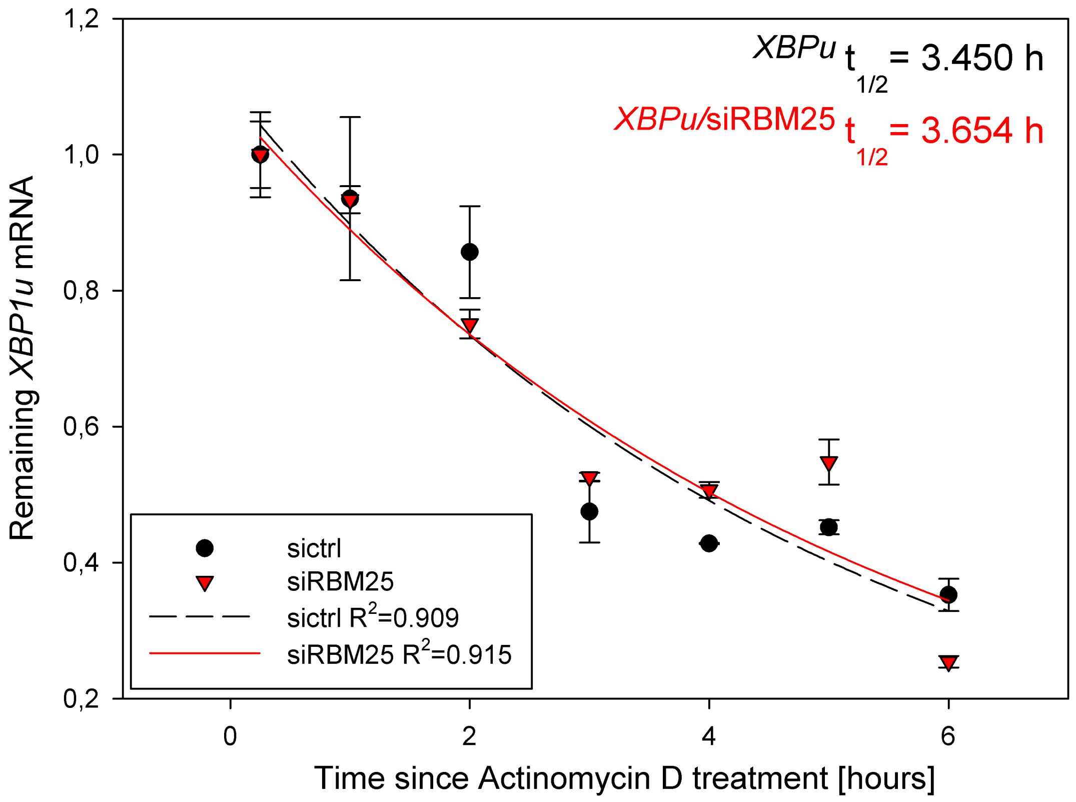

XBPu t1/2 = 3.450 h
XBPu/siRBM25 t1/2 = 3.654 h
Remaining XBP1u mRNA
Time since Actinomycin D treatment [hours]
sictrl
siRBM25
sictrl R2=0.909
siRBM25 R2=0.915
1,2
1,0
0,8
0,6
0,4
0,2
0
2
4
6

| | 1 | 2 | 3 | 4 | 5 | 6 |
|---|---|---|---|---|---|---|
| 1 | 0,2500 | 1,0000 | 0,0625 | 1,0000 | 0,0490 | 0,2500 |
| 2 | 1,0000 | 0,9352 | 0,1200 | 0,9335 | 0,0200 | 1,0000 |
| 3 | 2,0000 | 0,8566 | 0,0674 | 0,7510 | 0,0210 | 2,0000 |
| 4 | 3,0000 | 0,4750 | 0,0454 | 0,5258 | 6,4000e-3 | 3,0000 |
| 5 | 4,0000 | 0,4282 | 9,0000e-4 | 0,5069 | 0,0115 | 4,0000 |
| 6 | 5,0000 | 0,4520 | 0,0103 | 0,5481 | 0,0332 | 5,0000 |
| 7 | 6,0000 | 0,3525 | 0,0237 | 0,2544 | 8,5420e-3 | 6,0000 |
| 8 | | | | | | |
| 9 | | | | | | |
| 10 | | | | | | |
| 11 | | | | | | |
| 12 | | | | | | |
| 13 | | | | | | |
| 14 | | | | | | |
| 15 | | | | | | |
| 16 | | | | | | |
| 17 | | | | | | |
| 18 | | | | | | |
| 19 | | | | | | |
| 20 | | | | | | |
| 21 | | | | | | |
| 22 | | | | | | |
| 23 | | | | | | |
| 24 | | | | | | |
| 25 | | | | | | |
| 26 | | | | | | |
| 27 | | | | | | |
| 28 | | | | | | |
| 29 | | | | | | |
| 30 | | | | | | |
| 31 | | | | | | |
| 32 | | | | | | |
| 33 | | | | | | |
| 34 | | | | | | |
| 35 | | | | | | |
| 36 | | | | | | |
| 37 | | | | | | |
| 38 | | | | | | |
| 39 | | | | | | |
| 40 | | | | | | |
| 41 | | | | | | |
| 42 | | | | | | |
| 43 | | | | | | |
| 44 | | | | | | |
| 45 | | | | | | |
| 46 | | | | | | |
| 47 | | | | | | |
| 48 | | | | | | |
| 49 | | | | | | |

| | 7-x column | 8-y column | 9 | 10-SumSq | 11-Iterations | 12-a-Final |
|---|---|---|---|---|---|---|
| 1 | 0,2500 | 1,0434 | | 0,0411 | 7,0000 | 1,0972 |
| 2 | 0,2725 | 1,0387 | | 0,0411 | 7,0000 | 1,0972 |
| 3 | 0,2949 | 1,0341 | | 0,0411 | 8,0000 | 1,0972 |
| 4 | 0,3174 | 1,0294 | | 0,0411 | 8,0000 | 1,0972 |
| 5 | 0,3398 | 1,0248 | | 0,0411 | 8,0000 | 1,0972 |
| 6 | 0,3623 | 1,0202 | | 0,0411 | 10,0000 | 1,0972 |
| 7 | 0,3848 | 1,0156 | | 0,0411 | 7,0000 | 1,0972 |
| 8 | 0,4072 | 1,0110 | | 0,0411 | 7,0000 | 1,0972 |
| 9 | 0,4297 | 1,0064 | | 0,0411 | 10,0000 | 1,0972 |
| 10 | 0,4521 | 1,0019 | | 0,0411 | 7,0000 | 1,0972 |
| 11 | 0,4746 | 0,9974 | | 0,0411 | 8,0000 | 1,0972 |
| 12 | 0,4971 | 0,9929 | | 0,0411 | 7,0000 | 1,0972 |
| 13 | 0,5195 | 0,9884 | | 0,0411 | 15,0000 | 1,0972 |
| 14 | 0,5420 | 0,9840 | | 0,0411 | 7,0000 | 1,0972 |
| 15 | 0,5645 | 0,9796 | | 0,0411 | 10,0000 | 1,0972 |
| 16 | 0,5869 | 0,9752 | | 0,0411 | 8,0000 | 1,0972 |
| 17 | 0,6094 | 0,9708 | | 0,0411 | 8,0000 | 1,0972 |
| 18 | 0,6318 | 0,9664 | | 0,0411 | 8,0000 | 1,0972 |
| 19 | 0,6543 | 0,9620 | | 0,0411 | 8,0000 | 1,0972 |
| 20 | 0,6768 | 0,9577 | | 0,0411 | 7,0000 | 1,0972 |
| 21 | 0,6992 | 0,9534 | | 0,0411 | 12,0000 | 1,0972 |
| 22 | 0,7217 | 0,9491 | | 0,0411 | 8,0000 | 1,0972 |
| 23 | 0,7441 | 0,9448 | | 0,0411 | 8,0000 | 1,0972 |
| 24 | 0,7666 | 0,9406 | | 0,0411 | 7,0000 | 1,0972 |
| 25 | 0,7891 | 0,9363 | | 0,0411 | 8,0000 | 1,0972 |
| 26 | 0,8115 | 0,9321 | | 0,0411 | 10,0000 | 1,0972 |
| 27 | 0,8340 | 0,9279 | | 0,0411 | 8,0000 | 1,0972 |
| 28 | 0,8564 | 0,9238 | | 0,0411 | 8,0000 | 1,0972 |
| 29 | 0,8789 | 0,9196 | | 0,0411 | 10,0000 | 1,0972 |
| 30 | 0,9014 | 0,9155 | | 0,0411 | 10,0000 | 1,0972 |
| 31 | 0,9238 | 0,9113 | | 0,0411 | 8,0000 | 1,0972 |
| 32 | 0,9463 | 0,9072 | | 0,0411 | 8,0000 | 1,0972 |
| 33 | 0,9688 | 0,9031 | | 0,0411 | 8,0000 | 1,0972 |
| 34 | 0,9912 | 0,8991 | | 0,0411 | 10,0000 | 1,0972 |
| 35 | 1,0137 | 0,8950 | | 0,0411 | 7,0000 | 1,0972 |
| 36 | 1,0361 | 0,8910 | | 0,0411 | 7,0000 | 1,0972 |
| 37 | 1,0586 | 0,8870 | | 0,0411 | 8,0000 | 1,0972 |
| 38 | 1,0811 | 0,8830 | | 0,0411 | 7,0000 | 1,0972 |
| 39 | 1,1035 | 0,8790 | | 0,0411 | 7,0000 | 1,0972 |
| 40 | 1,1260 | 0,8751 | | 0,0411 | 7,0000 | 1,0972 |
| 41 | 1,1484 | 0,8711 | | 0,0411 | 8,0000 | 1,0972 |
| 42 | 1,1709 | 0,8672 | | 0,0411 | 10,0000 | 1,0972 |
| 43 | 1,1934 | 0,8633 | | 0,0411 | 10,0000 | 1,0972 |
| 44 | 1,2158 | 0,8594 | | 0,0411 | 8,0000 | 1,0972 |
| 45 | 1,2383 | 0,8555 | | 0,0411 | 10,0000 | 1,0972 |
| 46 | 1,2607 | 0,8517 | | 0,0411 | 10,0000 | 1,0972 |
| 47 | 1,2832 | 0,8479 | | 0,0411 | 7,0000 | 1,0972 |
| 48 | 1,3057 | 0,8440 | | 0,0411 | 12,0000 | 1,0972 |
| 49 | 1,3281 | 0,8402 | | 0,0411 | 7,0000 | 1,0972 |

| | 13-b-Final | 14-x column 1 | 15-y column 1 | 16 | 17-SumSq 1 | 18-Iterations 1 |
|---|---|---|---|---|---|---|
| 1 | 0,2009 | 0,2500 | 1,0253 | | 0,0352 | 8,0000 |
| 2 | 0,2009 | 0,2725 | 1,0210 | | 0,0352 | 7,0000 |
| 3 | 0,2009 | 0,2949 | 1,0166 | | 0,0352 | 7,0000 |
| 4 | 0,2009 | 0,3174 | 1,0123 | | 0,0352 | 7,0000 |
| 5 | 0,2009 | 0,3398 | 1,0080 | | 0,0352 | 7,0000 |
| 6 | 0,2009 | 0,3623 | 1,0037 | | 0,0352 | 7,0000 |
| 7 | 0,2009 | 0,3848 | 0,9994 | | 0,0352 | 7,0000 |
| 8 | 0,2009 | 0,4072 | 0,9952 | | 0,0352 | 10,0000 |
| 9 | 0,2009 | 0,4297 | 0,9909 | | 0,0352 | 7,0000 |
| 10 | 0,2009 | 0,4521 | 0,9867 | | 0,0352 | 7,0000 |
| 11 | 0,2009 | 0,4746 | 0,9825 | | 0,0352 | 7,0000 |
| 12 | 0,2009 | 0,4971 | 0,9784 | | 0,0352 | 7,0000 |
| 13 | 0,2009 | 0,5195 | 0,9742 | | 0,0352 | 10,0000 |
| 14 | 0,2009 | 0,5420 | 0,9701 | | 0,0352 | 7,0000 |
| 15 | 0,2009 | 0,5645 | 0,9659 | | 0,0352 | 7,0000 |
| 16 | 0,2009 | 0,5869 | 0,9618 | | 0,0352 | 10,0000 |
| 17 | 0,2009 | 0,6094 | 0,9577 | | 0,0352 | 7,0000 |
| 18 | 0,2009 | 0,6318 | 0,9537 | | 0,0352 | 10,0000 |
| 19 | 0,2009 | 0,6543 | 0,9496 | | 0,0352 | 7,0000 |
| 20 | 0,2009 | 0,6768 | 0,9456 | | 0,0352 | 7,0000 |
| 21 | 0,2009 | 0,6992 | 0,9415 | | 0,0352 | 10,0000 |
| 22 | 0,2009 | 0,7217 | 0,9375 | | 0,0352 | 7,0000 |
| 23 | 0,2009 | 0,7441 | 0,9335 | | 0,0352 | 10,0000 |
| 24 | 0,2009 | 0,7666 | 0,9296 | | 0,0352 | 12,0000 |
| 25 | 0,2009 | 0,7891 | 0,9256 | | 0,0352 | 7,0000 |
| 26 | 0,2009 | 0,8115 | 0,9217 | | 0,0352 | 7,0000 |
| 27 | 0,2009 | 0,8340 | 0,9178 | | 0,0352 | 7,0000 |
| 28 | 0,2009 | 0,8564 | 0,9139 | | 0,0352 | 10,0000 |
| 29 | 0,2009 | 0,8789 | 0,9100 | | 0,0352 | 7,0000 |
| 30 | 0,2009 | 0,9014 | 0,9061 | | 0,0352 | 7,0000 |
| 31 | 0,2009 | 0,9238 | 0,9023 | | 0,0352 | 10,0000 |
| 32 | 0,2009 | 0,9463 | 0,8984 | | 0,0352 | 10,0000 |
| 33 | 0,2009 | 0,9688 | 0,8946 | | 0,0352 | 7,0000 |
| 34 | 0,2009 | 0,9912 | 0,8908 | | 0,0352 | 7,0000 |
| 35 | 0,2009 | 1,0137 | 0,8870 | | 0,0352 | 7,0000 |
| 36 | 0,2009 | 1,0361 | 0,8832 | | 0,0352 | 10,0000 |
| 37 | 0,2009 | 1,0586 | 0,8795 | | 0,0352 | 7,0000 |
| 38 | 0,2009 | 1,0811 | 0,8757 | | 0,0352 | 7,0000 |
| 39 | 0,2009 | 1,1035 | 0,8720 | | 0,0352 | 8,0000 |
| 40 | 0,2009 | 1,1260 | 0,8683 | | 0,0352 | 7,0000 |
| 41 | 0,2009 | 1,1484 | 0,8646 | | 0,0352 | 7,0000 |
| 42 | 0,2009 | 1,1709 | 0,8609 | | 0,0352 | 8,0000 |
| 43 | 0,2009 | 1,1934 | 0,8573 | | 0,0352 | 7,0000 |
| 44 | 0,2009 | 1,2158 | 0,8536 | | 0,0352 | 7,0000 |
| 45 | 0,2009 | 1,2383 | 0,8500 | | 0,0352 | 7,0000 |
| 46 | 0,2009 | 1,2607 | 0,8464 | | 0,0352 | 7,0000 |
| 47 | 0,2009 | 1,2832 | 0,8428 | | 0,0352 | 7,0000 |
| 48 | 0,2009 | 1,3057 | 0,8392 | | 0,0352 | 7,0000 |
| 49 | 0,2009 | 1,3281 | 0,8356 | | 0,0352 | 7,0000 |

| | 19-a-Final 1 | 20-b-Final 1 |
|---|---|---|
| 1 | 1,0751 | 0,1897 |
| 2 | 1,0751 | 0,1897 |
| 3 | 1,0751 | 0,1897 |
| 4 | 1,0751 | 0,1897 |
| 5 | 1,0751 | 0,1897 |
| 6 | 1,0751 | 0,1897 |
| 7 | 1,0751 | 0,1897 |
| 8 | 1,0751 | 0,1897 |
| 9 | 1,0751 | 0,1897 |
| 10 | 1,0751 | 0,1897 |
| 11 | 1,0751 | 0,1897 |
| 12 | 1,0751 | 0,1897 |
| 13 | 1,0751 | 0,1897 |
| 14 | 1,0751 | 0,1897 |
| 15 | 1,0751 | 0,1897 |
| 16 | 1,0751 | 0,1897 |
| 17 | 1,0751 | 0,1897 |
| 18 | 1,0751 | 0,1897 |
| 19 | 1,0751 | 0,1897 |
| 20 | 1,0751 | 0,1897 |
| 21 | 1,0751 | 0,1897 |
| 22 | 1,0751 | 0,1897 |
| 23 | 1,0751 | 0,1897 |
| 24 | 1,0751 | 0,1897 |
| 25 | 1,0751 | 0,1897 |
| 26 | 1,0751 | 0,1897 |
| 27 | 1,0751 | 0,1897 |
| 28 | 1,0751 | 0,1897 |
| 29 | 1,0751 | 0,1897 |
| 30 | 1,0751 | 0,1897 |
| 31 | 1,0751 | 0,1897 |
| 32 | 1,0751 | 0,1897 |
| 33 | 1,0751 | 0,1897 |
| 34 | 1,0751 | 0,1897 |
| 35 | 1,0751 | 0,1897 |
| 36 | 1,0751 | 0,1897 |
| 37 | 1,0751 | 0,1897 |
| 38 | 1,0751 | 0,1897 |
| 39 | 1,0751 | 0,1897 |
| 40 | 1,0751 | 0,1897 |
| 41 | 1,0751 | 0,1897 |
| 42 | 1,0751 | 0,1897 |
| 43 | 1,0751 | 0,1897 |
| 44 | 1,0751 | 0,1897 |
| 45 | 1,0751 | 0,1897 |
| 46 | 1,0751 | 0,1897 |
| 47 | 1,0751 | 0,1897 |
| 48 | 1,0751 | 0,1897 |
| 49 | 1,0751 | 0,1897 |

| | 1 | 2 | 3 | 4 | 5 | 6 |
|---|---|---|---|---|---|---|
| 50 | | | | | | |
| 51 | | | | | | |
| 52 | | | | | | |
| 53 | | | | | | |
| 54 | | | | | | |
| 55 | | | | | | |
| 56 | | | | | | |
| 57 | | | | | | |
| 58 | | | | | | |
| 59 | | | | | | |
| 60 | | | | | | |
| 61 | | | | | | |
| 62 | | | | | | |
| 63 | | | | | | |
| 64 | | | | | | |
| 65 | | | | | | |
| 66 | | | | | | |
| 67 | | | | | | |
| 68 | | | | | | |
| 69 | | | | | | |
| 70 | | | | | | |
| 71 | | | | | | |
| 72 | | | | | | |
| 73 | | | | | | |
| 74 | | | | | | |
| 75 | | | | | | |
| 76 | | | | | | |
| 77 | | | | | | |
| 78 | | | | | | |
| 79 | | | | | | |
| 80 | | | | | | |
| 81 | | | | | | |
| 82 | | | | | | |
| 83 | | | | | | |
| 84 | | | | | | |
| 85 | | | | | | |
| 86 | | | | | | |
| 87 | | | | | | |
| 88 | | | | | | |
| 89 | | | | | | |
| 90 | | | | | | |
| 91 | | | | | | |
| 92 | | | | | | |
| 93 | | | | | | |
| 94 | | | | | | |
| 95 | | | | | | |
| 96 | | | | | | |
| 97 | | | | | | |
| 98 | | | | | | |

| | 7-x column | 8-y column | 9 | 10-SumSq | 11-Iterations | 12-a-Final |
|---|---|---|---|---|---|---|
| 50 | 1,3506 | 0,8365 | | 0,0411 | 7,0000 | 1,0972 |
| 51 | 1,3730 | 0,8327 | | 0,0411 | 7,0000 | 1,0972 |
| 52 | 1,3955 | 0,8289 | | 0,0411 | 10,0000 | 1,0972 |
| 53 | 1,4180 | 0,8252 | | 0,0411 | 7,0000 | 1,0972 |
| 54 | 1,4404 | 0,8215 | | 0,0411 | 7,0000 | 1,0972 |
| 55 | 1,4629 | 0,8178 | | 0,0411 | 8,0000 | 1,0972 |
| 56 | 1,4854 | 0,8141 | | 0,0411 | 10,0000 | 1,0972 |
| 57 | 1,5078 | 0,8105 | | 0,0411 | 8,0000 | 1,0972 |
| 58 | 1,5303 | 0,8068 | | 0,0411 | 12,0000 | 1,0972 |
| 59 | 1,5527 | 0,8032 | | 0,0411 | 8,0000 | 1,0972 |
| 60 | 1,5752 | 0,7996 | | 0,0411 | 10,0000 | 1,0972 |
| 61 | 1,5977 | 0,7960 | | 0,0411 | 10,0000 | 1,0972 |
| 62 | 1,6201 | 0,7924 | | 0,0411 | 7,0000 | 1,0972 |
| 63 | 1,6426 | 0,7888 | | 0,0411 | 10,0000 | 1,0972 |
| 64 | 1,6650 | 0,7853 | | 0,0411 | 8,0000 | 1,0972 |
| 65 | 1,6875 | 0,7817 | | 0,0411 | 10,0000 | 1,0972 |
| 66 | 1,7100 | 0,7782 | | 0,0411 | 10,0000 | 1,0972 |
| 67 | 1,7324 | 0,7747 | | 0,0411 | 10,0000 | 1,0972 |
| 68 | 1,7549 | 0,7712 | | 0,0411 | 8,0000 | 1,0972 |
| 69 | 1,7773 | 0,7677 | | 0,0411 | 7,0000 | 1,0972 |
| 70 | 1,7998 | 0,7643 | | 0,0411 | 7,0000 | 1,0972 |
| 71 | 1,8223 | 0,7608 | | 0,0411 | 8,0000 | 1,0972 |
| 72 | 1,8447 | 0,7574 | | 0,0411 | 7,0000 | 1,0972 |
| 73 | 1,8672 | 0,7540 | | 0,0411 | 8,0000 | 1,0972 |
| 74 | 1,8896 | 0,7506 | | 0,0411 | 10,0000 | 1,0972 |
| 75 | 1,9121 | 0,7472 | | 0,0411 | 7,0000 | 1,0972 |
| 76 | 1,9346 | 0,7439 | | 0,0411 | 7,0000 | 1,0972 |
| 77 | 1,9570 | 0,7405 | | 0,0411 | 7,0000 | 1,0972 |
| 78 | 1,9795 | 0,7372 | | 0,0411 | 8,0000 | 1,0972 |
| 79 | 2,0020 | 0,7339 | | 0,0411 | 7,0000 | 1,0972 |
| 80 | 2,0244 | 0,7306 | | 0,0411 | 8,0000 | 1,0972 |
| 81 | 2,0469 | 0,7273 | | 0,0411 | 8,0000 | 1,0972 |
| 82 | 2,0693 | 0,7240 | | 0,0411 | 8,0000 | 1,0972 |
| 83 | 2,0918 | 0,7207 | | 0,0411 | 7,0000 | 1,0972 |
| 84 | 2,1143 | 0,7175 | | 0,0411 | 10,0000 | 1,0972 |
| 85 | 2,1367 | 0,7143 | | 0,0411 | 7,0000 | 1,0972 |
| 86 | 2,1592 | 0,7110 | | 0,0411 | 7,0000 | 1,0972 |
| 87 | 2,1816 | 0,7078 | | 0,0411 | 7,0000 | 1,0972 |
| 88 | 2,2041 | 0,7047 | | 0,0411 | 7,0000 | 1,0972 |
| 89 | 2,2266 | 0,7015 | | 0,0411 | 8,0000 | 1,0972 |
| 90 | 2,2490 | 0,6983 | | 0,0411 | 7,0000 | 1,0972 |
| 91 | 2,2715 | 0,6952 | | 0,0411 | 7,0000 | 1,0972 |
| 92 | 2,2939 | 0,6921 | | 0,0411 | 8,0000 | 1,0972 |
| 93 | 2,3164 | 0,6889 | | 0,0411 | 7,0000 | 1,0972 |
| 94 | 2,3389 | 0,6858 | | 0,0411 | 8,0000 | 1,0972 |
| 95 | 2,3613 | 0,6827 | | 0,0411 | 10,0000 | 1,0972 |
| 96 | 2,3838 | 0,6797 | | 0,0411 | 10,0000 | 1,0972 |
| 97 | 2,4063 | 0,6766 | | 0,0411 | 8,0000 | 1,0972 |
| 98 | 2,4287 | 0,6736 | | 0,0411 | 22,0000 | 1,0972 |

| | 13-b-Final | 14-x column 1 | 15-y column 1 | 16 | 17-SumSq 1 | 18-Iterations 1 |
|---|---|---|---|---|---|---|
| 50 | 0,2009 | 1,3506 | 0,8321 | | 0,0352 | 7,0000 |
| 51 | 0,2009 | 1,3730 | 0,8285 | | 0,0352 | 7,0000 |
| 52 | 0,2009 | 1,3955 | 0,8250 | | 0,0352 | 10,0000 |
| 53 | 0,2009 | 1,4180 | 0,8215 | | 0,0352 | 7,0000 |
| 54 | 0,2009 | 1,4404 | 0,8180 | | 0,0352 | 7,0000 |
| 55 | 0,2009 | 1,4629 | 0,8145 | | 0,0352 | 7,0000 |
| 56 | 0,2009 | 1,4854 | 0,8111 | | 0,0352 | 7,0000 |
| 57 | 0,2009 | 1,5078 | 0,8076 | | 0,0352 | 7,0000 |
| 58 | 0,2009 | 1,5303 | 0,8042 | | 0,0352 | 10,0000 |
| 59 | 0,2009 | 1,5527 | 0,8008 | | 0,0352 | 7,0000 |
| 60 | 0,2009 | 1,5752 | 0,7974 | | 0,0352 | 7,0000 |
| 61 | 0,2009 | 1,5977 | 0,7940 | | 0,0352 | 10,0000 |
| 62 | 0,2009 | 1,6201 | 0,7906 | | 0,0352 | 7,0000 |
| 63 | 0,2009 | 1,6426 | 0,7872 | | 0,0352 | 7,0000 |
| 64 | 0,2009 | 1,6650 | 0,7839 | | 0,0352 | 7,0000 |
| 65 | 0,2009 | 1,6875 | 0,7806 | | 0,0352 | 10,0000 |
| 66 | 0,2009 | 1,7100 | 0,7772 | | 0,0352 | 7,0000 |
| 67 | 0,2009 | 1,7324 | 0,7739 | | 0,0352 | 10,0000 |
| 68 | 0,2009 | 1,7549 | 0,7706 | | 0,0352 | 10,0000 |
| 69 | 0,2009 | 1,7773 | 0,7674 | | 0,0352 | 7,0000 |
| 70 | 0,2009 | 1,7998 | 0,7641 | | 0,0352 | 7,0000 |
| 71 | 0,2009 | 1,8223 | 0,7608 | | 0,0352 | 7,0000 |
| 72 | 0,2009 | 1,8447 | 0,7576 | | 0,0352 | 7,0000 |
| 73 | 0,2009 | 1,8672 | 0,7544 | | 0,0352 | 10,0000 |
| 74 | 0,2009 | 1,8896 | 0,7512 | | 0,0352 | 7,0000 |
| 75 | 0,2009 | 1,9121 | 0,7480 | | 0,0352 | 10,0000 |
| 76 | 0,2009 | 1,9346 | 0,7448 | | 0,0352 | 7,0000 |
| 77 | 0,2009 | 1,9570 | 0,7416 | | 0,0352 | 12,0000 |
| 78 | 0,2009 | 1,9795 | 0,7385 | | 0,0352 | 7,0000 |
| 79 | 0,2009 | 2,0020 | 0,7353 | | 0,0352 | 7,0000 |
| 80 | 0,2009 | 2,0244 | 0,7322 | | 0,0352 | 10,0000 |
| 81 | 0,2009 | 2,0469 | 0,7291 | | 0,0352 | 7,0000 |
| 82 | 0,2009 | 2,0693 | 0,7260 | | 0,0352 | 7,0000 |
| 83 | 0,2009 | 2,0918 | 0,7229 | | 0,0352 | 7,0000 |
| 84 | 0,2009 | 2,1143 | 0,7198 | | 0,0352 | 7,0000 |
| 85 | 0,2009 | 2,1367 | 0,7168 | | 0,0352 | 7,0000 |
| 86 | 0,2009 | 2,1592 | 0,7137 | | 0,0352 | 7,0000 |
| 87 | 0,2009 | 2,1816 | 0,7107 | | 0,0352 | 7,0000 |
| 88 | 0,2009 | 2,2041 | 0,7077 | | 0,0352 | 7,0000 |
| 89 | 0,2009 | 2,2266 | 0,7047 | | 0,0352 | 7,0000 |
| 90 | 0,2009 | 2,2490 | 0,7017 | | 0,0352 | 7,0000 |
| 91 | 0,2009 | 2,2715 | 0,6987 | | 0,0352 | 10,0000 |
| 92 | 0,2009 | 2,2939 | 0,6957 | | 0,0352 | 10,0000 |
| 93 | 0,2009 | 2,3164 | 0,6928 | | 0,0352 | 7,0000 |
| 94 | 0,2009 | 2,3389 | 0,6898 | | 0,0352 | 7,0000 |
| 95 | 0,2009 | 2,3613 | 0,6869 | | 0,0352 | 7,0000 |
| 96 | 0,2009 | 2,3838 | 0,6840 | | 0,0352 | 7,0000 |
| 97 | 0,2009 | 2,4063 | 0,6810 | | 0,0352 | 7,0000 |
| 98 | 0,2009 | 2,4287 | 0,6781 | | 0,0352 | 8,0000 |

| | 19-a-Final 1 | 20-b-Final 1 |
|---|---|---|
| 50 | 1,0751 | 0,1897 |
| 51 | 1,0751 | 0,1897 |
| 52 | 1,0751 | 0,1897 |
| 53 | 1,0751 | 0,1897 |
| 54 | 1,0751 | 0,1897 |
| 55 | 1,0751 | 0,1897 |
| 56 | 1,0751 | 0,1897 |
| 57 | 1,0751 | 0,1897 |
| 58 | 1,0751 | 0,1897 |
| 59 | 1,0751 | 0,1897 |
| 60 | 1,0751 | 0,1897 |
| 61 | 1,0751 | 0,1897 |
| 62 | 1,0751 | 0,1897 |
| 63 | 1,0751 | 0,1897 |
| 64 | 1,0751 | 0,1897 |
| 65 | 1,0751 | 0,1897 |
| 66 | 1,0751 | 0,1897 |
| 67 | 1,0751 | 0,1897 |
| 68 | 1,0751 | 0,1897 |
| 69 | 1,0751 | 0,1897 |
| 70 | 1,0751 | 0,1897 |
| 71 | 1,0751 | 0,1897 |
| 72 | 1,0751 | 0,1897 |
| 73 | 1,0751 | 0,1897 |
| 74 | 1,0751 | 0,1897 |
| 75 | 1,0751 | 0,1897 |
| 76 | 1,0751 | 0,1897 |
| 77 | 1,0751 | 0,1897 |
| 78 | 1,0751 | 0,1897 |
| 79 | 1,0751 | 0,1897 |
| 80 | 1,0751 | 0,1897 |
| 81 | 1,0751 | 0,1897 |
| 82 | 1,0751 | 0,1897 |
| 83 | 1,0751 | 0,1897 |
| 84 | 1,0751 | 0,1897 |
| 85 | 1,0751 | 0,1897 |
| 86 | 1,0751 | 0,1897 |
| 87 | 1,0751 | 0,1897 |
| 88 | 1,0751 | 0,1897 |
| 89 | 1,0751 | 0,1897 |
| 90 | 1,0751 | 0,1897 |
| 91 | 1,0751 | 0,1897 |
| 92 | 1,0751 | 0,1897 |
| 93 | 1,0751 | 0,1897 |
| 94 | 1,0751 | 0,1897 |
| 95 | 1,0751 | 0,1897 |
| 96 | 1,0751 | 0,1897 |
| 97 | 1,0751 | 0,1897 |
| 98 | 1,0751 | 0,1897 |

| | 1 | 2 | 3 | 4 | 5 | 6 |
|---|---|---|---|---|---|---|
| 99 | | | | | | |
| 100 | | | | | | |
| 101 | | | | | | |
| 102 | | | | | | |
| 103 | | | | | | |
| 104 | | | | | | |
| 105 | | | | | | |
| 106 | | | | | | |
| 107 | | | | | | |
| 108 | | | | | | |
| 109 | | | | | | |
| 110 | | | | | | |
| 111 | | | | | | |
| 112 | | | | | | |
| 113 | | | | | | |
| 114 | | | | | | |
| 115 | | | | | | |
| 116 | | | | | | |
| 117 | | | | | | |
| 118 | | | | | | |
| 119 | | | | | | |
| 120 | | | | | | |
| 121 | | | | | | |
| 122 | | | | | | |
| 123 | | | | | | |
| 124 | | | | | | |
| 125 | | | | | | |
| 126 | | | | | | |
| 127 | | | | | | |
| 128 | | | | | | |
| 129 | | | | | | |
| 130 | | | | | | |
| 131 | | | | | | |
| 132 | | | | | | |
| 133 | | | | | | |
| 134 | | | | | | |
| 135 | | | | | | |
| 136 | | | | | | |
| 137 | | | | | | |
| 138 | | | | | | |
| 139 | | | | | | |
| 140 | | | | | | |
| 141 | | | | | | |
| 142 | | | | | | |
| 143 | | | | | | |
| 144 | | | | | | |
| 145 | | | | | | |
| 146 | | | | | | |
| 147 | | | | | | |

| | 7-x column | 8-y column | 9 | 10-SumSq | 11-Iterations | 12-a-Final |
|---|---|---|---|---|---|---|
| 99 | 2,4512 | 0,6705 | | 0,0411 | 7,0000 | 1,0972 |
| 100 | 2,4736 | 0,6675 | | 0,0411 | 10,0000 | 1,0972 |
| 101 | 2,4961 | 0,6645 | | 0,0411 | 8,0000 | 1,0972 |
| 102 | 2,5186 | 0,6615 | | 0,0411 | 7,0000 | 1,0972 |
| 103 | 2,5410 | 0,6585 | | 0,0411 | 10,0000 | 1,0972 |
| 104 | 2,5635 | 0,6556 | | 0,0411 | 7,0000 | 1,0972 |
| 105 | 2,5859 | 0,6526 | | 0,0411 | 7,0000 | 1,0972 |
| 106 | 2,6084 | 0,6497 | | 0,0411 | 10,0000 | 1,0972 |
| 107 | 2,6309 | 0,6468 | | 0,0411 | 7,0000 | 1,0972 |
| 108 | 2,6533 | 0,6439 | | 0,0411 | 7,0000 | 1,0972 |
| 109 | 2,6758 | 0,6410 | | 0,0411 | 7,0000 | 1,0972 |
| 110 | 2,6982 | 0,6381 | | 0,0411 | 10,0000 | 1,0972 |
| 111 | 2,7207 | 0,6352 | | 0,0411 | 10,0000 | 1,0972 |
| 112 | 2,7432 | 0,6323 | | 0,0411 | 10,0000 | 1,0972 |
| 113 | 2,7656 | 0,6295 | | 0,0411 | 7,0000 | 1,0972 |
| 114 | 2,7881 | 0,6267 | | 0,0411 | 7,0000 | 1,0972 |
| 115 | 2,8105 | 0,6238 | | 0,0411 | 8,0000 | 1,0972 |
| 116 | 2,8330 | 0,6210 | | 0,0411 | 7,0000 | 1,0972 |
| 117 | 2,8555 | 0,6182 | | 0,0411 | 10,0000 | 1,0972 |
| 118 | 2,8779 | 0,6154 | | 0,0411 | 10,0000 | 1,0972 |
| 119 | 2,9004 | 0,6127 | | 0,0411 | 7,0000 | 1,0972 |
| 120 | 2,9229 | 0,6099 | | 0,0411 | 7,0000 | 1,0972 |
| 121 | 2,9453 | 0,6072 | | 0,0411 | 7,0000 | 1,0972 |
| 122 | 2,9678 | 0,6044 | | 0,0411 | 7,0000 | 1,0972 |
| 123 | 2,9902 | 0,6017 | | 0,0411 | 7,0000 | 1,0972 |
| 124 | 3,0127 | 0,5990 | | 0,0411 | 7,0000 | 1,0972 |
| 125 | 3,0352 | 0,5963 | | 0,0411 | 10,0000 | 1,0972 |
| 126 | 3,0576 | 0,5936 | | 0,0411 | 8,0000 | 1,0972 |
| 127 | 3,0801 | 0,5910 | | 0,0411 | 8,0000 | 1,0972 |
| 128 | 3,1025 | 0,5883 | | 0,0411 | 7,0000 | 1,0972 |
| 129 | 3,1250 | 0,5856 | | 0,0411 | 10,0000 | 1,0972 |
| 130 | 3,1475 | 0,5830 | | 0,0411 | 7,0000 | 1,0972 |
| 131 | 3,1699 | 0,5804 | | 0,0411 | 7,0000 | 1,0972 |
| 132 | 3,1924 | 0,5778 | | 0,0411 | 7,0000 | 1,0972 |
| 133 | 3,2148 | 0,5752 | | 0,0411 | 8,0000 | 1,0972 |
| 134 | 3,2373 | 0,5726 | | 0,0411 | 10,0000 | 1,0972 |
| 135 | 3,2598 | 0,5700 | | 0,0411 | 7,0000 | 1,0972 |
| 136 | 3,2822 | 0,5674 | | 0,0411 | 7,0000 | 1,0972 |
| 137 | 3,3047 | 0,5649 | | 0,0411 | 33,0000 | 1,0972 |
| 138 | 3,3271 | 0,5623 | | 0,0411 | 8,0000 | 1,0972 |
| 139 | 3,3496 | 0,5598 | | 0,0411 | 7,0000 | 1,0972 |
| 140 | 3,3721 | 0,5573 | | 0,0411 | 7,0000 | 1,0972 |
| 141 | 3,3945 | 0,5548 | | 0,0411 | 8,0000 | 1,0972 |
| 142 | 3,4170 | 0,5523 | | 0,0411 | 7,0000 | 1,0972 |
| 143 | 3,4395 | 0,5498 | | 0,0411 | 8,0000 | 1,0972 |
| 144 | 3,4619 | 0,5473 | | 0,0411 | 7,0000 | 1,0972 |
| 145 | 3,4844 | 0,5449 | | 0,0411 | 7,0000 | 1,0972 |
| 146 | 3,5068 | 0,5424 | | 0,0411 | 7,0000 | 1,0972 |
| 147 | 3,5293 | 0,5400 | | 0,0411 | 8,0000 | 1,0972 |

| | 13-b-Final | 14-x column 1 | 15-y column 1 | 16 | 17-SumSq 1 | 18-Iterations 1 |
|---|---|---|---|---|---|---|
| 99 | 0,2009 | 2,4512 | 0,6753 | | 0,0352 | 7,0000 |
| 100 | 0,2009 | 2,4736 | 0,6724 | | 0,0352 | 19,0000 |
| 101 | 0,2009 | 2,4961 | 0,6695 | | 0,0352 | 10,0000 |
| 102 | 0,2009 | 2,5186 | 0,6667 | | 0,0352 | 10,0000 |
| 103 | 0,2009 | 2,5410 | 0,6638 | | 0,0352 | 7,0000 |
| 104 | 0,2009 | 2,5635 | 0,6610 | | 0,0352 | 7,0000 |
| 105 | 0,2009 | 2,5859 | 0,6582 | | 0,0352 | 7,0000 |
| 106 | 0,2009 | 2,6084 | 0,6554 | | 0,0352 | 10,0000 |
| 107 | 0,2009 | 2,6309 | 0,6526 | | 0,0352 | 7,0000 |
| 108 | 0,2009 | 2,6533 | 0,6499 | | 0,0352 | 7,0000 |
| 109 | 0,2009 | 2,6758 | 0,6471 | | 0,0352 | 10,0000 |
| 110 | 0,2009 | 2,6982 | 0,6443 | | 0,0352 | 7,0000 |
| 111 | 0,2009 | 2,7207 | 0,6416 | | 0,0352 | 7,0000 |
| 112 | 0,2009 | 2,7432 | 0,6389 | | 0,0352 | 10,0000 |
| 113 | 0,2009 | 2,7656 | 0,6362 | | 0,0352 | 7,0000 |
| 114 | 0,2009 | 2,7881 | 0,6334 | | 0,0352 | 8,0000 |
| 115 | 0,2009 | 2,8105 | 0,6308 | | 0,0352 | 7,0000 |
| 116 | 0,2009 | 2,8330 | 0,6281 | | 0,0352 | 7,0000 |
| 117 | 0,2009 | 2,8555 | 0,6254 | | 0,0352 | 7,0000 |
| 118 | 0,2009 | 2,8779 | 0,6227 | | 0,0352 | 7,0000 |
| 119 | 0,2009 | 2,9004 | 0,6201 | | 0,0352 | 10,0000 |
| 120 | 0,2009 | 2,9229 | 0,6175 | | 0,0352 | 7,0000 |
| 121 | 0,2009 | 2,9453 | 0,6148 | | 0,0352 | 7,0000 |
| 122 | 0,2009 | 2,9678 | 0,6122 | | 0,0352 | 7,0000 |
| 123 | 0,2009 | 2,9902 | 0,6096 | | 0,0352 | 7,0000 |
| 124 | 0,2009 | 3,0127 | 0,6070 | | 0,0352 | 7,0000 |
| 125 | 0,2009 | 3,0352 | 0,6044 | | 0,0352 | 7,0000 |
| 126 | 0,2009 | 3,0576 | 0,6019 | | 0,0352 | 7,0000 |
| 127 | 0,2009 | 3,0801 | 0,5993 | | 0,0352 | 10,0000 |
| 128 | 0,2009 | 3,1025 | 0,5968 | | 0,0352 | 7,0000 |
| 129 | 0,2009 | 3,1250 | 0,5942 | | 0,0352 | 7,0000 |
| 130 | 0,2009 | 3,1475 | 0,5917 | | 0,0352 | 7,0000 |
| 131 | 0,2009 | 3,1699 | 0,5892 | | 0,0352 | 7,0000 |
| 132 | 0,2009 | 3,1924 | 0,5867 | | 0,0352 | 8,0000 |
| 133 | 0,2009 | 3,2148 | 0,5842 | | 0,0352 | 7,0000 |
| 134 | 0,2009 | 3,2373 | 0,5817 | | 0,0352 | 7,0000 |
| 135 | 0,2009 | 3,2598 | 0,5792 | | 0,0352 | 7,0000 |
| 136 | 0,2009 | 3,2822 | 0,5768 | | 0,0352 | 7,0000 |
| 137 | 0,2009 | 3,3047 | 0,5743 | | 0,0352 | 10,0000 |
| 138 | 0,2009 | 3,3271 | 0,5719 | | 0,0352 | 7,0000 |
| 139 | 0,2009 | 3,3496 | 0,5694 | | 0,0352 | 7,0000 |
| 140 | 0,2009 | 3,3721 | 0,5670 | | 0,0352 | 7,0000 |
| 141 | 0,2009 | 3,3945 | 0,5646 | | 0,0352 | 8,0000 |
| 142 | 0,2009 | 3,4170 | 0,5622 | | 0,0352 | 8,0000 |
| 143 | 0,2009 | 3,4395 | 0,5598 | | 0,0352 | 7,0000 |
| 144 | 0,2009 | 3,4619 | 0,5574 | | 0,0352 | 8,0000 |
| 145 | 0,2009 | 3,4844 | 0,5551 | | 0,0352 | 7,0000 |
| 146 | 0,2009 | 3,5068 | 0,5527 | | 0,0352 | 15,0000 |
| 147 | 0,2009 | 3,5293 | 0,5503 | | 0,0352 | 7,0000 |

| | 19-a-Final 1 | 20-b-Final 1 |
|---|---|---|
| 99 | 1,0751 | 0,1897 |
| 100 | 1,0751 | 0,1897 |
| 101 | 1,0751 | 0,1897 |
| 102 | 1,0751 | 0,1897 |
| 103 | 1,0751 | 0,1897 |
| 104 | 1,0751 | 0,1897 |
| 105 | 1,0751 | 0,1897 |
| 106 | 1,0751 | 0,1897 |
| 107 | 1,0751 | 0,1897 |
| 108 | 1,0751 | 0,1897 |
| 109 | 1,0751 | 0,1897 |
| 110 | 1,0751 | 0,1897 |
| 111 | 1,0751 | 0,1897 |
| 112 | 1,0751 | 0,1897 |
| 113 | 1,0751 | 0,1897 |
| 114 | 1,0751 | 0,1897 |
| 115 | 1,0751 | 0,1897 |
| 116 | 1,0751 | 0,1897 |
| 117 | 1,0751 | 0,1897 |
| 118 | 1,0751 | 0,1897 |
| 119 | 1,0751 | 0,1897 |
| 120 | 1,0751 | 0,1897 |
| 121 | 1,0751 | 0,1897 |
| 122 | 1,0751 | 0,1897 |
| 123 | 1,0751 | 0,1897 |
| 124 | 1,0751 | 0,1897 |
| 125 | 1,0751 | 0,1897 |
| 126 | 1,0751 | 0,1897 |
| 127 | 1,0751 | 0,1897 |
| 128 | 1,0751 | 0,1897 |
| 129 | 1,0751 | 0,1897 |
| 130 | 1,0751 | 0,1897 |
| 131 | 1,0751 | 0,1897 |
| 132 | 1,0751 | 0,1897 |
| 133 | 1,0751 | 0,1897 |
| 134 | 1,0751 | 0,1897 |
| 135 | 1,0751 | 0,1897 |
| 136 | 1,0751 | 0,1897 |
| 137 | 1,0751 | 0,1897 |
| 138 | 1,0751 | 0,1897 |
| 139 | 1,0751 | 0,1897 |
| 140 | 1,0751 | 0,1897 |
| 141 | 1,0751 | 0,1897 |
| 142 | 1,0751 | 0,1897 |
| 143 | 1,0751 | 0,1897 |
| 144 | 1,0751 | 0,1897 |
| 145 | 1,0751 | 0,1897 |
| 146 | 1,0751 | 0,1897 |
| 147 | 1,0751 | 0,1897 |

| | 1 | 2 | 3 | 4 | 5 | 6 |
|---|---|---|---|---|---|---|
| 148 | | | | | | |
| 149 | | | | | | |
| 150 | | | | | | |
| 151 | | | | | | |
| 152 | | | | | | |
| 153 | | | | | | |
| 154 | | | | | | |
| 155 | | | | | | |
| 156 | | | | | | |
| 157 | | | | | | |
| 158 | | | | | | |
| 159 | | | | | | |
| 160 | | | | | | |
| 161 | | | | | | |
| 162 | | | | | | |
| 163 | | | | | | |
| 164 | | | | | | |
| 165 | | | | | | |
| 166 | | | | | | |
| 167 | | | | | | |
| 168 | | | | | | |
| 169 | | | | | | |
| 170 | | | | | | |
| 171 | | | | | | |
| 172 | | | | | | |
| 173 | | | | | | |
| 174 | | | | | | |
| 175 | | | | | | |
| 176 | | | | | | |
| 177 | | | | | | |
| 178 | | | | | | |
| 179 | | | | | | |
| 180 | | | | | | |
| 181 | | | | | | |
| 182 | | | | | | |
| 183 | | | | | | |
| 184 | | | | | | |
| 185 | | | | | | |
| 186 | | | | | | |
| 187 | | | | | | |
| 188 | | | | | | |
| 189 | | | | | | |
| 190 | | | | | | |
| 191 | | | | | | |
| 192 | | | | | | |
| 193 | | | | | | |
| 194 | | | | | | |
| 195 | | | | | | |
| 196 | | | | | | |

| | 7-x column | 8-y column | 9 | 10-SumSq | 11-Iterations | 12-a-Final |
|---|---|---|---|---|---|---|
| 148 | 3,5518 | 0,5375 | | 0,0411 | 8,0000 | 1,0972 |
| 149 | 3,5742 | 0,5351 | | 0,0411 | 8,0000 | 1,0972 |
| 150 | 3,5967 | 0,5327 | | 0,0411 | 7,0000 | 1,0972 |
| 151 | 3,6191 | 0,5303 | | 0,0411 | 8,0000 | 1,0972 |
| 152 | 3,6416 | 0,5279 | | 0,0411 | 7,0000 | 1,0972 |
| 153 | 3,6641 | 0,5255 | | 0,0411 | 7,0000 | 1,0972 |
| 154 | 3,6865 | 0,5232 | | 0,0411 | 10,0000 | 1,0972 |
| 155 | 3,7090 | 0,5208 | | 0,0411 | 7,0000 | 1,0972 |
| 156 | 3,7314 | 0,5185 | | 0,0411 | 7,0000 | 1,0972 |
| 157 | 3,7539 | 0,5161 | | 0,0411 | 7,0000 | 1,0972 |
| 158 | 3,7764 | 0,5138 | | 0,0411 | 7,0000 | 1,0972 |
| 159 | 3,7988 | 0,5115 | | 0,0411 | 7,0000 | 1,0972 |
| 160 | 3,8213 | 0,5092 | | 0,0411 | 7,0000 | 1,0972 |
| 161 | 3,8438 | 0,5069 | | 0,0411 | 7,0000 | 1,0972 |
| 162 | 3,8662 | 0,5046 | | 0,0411 | 7,0000 | 1,0972 |
| 163 | 3,8887 | 0,5023 | | 0,0411 | 7,0000 | 1,0972 |
| 164 | 3,9111 | 0,5001 | | 0,0411 | 7,0000 | 1,0972 |
| 165 | 3,9336 | 0,4978 | | 0,0411 | 7,0000 | 1,0972 |
| 166 | 3,9561 | 0,4956 | | 0,0411 | 7,0000 | 1,0972 |
| 167 | 3,9785 | 0,4934 | | 0,0411 | 7,0000 | 1,0972 |
| 168 | 4,0010 | 0,4911 | | 0,0411 | 7,0000 | 1,0972 |
| 169 | 4,0234 | 0,4889 | | 0,0411 | 7,0000 | 1,0972 |
| 170 | 4,0459 | 0,4867 | | 0,0411 | 7,0000 | 1,0972 |
| 171 | 4,0684 | 0,4845 | | 0,0411 | 7,0000 | 1,0972 |
| 172 | 4,0908 | 0,4824 | | 0,0411 | 7,0000 | 1,0972 |
| 173 | 4,1133 | 0,4802 | | 0,0411 | 7,0000 | 1,0972 |
| 174 | 4,1357 | 0,4780 | | 0,0411 | 7,0000 | 1,0972 |
| 175 | 4,1582 | 0,4759 | | 0,0411 | 7,0000 | 1,0972 |
| 176 | 4,1807 | 0,4737 | | 0,0411 | 7,0000 | 1,0972 |
| 177 | 4,2031 | 0,4716 | | 0,0411 | 7,0000 | 1,0972 |
| 178 | 4,2256 | 0,4695 | | 0,0411 | 7,0000 | 1,0972 |
| 179 | 4,2480 | 0,4674 | | 0,0411 | 7,0000 | 1,0972 |
| 180 | 4,2705 | 0,4653 | | 0,0411 | 7,0000 | 1,0972 |
| 181 | 4,2930 | 0,4632 | | 0,0411 | 7,0000 | 1,0972 |
| 182 | 4,3154 | 0,4611 | | 0,0411 | 7,0000 | 1,0972 |
| 183 | 4,3379 | 0,4590 | | 0,0411 | 7,0000 | 1,0972 |
| 184 | 4,3604 | 0,4569 | | 0,0411 | 7,0000 | 1,0972 |
| 185 | 4,3828 | 0,4549 | | 0,0411 | 7,0000 | 1,0972 |
| 186 | 4,4053 | 0,4528 | | 0,0411 | 7,0000 | 1,0972 |
| 187 | 4,4277 | 0,4508 | | 0,0411 | 7,0000 | 1,0972 |
| 188 | 4,4502 | 0,4488 | | 0,0411 | 7,0000 | 1,0972 |
| 189 | 4,4727 | 0,4467 | | 0,0411 | 7,0000 | 1,0972 |
| 190 | 4,4951 | 0,4447 | | 0,0411 | 7,0000 | 1,0972 |
| 191 | 4,5176 | 0,4427 | | 0,0411 | 7,0000 | 1,0972 |
| 192 | 4,5400 | 0,4407 | | 0,0411 | 7,0000 | 1,0972 |
| 193 | 4,5625 | 0,4387 | | 0,0411 | 7,0000 | 1,0972 |
| 194 | 4,5850 | 0,4368 | | 0,0411 | 7,0000 | 1,0972 |
| 195 | 4,6074 | 0,4348 | | 0,0411 | 7,0000 | 1,0972 |
| 196 | 4,6299 | 0,4328 | | 0,0411 | 7,0000 | 1,0972 |

| | 13-b-Final | 14-x column 1 | 15-y column 1 | 16 | 17-SumSq 1 | 18-Iterations 1 |
|---|---|---|---|---|---|---|
| 148 | 0,2009 | 3,5518 | 0,5480 | | 0,0352 | 7,0000 |
| 149 | 0,2009 | 3,5742 | 0,5457 | | 0,0352 | 12,0000 |
| 150 | 0,2009 | 3,5967 | 0,5434 | | 0,0352 | 33,0000 |
| 151 | 0,2009 | 3,6191 | 0,5410 | | 0,0352 | 7,0000 |
| 152 | 0,2009 | 3,6416 | 0,5387 | | 0,0352 | 7,0000 |
| 153 | 0,2009 | 3,6641 | 0,5364 | | 0,0352 | 7,0000 |
| 154 | 0,2009 | 3,6865 | 0,5342 | | 0,0352 | 10,0000 |
| 155 | 0,2009 | 3,7090 | 0,5319 | | 0,0352 | 7,0000 |
| 156 | 0,2009 | 3,7314 | 0,5296 | | 0,0352 | 7,0000 |
| 157 | 0,2009 | 3,7539 | 0,5274 | | 0,0352 | 7,0000 |
| 158 | 0,2009 | 3,7764 | 0,5251 | | 0,0352 | 7,0000 |
| 159 | 0,2009 | 3,7988 | 0,5229 | | 0,0352 | 7,0000 |
| 160 | 0,2009 | 3,8213 | 0,5207 | | 0,0352 | 7,0000 |
| 161 | 0,2009 | 3,8438 | 0,5185 | | 0,0352 | 7,0000 |
| 162 | 0,2009 | 3,8662 | 0,5163 | | 0,0352 | 10,0000 |
| 163 | 0,2009 | 3,8887 | 0,5141 | | 0,0352 | 7,0000 |
| 164 | 0,2009 | 3,9111 | 0,5119 | | 0,0352 | 7,0000 |
| 165 | 0,2009 | 3,9336 | 0,5097 | | 0,0352 | 7,0000 |
| 166 | 0,2009 | 3,9561 | 0,5075 | | 0,0352 | 7,0000 |
| 167 | 0,2009 | 3,9785 | 0,5054 | | 0,0352 | 7,0000 |
| 168 | 0,2009 | 4,0010 | 0,5032 | | 0,0352 | 7,0000 |
| 169 | 0,2009 | 4,0234 | 0,5011 | | 0,0352 | 7,0000 |
| 170 | 0,2009 | 4,0459 | 0,4990 | | 0,0352 | 7,0000 |
| 171 | 0,2009 | 4,0684 | 0,4968 | | 0,0352 | 7,0000 |
| 172 | 0,2009 | 4,0908 | 0,4947 | | 0,0352 | 7,0000 |
| 173 | 0,2009 | 4,1133 | 0,4926 | | 0,0352 | 7,0000 |
| 174 | 0,2009 | 4,1357 | 0,4905 | | 0,0352 | 10,0000 |
| 175 | 0,2009 | 4,1582 | 0,4884 | | 0,0352 | 7,0000 |
| 176 | 0,2009 | 4,1807 | 0,4864 | | 0,0352 | 7,0000 |
| 177 | 0,2009 | 4,2031 | 0,4843 | | 0,0352 | 10,0000 |
| 178 | 0,2009 | 4,2256 | 0,4822 | | 0,0352 | 7,0000 |
| 179 | 0,2009 | 4,2480 | 0,4802 | | 0,0352 | 7,0000 |
| 180 | 0,2009 | 4,2705 | 0,4781 | | 0,0352 | 7,0000 |
| 181 | 0,2009 | 4,2930 | 0,4761 | | 0,0352 | 7,0000 |
| 182 | 0,2009 | 4,3154 | 0,4741 | | 0,0352 | 10,0000 |
| 183 | 0,2009 | 4,3379 | 0,4721 | | 0,0352 | 7,0000 |
| 184 | 0,2009 | 4,3604 | 0,4701 | | 0,0352 | 7,0000 |
| 185 | 0,2009 | 4,3828 | 0,4681 | | 0,0352 | 7,0000 |
| 186 | 0,2009 | 4,4053 | 0,4661 | | 0,0352 | 7,0000 |
| 187 | 0,2009 | 4,4277 | 0,4641 | | 0,0352 | 7,0000 |
| 188 | 0,2009 | 4,4502 | 0,4621 | | 0,0352 | 7,0000 |
| 189 | 0,2009 | 4,4727 | 0,4601 | | 0,0352 | 7,0000 |
| 190 | 0,2009 | 4,4951 | 0,4582 | | 0,0352 | 7,0000 |
| 191 | 0,2009 | 4,5176 | 0,4562 | | 0,0352 | 7,0000 |
| 192 | 0,2009 | 4,5400 | 0,4543 | | 0,0352 | 9,0000 |
| 193 | 0,2009 | 4,5625 | 0,4524 | | 0,0352 | 7,0000 |
| 194 | 0,2009 | 4,5850 | 0,4504 | | 0,0352 | 7,0000 |
| 195 | 0,2009 | 4,6074 | 0,4485 | | 0,0352 | 7,0000 |
| 196 | 0,2009 | 4,6299 | 0,4466 | | 0,0352 | 7,0000 |

| | 19-a-Final 1 | 20-b-Final 1 |
|---|---|---|
| 148 | 1,0751 | 0,1897 |
| 149 | 1,0751 | 0,1897 |
| 150 | 1,0751 | 0,1897 |
| 151 | 1,0751 | 0,1897 |
| 152 | 1,0751 | 0,1897 |
| 153 | 1,0751 | 0,1897 |
| 154 | 1,0751 | 0,1897 |
| 155 | 1,0751 | 0,1897 |
| 156 | 1,0751 | 0,1897 |
| 157 | 1,0751 | 0,1897 |
| 158 | 1,0751 | 0,1897 |
| 159 | 1,0751 | 0,1897 |
| 160 | 1,0751 | 0,1897 |
| 161 | 1,0751 | 0,1897 |
| 162 | 1,0751 | 0,1897 |
| 163 | 1,0751 | 0,1897 |
| 164 | 1,0751 | 0,1897 |
| 165 | 1,0751 | 0,1897 |
| 166 | 1,0751 | 0,1897 |
| 167 | 1,0751 | 0,1897 |
| 168 | 1,0751 | 0,1897 |
| 169 | 1,0751 | 0,1897 |
| 170 | 1,0751 | 0,1897 |
| 171 | 1,0751 | 0,1897 |
| 172 | 1,0751 | 0,1897 |
| 173 | 1,0751 | 0,1897 |
| 174 | 1,0751 | 0,1897 |
| 175 | 1,0751 | 0,1897 |
| 176 | 1,0751 | 0,1897 |
| 177 | 1,0751 | 0,1897 |
| 178 | 1,0751 | 0,1897 |
| 179 | 1,0751 | 0,1897 |
| 180 | 1,0751 | 0,1897 |
| 181 | 1,0751 | 0,1897 |
| 182 | 1,0751 | 0,1897 |
| 183 | 1,0751 | 0,1897 |
| 184 | 1,0751 | 0,1897 |
| 185 | 1,0751 | 0,1897 |
| 186 | 1,0751 | 0,1897 |
| 187 | 1,0751 | 0,1897 |
| 188 | 1,0751 | 0,1897 |
| 189 | 1,0751 | 0,1897 |
| 190 | 1,0751 | 0,1897 |
| 191 | 1,0751 | 0,1897 |
| 192 | 1,0751 | 0,1897 |
| 193 | 1,0751 | 0,1897 |
| 194 | 1,0751 | 0,1897 |
| 195 | 1,0751 | 0,1897 |
| 196 | 1,0751 | 0,1897 |

| | 1 | 2 | 3 | 4 | 5 | 6 |
|---|---|---|---|---|---|---|
| 197 | | | | | | |
| 198 | | | | | | |
| 199 | | | | | | |
| 200 | | | | | | |
| 201 | | | | | | |
| 202 | | | | | | |
| 203 | | | | | | |
| 204 | | | | | | |
| 205 | | | | | | |
| 206 | | | | | | |
| 207 | | | | | | |
| 208 | | | | | | |
| 209 | | | | | | |
| 210 | | | | | | |
| 211 | | | | | | |
| 212 | | | | | | |
| 213 | | | | | | |
| 214 | | | | | | |
| 215 | | | | | | |
| 216 | | | | | | |
| 217 | | | | | | |
| 218 | | | | | | |
| 219 | | | | | | |
| 220 | | | | | | |
| 221 | | | | | | |
| 222 | | | | | | |
| 223 | | | | | | |
| 224 | | | | | | |
| 225 | | | | | | |
| 226 | | | | | | |
| 227 | | | | | | |
| 228 | | | | | | |
| 229 | | | | | | |
| 230 | | | | | | |
| 231 | | | | | | |
| 232 | | | | | | |
| 233 | | | | | | |
| 234 | | | | | | |
| 235 | | | | | | |
| 236 | | | | | | |
| 237 | | | | | | |
| 238 | | | | | | |
| 239 | | | | | | |
| 240 | | | | | | |
| 241 | | | | | | |
| 242 | | | | | | |
| 243 | | | | | | |
| 244 | | | | | | |
| 245 | | | | | | |

| | 7-x column | 8-y column | 9 | 10-SumSq | 11-Iterations | 12-a-Final |
|---|---|---|---|---|---|---|
| 197 | 4,6523 | 0,4309 | | 0,0411 | 7,0000 | 1,0972 |
| 198 | 4,6748 | 0,4290 | | 0,0411 | 7,0000 | 1,0972 |
| 199 | 4,6973 | 0,4270 | | 0,0411 | 7,0000 | 1,0972 |
| 200 | 4,7197 | 0,4251 | | 0,0411 | 6,0000 | 1,0972 |
| 201 | 4,7422 | 0,4232 | | | | |
| 202 | 4,7646 | 0,4213 | | | | |
| 203 | 4,7871 | 0,4194 | | | | |
| 204 | 4,8096 | 0,4175 | | | | |
| 205 | 4,8320 | 0,4156 | | | | |
| 206 | 4,8545 | 0,4138 | | | | |
| 207 | 4,8770 | 0,4119 | | | | |
| 208 | 4,8994 | 0,4100 | | | | |
| 209 | 4,9219 | 0,4082 | | | | |
| 210 | 4,9443 | 0,4064 | | | | |
| 211 | 4,9668 | 0,4045 | | | | |
| 212 | 4,9893 | 0,4027 | | | | |
| 213 | 5,0117 | 0,4009 | | | | |
| 214 | 5,0342 | 0,3991 | | | | |
| 215 | 5,0566 | 0,3973 | | | | |
| 216 | 5,0791 | 0,3955 | | | | |
| 217 | 5,1016 | 0,3937 | | | | |
| 218 | 5,1240 | 0,3919 | | | | |
| 219 | 5,1465 | 0,3902 | | | | |
| 220 | 5,1689 | 0,3884 | | | | |
| 221 | 5,1914 | 0,3867 | | | | |
| 222 | 5,2139 | 0,3849 | | | | |
| 223 | 5,2363 | 0,3832 | | | | |
| 224 | 5,2588 | 0,3815 | | | | |
| 225 | 5,2813 | 0,3798 | | | | |
| 226 | 5,3037 | 0,3780 | | | | |
| 227 | 5,3262 | 0,3763 | | | | |
| 228 | 5,3486 | 0,3747 | | | | |
| 229 | 5,3711 | 0,3730 | | | | |
| 230 | 5,3936 | 0,3713 | | | | |
| 231 | 5,4160 | 0,3696 | | | | |
| 232 | 5,4385 | 0,3680 | | | | |
| 233 | 5,4609 | 0,3663 | | | | |
| 234 | 5,4834 | 0,3646 | | | | |
| 235 | 5,5059 | 0,3630 | | | | |
| 236 | 5,5283 | 0,3614 | | | | |
| 237 | 5,5508 | 0,3597 | | | | |
| 238 | 5,5732 | 0,3581 | | | | |
| 239 | 5,5957 | 0,3565 | | | | |
| 240 | 5,6182 | 0,3549 | | | | |
| 241 | 5,6406 | 0,3533 | | | | |
| 242 | 5,6631 | 0,3517 | | | | |
| 243 | 5,6855 | 0,3501 | | | | |
| 244 | 5,7080 | 0,3486 | | | | |
| 245 | 5,7305 | 0,3470 | | | | |

| | 13-b-Final | 14-x column 1 | 15-y column 1 | 16 | 17-SumSq 1 | 18-Iterations 1 |
|---|---|---|---|---|---|---|
| 197 | 0,2009 | 4,6523 | 0,4447 | | 0,0352 | 7,0000 |
| 198 | 0,2009 | 4,6748 | 0,4428 | | 0,0352 | 7,0000 |
| 199 | 0,2009 | 4,6973 | 0,4409 | | 0,0352 | 7,0000 |
| 200 | 0,2009 | 4,7197 | 0,4391 | | 0,0352 | 6,0000 |
| 201 | | 4,7422 | 0,4372 | | | |
| 202 | | 4,7646 | 0,4353 | | | |
| 203 | | 4,7871 | 0,4335 | | | |
| 204 | | 4,8096 | 0,4317 | | | |
| 205 | | 4,8320 | 0,4298 | | | |
| 206 | | 4,8545 | 0,4280 | | | |
| 207 | | 4,8770 | 0,4262 | | | |
| 208 | | 4,8994 | 0,4244 | | | |
| 209 | | 4,9219 | 0,4226 | | | |
| 210 | | 4,9443 | 0,4208 | | | |
| 211 | | 4,9668 | 0,4190 | | | |
| 212 | | 4,9893 | 0,4172 | | | |
| 213 | | 5,0117 | 0,4154 | | | |
| 214 | | 5,0342 | 0,4136 | | | |
| 215 | | 5,0566 | 0,4119 | | | |
| 216 | | 5,0791 | 0,4101 | | | |
| 217 | | 5,1016 | 0,4084 | | | |
| 218 | | 5,1240 | 0,4067 | | | |
| 219 | | 5,1465 | 0,4049 | | | |
| 220 | | 5,1689 | 0,4032 | | | |
| 221 | | 5,1914 | 0,4015 | | | |
| 222 | | 5,2139 | 0,3998 | | | |
| 223 | | 5,2363 | 0,3981 | | | |
| 224 | | 5,2588 | 0,3964 | | | |
| 225 | | 5,2813 | 0,3947 | | | |
| 226 | | 5,3037 | 0,3930 | | | |
| 227 | | 5,3262 | 0,3914 | | | |
| 228 | | 5,3486 | 0,3897 | | | |
| 229 | | 5,3711 | 0,3880 | | | |
| 230 | | 5,3936 | 0,3864 | | | |
| 231 | | 5,4160 | 0,3847 | | | |
| 232 | | 5,4385 | 0,3831 | | | |
| 233 | | 5,4609 | 0,3815 | | | |
| 234 | | 5,4834 | 0,3798 | | | |
| 235 | | 5,5059 | 0,3782 | | | |
| 236 | | 5,5283 | 0,3766 | | | |
| 237 | | 5,5508 | 0,3750 | | | |
| 238 | | 5,5732 | 0,3734 | | | |
| 239 | | 5,5957 | 0,3718 | | | |
| 240 | | 5,6182 | 0,3703 | | | |
| 241 | | 5,6406 | 0,3687 | | | |
| 242 | | 5,6631 | 0,3671 | | | |
| 243 | | 5,6855 | 0,3656 | | | |
| 244 | | 5,7080 | 0,3640 | | | |
| 245 | | 5,7305 | 0,3625 | | | |

| | 19-a-Final 1 | 20-b-Final 1 |
|---|---|---|
| 197 | 1,0751 | 0,1897 |
| 198 | 1,0751 | 0,1897 |
| 199 | 1,0751 | 0,1897 |
| 200 | 1,0751 | 0,1897 |
| 201 | | |
| 202 | | |
| 203 | | |
| 204 | | |
| 205 | | |
| 206 | | |
| 207 | | |
| 208 | | |
| 209 | | |
| 210 | | |
| 211 | | |
| 212 | | |
| 213 | | |
| 214 | | |
| 215 | | |
| 216 | | |
| 217 | | |
| 218 | | |
| 219 | | |
| 220 | | |
| 221 | | |
| 222 | | |
| 223 | | |
| 224 | | |
| 225 | | |
| 226 | | |
| 227 | | |
| 228 | | |
| 229 | | |
| 230 | | |
| 231 | | |
| 232 | | |
| 233 | | |
| 234 | | |
| 235 | | |
| 236 | | |
| 237 | | |
| 238 | | |
| 239 | | |
| 240 | | |
| 241 | | |
| 242 | | |
| 243 | | |
| 244 | | |
| 245 | | |

| | 1 | 2 | 3 | 4 | 5 | 6 |
|---|---|---|---|---|---|---|
| 246 | | | | | | |
| 247 | | | | | | |
| 248 | | | | | | |
| 249 | | | | | | |
| 250 | | | | | | |
| 251 | | | | | | |
| 252 | | | | | | |
| 253 | | | | | | |
| 254 | | | | | | |
| 255 | | | | | | |
| 256 | | | | | | |
| 257 | | | | | | |

| | 7-x column | 8-y column | 9 | 10-SumSq | 11-Iterations | 12-a-Final |
|---|---|---|---|---|---|---|
| 246 | 5,7529 | 0,3454 | | | | |
| 247 | 5,7754 | 0,3439 | | | | |
| 248 | 5,7979 | 0,3423 | | | | |
| 249 | 5,8203 | 0,3408 | | | | |
| 250 | 5,8428 | 0,3392 | | | | |
| 251 | 5,8652 | 0,3377 | | | | |
| 252 | 5,8877 | 0,3362 | | | | |
| 253 | 5,9102 | 0,3347 | | | | |
| 254 | 5,9326 | 0,3332 | | | | |
| 255 | 5,9551 | 0,3317 | | | | |
| 256 | 5,9775 | 0,3302 | | | | |
| 257 | 6,0000 | 0,3287 | | | | |

| | 13-b-Final | 14-x column 1 | 15-y column 1 | 16 | 17-SumSq 1 | 18-Iterations 1 |
|---|---|---|---|---|---|---|
| 246 | | 5,7529 | 0,3609 | | | |
| 247 | | 5,7754 | 0,3594 | | | |
| 248 | | 5,7979 | 0,3578 | | | |
| 249 | | 5,8203 | 0,3563 | | | |
| 250 | | 5,8428 | 0,3548 | | | |
| 251 | | 5,8652 | 0,3533 | | | |
| 252 | | 5,8877 | 0,3518 | | | |
| 253 | | 5,9102 | 0,3503 | | | |
| 254 | | 5,9326 | 0,3488 | | | |
| 255 | | 5,9551 | 0,3473 | | | |
| 256 | | 5,9775 | 0,3459 | | | |
| 257 | | 6,0000 | 0,3444 | | | |

| | 19-a-Final 1 | 20-b-Final 1 |
|---|---|---|
| 246 | | |
| 247 | | |
| 248 | | |
| 249 | | |
| 250 | | |
| 251 | | |
| 252 | | |
| 253 | | |
| 254 | | |
| 255 | | |
| 256 | | |
| 257 | | |

**Nonlinear Regression - Dynamic Fitting** **czwartek, maj 07, 2026, 15:32:02**

**Data Source: Data 1 in Notebook1**
**Equation: Exponential Decay; Single, 2 Parameter**
f = a*exp(-b*x)

**Dynamic Fit Options:**

| | |
|---|---|
| Total Number of Fits | 200 |
| Maximum Number of Iterations | 200 |

**Parameter Ranges for Initial Estimates:**

| | **Minimum** | **Maximum** |
|---|---|---|
| a | -1,0689 | 3,2067 |
| b | 0,0000 | 0,9356 |

**Summary of Fit Results:**

| | |
|---|---|
| Converged | 100,0% |

**Results for the Overall Best-Fit Solution:**

| **R** | **Rsqr** | **Adj Rsqr** | **Standard Error of Estimate** |
|---|---|---|---|
| 0,9536 | 0,9094 | 0,8913 | 0,0907 |

| | **Coefficient** | **Std. Error** | **t** | **P** |
|---|---|---|---|---|
| a | 1,0972 | 0,0804 | 13,6544 | <0,0001 |
| b | 0,2009 | 0,0313 | 6,4130 | 0,0014 |

**Analysis of Variance:**

Analysis of Variance:

| | **DF** | **SS** | **MS** |
|---|---|---|---|
| Regression | 2 | 3,3047 | 1,6524 |
| Residual | 5 | 0,0411 | 0,0082 |
| Total | 7 | 3,3458 | 0,4780 |

Corrected for the mean of the observations:

| | **DF** | **SS** | **MS** | **F** | **P** |
|---|---|---|---|---|---|
| Regression | 1 | 0,4125 | 0,4125 | 50,2027 | 0,0009 |
| Residual | 5 | 0,0411 | 0,0082 | | |
| Total | 6 | 0,4536 | 0,0756 | | |

**Statistical Tests:**

**Normality Test (Shapiro-Wilk)** Passed (P = 0,9353)

W Statistic= 0,9756 Significance Level = <0,0001

**Constant Variance Test** Passed (P = 0,7810)

**Fit Equation Description:**

```
[Variables]
x = col(1)
y = col(2)
reciprocal_y = 1/abs(y)
reciprocal_ysquare = 1/y^2
'Automatic Initial Parameter Estimate Functions
F(q)=if(size(x)>1; if(total(abs(y))>0; ape(x;log(abs(y));1;0;1); -306); 0)
asign(q)=if(mean(q)>=0;1;-1)
[Parameters]
a = if(F(0)[1]< 307; if(F(0)[1]>-307; asign(y)*10^F(0)[1]; asign(y)*10^(-307)); asign(y)*10^307) "Auto {{previous: 1,09719}} {{MinRange: 0,99}} {{MaxRange: 1}}
b = if(x50(x;y)-min(x)=0; 1;  -ln(,5)/(x50(x;y)-min(x)))  "Auto {{previous: 0,200894}} {{MinRange: 0}} {{MaxRange: 1}}
[Equation]
f = a*exp(-b*x)
fit f to y
"fit f to y with weight reciprocal_y
"fit f to y with weight reciprocal_ysquare
[Constraints]
b>0
[Options]
tolerance=1e-10
stepsize=1
iterations=200
```

Number of Iterations Performed = 7

**Nonlinear Regression - Dynamic Fitting** **czwartek, maj 07, 2026, 15:33:06**

**Data Source: Data 1 in Notebook1**
**Equation: Exponential Decay; Single, 2 Parameter**
f = a*exp(-b*x)

**Dynamic Fit Options:**

| | |
|---|---|
| Total Number of Fits | 200 |
| Maximum Number of Iterations | 200 |

**Parameter Ranges for Initial Estimates:**

| | **Minimum** | **Maximum** |
|---|---|---|
| a | -1,1054 | 3,3162 |
| b | 0,0000 | 0,9042 |

**Summary of Fit Results:**
Converged 100,0%

**Results for the Overall Best-Fit Solution:**

| **R** | **Rsqr** | **Adj Rsqr** | **Standard Error of Estimate** |
|---|---|---|---|
| 0,9568 | 0,9154 | 0,8985 | 0,0838 |

| | **Coefficient** | **Std. Error** | **t** | **P** |
|---|---|---|---|---|
| a | 1,0751 | 0,0735 | 14,6347 | <0,0001 |
| b | 0,1897 | 0,0285 | 6,6657 | 0,0011 |

**Analysis of Variance:**

Analysis of Variance:

| | **DF** | **SS** | **MS** |
|---|---|---|---|
| Regression | 2 | 3,2988 | 1,6494 |
| Residual | 5 | 0,0352 | 0,0070 |
| Total | 7 | 3,3339 | 0,4763 |

Corrected for the mean of the observations:

| | **DF** | **SS** | **MS** | **F** | **P** |
|---|---|---|---|---|---|
| Regression | 1 | 0,3806 | 0,3806 | 54,1365 | 0,0007 |
| Residual | 5 | 0,0352 | 0,0070 | | |
| Total | 6 | 0,4157 | 0,0693 | | |

**Statistical Tests:**

**Normality Test (Shapiro-Wilk)** Passed (P = 0,6973)

W Statistic= 0,9464 Significance Level = <0,0001

**Constant Variance Test** Passed (P = 0,2166)

**Fit Equation Description:**

```
[Variables]
x = col(6)
y = col(4)
reciprocal_y = 1/abs(y)
reciprocal_ysquare = 1/y^2
'Automatic Initial Parameter Estimate Functions
F(q)=if(size(x)>1; if(total(abs(y))>0; ape(x;log(abs(y));1;0;1); -306); 0)
asign(q)=if(mean(q)>=0;1;-1)
[Parameters]
a = if(F(0)[1]< 307; if(F(0)[1]>-307; asign(y)*10^F(0)[1]; asign(y)*10^(-307)); asign(y)*10^307) "Auto {{previous: 1,07512}} {{MinRange: 0,99}} {{MaxRange: 1}}
b = if(x50(x;y)-min(x)=0; 1;  -ln(,5)/(x50(x;y)-min(x)))  "Auto {{previous: 0,189739}} {{MinRange: 0}} {{MaxRange: 1}}
[Equation]
f = a*exp(-b*x)
fit f to y
"fit f to y with weight reciprocal_y
"fit f to y with weight reciprocal_ysquare
[Constraints]
b>0
[Options]
tolerance=1e-10
stepsize=1
iterations=200
```

Number of Iterations Performed = 7

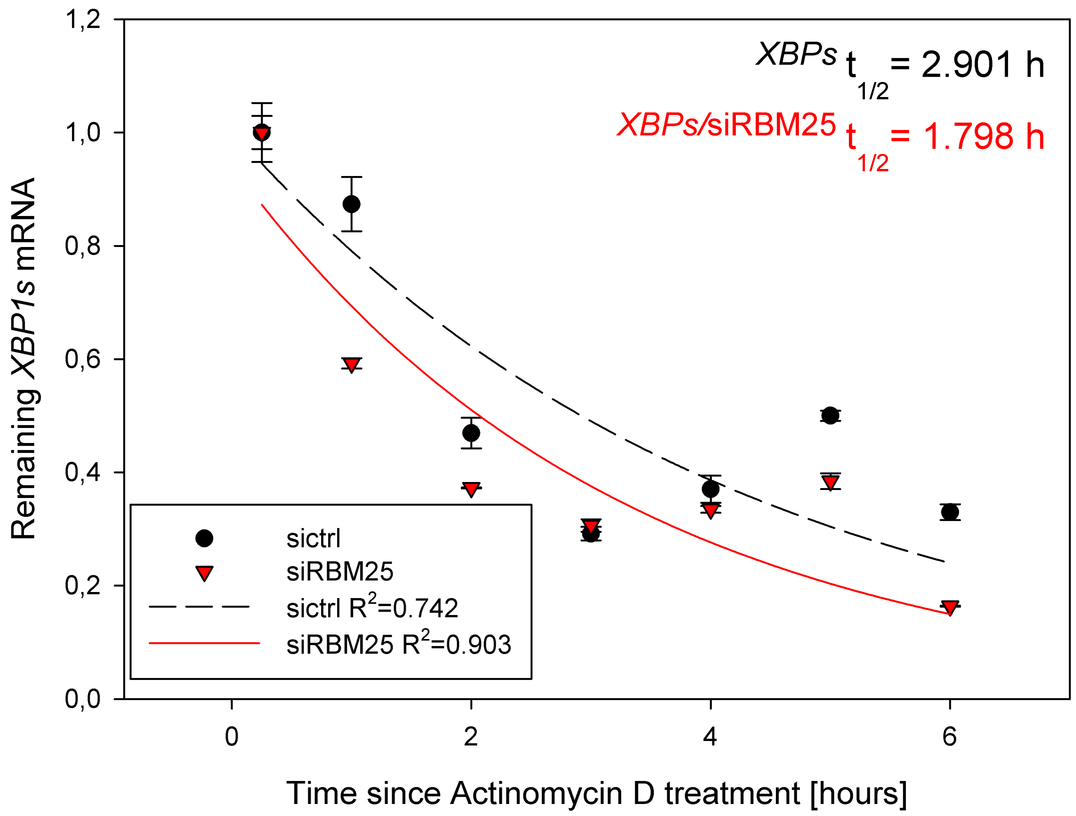

XBPs t1/2 = 2.901 h
XBPs/siRBM25 t1/2 = 1.798 h
Remaining XBP1s mRNA
Time since Actinomycin D treatment [hours]
sictrl
siRBM25
sictrl R2=0.742
siRBM25 R2=0.903
1,2
1,0
0,8
0,6
0,4
0,2
0,0
0
2
4
6

**Nonlinear Regression - Dynamic Fitting** **czwartek, maj 07, 2026, 15:50:15**

**Data Source: Copy of Data 1 in Notebook1 half life calculations.JNB**
**Equation: Exponential Decay; Single, 2 Parameter**
f = a*exp(-b*x)

**Dynamic Fit Options:**

| | |
|---|---|
| Total Number of Fits | 200 |
| Maximum Number of Iterations | 200 |

**Parameter Ranges for Initial Estimates:**

| | **Minimum** | **Maximum** |
|---|---|---|
| a | 0,9900 | 1,0000 |
| b | 0,0000 | 1,0000 |

**Summary of Fit Results:**
Converged 100,0%

**Results for the Overall Best-Fit Solution:**

| **R** | **Rsqr** | **Adj Rsqr** | **Standard Error of Estimate** |
|---|---|---|---|
| 0,8616 | 0,7424 | 0,6909 | 0,1545 |

| | **Coefficient** | **Std. Error** | **t** | **P** |
|---|---|---|---|---|
| a | 1,0036 | 0,1424 | 7,0489 | 0,0009 |
| b | 0,2389 | 0,0664 | 3,5959 | 0,0156 |

**Analysis of Variance:**

Analysis of Variance:

| | **DF** | **SS** | **MS** |
|---|---|---|---|
| Regression | 2 | 2,4448 | 1,2224 |
| Residual | 5 | 0,1194 | 0,0239 |
| Total | 7 | 2,5642 | 0,3663 |

Corrected for the mean of the observations:

| | **DF** | **SS** | **MS** | **F** | **P** |
|---|---|---|---|---|---|
| Regression | 1 | 0,3441 | 0,3441 | 14,4086 | 0,0127 |
| Residual | 5 | 0,1194 | 0,0239 | | |
| Total | 6 | 0,4635 | 0,0773 | | |

**Statistical Tests:**

**Normality Test (Shapiro-Wilk)** Passed (P = 0,5862)

W Statistic= 0,9341 Significance Level = <0,0001

**Constant Variance Test** Passed (P = 0,4907)

**Fit Equation Description:**

```
[Variables]
x = col(1)
y = col(2)
reciprocal_y = 1/abs(y)
reciprocal_ysquare = 1/y^2
'Automatic Initial Parameter Estimate Functions
F(q)=if(size(x)>1; if(total(abs(y))>0; ape(x;log(abs(y));1;0;1); -306); 0)
asign(q)=if(mean(q)>=0;1;-1)
[Parameters]
a = if(F(0)[1]< 307; if(F(0)[1]>-307; asign(y)*10^F(0)[1]; asign(y)*10^(-307)); asign(y)*10^307) "Auto {{previous: 1,00361}} {{MinRange: 0,99}} {{MaxRange: 1}}
b = if(x50(x;y)-min(x)=0; 1;  -ln(,5)/(x50(x;y)-min(x)))  "Auto {{previous: 0,238884}} {{MinRange: 0}} {{MaxRange: 1}}
[Equation]
f = a*exp(-b*x)
fit f to y
"fit f to y with weight reciprocal_y
"fit f to y with weight reciprocal_ysquare
[Constraints]
b>0
[Options]
tolerance=1e-10
stepsize=1
iterations=200
```

Number of Iterations Performed = 11

### Copy of Data 1

| | 1 | 2 | 3 | 4 | 5 | 6 |
|---|---|---|---|---|---|---|
| 1 | 0,2500 | 1,0000 | 0,0520 | 1,0000 | 0,0290 | 0,2500 |
| 2 | 1,0000 | 0,8734 | 0,0480 | 0,5925 | 9,0000e-3 | 1,0000 |
| 3 | 2,0000 | 0,4694 | 0,0270 | 0,3726 | 1,0000e-3 | 2,0000 |
| 4 | 3,0000 | 0,2919 | 0,0120 | 0,3062 | 0,0110 | 3,0000 |
| 5 | 4,0000 | 0,3703 | 0,0240 | 0,3346 | 6,0000e-3 | 4,0000 |
| 6 | 5,0000 | 0,5000 | 9,0000e-3 | 0,3844 | 0,0140 | 5,0000 |
| 7 | 6,0000 | 0,3297 | 0,0140 | 0,1637 | 1,0000e-3 | 6,0000 |
| 8 | | | | | | |
| 9 | | | | | | |
| 10 | | | | | | |
| 11 | | | | | | |
| 12 | | | | | | |
| 13 | | | | | | |
| 14 | | | | | | |
| 15 | | | | | | |
| 16 | | | | | | |
| 17 | | | | | | |
| 18 | | | | | | |
| 19 | | | | | | |
| 20 | | | | | | |
| 21 | | | | | | |
| 22 | | | | | | |
| 23 | | | | | | |
| 24 | | | | | | |
| 25 | | | | | | |
| 26 | | | | | | |
| 27 | | | | | | |
| 28 | | | | | | |
| 29 | | | | | | |
| 30 | | | | | | |
| 31 | | | | | | |
| 32 | | | | | | |
| 33 | | | | | | |
| 34 | | | | | | |
| 35 | | | | | | |
| 36 | | | | | | |
| 37 | | | | | | |
| 38 | | | | | | |
| 39 | | | | | | |
| 40 | | | | | | |
| 41 | | | | | | |
| 42 | | | | | | |
| 43 | | | | | | |
| 44 | | | | | | |
| 45 | | | | | | |
| 46 | | | | | | |
| 47 | | | | | | |
| 48 | | | | | | |
| 49 | | | | | | |

| | 7-x column | 8-y column | 9 | 10-SumSq | 11-Iterations | 12-a-Final |
|---|---|---|---|---|---|---|
| 1 | 0,2500 | 1,0434 | | 0,0411 | 7,0000 | 1,0972 |
| 2 | 0,2725 | 1,0387 | | 0,0411 | 7,0000 | 1,0972 |
| 3 | 0,2949 | 1,0341 | | 0,0411 | 8,0000 | 1,0972 |
| 4 | 0,3174 | 1,0294 | | 0,0411 | 8,0000 | 1,0972 |
| 5 | 0,3398 | 1,0248 | | 0,0411 | 8,0000 | 1,0972 |
| 6 | 0,3623 | 1,0202 | | 0,0411 | 10,0000 | 1,0972 |
| 7 | 0,3848 | 1,0156 | | 0,0411 | 7,0000 | 1,0972 |
| 8 | 0,4072 | 1,0110 | | 0,0411 | 7,0000 | 1,0972 |
| 9 | 0,4297 | 1,0064 | | 0,0411 | 10,0000 | 1,0972 |
| 10 | 0,4521 | 1,0019 | | 0,0411 | 7,0000 | 1,0972 |
| 11 | 0,4746 | 0,9974 | | 0,0411 | 8,0000 | 1,0972 |
| 12 | 0,4971 | 0,9929 | | 0,0411 | 7,0000 | 1,0972 |
| 13 | 0,5195 | 0,9884 | | 0,0411 | 15,0000 | 1,0972 |
| 14 | 0,5420 | 0,9840 | | 0,0411 | 7,0000 | 1,0972 |
| 15 | 0,5645 | 0,9796 | | 0,0411 | 10,0000 | 1,0972 |
| 16 | 0,5869 | 0,9752 | | 0,0411 | 8,0000 | 1,0972 |
| 17 | 0,6094 | 0,9708 | | 0,0411 | 8,0000 | 1,0972 |
| 18 | 0,6318 | 0,9664 | | 0,0411 | 8,0000 | 1,0972 |
| 19 | 0,6543 | 0,9620 | | 0,0411 | 8,0000 | 1,0972 |
| 20 | 0,6768 | 0,9577 | | 0,0411 | 7,0000 | 1,0972 |
| 21 | 0,6992 | 0,9534 | | 0,0411 | 12,0000 | 1,0972 |
| 22 | 0,7217 | 0,9491 | | 0,0411 | 8,0000 | 1,0972 |
| 23 | 0,7441 | 0,9448 | | 0,0411 | 8,0000 | 1,0972 |
| 24 | 0,7666 | 0,9406 | | 0,0411 | 7,0000 | 1,0972 |
| 25 | 0,7891 | 0,9363 | | 0,0411 | 8,0000 | 1,0972 |
| 26 | 0,8115 | 0,9321 | | 0,0411 | 10,0000 | 1,0972 |
| 27 | 0,8340 | 0,9279 | | 0,0411 | 8,0000 | 1,0972 |
| 28 | 0,8564 | 0,9238 | | 0,0411 | 8,0000 | 1,0972 |
| 29 | 0,8789 | 0,9196 | | 0,0411 | 10,0000 | 1,0972 |
| 30 | 0,9014 | 0,9155 | | 0,0411 | 10,0000 | 1,0972 |
| 31 | 0,9238 | 0,9113 | | 0,0411 | 8,0000 | 1,0972 |
| 32 | 0,9463 | 0,9072 | | 0,0411 | 8,0000 | 1,0972 |
| 33 | 0,9688 | 0,9031 | | 0,0411 | 8,0000 | 1,0972 |
| 34 | 0,9912 | 0,8991 | | 0,0411 | 10,0000 | 1,0972 |
| 35 | 1,0137 | 0,8950 | | 0,0411 | 7,0000 | 1,0972 |
| 36 | 1,0361 | 0,8910 | | 0,0411 | 7,0000 | 1,0972 |
| 37 | 1,0586 | 0,8870 | | 0,0411 | 8,0000 | 1,0972 |
| 38 | 1,0811 | 0,8830 | | 0,0411 | 7,0000 | 1,0972 |
| 39 | 1,1035 | 0,8790 | | 0,0411 | 7,0000 | 1,0972 |
| 40 | 1,1260 | 0,8751 | | 0,0411 | 7,0000 | 1,0972 |
| 41 | 1,1484 | 0,8711 | | 0,0411 | 8,0000 | 1,0972 |
| 42 | 1,1709 | 0,8672 | | 0,0411 | 10,0000 | 1,0972 |
| 43 | 1,1934 | 0,8633 | | 0,0411 | 10,0000 | 1,0972 |
| 44 | 1,2158 | 0,8594 | | 0,0411 | 8,0000 | 1,0972 |
| 45 | 1,2383 | 0,8555 | | 0,0411 | 10,0000 | 1,0972 |
| 46 | 1,2607 | 0,8517 | | 0,0411 | 10,0000 | 1,0972 |
| 47 | 1,2832 | 0,8479 | | 0,0411 | 7,0000 | 1,0972 |
| 48 | 1,3057 | 0,8440 | | 0,0411 | 12,0000 | 1,0972 |
| 49 | 1,3281 | 0,8402 | | 0,0411 | 7,0000 | 1,0972 |

| | 13-b-Final | 14-x column 1 | 15-y column 1 | 16 | 17-SumSq 1 | 18-Iterations 1 |
|---|---|---|---|---|---|---|
| 1 | 0,2009 | 0,2500 | 1,0253 | | 0,0352 | 8,0000 |
| 2 | 0,2009 | 0,2725 | 1,0210 | | 0,0352 | 7,0000 |
| 3 | 0,2009 | 0,2949 | 1,0166 | | 0,0352 | 7,0000 |
| 4 | 0,2009 | 0,3174 | 1,0123 | | 0,0352 | 7,0000 |
| 5 | 0,2009 | 0,3398 | 1,0080 | | 0,0352 | 7,0000 |
| 6 | 0,2009 | 0,3623 | 1,0037 | | 0,0352 | 7,0000 |
| 7 | 0,2009 | 0,3848 | 0,9994 | | 0,0352 | 7,0000 |
| 8 | 0,2009 | 0,4072 | 0,9952 | | 0,0352 | 10,0000 |
| 9 | 0,2009 | 0,4297 | 0,9909 | | 0,0352 | 7,0000 |
| 10 | 0,2009 | 0,4521 | 0,9867 | | 0,0352 | 7,0000 |
| 11 | 0,2009 | 0,4746 | 0,9825 | | 0,0352 | 7,0000 |
| 12 | 0,2009 | 0,4971 | 0,9784 | | 0,0352 | 7,0000 |
| 13 | 0,2009 | 0,5195 | 0,9742 | | 0,0352 | 10,0000 |
| 14 | 0,2009 | 0,5420 | 0,9701 | | 0,0352 | 7,0000 |
| 15 | 0,2009 | 0,5645 | 0,9659 | | 0,0352 | 7,0000 |
| 16 | 0,2009 | 0,5869 | 0,9618 | | 0,0352 | 10,0000 |
| 17 | 0,2009 | 0,6094 | 0,9577 | | 0,0352 | 7,0000 |
| 18 | 0,2009 | 0,6318 | 0,9537 | | 0,0352 | 10,0000 |
| 19 | 0,2009 | 0,6543 | 0,9496 | | 0,0352 | 7,0000 |
| 20 | 0,2009 | 0,6768 | 0,9456 | | 0,0352 | 7,0000 |
| 21 | 0,2009 | 0,6992 | 0,9415 | | 0,0352 | 10,0000 |
| 22 | 0,2009 | 0,7217 | 0,9375 | | 0,0352 | 7,0000 |
| 23 | 0,2009 | 0,7441 | 0,9335 | | 0,0352 | 10,0000 |
| 24 | 0,2009 | 0,7666 | 0,9296 | | 0,0352 | 12,0000 |
| 25 | 0,2009 | 0,7891 | 0,9256 | | 0,0352 | 7,0000 |
| 26 | 0,2009 | 0,8115 | 0,9217 | | 0,0352 | 7,0000 |
| 27 | 0,2009 | 0,8340 | 0,9178 | | 0,0352 | 7,0000 |
| 28 | 0,2009 | 0,8564 | 0,9139 | | 0,0352 | 10,0000 |
| 29 | 0,2009 | 0,8789 | 0,9100 | | 0,0352 | 7,0000 |
| 30 | 0,2009 | 0,9014 | 0,9061 | | 0,0352 | 7,0000 |
| 31 | 0,2009 | 0,9238 | 0,9023 | | 0,0352 | 10,0000 |
| 32 | 0,2009 | 0,9463 | 0,8984 | | 0,0352 | 10,0000 |
| 33 | 0,2009 | 0,9688 | 0,8946 | | 0,0352 | 7,0000 |
| 34 | 0,2009 | 0,9912 | 0,8908 | | 0,0352 | 7,0000 |
| 35 | 0,2009 | 1,0137 | 0,8870 | | 0,0352 | 7,0000 |
| 36 | 0,2009 | 1,0361 | 0,8832 | | 0,0352 | 10,0000 |
| 37 | 0,2009 | 1,0586 | 0,8795 | | 0,0352 | 7,0000 |
| 38 | 0,2009 | 1,0811 | 0,8757 | | 0,0352 | 7,0000 |
| 39 | 0,2009 | 1,1035 | 0,8720 | | 0,0352 | 8,0000 |
| 40 | 0,2009 | 1,1260 | 0,8683 | | 0,0352 | 7,0000 |
| 41 | 0,2009 | 1,1484 | 0,8646 | | 0,0352 | 7,0000 |
| 42 | 0,2009 | 1,1709 | 0,8609 | | 0,0352 | 8,0000 |
| 43 | 0,2009 | 1,1934 | 0,8573 | | 0,0352 | 7,0000 |
| 44 | 0,2009 | 1,2158 | 0,8536 | | 0,0352 | 7,0000 |
| 45 | 0,2009 | 1,2383 | 0,8500 | | 0,0352 | 7,0000 |
| 46 | 0,2009 | 1,2607 | 0,8464 | | 0,0352 | 7,0000 |
| 47 | 0,2009 | 1,2832 | 0,8428 | | 0,0352 | 7,0000 |
| 48 | 0,2009 | 1,3057 | 0,8392 | | 0,0352 | 7,0000 |
| 49 | 0,2009 | 1,3281 | 0,8356 | | 0,0352 | 7,0000 |

| | 19-a-Final 1 | 20-b-Final 1 | 21-x column 2 | 22-y column 2 | 23 | 24-SumSq 2 |
|---|---|---|---|---|---|---|
| 1 | 1,0751 | 0,1897 | 0,2500 | 0,9899 | | 0,0664 |
| 2 | 1,0751 | 0,1897 | 0,2725 | 0,9831 | | 0,0664 |
| 3 | 1,0751 | 0,1897 | 0,2949 | 0,9764 | | 0,0664 |
| 4 | 1,0751 | 0,1897 | 0,3174 | 0,9697 | | 0,0664 |
| 5 | 1,0751 | 0,1897 | 0,3398 | 0,9631 | | 0,0664 |
| 6 | 1,0751 | 0,1897 | 0,3623 | 0,9565 | | 0,0664 |
| 7 | 1,0751 | 0,1897 | 0,3848 | 0,9500 | | 0,0664 |
| 8 | 1,0751 | 0,1897 | 0,4072 | 0,9435 | | 0,0664 |
| 9 | 1,0751 | 0,1897 | 0,4297 | 0,9371 | | 0,0664 |
| 10 | 1,0751 | 0,1897 | 0,4521 | 0,9307 | | 0,0664 |
| 11 | 1,0751 | 0,1897 | 0,4746 | 0,9243 | | 0,0664 |
| 12 | 1,0751 | 0,1897 | 0,4971 | 0,9180 | | 0,0664 |
| 13 | 1,0751 | 0,1897 | 0,5195 | 0,9117 | | 0,0664 |
| 14 | 1,0751 | 0,1897 | 0,5420 | 0,9055 | | 0,0664 |
| 15 | 1,0751 | 0,1897 | 0,5645 | 0,8993 | | 0,0664 |
| 16 | 1,0751 | 0,1897 | 0,5869 | 0,8931 | | 0,0664 |
| 17 | 1,0751 | 0,1897 | 0,6094 | 0,8870 | | 0,0664 |
| 18 | 1,0751 | 0,1897 | 0,6318 | 0,8810 | | 0,0664 |
| 19 | 1,0751 | 0,1897 | 0,6543 | 0,8750 | | 0,0664 |
| 20 | 1,0751 | 0,1897 | 0,6768 | 0,8690 | | 0,0664 |
| 21 | 1,0751 | 0,1897 | 0,6992 | 0,8630 | | 0,0664 |
| 22 | 1,0751 | 0,1897 | 0,7217 | 0,8572 | | 0,0664 |
| 23 | 1,0751 | 0,1897 | 0,7441 | 0,8513 | | 0,0664 |
| 24 | 1,0751 | 0,1897 | 0,7666 | 0,8455 | | 0,0664 |
| 25 | 1,0751 | 0,1897 | 0,7891 | 0,8397 | | 0,0664 |
| 26 | 1,0751 | 0,1897 | 0,8115 | 0,8340 | | 0,0664 |
| 27 | 1,0751 | 0,1897 | 0,8340 | 0,8283 | | 0,0664 |
| 28 | 1,0751 | 0,1897 | 0,8564 | 0,8226 | | 0,0664 |
| 29 | 1,0751 | 0,1897 | 0,8789 | 0,8170 | | 0,0664 |
| 30 | 1,0751 | 0,1897 | 0,9014 | 0,8114 | | 0,0664 |
| 31 | 1,0751 | 0,1897 | 0,9238 | 0,8059 | | 0,0664 |
| 32 | 1,0751 | 0,1897 | 0,9463 | 0,8004 | | 0,0664 |
| 33 | 1,0751 | 0,1897 | 0,9688 | 0,7949 | | 0,0664 |
| 34 | 1,0751 | 0,1897 | 0,9912 | 0,7895 | | 0,0664 |
| 35 | 1,0751 | 0,1897 | 1,0137 | 0,7841 | | 0,0664 |
| 36 | 1,0751 | 0,1897 | 1,0361 | 0,7787 | | 0,0664 |
| 37 | 1,0751 | 0,1897 | 1,0586 | 0,7734 | | 0,0664 |
| 38 | 1,0751 | 0,1897 | 1,0811 | 0,7681 | | 0,0664 |
| 39 | 1,0751 | 0,1897 | 1,1035 | 0,7628 | | 0,0664 |
| 40 | 1,0751 | 0,1897 | 1,1260 | 0,7576 | | 0,0664 |
| 41 | 1,0751 | 0,1897 | 1,1484 | 0,7525 | | 0,0664 |
| 42 | 1,0751 | 0,1897 | 1,1709 | 0,7473 | | 0,0664 |
| 43 | 1,0751 | 0,1897 | 1,1934 | 0,7422 | | 0,0664 |
| 44 | 1,0751 | 0,1897 | 1,2158 | 0,7371 | | 0,0664 |
| 45 | 1,0751 | 0,1897 | 1,2383 | 0,7321 | | 0,0664 |
| 46 | 1,0751 | 0,1897 | 1,2607 | 0,7271 | | 0,0664 |
| 47 | 1,0751 | 0,1897 | 1,2832 | 0,7221 | | 0,0664 |
| 48 | 1,0751 | 0,1897 | 1,3057 | 0,7172 | | 0,0664 |
| 49 | 1,0751 | 0,1897 | 1,3281 | 0,7123 | | 0,0664 |

| | 25-Iterations 2 | 26-a-Final 2 | 27-b-Final 2 | 28-x column 3 | 29-y column 3 | 30 |
|---|---|---|---|---|---|---|
| 1 | 11,0000 | 1,0684 | 0,3052 | 0,2500 | 0,9145 | |
| 2 | 10,0000 | 1,0684 | 0,3052 | 0,2725 | 0,9067 | |
| 3 | 14,0000 | 1,0684 | 0,3052 | 0,2949 | 0,8988 | |
| 4 | 10,0000 | 1,0684 | 0,3052 | 0,3174 | 0,8911 | |
| 5 | 13,0000 | 1,0684 | 0,3052 | 0,3398 | 0,8834 | |
| 6 | 11,0000 | 1,0684 | 0,3052 | 0,3623 | 0,8758 | |
| 7 | 11,0000 | 1,0684 | 0,3052 | 0,3848 | 0,8683 | |
| 8 | 10,0000 | 1,0684 | 0,3052 | 0,4072 | 0,8608 | |
| 9 | 15,0000 | 1,0684 | 0,3052 | 0,4297 | 0,8534 | |
| 10 | 10,0000 | 1,0684 | 0,3052 | 0,4521 | 0,8460 | |
| 11 | 10,0000 | 1,0684 | 0,3052 | 0,4746 | 0,8387 | |
| 12 | 10,0000 | 1,0684 | 0,3052 | 0,4971 | 0,8315 | |
| 13 | 11,0000 | 1,0684 | 0,3052 | 0,5195 | 0,8243 | |
| 14 | 10,0000 | 1,0684 | 0,3052 | 0,5420 | 0,8172 | |
| 15 | 15,0000 | 1,0684 | 0,3052 | 0,5645 | 0,8102 | |
| 16 | 10,0000 | 1,0684 | 0,3052 | 0,5869 | 0,8032 | |
| 17 | 15,0000 | 1,0684 | 0,3052 | 0,6094 | 0,7963 | |
| 18 | 25,0000 | 1,0684 | 0,3052 | 0,6318 | 0,7894 | |
| 19 | 10,0000 | 1,0684 | 0,3052 | 0,6543 | 0,7826 | |
| 20 | 15,0000 | 1,0684 | 0,3052 | 0,6768 | 0,7758 | |
| 21 | 10,0000 | 1,0684 | 0,3052 | 0,6992 | 0,7692 | |
| 22 | 11,0000 | 1,0684 | 0,3052 | 0,7217 | 0,7625 | |
| 23 | 15,0000 | 1,0684 | 0,3052 | 0,7441 | 0,7560 | |
| 24 | 13,0000 | 1,0684 | 0,3052 | 0,7666 | 0,7494 | |
| 25 | 10,0000 | 1,0684 | 0,3052 | 0,7891 | 0,7430 | |
| 26 | 11,0000 | 1,0684 | 0,3052 | 0,8115 | 0,7366 | |
| 27 | 10,0000 | 1,0684 | 0,3052 | 0,8340 | 0,7302 | |
| 28 | 10,0000 | 1,0684 | 0,3052 | 0,8564 | 0,7239 | |
| 29 | 10,0000 | 1,0684 | 0,3052 | 0,8789 | 0,7177 | |
| 30 | 12,0000 | 1,0684 | 0,3052 | 0,9014 | 0,7115 | |
| 31 | 11,0000 | 1,0684 | 0,3052 | 0,9238 | 0,7054 | |
| 32 | 11,0000 | 1,0684 | 0,3052 | 0,9463 | 0,6993 | |
| 33 | 11,0000 | 1,0684 | 0,3052 | 0,9688 | 0,6933 | |
| 34 | 15,0000 | 1,0684 | 0,3052 | 0,9912 | 0,6873 | |
| 35 | 11,0000 | 1,0684 | 0,3052 | 1,0137 | 0,6814 | |
| 36 | 10,0000 | 1,0684 | 0,3052 | 1,0361 | 0,6755 | |
| 37 | 10,0000 | 1,0684 | 0,3052 | 1,0586 | 0,6697 | |
| 38 | 15,0000 | 1,0684 | 0,3052 | 1,0811 | 0,6639 | |
| 39 | 9,0000 | 1,0684 | 0,3052 | 1,1035 | 0,6582 | |
| 40 | 10,0000 | 1,0684 | 0,3052 | 1,1260 | 0,6525 | |
| 41 | 10,0000 | 1,0684 | 0,3052 | 1,1484 | 0,6469 | |
| 42 | 10,0000 | 1,0684 | 0,3052 | 1,1709 | 0,6413 | |
| 43 | 15,0000 | 1,0684 | 0,3052 | 1,1934 | 0,6358 | |
| 44 | 10,0000 | 1,0684 | 0,3052 | 1,2158 | 0,6303 | |
| 45 | 14,0000 | 1,0684 | 0,3052 | 1,2383 | 0,6249 | |
| 46 | 10,0000 | 1,0684 | 0,3052 | 1,2607 | 0,6195 | |
| 47 | 15,0000 | 1,0684 | 0,3052 | 1,2832 | 0,6142 | |
| 48 | 10,0000 | 1,0684 | 0,3052 | 1,3057 | 0,6089 | |
| 49 | 10,0000 | 1,0684 | 0,3052 | 1,3281 | 0,6036 | |

| | 31-SumSq 3 | 32-Iterations 3 | 33-a-Final 3 | 34-b-Final 3 | 35-x column 4 | 36-y column 4 |
|---|---|---|---|---|---|---|
| 1 | 0,0429 | 11,0000 | 1,0070 | 0,3854 | 0,2500 | 0,9145 |
| 2 | 0,0429 | 12,0000 | 1,0070 | 0,3854 | 0,2725 | 0,9067 |
| 3 | 0,0429 | 15,0000 | 1,0070 | 0,3854 | 0,2949 | 0,8988 |
| 4 | 0,0429 | 16,0000 | 1,0070 | 0,3854 | 0,3174 | 0,8911 |
| 5 | 0,0429 | 14,0000 | 1,0070 | 0,3854 | 0,3398 | 0,8834 |
| 6 | 0,0429 | 12,0000 | 1,0070 | 0,3854 | 0,3623 | 0,8758 |
| 7 | 0,0429 | 11,0000 | 1,0070 | 0,3854 | 0,3848 | 0,8683 |
| 8 | 0,0429 | 12,0000 | 1,0070 | 0,3854 | 0,4072 | 0,8608 |
| 9 | 0,0429 | 11,0000 | 1,0070 | 0,3854 | 0,4297 | 0,8534 |
| 10 | 0,0429 | 11,0000 | 1,0070 | 0,3854 | 0,4521 | 0,8460 |
| 11 | 0,0429 | 16,0000 | 1,0070 | 0,3854 | 0,4746 | 0,8387 |
| 12 | 0,0429 | 12,0000 | 1,0070 | 0,3854 | 0,4971 | 0,8315 |
| 13 | 0,0429 | 16,0000 | 1,0070 | 0,3854 | 0,5195 | 0,8243 |
| 14 | 0,0429 | 15,0000 | 1,0070 | 0,3854 | 0,5420 | 0,8172 |
| 15 | 0,0429 | 12,0000 | 1,0070 | 0,3854 | 0,5645 | 0,8102 |
| 16 | 0,0429 | 12,0000 | 1,0070 | 0,3854 | 0,5869 | 0,8032 |
| 17 | 0,0429 | 11,0000 | 1,0070 | 0,3854 | 0,6094 | 0,7963 |
| 18 | 0,0429 | 16,0000 | 1,0070 | 0,3854 | 0,6318 | 0,7894 |
| 19 | 0,0429 | 16,0000 | 1,0070 | 0,3854 | 0,6543 | 0,7826 |
| 20 | 0,0429 | 11,0000 | 1,0070 | 0,3854 | 0,6768 | 0,7758 |
| 21 | 0,0429 | 16,0000 | 1,0070 | 0,3854 | 0,6992 | 0,7692 |
| 22 | 0,0429 | 8,0000 | 1,0070 | 0,3854 | 0,7217 | 0,7625 |
| 23 | 0,0429 | 24,0000 | 1,0070 | 0,3854 | 0,7441 | 0,7560 |
| 24 | 0,0429 | 14,0000 | 1,0070 | 0,3854 | 0,7666 | 0,7494 |
| 25 | 0,0429 | 16,0000 | 1,0070 | 0,3854 | 0,7891 | 0,7430 |
| 26 | 0,0429 | 16,0000 | 1,0070 | 0,3854 | 0,8115 | 0,7366 |
| 27 | 0,0429 | 11,0000 | 1,0070 | 0,3854 | 0,8340 | 0,7302 |
| 28 | 0,0429 | 10,0000 | 1,0070 | 0,3854 | 0,8564 | 0,7239 |
| 29 | 0,0429 | 11,0000 | 1,0070 | 0,3854 | 0,8789 | 0,7177 |
| 30 | 0,0429 | 11,0000 | 1,0070 | 0,3854 | 0,9014 | 0,7115 |
| 31 | 0,0429 | 11,0000 | 1,0070 | 0,3854 | 0,9238 | 0,7054 |
| 32 | 0,0429 | 11,0000 | 1,0070 | 0,3854 | 0,9463 | 0,6993 |
| 33 | 0,0429 | 15,0000 | 1,0070 | 0,3854 | 0,9688 | 0,6933 |
| 34 | 0,0429 | 11,0000 | 1,0070 | 0,3854 | 0,9912 | 0,6873 |
| 35 | 0,0429 | 11,0000 | 1,0070 | 0,3854 | 1,0137 | 0,6814 |
| 36 | 0,0429 | 33,0000 | 1,0070 | 0,3854 | 1,0361 | 0,6755 |
| 37 | 0,0429 | 11,0000 | 1,0070 | 0,3854 | 1,0586 | 0,6697 |
| 38 | 0,0429 | 14,0000 | 1,0070 | 0,3854 | 1,0811 | 0,6639 |
| 39 | 0,0429 | 6,0000 | 1,0070 | 0,3854 | 1,1035 | 0,6582 |
| 40 | 0,0429 | 11,0000 | 1,0070 | 0,3854 | 1,1260 | 0,6525 |
| 41 | 0,0429 | 11,0000 | 1,0070 | 0,3854 | 1,1484 | 0,6469 |
| 42 | 0,0429 | 16,0000 | 1,0070 | 0,3854 | 1,1709 | 0,6413 |
| 43 | 0,0429 | 14,0000 | 1,0070 | 0,3854 | 1,1934 | 0,6358 |
| 44 | 0,0429 | 11,0000 | 1,0070 | 0,3854 | 1,2158 | 0,6303 |
| 45 | 0,0429 | 14,0000 | 1,0070 | 0,3854 | 1,2383 | 0,6249 |
| 46 | 0,0429 | 14,0000 | 1,0070 | 0,3854 | 1,2607 | 0,6195 |
| 47 | 0,0429 | 11,0000 | 1,0070 | 0,3854 | 1,2832 | 0,6142 |
| 48 | 0,0429 | 11,0000 | 1,0070 | 0,3854 | 1,3057 | 0,6089 |
| 49 | 0,0429 | 11,0000 | 1,0070 | 0,3854 | 1,3281 | 0,6036 |

| | 37 | 38-SumSq 4 | 39-Iterations 4 | 40-a-Final 4 | 41-b-Final 4 | 42-x column 5 |
|---|---|---|---|---|---|---|
| 1 | | 0,0429 | 12,0000 | 1,0070 | 0,3854 | 0,2500 |
| 2 | | 0,0429 | 11,0000 | 1,0070 | 0,3854 | 0,2725 |
| 3 | | 0,0429 | 12,0000 | 1,0070 | 0,3854 | 0,2949 |
| 4 | | 0,0429 | 10,0000 | 1,0070 | 0,3854 | 0,3174 |
| 5 | | 0,0429 | 10,0000 | 1,0070 | 0,3854 | 0,3398 |
| 6 | | 0,0429 | 11,0000 | 1,0070 | 0,3854 | 0,3623 |
| 7 | | 0,0429 | 12,0000 | 1,0070 | 0,3854 | 0,3848 |
| 8 | | 0,0429 | 12,0000 | 1,0070 | 0,3854 | 0,4072 |
| 9 | | 0,0429 | 10,0000 | 1,0070 | 0,3854 | 0,4297 |
| 10 | | 0,0429 | 12,0000 | 1,0070 | 0,3854 | 0,4521 |
| 11 | | 0,0429 | 10,0000 | 1,0070 | 0,3854 | 0,4746 |
| 12 | | 0,0429 | 9,0000 | 1,0070 | 0,3854 | 0,4971 |
| 13 | | 0,0429 | 11,0000 | 1,0070 | 0,3854 | 0,5195 |
| 14 | | 0,0429 | 12,0000 | 1,0070 | 0,3854 | 0,5420 |
| 15 | | 0,0429 | 10,0000 | 1,0070 | 0,3854 | 0,5645 |
| 16 | | 0,0429 | 11,0000 | 1,0070 | 0,3854 | 0,5869 |
| 17 | | 0,0429 | 10,0000 | 1,0070 | 0,3854 | 0,6094 |
| 18 | | 0,0429 | 10,0000 | 1,0070 | 0,3854 | 0,6318 |
| 19 | | 0,0429 | 11,0000 | 1,0070 | 0,3854 | 0,6543 |
| 20 | | 0,0429 | 9,0000 | 1,0070 | 0,3854 | 0,6768 |
| 21 | | 0,0429 | 10,0000 | 1,0070 | 0,3854 | 0,6992 |
| 22 | | 0,0429 | 11,0000 | 1,0070 | 0,3854 | 0,7217 |
| 23 | | 0,0429 | 11,0000 | 1,0070 | 0,3854 | 0,7441 |
| 24 | | 0,0429 | 10,0000 | 1,0070 | 0,3854 | 0,7666 |
| 25 | | 0,0429 | 10,0000 | 1,0070 | 0,3854 | 0,7891 |
| 26 | | 0,0429 | 11,0000 | 1,0070 | 0,3854 | 0,8115 |
| 27 | | 0,0429 | 10,0000 | 1,0070 | 0,3854 | 0,8340 |
| 28 | | 0,0429 | 11,0000 | 1,0070 | 0,3854 | 0,8564 |
| 29 | | 0,0429 | 10,0000 | 1,0070 | 0,3854 | 0,8789 |
| 30 | | 0,0429 | 9,0000 | 1,0070 | 0,3854 | 0,9014 |
| 31 | | 0,0429 | 10,0000 | 1,0070 | 0,3854 | 0,9238 |
| 32 | | 0,0429 | 10,0000 | 1,0070 | 0,3854 | 0,9463 |
| 33 | | 0,0429 | 9,0000 | 1,0070 | 0,3854 | 0,9688 |
| 34 | | 0,0429 | 11,0000 | 1,0070 | 0,3854 | 0,9912 |
| 35 | | 0,0429 | 10,0000 | 1,0070 | 0,3854 | 1,0137 |
| 36 | | 0,0429 | 10,0000 | 1,0070 | 0,3854 | 1,0361 |
| 37 | | 0,0429 | 8,0000 | 1,0070 | 0,3854 | 1,0586 |
| 38 | | 0,0429 | 11,0000 | 1,0070 | 0,3854 | 1,0811 |
| 39 | | 0,0429 | 10,0000 | 1,0070 | 0,3854 | 1,1035 |
| 40 | | 0,0429 | 11,0000 | 1,0070 | 0,3854 | 1,1260 |
| 41 | | 0,0429 | 10,0000 | 1,0070 | 0,3854 | 1,1484 |
| 42 | | 0,0429 | 10,0000 | 1,0070 | 0,3854 | 1,1709 |
| 43 | | 0,0429 | 10,0000 | 1,0070 | 0,3854 | 1,1934 |
| 44 | | 0,0429 | 11,0000 | 1,0070 | 0,3854 | 1,2158 |
| 45 | | 0,0429 | 10,0000 | 1,0070 | 0,3854 | 1,2383 |
| 46 | | 0,0429 | 10,0000 | 1,0070 | 0,3854 | 1,2607 |
| 47 | | 0,0429 | 10,0000 | 1,0070 | 0,3854 | 1,2832 |
| 48 | | 0,0429 | 11,0000 | 1,0070 | 0,3854 | 1,3057 |
| 49 | | 0,0429 | 9,0000 | 1,0070 | 0,3854 | 1,3281 |

| | 43-y column 5 | 44 | 45-SumSq 5 | 46-Iterations 5 | 47-a-Final 5 | 48-b-Final 5 |
|---|---|---|---|---|---|---|
| 1 | 0,9454 | | 0,1194 | 11,0000 | 1,0036 | 0,2389 |
| 2 | 0,9404 | | 0,1194 | 9,0000 | 1,0036 | 0,2389 |
| 3 | 0,9353 | | 0,1194 | 10,0000 | 1,0036 | 0,2389 |
| 4 | 0,9303 | | 0,1194 | 10,0000 | 1,0036 | 0,2389 |
| 5 | 0,9254 | | 0,1194 | 11,0000 | 1,0036 | 0,2389 |
| 6 | 0,9204 | | 0,1194 | 11,0000 | 1,0036 | 0,2389 |
| 7 | 0,9155 | | 0,1194 | 9,0000 | 1,0036 | 0,2389 |
| 8 | 0,9106 | | 0,1194 | 10,0000 | 1,0036 | 0,2389 |
| 9 | 0,9057 | | 0,1194 | 8,0000 | 1,0036 | 0,2389 |
| 10 | 0,9009 | | 0,1194 | 9,0000 | 1,0036 | 0,2389 |
| 11 | 0,8960 | | 0,1194 | 9,0000 | 1,0036 | 0,2389 |
| 12 | 0,8912 | | 0,1194 | 10,0000 | 1,0036 | 0,2389 |
| 13 | 0,8865 | | 0,1194 | 10,0000 | 1,0036 | 0,2389 |
| 14 | 0,8817 | | 0,1194 | 9,0000 | 1,0036 | 0,2389 |
| 15 | 0,8770 | | 0,1194 | 11,0000 | 1,0036 | 0,2389 |
| 16 | 0,8723 | | 0,1194 | 11,0000 | 1,0036 | 0,2389 |
| 17 | 0,8677 | | 0,1194 | 9,0000 | 1,0036 | 0,2389 |
| 18 | 0,8630 | | 0,1194 | 11,0000 | 1,0036 | 0,2389 |
| 19 | 0,8584 | | 0,1194 | 10,0000 | 1,0036 | 0,2389 |
| 20 | 0,8538 | | 0,1194 | 10,0000 | 1,0036 | 0,2389 |
| 21 | 0,8492 | | 0,1194 | 9,0000 | 1,0036 | 0,2389 |
| 22 | 0,8447 | | 0,1194 | 10,0000 | 1,0036 | 0,2389 |
| 23 | 0,8402 | | 0,1194 | 11,0000 | 1,0036 | 0,2389 |
| 24 | 0,8357 | | 0,1194 | 10,0000 | 1,0036 | 0,2389 |
| 25 | 0,8312 | | 0,1194 | 11,0000 | 1,0036 | 0,2389 |
| 26 | 0,8268 | | 0,1194 | 10,0000 | 1,0036 | 0,2389 |
| 27 | 0,8223 | | 0,1194 | 11,0000 | 1,0036 | 0,2389 |
| 28 | 0,8179 | | 0,1194 | 10,0000 | 1,0036 | 0,2389 |
| 29 | 0,8135 | | 0,1194 | 9,0000 | 1,0036 | 0,2389 |
| 30 | 0,8092 | | 0,1194 | 9,0000 | 1,0036 | 0,2389 |
| 31 | 0,8049 | | 0,1194 | 10,0000 | 1,0036 | 0,2389 |
| 32 | 0,8006 | | 0,1194 | 10,0000 | 1,0036 | 0,2389 |
| 33 | 0,7963 | | 0,1194 | 11,0000 | 1,0036 | 0,2389 |
| 34 | 0,7920 | | 0,1194 | 10,0000 | 1,0036 | 0,2389 |
| 35 | 0,7878 | | 0,1194 | 7,0000 | 1,0036 | 0,2389 |
| 36 | 0,7836 | | 0,1194 | 11,0000 | 1,0036 | 0,2389 |
| 37 | 0,7794 | | 0,1194 | 10,0000 | 1,0036 | 0,2389 |
| 38 | 0,7752 | | 0,1194 | 10,0000 | 1,0036 | 0,2389 |
| 39 | 0,7710 | | 0,1194 | 9,0000 | 1,0036 | 0,2389 |
| 40 | 0,7669 | | 0,1194 | 10,0000 | 1,0036 | 0,2389 |
| 41 | 0,7628 | | 0,1194 | 9,0000 | 1,0036 | 0,2389 |
| 42 | 0,7587 | | 0,1194 | 9,0000 | 1,0036 | 0,2389 |
| 43 | 0,7547 | | 0,1194 | 10,0000 | 1,0036 | 0,2389 |
| 44 | 0,7506 | | 0,1194 | 10,0000 | 1,0036 | 0,2389 |
| 45 | 0,7466 | | 0,1194 | 9,0000 | 1,0036 | 0,2389 |
| 46 | 0,7426 | | 0,1194 | 8,0000 | 1,0036 | 0,2389 |
| 47 | 0,7387 | | 0,1194 | 10,0000 | 1,0036 | 0,2389 |
| 48 | 0,7347 | | 0,1194 | 9,0000 | 1,0036 | 0,2389 |
| 49 | 0,7308 | | 0,1194 | 10,0000 | 1,0036 | 0,2389 |

| | 49-x column 6 | 50-y column 6 | 51 | 52-SumSq 6 | 53-Iterations 6 | 54-a-Final 6 |
|---|---|---|---|---|---|---|
| 1 | 0,2500 | 0,8725 | | 0,0865 | 11,0000 | 0,9419 |
| 2 | 0,2725 | 0,8665 | | 0,0865 | 10,0000 | 0,9419 |
| 3 | 0,2949 | 0,8605 | | 0,0865 | 11,0000 | 0,9419 |
| 4 | 0,3174 | 0,8546 | | 0,0865 | 9,0000 | 0,9419 |
| 5 | 0,3398 | 0,8488 | | 0,0865 | 10,0000 | 0,9419 |
| 6 | 0,3623 | 0,8429 | | 0,0865 | 11,0000 | 0,9419 |
| 7 | 0,3848 | 0,8372 | | 0,0865 | 10,0000 | 0,9419 |
| 8 | 0,4072 | 0,8314 | | 0,0865 | 8,0000 | 0,9419 |
| 9 | 0,4297 | 0,8257 | | 0,0865 | 11,0000 | 0,9419 |
| 10 | 0,4521 | 0,8201 | | 0,0865 | 10,0000 | 0,9419 |
| 11 | 0,4746 | 0,8144 | | 0,0865 | 10,0000 | 0,9419 |
| 12 | 0,4971 | 0,8088 | | 0,0865 | 11,0000 | 0,9419 |
| 13 | 0,5195 | 0,8033 | | 0,0865 | 10,0000 | 0,9419 |
| 14 | 0,5420 | 0,7978 | | 0,0865 | 10,0000 | 0,9419 |
| 15 | 0,5645 | 0,7923 | | 0,0865 | 11,0000 | 0,9419 |
| 16 | 0,5869 | 0,7869 | | 0,0865 | 11,0000 | 0,9419 |
| 17 | 0,6094 | 0,7815 | | 0,0865 | 10,0000 | 0,9419 |
| 18 | 0,6318 | 0,7761 | | 0,0865 | 10,0000 | 0,9419 |
| 19 | 0,6543 | 0,7708 | | 0,0865 | 11,0000 | 0,9419 |
| 20 | 0,6768 | 0,7655 | | 0,0865 | 10,0000 | 0,9419 |
| 21 | 0,6992 | 0,7603 | | 0,0865 | 9,0000 | 0,9419 |
| 22 | 0,7217 | 0,7550 | | 0,0865 | 11,0000 | 0,9419 |
| 23 | 0,7441 | 0,7499 | | 0,0865 | 9,0000 | 0,9419 |
| 24 | 0,7666 | 0,7447 | | 0,0865 | 11,0000 | 0,9419 |
| 25 | 0,7891 | 0,7396 | | 0,0865 | 10,0000 | 0,9419 |
| 26 | 0,8115 | 0,7345 | | 0,0865 | 10,0000 | 0,9419 |
| 27 | 0,8340 | 0,7295 | | 0,0865 | 11,0000 | 0,9419 |
| 28 | 0,8564 | 0,7245 | | 0,0865 | 11,0000 | 0,9419 |
| 29 | 0,8789 | 0,7195 | | 0,0865 | 10,0000 | 0,9419 |
| 30 | 0,9014 | 0,7146 | | 0,0865 | 10,0000 | 0,9419 |
| 31 | 0,9238 | 0,7097 | | 0,0865 | 10,0000 | 0,9419 |
| 32 | 0,9463 | 0,7048 | | 0,0865 | 10,0000 | 0,9419 |
| 33 | 0,9688 | 0,7000 | | 0,0865 | 11,0000 | 0,9419 |
| 34 | 0,9912 | 0,6952 | | 0,0865 | 10,0000 | 0,9419 |
| 35 | 1,0137 | 0,6904 | | 0,0865 | 10,0000 | 0,9419 |
| 36 | 1,0361 | 0,6857 | | 0,0865 | 11,0000 | 0,9419 |
| 37 | 1,0586 | 0,6810 | | 0,0865 | 10,0000 | 0,9419 |
| 38 | 1,0811 | 0,6763 | | 0,0865 | 10,0000 | 0,9419 |
| 39 | 1,1035 | 0,6717 | | 0,0865 | 10,0000 | 0,9419 |
| 40 | 1,1260 | 0,6671 | | 0,0865 | 10,0000 | 0,9419 |
| 41 | 1,1484 | 0,6625 | | 0,0865 | 11,0000 | 0,9419 |
| 42 | 1,1709 | 0,6579 | | 0,0865 | 9,0000 | 0,9419 |
| 43 | 1,1934 | 0,6534 | | 0,0865 | 11,0000 | 0,9419 |
| 44 | 1,2158 | 0,6490 | | 0,0865 | 10,0000 | 0,9419 |
| 45 | 1,2383 | 0,6445 | | 0,0865 | 10,0000 | 0,9419 |
| 46 | 1,2607 | 0,6401 | | 0,0865 | 11,0000 | 0,9419 |
| 47 | 1,2832 | 0,6357 | | 0,0865 | 9,0000 | 0,9419 |
| 48 | 1,3057 | 0,6313 | | 0,0865 | 10,0000 | 0,9419 |
| 49 | 1,3281 | 0,6270 | | 0,0865 | 10,0000 | 0,9419 |

| | 55-b-Final 6 |
|---|---|
| 1 | 0,3064 |
| 2 | 0,3064 |
| 3 | 0,3064 |
| 4 | 0,3064 |
| 5 | 0,3064 |
| 6 | 0,3064 |
| 7 | 0,3064 |
| 8 | 0,3064 |
| 9 | 0,3064 |
| 10 | 0,3064 |
| 11 | 0,3064 |
| 12 | 0,3064 |
| 13 | 0,3064 |
| 14 | 0,3064 |
| 15 | 0,3064 |
| 16 | 0,3064 |
| 17 | 0,3064 |
| 18 | 0,3064 |
| 19 | 0,3064 |
| 20 | 0,3064 |
| 21 | 0,3064 |
| 22 | 0,3064 |
| 23 | 0,3064 |
| 24 | 0,3064 |
| 25 | 0,3064 |
| 26 | 0,3064 |
| 27 | 0,3064 |
| 28 | 0,3064 |
| 29 | 0,3064 |
| 30 | 0,3064 |
| 31 | 0,3064 |
| 32 | 0,3064 |
| 33 | 0,3064 |
| 34 | 0,3064 |
| 35 | 0,3064 |
| 36 | 0,3064 |
| 37 | 0,3064 |
| 38 | 0,3064 |
| 39 | 0,3064 |
| 40 | 0,3064 |
| 41 | 0,3064 |
| 42 | 0,3064 |
| 43 | 0,3064 |
| 44 | 0,3064 |
| 45 | 0,3064 |
| 46 | 0,3064 |
| 47 | 0,3064 |
| 48 | 0,3064 |
| 49 | 0,3064 |

| | 1 | 2 | 3 | 4 | 5 | 6 |
|---|---|---|---|---|---|---|
| 50 | | | | | | |
| 51 | | | | | | |
| 52 | | | | | | |
| 53 | | | | | | |
| 54 | | | | | | |
| 55 | | | | | | |
| 56 | | | | | | |
| 57 | | | | | | |
| 58 | | | | | | |
| 59 | | | | | | |
| 60 | | | | | | |
| 61 | | | | | | |
| 62 | | | | | | |
| 63 | | | | | | |
| 64 | | | | | | |
| 65 | | | | | | |
| 66 | | | | | | |
| 67 | | | | | | |
| 68 | | | | | | |
| 69 | | | | | | |
| 70 | | | | | | |
| 71 | | | | | | |
| 72 | | | | | | |
| 73 | | | | | | |
| 74 | | | | | | |
| 75 | | | | | | |
| 76 | | | | | | |
| 77 | | | | | | |
| 78 | | | | | | |
| 79 | | | | | | |
| 80 | | | | | | |
| 81 | | | | | | |
| 82 | | | | | | |
| 83 | | | | | | |
| 84 | | | | | | |
| 85 | | | | | | |
| 86 | | | | | | |
| 87 | | | | | | |
| 88 | | | | | | |
| 89 | | | | | | |
| 90 | | | | | | |
| 91 | | | | | | |
| 92 | | | | | | |
| 93 | | | | | | |
| 94 | | | | | | |
| 95 | | | | | | |
| 96 | | | | | | |
| 97 | | | | | | |
| 98 | | | | | | |

| | 7-x column | 8-y column | 9 | 10-SumSq | 11-Iterations | 12-a-Final |
|---|---|---|---|---|---|---|
| 50 | 1,3506 | 0,8365 | | 0,0411 | 7,0000 | 1,0972 |
| 51 | 1,3730 | 0,8327 | | 0,0411 | 7,0000 | 1,0972 |
| 52 | 1,3955 | 0,8289 | | 0,0411 | 10,0000 | 1,0972 |
| 53 | 1,4180 | 0,8252 | | 0,0411 | 7,0000 | 1,0972 |
| 54 | 1,4404 | 0,8215 | | 0,0411 | 7,0000 | 1,0972 |
| 55 | 1,4629 | 0,8178 | | 0,0411 | 8,0000 | 1,0972 |
| 56 | 1,4854 | 0,8141 | | 0,0411 | 10,0000 | 1,0972 |
| 57 | 1,5078 | 0,8105 | | 0,0411 | 8,0000 | 1,0972 |
| 58 | 1,5303 | 0,8068 | | 0,0411 | 12,0000 | 1,0972 |
| 59 | 1,5527 | 0,8032 | | 0,0411 | 8,0000 | 1,0972 |
| 60 | 1,5752 | 0,7996 | | 0,0411 | 10,0000 | 1,0972 |
| 61 | 1,5977 | 0,7960 | | 0,0411 | 10,0000 | 1,0972 |
| 62 | 1,6201 | 0,7924 | | 0,0411 | 7,0000 | 1,0972 |
| 63 | 1,6426 | 0,7888 | | 0,0411 | 10,0000 | 1,0972 |
| 64 | 1,6650 | 0,7853 | | 0,0411 | 8,0000 | 1,0972 |
| 65 | 1,6875 | 0,7817 | | 0,0411 | 10,0000 | 1,0972 |
| 66 | 1,7100 | 0,7782 | | 0,0411 | 10,0000 | 1,0972 |
| 67 | 1,7324 | 0,7747 | | 0,0411 | 10,0000 | 1,0972 |
| 68 | 1,7549 | 0,7712 | | 0,0411 | 8,0000 | 1,0972 |
| 69 | 1,7773 | 0,7677 | | 0,0411 | 7,0000 | 1,0972 |
| 70 | 1,7998 | 0,7643 | | 0,0411 | 7,0000 | 1,0972 |
| 71 | 1,8223 | 0,7608 | | 0,0411 | 8,0000 | 1,0972 |
| 72 | 1,8447 | 0,7574 | | 0,0411 | 7,0000 | 1,0972 |
| 73 | 1,8672 | 0,7540 | | 0,0411 | 8,0000 | 1,0972 |
| 74 | 1,8896 | 0,7506 | | 0,0411 | 10,0000 | 1,0972 |
| 75 | 1,9121 | 0,7472 | | 0,0411 | 7,0000 | 1,0972 |
| 76 | 1,9346 | 0,7439 | | 0,0411 | 7,0000 | 1,0972 |
| 77 | 1,9570 | 0,7405 | | 0,0411 | 7,0000 | 1,0972 |
| 78 | 1,9795 | 0,7372 | | 0,0411 | 8,0000 | 1,0972 |
| 79 | 2,0020 | 0,7339 | | 0,0411 | 7,0000 | 1,0972 |
| 80 | 2,0244 | 0,7306 | | 0,0411 | 8,0000 | 1,0972 |
| 81 | 2,0469 | 0,7273 | | 0,0411 | 8,0000 | 1,0972 |
| 82 | 2,0693 | 0,7240 | | 0,0411 | 8,0000 | 1,0972 |
| 83 | 2,0918 | 0,7207 | | 0,0411 | 7,0000 | 1,0972 |
| 84 | 2,1143 | 0,7175 | | 0,0411 | 10,0000 | 1,0972 |
| 85 | 2,1367 | 0,7143 | | 0,0411 | 7,0000 | 1,0972 |
| 86 | 2,1592 | 0,7110 | | 0,0411 | 7,0000 | 1,0972 |
| 87 | 2,1816 | 0,7078 | | 0,0411 | 7,0000 | 1,0972 |
| 88 | 2,2041 | 0,7047 | | 0,0411 | 7,0000 | 1,0972 |
| 89 | 2,2266 | 0,7015 | | 0,0411 | 8,0000 | 1,0972 |
| 90 | 2,2490 | 0,6983 | | 0,0411 | 7,0000 | 1,0972 |
| 91 | 2,2715 | 0,6952 | | 0,0411 | 7,0000 | 1,0972 |
| 92 | 2,2939 | 0,6921 | | 0,0411 | 8,0000 | 1,0972 |
| 93 | 2,3164 | 0,6889 | | 0,0411 | 7,0000 | 1,0972 |
| 94 | 2,3389 | 0,6858 | | 0,0411 | 8,0000 | 1,0972 |
| 95 | 2,3613 | 0,6827 | | 0,0411 | 10,0000 | 1,0972 |
| 96 | 2,3838 | 0,6797 | | 0,0411 | 10,0000 | 1,0972 |
| 97 | 2,4063 | 0,6766 | | 0,0411 | 8,0000 | 1,0972 |
| 98 | 2,4287 | 0,6736 | | 0,0411 | 22,0000 | 1,0972 |

| | 13-b-Final | 14-x column 1 | 15-y column 1 | 16 | 17-SumSq 1 | 18-Iterations 1 |
|---|---|---|---|---|---|---|
| 50 | 0,2009 | 1,3506 | 0,8321 | | 0,0352 | 7,0000 |
| 51 | 0,2009 | 1,3730 | 0,8285 | | 0,0352 | 7,0000 |
| 52 | 0,2009 | 1,3955 | 0,8250 | | 0,0352 | 10,0000 |
| 53 | 0,2009 | 1,4180 | 0,8215 | | 0,0352 | 7,0000 |
| 54 | 0,2009 | 1,4404 | 0,8180 | | 0,0352 | 7,0000 |
| 55 | 0,2009 | 1,4629 | 0,8145 | | 0,0352 | 7,0000 |
| 56 | 0,2009 | 1,4854 | 0,8111 | | 0,0352 | 7,0000 |
| 57 | 0,2009 | 1,5078 | 0,8076 | | 0,0352 | 7,0000 |
| 58 | 0,2009 | 1,5303 | 0,8042 | | 0,0352 | 10,0000 |
| 59 | 0,2009 | 1,5527 | 0,8008 | | 0,0352 | 7,0000 |
| 60 | 0,2009 | 1,5752 | 0,7974 | | 0,0352 | 7,0000 |
| 61 | 0,2009 | 1,5977 | 0,7940 | | 0,0352 | 10,0000 |
| 62 | 0,2009 | 1,6201 | 0,7906 | | 0,0352 | 7,0000 |
| 63 | 0,2009 | 1,6426 | 0,7872 | | 0,0352 | 7,0000 |
| 64 | 0,2009 | 1,6650 | 0,7839 | | 0,0352 | 7,0000 |
| 65 | 0,2009 | 1,6875 | 0,7806 | | 0,0352 | 10,0000 |
| 66 | 0,2009 | 1,7100 | 0,7772 | | 0,0352 | 7,0000 |
| 67 | 0,2009 | 1,7324 | 0,7739 | | 0,0352 | 10,0000 |
| 68 | 0,2009 | 1,7549 | 0,7706 | | 0,0352 | 10,0000 |
| 69 | 0,2009 | 1,7773 | 0,7674 | | 0,0352 | 7,0000 |
| 70 | 0,2009 | 1,7998 | 0,7641 | | 0,0352 | 7,0000 |
| 71 | 0,2009 | 1,8223 | 0,7608 | | 0,0352 | 7,0000 |
| 72 | 0,2009 | 1,8447 | 0,7576 | | 0,0352 | 7,0000 |
| 73 | 0,2009 | 1,8672 | 0,7544 | | 0,0352 | 10,0000 |
| 74 | 0,2009 | 1,8896 | 0,7512 | | 0,0352 | 7,0000 |
| 75 | 0,2009 | 1,9121 | 0,7480 | | 0,0352 | 10,0000 |
| 76 | 0,2009 | 1,9346 | 0,7448 | | 0,0352 | 7,0000 |
| 77 | 0,2009 | 1,9570 | 0,7416 | | 0,0352 | 12,0000 |
| 78 | 0,2009 | 1,9795 | 0,7385 | | 0,0352 | 7,0000 |
| 79 | 0,2009 | 2,0020 | 0,7353 | | 0,0352 | 7,0000 |
| 80 | 0,2009 | 2,0244 | 0,7322 | | 0,0352 | 10,0000 |
| 81 | 0,2009 | 2,0469 | 0,7291 | | 0,0352 | 7,0000 |
| 82 | 0,2009 | 2,0693 | 0,7260 | | 0,0352 | 7,0000 |
| 83 | 0,2009 | 2,0918 | 0,7229 | | 0,0352 | 7,0000 |
| 84 | 0,2009 | 2,1143 | 0,7198 | | 0,0352 | 7,0000 |
| 85 | 0,2009 | 2,1367 | 0,7168 | | 0,0352 | 7,0000 |
| 86 | 0,2009 | 2,1592 | 0,7137 | | 0,0352 | 7,0000 |
| 87 | 0,2009 | 2,1816 | 0,7107 | | 0,0352 | 7,0000 |
| 88 | 0,2009 | 2,2041 | 0,7077 | | 0,0352 | 7,0000 |
| 89 | 0,2009 | 2,2266 | 0,7047 | | 0,0352 | 7,0000 |
| 90 | 0,2009 | 2,2490 | 0,7017 | | 0,0352 | 7,0000 |
| 91 | 0,2009 | 2,2715 | 0,6987 | | 0,0352 | 10,0000 |
| 92 | 0,2009 | 2,2939 | 0,6957 | | 0,0352 | 10,0000 |
| 93 | 0,2009 | 2,3164 | 0,6928 | | 0,0352 | 7,0000 |
| 94 | 0,2009 | 2,3389 | 0,6898 | | 0,0352 | 7,0000 |
| 95 | 0,2009 | 2,3613 | 0,6869 | | 0,0352 | 7,0000 |
| 96 | 0,2009 | 2,3838 | 0,6840 | | 0,0352 | 7,0000 |
| 97 | 0,2009 | 2,4063 | 0,6810 | | 0,0352 | 7,0000 |
| 98 | 0,2009 | 2,4287 | 0,6781 | | 0,0352 | 8,0000 |

| | 19-a-Final 1 | 20-b-Final 1 | 21-x column 2 | 22-y column 2 | 23 | 24-SumSq 2 |
|---|---|---|---|---|---|---|
| 50 | 1,0751 | 0,1897 | 1,3506 | 0,7074 | | 0,0664 |
| 51 | 1,0751 | 0,1897 | 1,3730 | 0,7026 | | 0,0664 |
| 52 | 1,0751 | 0,1897 | 1,3955 | 0,6978 | | 0,0664 |
| 53 | 1,0751 | 0,1897 | 1,4180 | 0,6930 | | 0,0664 |
| 54 | 1,0751 | 0,1897 | 1,4404 | 0,6883 | | 0,0664 |
| 55 | 1,0751 | 0,1897 | 1,4629 | 0,6836 | | 0,0664 |
| 56 | 1,0751 | 0,1897 | 1,4854 | 0,6789 | | 0,0664 |
| 57 | 1,0751 | 0,1897 | 1,5078 | 0,6743 | | 0,0664 |
| 58 | 1,0751 | 0,1897 | 1,5303 | 0,6697 | | 0,0664 |
| 59 | 1,0751 | 0,1897 | 1,5527 | 0,6651 | | 0,0664 |
| 60 | 1,0751 | 0,1897 | 1,5752 | 0,6606 | | 0,0664 |
| 61 | 1,0751 | 0,1897 | 1,5977 | 0,6560 | | 0,0664 |
| 62 | 1,0751 | 0,1897 | 1,6201 | 0,6516 | | 0,0664 |
| 63 | 1,0751 | 0,1897 | 1,6426 | 0,6471 | | 0,0664 |
| 64 | 1,0751 | 0,1897 | 1,6650 | 0,6427 | | 0,0664 |
| 65 | 1,0751 | 0,1897 | 1,6875 | 0,6383 | | 0,0664 |
| 66 | 1,0751 | 0,1897 | 1,7100 | 0,6339 | | 0,0664 |
| 67 | 1,0751 | 0,1897 | 1,7324 | 0,6296 | | 0,0664 |
| 68 | 1,0751 | 0,1897 | 1,7549 | 0,6253 | | 0,0664 |
| 69 | 1,0751 | 0,1897 | 1,7773 | 0,6210 | | 0,0664 |
| 70 | 1,0751 | 0,1897 | 1,7998 | 0,6168 | | 0,0664 |
| 71 | 1,0751 | 0,1897 | 1,8223 | 0,6126 | | 0,0664 |
| 72 | 1,0751 | 0,1897 | 1,8447 | 0,6084 | | 0,0664 |
| 73 | 1,0751 | 0,1897 | 1,8672 | 0,6042 | | 0,0664 |
| 74 | 1,0751 | 0,1897 | 1,8896 | 0,6001 | | 0,0664 |
| 75 | 1,0751 | 0,1897 | 1,9121 | 0,5960 | | 0,0664 |
| 76 | 1,0751 | 0,1897 | 1,9346 | 0,5919 | | 0,0664 |
| 77 | 1,0751 | 0,1897 | 1,9570 | 0,5879 | | 0,0664 |
| 78 | 1,0751 | 0,1897 | 1,9795 | 0,5839 | | 0,0664 |
| 79 | 1,0751 | 0,1897 | 2,0020 | 0,5799 | | 0,0664 |
| 80 | 1,0751 | 0,1897 | 2,0244 | 0,5759 | | 0,0664 |
| 81 | 1,0751 | 0,1897 | 2,0469 | 0,5720 | | 0,0664 |
| 82 | 1,0751 | 0,1897 | 2,0693 | 0,5681 | | 0,0664 |
| 83 | 1,0751 | 0,1897 | 2,0918 | 0,5642 | | 0,0664 |
| 84 | 1,0751 | 0,1897 | 2,1143 | 0,5603 | | 0,0664 |
| 85 | 1,0751 | 0,1897 | 2,1367 | 0,5565 | | 0,0664 |
| 86 | 1,0751 | 0,1897 | 2,1592 | 0,5527 | | 0,0664 |
| 87 | 1,0751 | 0,1897 | 2,1816 | 0,5489 | | 0,0664 |
| 88 | 1,0751 | 0,1897 | 2,2041 | 0,5452 | | 0,0664 |
| 89 | 1,0751 | 0,1897 | 2,2266 | 0,5415 | | 0,0664 |
| 90 | 1,0751 | 0,1897 | 2,2490 | 0,5378 | | 0,0664 |
| 91 | 1,0751 | 0,1897 | 2,2715 | 0,5341 | | 0,0664 |
| 92 | 1,0751 | 0,1897 | 2,2939 | 0,5304 | | 0,0664 |
| 93 | 1,0751 | 0,1897 | 2,3164 | 0,5268 | | 0,0664 |
| 94 | 1,0751 | 0,1897 | 2,3389 | 0,5232 | | 0,0664 |
| 95 | 1,0751 | 0,1897 | 2,3613 | 0,5196 | | 0,0664 |
| 96 | 1,0751 | 0,1897 | 2,3838 | 0,5161 | | 0,0664 |
| 97 | 1,0751 | 0,1897 | 2,4063 | 0,5126 | | 0,0664 |
| 98 | 1,0751 | 0,1897 | 2,4287 | 0,5091 | | 0,0664 |

| | 25-Iterations 2 | 26-a-Final 2 | 27-b-Final 2 | 28-x column 3 | 29-y column 3 | 30 |
|---|---|---|---|---|---|---|
| 50 | 10,0000 | 1,0684 | 0,3052 | 1,3506 | 0,5984 | |
| 51 | 10,0000 | 1,0684 | 0,3052 | 1,3730 | 0,5933 | |
| 52 | 10,0000 | 1,0684 | 0,3052 | 1,3955 | 0,5881 | |
| 53 | 11,0000 | 1,0684 | 0,3052 | 1,4180 | 0,5831 | |
| 54 | 14,0000 | 1,0684 | 0,3052 | 1,4404 | 0,5781 | |
| 55 | 10,0000 | 1,0684 | 0,3052 | 1,4629 | 0,5731 | |
| 56 | 10,0000 | 1,0684 | 0,3052 | 1,4854 | 0,5681 | |
| 57 | 10,0000 | 1,0684 | 0,3052 | 1,5078 | 0,5632 | |
| 58 | 10,0000 | 1,0684 | 0,3052 | 1,5303 | 0,5584 | |
| 59 | 14,0000 | 1,0684 | 0,3052 | 1,5527 | 0,5536 | |
| 60 | 11,0000 | 1,0684 | 0,3052 | 1,5752 | 0,5488 | |
| 61 | 10,0000 | 1,0684 | 0,3052 | 1,5977 | 0,5441 | |
| 62 | 10,0000 | 1,0684 | 0,3052 | 1,6201 | 0,5394 | |
| 63 | 10,0000 | 1,0684 | 0,3052 | 1,6426 | 0,5347 | |
| 64 | 10,0000 | 1,0684 | 0,3052 | 1,6650 | 0,5301 | |
| 65 | 10,0000 | 1,0684 | 0,3052 | 1,6875 | 0,5256 | |
| 66 | 10,0000 | 1,0684 | 0,3052 | 1,7100 | 0,5210 | |
| 67 | 13,0000 | 1,0684 | 0,3052 | 1,7324 | 0,5165 | |
| 68 | 13,0000 | 1,0684 | 0,3052 | 1,7549 | 0,5121 | |
| 69 | 13,0000 | 1,0684 | 0,3052 | 1,7773 | 0,5077 | |
| 70 | 13,0000 | 1,0684 | 0,3052 | 1,7998 | 0,5033 | |
| 71 | 13,0000 | 1,0684 | 0,3052 | 1,8223 | 0,4990 | |
| 72 | 13,0000 | 1,0684 | 0,3052 | 1,8447 | 0,4947 | |
| 73 | 10,0000 | 1,0684 | 0,3052 | 1,8672 | 0,4904 | |
| 74 | 13,0000 | 1,0684 | 0,3052 | 1,8896 | 0,4862 | |
| 75 | 13,0000 | 1,0684 | 0,3052 | 1,9121 | 0,4820 | |
| 76 | 13,0000 | 1,0684 | 0,3052 | 1,9346 | 0,4778 | |
| 77 | 10,0000 | 1,0684 | 0,3052 | 1,9570 | 0,4737 | |
| 78 | 13,0000 | 1,0684 | 0,3052 | 1,9795 | 0,4696 | |
| 79 | 10,0000 | 1,0684 | 0,3052 | 2,0020 | 0,4656 | |
| 80 | 13,0000 | 1,0684 | 0,3052 | 2,0244 | 0,4616 | |
| 81 | 10,0000 | 1,0684 | 0,3052 | 2,0469 | 0,4576 | |
| 82 | 13,0000 | 1,0684 | 0,3052 | 2,0693 | 0,4536 | |
| 83 | 10,0000 | 1,0684 | 0,3052 | 2,0918 | 0,4497 | |
| 84 | 10,0000 | 1,0684 | 0,3052 | 2,1143 | 0,4459 | |
| 85 | 10,0000 | 1,0684 | 0,3052 | 2,1367 | 0,4420 | |
| 86 | 10,0000 | 1,0684 | 0,3052 | 2,1592 | 0,4382 | |
| 87 | 10,0000 | 1,0684 | 0,3052 | 2,1816 | 0,4344 | |
| 88 | 11,0000 | 1,0684 | 0,3052 | 2,2041 | 0,4307 | |
| 89 | 14,0000 | 1,0684 | 0,3052 | 2,2266 | 0,4270 | |
| 90 | 10,0000 | 1,0684 | 0,3052 | 2,2490 | 0,4233 | |
| 91 | 10,0000 | 1,0684 | 0,3052 | 2,2715 | 0,4196 | |
| 92 | 10,0000 | 1,0684 | 0,3052 | 2,2939 | 0,4160 | |
| 93 | 11,0000 | 1,0684 | 0,3052 | 2,3164 | 0,4124 | |
| 94 | 11,0000 | 1,0684 | 0,3052 | 2,3389 | 0,4089 | |
| 95 | 10,0000 | 1,0684 | 0,3052 | 2,3613 | 0,4054 | |
| 96 | 14,0000 | 1,0684 | 0,3052 | 2,3838 | 0,4019 | |
| 97 | 9,0000 | 1,0684 | 0,3052 | 2,4063 | 0,3984 | |
| 98 | 11,0000 | 1,0684 | 0,3052 | 2,4287 | 0,3950 | |

| | 31-SumSq 3 | 32-Iterations 3 | 33-a-Final 3 | 34-b-Final 3 | 35-x column 4 | 36-y column 4 |
|---|---|---|---|---|---|---|
| 50 | 0,0429 | 11,0000 | 1,0070 | 0,3854 | 1,3506 | 0,5984 |
| 51 | 0,0429 | 11,0000 | 1,0070 | 0,3854 | 1,3730 | 0,5933 |
| 52 | 0,0429 | 15,0000 | 1,0070 | 0,3854 | 1,3955 | 0,5881 |
| 53 | 0,0429 | 11,0000 | 1,0070 | 0,3854 | 1,4180 | 0,5831 |
| 54 | 0,0429 | 11,0000 | 1,0070 | 0,3854 | 1,4404 | 0,5781 |
| 55 | 0,0429 | 12,0000 | 1,0070 | 0,3854 | 1,4629 | 0,5731 |
| 56 | 0,0429 | 11,0000 | 1,0070 | 0,3854 | 1,4854 | 0,5681 |
| 57 | 0,0429 | 11,0000 | 1,0070 | 0,3854 | 1,5078 | 0,5632 |
| 58 | 0,0429 | 11,0000 | 1,0070 | 0,3854 | 1,5303 | 0,5584 |
| 59 | 0,0429 | 11,0000 | 1,0070 | 0,3854 | 1,5527 | 0,5536 |
| 60 | 0,0429 | 12,0000 | 1,0070 | 0,3854 | 1,5752 | 0,5488 |
| 61 | 0,0429 | 11,0000 | 1,0070 | 0,3854 | 1,5977 | 0,5441 |
| 62 | 0,0429 | 16,0000 | 1,0070 | 0,3854 | 1,6201 | 0,5394 |
| 63 | 0,0429 | 12,0000 | 1,0070 | 0,3854 | 1,6426 | 0,5347 |
| 64 | 0,0429 | 16,0000 | 1,0070 | 0,3854 | 1,6650 | 0,5301 |
| 65 | 0,0429 | 12,0000 | 1,0070 | 0,3854 | 1,6875 | 0,5256 |
| 66 | 0,0429 | 16,0000 | 1,0070 | 0,3854 | 1,7100 | 0,5210 |
| 67 | 0,0429 | 11,0000 | 1,0070 | 0,3854 | 1,7324 | 0,5165 |
| 68 | 0,0429 | 11,0000 | 1,0070 | 0,3854 | 1,7549 | 0,5121 |
| 69 | 0,0429 | 16,0000 | 1,0070 | 0,3854 | 1,7773 | 0,5077 |
| 70 | 0,0429 | 11,0000 | 1,0070 | 0,3854 | 1,7998 | 0,5033 |
| 71 | 0,0429 | 11,0000 | 1,0070 | 0,3854 | 1,8223 | 0,4990 |
| 72 | 0,0429 | 10,0000 | 1,0070 | 0,3854 | 1,8447 | 0,4947 |
| 73 | 0,0429 | 11,0000 | 1,0070 | 0,3854 | 1,8672 | 0,4904 |
| 74 | 0,0429 | 9,0000 | 1,0070 | 0,3854 | 1,8896 | 0,4862 |
| 75 | 0,0429 | 15,0000 | 1,0070 | 0,3854 | 1,9121 | 0,4820 |
| 76 | 0,0429 | 16,0000 | 1,0070 | 0,3854 | 1,9346 | 0,4778 |
| 77 | 0,0429 | 11,0000 | 1,0070 | 0,3854 | 1,9570 | 0,4737 |
| 78 | 0,0429 | 11,0000 | 1,0070 | 0,3854 | 1,9795 | 0,4696 |
| 79 | 0,0429 | 11,0000 | 1,0070 | 0,3854 | 2,0020 | 0,4656 |
| 80 | 0,0429 | 11,0000 | 1,0070 | 0,3854 | 2,0244 | 0,4616 |
| 81 | 0,0429 | 11,0000 | 1,0070 | 0,3854 | 2,0469 | 0,4576 |
| 82 | 0,0429 | 11,0000 | 1,0070 | 0,3854 | 2,0693 | 0,4536 |
| 83 | 0,0429 | 12,0000 | 1,0070 | 0,3854 | 2,0918 | 0,4497 |
| 84 | 0,0429 | 11,0000 | 1,0070 | 0,3854 | 2,1143 | 0,4459 |
| 85 | 0,0429 | 11,0000 | 1,0070 | 0,3854 | 2,1367 | 0,4420 |
| 86 | 0,0429 | 15,0000 | 1,0070 | 0,3854 | 2,1592 | 0,4382 |
| 87 | 0,0429 | 12,0000 | 1,0070 | 0,3854 | 2,1816 | 0,4344 |
| 88 | 0,0429 | 11,0000 | 1,0070 | 0,3854 | 2,2041 | 0,4307 |
| 89 | 0,0429 | 11,0000 | 1,0070 | 0,3854 | 2,2266 | 0,4270 |
| 90 | 0,0429 | 12,0000 | 1,0070 | 0,3854 | 2,2490 | 0,4233 |
| 91 | 0,0429 | 15,0000 | 1,0070 | 0,3854 | 2,2715 | 0,4196 |
| 92 | 0,0429 | 10,0000 | 1,0070 | 0,3854 | 2,2939 | 0,4160 |
| 93 | 0,0429 | 10,0000 | 1,0070 | 0,3854 | 2,3164 | 0,4124 |
| 94 | 0,0429 | 11,0000 | 1,0070 | 0,3854 | 2,3389 | 0,4089 |
| 95 | 0,0429 | 14,0000 | 1,0070 | 0,3854 | 2,3613 | 0,4054 |
| 96 | 0,0429 | 12,0000 | 1,0070 | 0,3854 | 2,3838 | 0,4019 |
| 97 | 0,0429 | 11,0000 | 1,0070 | 0,3854 | 2,4063 | 0,3984 |
| 98 | 0,0429 | 11,0000 | 1,0070 | 0,3854 | 2,4287 | 0,3950 |

| | 37 | 38-SumSq 4 | 39-Iterations 4 | 40-a-Final 4 | 41-b-Final 4 | 42-x column 5 |
|---|---|---|---|---|---|---|
| 50 | | 0,0429 | 10,0000 | 1,0070 | 0,3854 | 1,3506 |
| 51 | | 0,0429 | 11,0000 | 1,0070 | 0,3854 | 1,3730 |
| 52 | | 0,0429 | 10,0000 | 1,0070 | 0,3854 | 1,3955 |
| 53 | | 0,0429 | 11,0000 | 1,0070 | 0,3854 | 1,4180 |
| 54 | | 0,0429 | 10,0000 | 1,0070 | 0,3854 | 1,4404 |
| 55 | | 0,0429 | 10,0000 | 1,0070 | 0,3854 | 1,4629 |
| 56 | | 0,0429 | 10,0000 | 1,0070 | 0,3854 | 1,4854 |
| 57 | | 0,0429 | 9,0000 | 1,0070 | 0,3854 | 1,5078 |
| 58 | | 0,0429 | 10,0000 | 1,0070 | 0,3854 | 1,5303 |
| 59 | | 0,0429 | 10,0000 | 1,0070 | 0,3854 | 1,5527 |
| 60 | | 0,0429 | 10,0000 | 1,0070 | 0,3854 | 1,5752 |
| 61 | | 0,0429 | 11,0000 | 1,0070 | 0,3854 | 1,5977 |
| 62 | | 0,0429 | 10,0000 | 1,0070 | 0,3854 | 1,6201 |
| 63 | | 0,0429 | 11,0000 | 1,0070 | 0,3854 | 1,6426 |
| 64 | | 0,0429 | 10,0000 | 1,0070 | 0,3854 | 1,6650 |
| 65 | | 0,0429 | 10,0000 | 1,0070 | 0,3854 | 1,6875 |
| 66 | | 0,0429 | 10,0000 | 1,0070 | 0,3854 | 1,7100 |
| 67 | | 0,0429 | 10,0000 | 1,0070 | 0,3854 | 1,7324 |
| 68 | | 0,0429 | 11,0000 | 1,0070 | 0,3854 | 1,7549 |
| 69 | | 0,0429 | 10,0000 | 1,0070 | 0,3854 | 1,7773 |
| 70 | | 0,0429 | 9,0000 | 1,0070 | 0,3854 | 1,7998 |
| 71 | | 0,0429 | 10,0000 | 1,0070 | 0,3854 | 1,8223 |
| 72 | | 0,0429 | 11,0000 | 1,0070 | 0,3854 | 1,8447 |
| 73 | | 0,0429 | 10,0000 | 1,0070 | 0,3854 | 1,8672 |
| 74 | | 0,0429 | 10,0000 | 1,0070 | 0,3854 | 1,8896 |
| 75 | | 0,0429 | 10,0000 | 1,0070 | 0,3854 | 1,9121 |
| 76 | | 0,0429 | 11,0000 | 1,0070 | 0,3854 | 1,9346 |
| 77 | | 0,0429 | 10,0000 | 1,0070 | 0,3854 | 1,9570 |
| 78 | | 0,0429 | 10,0000 | 1,0070 | 0,3854 | 1,9795 |
| 79 | | 0,0429 | 10,0000 | 1,0070 | 0,3854 | 2,0020 |
| 80 | | 0,0429 | 10,0000 | 1,0070 | 0,3854 | 2,0244 |
| 81 | | 0,0429 | 11,0000 | 1,0070 | 0,3854 | 2,0469 |
| 82 | | 0,0429 | 10,0000 | 1,0070 | 0,3854 | 2,0693 |
| 83 | | 0,0429 | 10,0000 | 1,0070 | 0,3854 | 2,0918 |
| 84 | | 0,0429 | 10,0000 | 1,0070 | 0,3854 | 2,1143 |
| 85 | | 0,0429 | 10,0000 | 1,0070 | 0,3854 | 2,1367 |
| 86 | | 0,0429 | 10,0000 | 1,0070 | 0,3854 | 2,1592 |
| 87 | | 0,0429 | 10,0000 | 1,0070 | 0,3854 | 2,1816 |
| 88 | | 0,0429 | 10,0000 | 1,0070 | 0,3854 | 2,2041 |
| 89 | | 0,0429 | 11,0000 | 1,0070 | 0,3854 | 2,2266 |
| 90 | | 0,0429 | 10,0000 | 1,0070 | 0,3854 | 2,2490 |
| 91 | | 0,0429 | 10,0000 | 1,0070 | 0,3854 | 2,2715 |
| 92 | | 0,0429 | 10,0000 | 1,0070 | 0,3854 | 2,2939 |
| 93 | | 0,0429 | 10,0000 | 1,0070 | 0,3854 | 2,3164 |
| 94 | | 0,0429 | 10,0000 | 1,0070 | 0,3854 | 2,3389 |
| 95 | | 0,0429 | 11,0000 | 1,0070 | 0,3854 | 2,3613 |
| 96 | | 0,0429 | 10,0000 | 1,0070 | 0,3854 | 2,3838 |
| 97 | | 0,0429 | 10,0000 | 1,0070 | 0,3854 | 2,4063 |
| 98 | | 0,0429 | 10,0000 | 1,0070 | 0,3854 | 2,4287 |

| | 43-y column 5 | 44 | 45-SumSq 5 | 46-Iterations 5 | 47-a-Final 5 | 48-b-Final 5 |
|---|---|---|---|---|---|---|
| 50 | 0,7269 | | 0,1194 | 9,0000 | 1,0036 | 0,2389 |
| 51 | 0,7230 | | 0,1194 | 9,0000 | 1,0036 | 0,2389 |
| 52 | 0,7191 | | 0,1194 | 10,0000 | 1,0036 | 0,2389 |
| 53 | 0,7153 | | 0,1194 | 9,0000 | 1,0036 | 0,2389 |
| 54 | 0,7114 | | 0,1194 | 10,0000 | 1,0036 | 0,2389 |
| 55 | 0,7076 | | 0,1194 | 10,0000 | 1,0036 | 0,2389 |
| 56 | 0,7038 | | 0,1194 | 10,0000 | 1,0036 | 0,2389 |
| 57 | 0,7001 | | 0,1194 | 9,0000 | 1,0036 | 0,2389 |
| 58 | 0,6963 | | 0,1194 | 9,0000 | 1,0036 | 0,2389 |
| 59 | 0,6926 | | 0,1194 | 9,0000 | 1,0036 | 0,2389 |
| 60 | 0,6889 | | 0,1194 | 10,0000 | 1,0036 | 0,2389 |
| 61 | 0,6852 | | 0,1194 | 9,0000 | 1,0036 | 0,2389 |
| 62 | 0,6815 | | 0,1194 | 10,0000 | 1,0036 | 0,2389 |
| 63 | 0,6779 | | 0,1194 | 9,0000 | 1,0036 | 0,2389 |
| 64 | 0,6743 | | 0,1194 | 10,0000 | 1,0036 | 0,2389 |
| 65 | 0,6706 | | 0,1194 | 9,0000 | 1,0036 | 0,2389 |
| 66 | 0,6671 | | 0,1194 | 10,0000 | 1,0036 | 0,2389 |
| 67 | 0,6635 | | 0,1194 | 9,0000 | 1,0036 | 0,2389 |
| 68 | 0,6599 | | 0,1194 | 9,0000 | 1,0036 | 0,2389 |
| 69 | 0,6564 | | 0,1194 | 9,0000 | 1,0036 | 0,2389 |
| 70 | 0,6529 | | 0,1194 | 9,0000 | 1,0036 | 0,2389 |
| 71 | 0,6494 | | 0,1194 | 8,0000 | 1,0036 | 0,2389 |
| 72 | 0,6459 | | 0,1194 | 10,0000 | 1,0036 | 0,2389 |
| 73 | 0,6425 | | 0,1194 | 9,0000 | 1,0036 | 0,2389 |
| 74 | 0,6390 | | 0,1194 | 10,0000 | 1,0036 | 0,2389 |
| 75 | 0,6356 | | 0,1194 | 9,0000 | 1,0036 | 0,2389 |
| 76 | 0,6322 | | 0,1194 | 9,0000 | 1,0036 | 0,2389 |
| 77 | 0,6288 | | 0,1194 | 10,0000 | 1,0036 | 0,2389 |
| 78 | 0,6255 | | 0,1194 | 10,0000 | 1,0036 | 0,2389 |
| 79 | 0,6221 | | 0,1194 | 10,0000 | 1,0036 | 0,2389 |
| 80 | 0,6188 | | 0,1194 | 9,0000 | 1,0036 | 0,2389 |
| 81 | 0,6155 | | 0,1194 | 9,0000 | 1,0036 | 0,2389 |
| 82 | 0,6122 | | 0,1194 | 9,0000 | 1,0036 | 0,2389 |
| 83 | 0,6089 | | 0,1194 | 10,0000 | 1,0036 | 0,2389 |
| 84 | 0,6056 | | 0,1194 | 10,0000 | 1,0036 | 0,2389 |
| 85 | 0,6024 | | 0,1194 | 8,0000 | 1,0036 | 0,2389 |
| 86 | 0,5992 | | 0,1194 | 9,0000 | 1,0036 | 0,2389 |
| 87 | 0,5960 | | 0,1194 | 9,0000 | 1,0036 | 0,2389 |
| 88 | 0,5928 | | 0,1194 | 7,0000 | 1,0036 | 0,2389 |
| 89 | 0,5896 | | 0,1194 | 9,0000 | 1,0036 | 0,2389 |
| 90 | 0,5865 | | 0,1194 | 10,0000 | 1,0036 | 0,2389 |
| 91 | 0,5833 | | 0,1194 | 9,0000 | 1,0036 | 0,2389 |
| 92 | 0,5802 | | 0,1194 | 10,0000 | 1,0036 | 0,2389 |
| 93 | 0,5771 | | 0,1194 | 9,0000 | 1,0036 | 0,2389 |
| 94 | 0,5740 | | 0,1194 | 9,0000 | 1,0036 | 0,2389 |
| 95 | 0,5709 | | 0,1194 | 9,0000 | 1,0036 | 0,2389 |
| 96 | 0,5679 | | 0,1194 | 9,0000 | 1,0036 | 0,2389 |
| 97 | 0,5648 | | 0,1194 | 9,0000 | 1,0036 | 0,2389 |
| 98 | 0,5618 | | 0,1194 | 10,0000 | 1,0036 | 0,2389 |

| | 49-x column 6 | 50-y column 6 | 51 | 52-SumSq 6 | 53-Iterations 6 | 54-a-Final 6 |
|---|---|---|---|---|---|---|
| 50 | 1,3506 | 0,6227 | | 0,0865 | 9,0000 | 0,9419 |
| 51 | 1,3730 | 0,6184 | | 0,0865 | 11,0000 | 0,9419 |
| 52 | 1,3955 | 0,6142 | | 0,0865 | 11,0000 | 0,9419 |
| 53 | 1,4180 | 0,6100 | | 0,0865 | 10,0000 | 0,9419 |
| 54 | 1,4404 | 0,6058 | | 0,0865 | 10,0000 | 0,9419 |
| 55 | 1,4629 | 0,6016 | | 0,0865 | 10,0000 | 0,9419 |
| 56 | 1,4854 | 0,5975 | | 0,0865 | 10,0000 | 0,9419 |
| 57 | 1,5078 | 0,5934 | | 0,0865 | 8,0000 | 0,9419 |
| 58 | 1,5303 | 0,5893 | | 0,0865 | 10,0000 | 0,9419 |
| 59 | 1,5527 | 0,5853 | | 0,0865 | 11,0000 | 0,9419 |
| 60 | 1,5752 | 0,5813 | | 0,0865 | 10,0000 | 0,9419 |
| 61 | 1,5977 | 0,5773 | | 0,0865 | 11,0000 | 0,9419 |
| 62 | 1,6201 | 0,5733 | | 0,0865 | 10,0000 | 0,9419 |
| 63 | 1,6426 | 0,5694 | | 0,0865 | 10,0000 | 0,9419 |
| 64 | 1,6650 | 0,5655 | | 0,0865 | 9,0000 | 0,9419 |
| 65 | 1,6875 | 0,5616 | | 0,0865 | 11,0000 | 0,9419 |
| 66 | 1,7100 | 0,5578 | | 0,0865 | 10,0000 | 0,9419 |
| 67 | 1,7324 | 0,5539 | | 0,0865 | 10,0000 | 0,9419 |
| 68 | 1,7549 | 0,5501 | | 0,0865 | 10,0000 | 0,9419 |
| 69 | 1,7773 | 0,5464 | | 0,0865 | 11,0000 | 0,9419 |
| 70 | 1,7998 | 0,5426 | | 0,0865 | 10,0000 | 0,9419 |
| 71 | 1,8223 | 0,5389 | | 0,0865 | 10,0000 | 0,9419 |
| 72 | 1,8447 | 0,5352 | | 0,0865 | 11,0000 | 0,9419 |
| 73 | 1,8672 | 0,5315 | | 0,0865 | 10,0000 | 0,9419 |
| 74 | 1,8896 | 0,5279 | | 0,0865 | 10,0000 | 0,9419 |
| 75 | 1,9121 | 0,5243 | | 0,0865 | 11,0000 | 0,9419 |
| 76 | 1,9346 | 0,5207 | | 0,0865 | 10,0000 | 0,9419 |
| 77 | 1,9570 | 0,5171 | | 0,0865 | 10,0000 | 0,9419 |
| 78 | 1,9795 | 0,5135 | | 0,0865 | 8,0000 | 0,9419 |
| 79 | 2,0020 | 0,5100 | | 0,0865 | 10,0000 | 0,9419 |
| 80 | 2,0244 | 0,5065 | | 0,0865 | 10,0000 | 0,9419 |
| 81 | 2,0469 | 0,5031 | | 0,0865 | 11,0000 | 0,9419 |
| 82 | 2,0693 | 0,4996 | | 0,0865 | 10,0000 | 0,9419 |
| 83 | 2,0918 | 0,4962 | | 0,0865 | 10,0000 | 0,9419 |
| 84 | 2,1143 | 0,4928 | | 0,0865 | 11,0000 | 0,9419 |
| 85 | 2,1367 | 0,4894 | | 0,0865 | 9,0000 | 0,9419 |
| 86 | 2,1592 | 0,4860 | | 0,0865 | 10,0000 | 0,9419 |
| 87 | 2,1816 | 0,4827 | | 0,0865 | 10,0000 | 0,9419 |
| 88 | 2,2041 | 0,4794 | | 0,0865 | 10,0000 | 0,9419 |
| 89 | 2,2266 | 0,4761 | | 0,0865 | 10,0000 | 0,9419 |
| 90 | 2,2490 | 0,4728 | | 0,0865 | 10,0000 | 0,9419 |
| 91 | 2,2715 | 0,4696 | | 0,0865 | 10,0000 | 0,9419 |
| 92 | 2,2939 | 0,4664 | | 0,0865 | 10,0000 | 0,9419 |
| 93 | 2,3164 | 0,4632 | | 0,0865 | 11,0000 | 0,9419 |
| 94 | 2,3389 | 0,4600 | | 0,0865 | 9,0000 | 0,9419 |
| 95 | 2,3613 | 0,4568 | | 0,0865 | 10,0000 | 0,9419 |
| 96 | 2,3838 | 0,4537 | | 0,0865 | 10,0000 | 0,9419 |
| 97 | 2,4063 | 0,4506 | | 0,0865 | 11,0000 | 0,9419 |
| 98 | 2,4287 | 0,4475 | | 0,0865 | 10,0000 | 0,9419 |

| | 55-b-Final 6 |
|---|---|
| 50 | 0,3064 |
| 51 | 0,3064 |
| 52 | 0,3064 |
| 53 | 0,3064 |
| 54 | 0,3064 |
| 55 | 0,3064 |
| 56 | 0,3064 |
| 57 | 0,3064 |
| 58 | 0,3064 |
| 59 | 0,3064 |
| 60 | 0,3064 |
| 61 | 0,3064 |
| 62 | 0,3064 |
| 63 | 0,3064 |
| 64 | 0,3064 |
| 65 | 0,3064 |
| 66 | 0,3064 |
| 67 | 0,3064 |
| 68 | 0,3064 |
| 69 | 0,3064 |
| 70 | 0,3064 |
| 71 | 0,3064 |
| 72 | 0,3064 |
| 73 | 0,3064 |
| 74 | 0,3064 |
| 75 | 0,3064 |
| 76 | 0,3064 |
| 77 | 0,3064 |
| 78 | 0,3064 |
| 79 | 0,3064 |
| 80 | 0,3064 |
| 81 | 0,3064 |
| 82 | 0,3064 |
| 83 | 0,3064 |
| 84 | 0,3064 |
| 85 | 0,3064 |
| 86 | 0,3064 |
| 87 | 0,3064 |
| 88 | 0,3064 |
| 89 | 0,3064 |
| 90 | 0,3064 |
| 91 | 0,3064 |
| 92 | 0,3064 |
| 93 | 0,3064 |
| 94 | 0,3064 |
| 95 | 0,3064 |
| 96 | 0,3064 |
| 97 | 0,3064 |
| 98 | 0,3064 |

| | 1 | 2 | 3 | 4 | 5 | 6 |
|---|---|---|---|---|---|---|
| 99 | | | | | | |
| 100 | | | | | | |
| 101 | | | | | | |
| 102 | | | | | | |
| 103 | | | | | | |
| 104 | | | | | | |
| 105 | | | | | | |
| 106 | | | | | | |
| 107 | | | | | | |
| 108 | | | | | | |
| 109 | | | | | | |
| 110 | | | | | | |
| 111 | | | | | | |
| 112 | | | | | | |
| 113 | | | | | | |
| 114 | | | | | | |
| 115 | | | | | | |
| 116 | | | | | | |
| 117 | | | | | | |
| 118 | | | | | | |
| 119 | | | | | | |
| 120 | | | | | | |
| 121 | | | | | | |
| 122 | | | | | | |
| 123 | | | | | | |
| 124 | | | | | | |
| 125 | | | | | | |
| 126 | | | | | | |
| 127 | | | | | | |
| 128 | | | | | | |
| 129 | | | | | | |
| 130 | | | | | | |
| 131 | | | | | | |
| 132 | | | | | | |
| 133 | | | | | | |
| 134 | | | | | | |
| 135 | | | | | | |
| 136 | | | | | | |
| 137 | | | | | | |
| 138 | | | | | | |
| 139 | | | | | | |
| 140 | | | | | | |
| 141 | | | | | | |
| 142 | | | | | | |
| 143 | | | | | | |
| 144 | | | | | | |
| 145 | | | | | | |
| 146 | | | | | | |
| 147 | | | | | | |

| | 7-x column | 8-y column | 9 | 10-SumSq | 11-Iterations | 12-a-Final |
|---|---|---|---|---|---|---|
| 99 | 2,4512 | 0,6705 | | 0,0411 | 7,0000 | 1,0972 |
| 100 | 2,4736 | 0,6675 | | 0,0411 | 10,0000 | 1,0972 |
| 101 | 2,4961 | 0,6645 | | 0,0411 | 8,0000 | 1,0972 |
| 102 | 2,5186 | 0,6615 | | 0,0411 | 7,0000 | 1,0972 |
| 103 | 2,5410 | 0,6585 | | 0,0411 | 10,0000 | 1,0972 |
| 104 | 2,5635 | 0,6556 | | 0,0411 | 7,0000 | 1,0972 |
| 105 | 2,5859 | 0,6526 | | 0,0411 | 7,0000 | 1,0972 |
| 106 | 2,6084 | 0,6497 | | 0,0411 | 10,0000 | 1,0972 |
| 107 | 2,6309 | 0,6468 | | 0,0411 | 7,0000 | 1,0972 |
| 108 | 2,6533 | 0,6439 | | 0,0411 | 7,0000 | 1,0972 |
| 109 | 2,6758 | 0,6410 | | 0,0411 | 7,0000 | 1,0972 |
| 110 | 2,6982 | 0,6381 | | 0,0411 | 10,0000 | 1,0972 |
| 111 | 2,7207 | 0,6352 | | 0,0411 | 10,0000 | 1,0972 |
| 112 | 2,7432 | 0,6323 | | 0,0411 | 10,0000 | 1,0972 |
| 113 | 2,7656 | 0,6295 | | 0,0411 | 7,0000 | 1,0972 |
| 114 | 2,7881 | 0,6267 | | 0,0411 | 7,0000 | 1,0972 |
| 115 | 2,8105 | 0,6238 | | 0,0411 | 8,0000 | 1,0972 |
| 116 | 2,8330 | 0,6210 | | 0,0411 | 7,0000 | 1,0972 |
| 117 | 2,8555 | 0,6182 | | 0,0411 | 10,0000 | 1,0972 |
| 118 | 2,8779 | 0,6154 | | 0,0411 | 10,0000 | 1,0972 |
| 119 | 2,9004 | 0,6127 | | 0,0411 | 7,0000 | 1,0972 |
| 120 | 2,9229 | 0,6099 | | 0,0411 | 7,0000 | 1,0972 |
| 121 | 2,9453 | 0,6072 | | 0,0411 | 7,0000 | 1,0972 |
| 122 | 2,9678 | 0,6044 | | 0,0411 | 7,0000 | 1,0972 |
| 123 | 2,9902 | 0,6017 | | 0,0411 | 7,0000 | 1,0972 |
| 124 | 3,0127 | 0,5990 | | 0,0411 | 7,0000 | 1,0972 |
| 125 | 3,0352 | 0,5963 | | 0,0411 | 10,0000 | 1,0972 |
| 126 | 3,0576 | 0,5936 | | 0,0411 | 8,0000 | 1,0972 |
| 127 | 3,0801 | 0,5910 | | 0,0411 | 8,0000 | 1,0972 |
| 128 | 3,1025 | 0,5883 | | 0,0411 | 7,0000 | 1,0972 |
| 129 | 3,1250 | 0,5856 | | 0,0411 | 10,0000 | 1,0972 |
| 130 | 3,1475 | 0,5830 | | 0,0411 | 7,0000 | 1,0972 |
| 131 | 3,1699 | 0,5804 | | 0,0411 | 7,0000 | 1,0972 |
| 132 | 3,1924 | 0,5778 | | 0,0411 | 7,0000 | 1,0972 |
| 133 | 3,2148 | 0,5752 | | 0,0411 | 8,0000 | 1,0972 |
| 134 | 3,2373 | 0,5726 | | 0,0411 | 10,0000 | 1,0972 |
| 135 | 3,2598 | 0,5700 | | 0,0411 | 7,0000 | 1,0972 |
| 136 | 3,2822 | 0,5674 | | 0,0411 | 7,0000 | 1,0972 |
| 137 | 3,3047 | 0,5649 | | 0,0411 | 33,0000 | 1,0972 |
| 138 | 3,3271 | 0,5623 | | 0,0411 | 8,0000 | 1,0972 |
| 139 | 3,3496 | 0,5598 | | 0,0411 | 7,0000 | 1,0972 |
| 140 | 3,3721 | 0,5573 | | 0,0411 | 7,0000 | 1,0972 |
| 141 | 3,3945 | 0,5548 | | 0,0411 | 8,0000 | 1,0972 |
| 142 | 3,4170 | 0,5523 | | 0,0411 | 7,0000 | 1,0972 |
| 143 | 3,4395 | 0,5498 | | 0,0411 | 8,0000 | 1,0972 |
| 144 | 3,4619 | 0,5473 | | 0,0411 | 7,0000 | 1,0972 |
| 145 | 3,4844 | 0,5449 | | 0,0411 | 7,0000 | 1,0972 |
| 146 | 3,5068 | 0,5424 | | 0,0411 | 7,0000 | 1,0972 |
| 147 | 3,5293 | 0,5400 | | 0,0411 | 8,0000 | 1,0972 |

| | 13-b-Final | 14-x column 1 | 15-y column 1 | 16 | 17-SumSq 1 | 18-Iterations 1 |
|---|---|---|---|---|---|---|
| 99 | 0,2009 | 2,4512 | 0,6753 | | 0,0352 | 7,0000 |
| 100 | 0,2009 | 2,4736 | 0,6724 | | 0,0352 | 19,0000 |
| 101 | 0,2009 | 2,4961 | 0,6695 | | 0,0352 | 10,0000 |
| 102 | 0,2009 | 2,5186 | 0,6667 | | 0,0352 | 10,0000 |
| 103 | 0,2009 | 2,5410 | 0,6638 | | 0,0352 | 7,0000 |
| 104 | 0,2009 | 2,5635 | 0,6610 | | 0,0352 | 7,0000 |
| 105 | 0,2009 | 2,5859 | 0,6582 | | 0,0352 | 7,0000 |
| 106 | 0,2009 | 2,6084 | 0,6554 | | 0,0352 | 10,0000 |
| 107 | 0,2009 | 2,6309 | 0,6526 | | 0,0352 | 7,0000 |
| 108 | 0,2009 | 2,6533 | 0,6499 | | 0,0352 | 7,0000 |
| 109 | 0,2009 | 2,6758 | 0,6471 | | 0,0352 | 10,0000 |
| 110 | 0,2009 | 2,6982 | 0,6443 | | 0,0352 | 7,0000 |
| 111 | 0,2009 | 2,7207 | 0,6416 | | 0,0352 | 7,0000 |
| 112 | 0,2009 | 2,7432 | 0,6389 | | 0,0352 | 10,0000 |
| 113 | 0,2009 | 2,7656 | 0,6362 | | 0,0352 | 7,0000 |
| 114 | 0,2009 | 2,7881 | 0,6334 | | 0,0352 | 8,0000 |
| 115 | 0,2009 | 2,8105 | 0,6308 | | 0,0352 | 7,0000 |
| 116 | 0,2009 | 2,8330 | 0,6281 | | 0,0352 | 7,0000 |
| 117 | 0,2009 | 2,8555 | 0,6254 | | 0,0352 | 7,0000 |
| 118 | 0,2009 | 2,8779 | 0,6227 | | 0,0352 | 7,0000 |
| 119 | 0,2009 | 2,9004 | 0,6201 | | 0,0352 | 10,0000 |
| 120 | 0,2009 | 2,9229 | 0,6175 | | 0,0352 | 7,0000 |
| 121 | 0,2009 | 2,9453 | 0,6148 | | 0,0352 | 7,0000 |
| 122 | 0,2009 | 2,9678 | 0,6122 | | 0,0352 | 7,0000 |
| 123 | 0,2009 | 2,9902 | 0,6096 | | 0,0352 | 7,0000 |
| 124 | 0,2009 | 3,0127 | 0,6070 | | 0,0352 | 7,0000 |
| 125 | 0,2009 | 3,0352 | 0,6044 | | 0,0352 | 7,0000 |
| 126 | 0,2009 | 3,0576 | 0,6019 | | 0,0352 | 7,0000 |
| 127 | 0,2009 | 3,0801 | 0,5993 | | 0,0352 | 10,0000 |
| 128 | 0,2009 | 3,1025 | 0,5968 | | 0,0352 | 7,0000 |
| 129 | 0,2009 | 3,1250 | 0,5942 | | 0,0352 | 7,0000 |
| 130 | 0,2009 | 3,1475 | 0,5917 | | 0,0352 | 7,0000 |
| 131 | 0,2009 | 3,1699 | 0,5892 | | 0,0352 | 7,0000 |
| 132 | 0,2009 | 3,1924 | 0,5867 | | 0,0352 | 8,0000 |
| 133 | 0,2009 | 3,2148 | 0,5842 | | 0,0352 | 7,0000 |
| 134 | 0,2009 | 3,2373 | 0,5817 | | 0,0352 | 7,0000 |
| 135 | 0,2009 | 3,2598 | 0,5792 | | 0,0352 | 7,0000 |
| 136 | 0,2009 | 3,2822 | 0,5768 | | 0,0352 | 7,0000 |
| 137 | 0,2009 | 3,3047 | 0,5743 | | 0,0352 | 10,0000 |
| 138 | 0,2009 | 3,3271 | 0,5719 | | 0,0352 | 7,0000 |
| 139 | 0,2009 | 3,3496 | 0,5694 | | 0,0352 | 7,0000 |
| 140 | 0,2009 | 3,3721 | 0,5670 | | 0,0352 | 7,0000 |
| 141 | 0,2009 | 3,3945 | 0,5646 | | 0,0352 | 8,0000 |
| 142 | 0,2009 | 3,4170 | 0,5622 | | 0,0352 | 8,0000 |
| 143 | 0,2009 | 3,4395 | 0,5598 | | 0,0352 | 7,0000 |
| 144 | 0,2009 | 3,4619 | 0,5574 | | 0,0352 | 8,0000 |
| 145 | 0,2009 | 3,4844 | 0,5551 | | 0,0352 | 7,0000 |
| 146 | 0,2009 | 3,5068 | 0,5527 | | 0,0352 | 15,0000 |
| 147 | 0,2009 | 3,5293 | 0,5503 | | 0,0352 | 7,0000 |

| | 19-a-Final 1 | 20-b-Final 1 | 21-x column 2 | 22-y column 2 | 23 | 24-SumSq 2 |
|---|---|---|---|---|---|---|
| 99 | 1,0751 | 0,1897 | 2,4512 | 0,5056 | | 0,0664 |
| 100 | 1,0751 | 0,1897 | 2,4736 | 0,5021 | | 0,0664 |
| 101 | 1,0751 | 0,1897 | 2,4961 | 0,4987 | | 0,0664 |
| 102 | 1,0751 | 0,1897 | 2,5186 | 0,4953 | | 0,0664 |
| 103 | 1,0751 | 0,1897 | 2,5410 | 0,4919 | | 0,0664 |
| 104 | 1,0751 | 0,1897 | 2,5635 | 0,4885 | | 0,0664 |
| 105 | 1,0751 | 0,1897 | 2,5859 | 0,4852 | | 0,0664 |
| 106 | 1,0751 | 0,1897 | 2,6084 | 0,4819 | | 0,0664 |
| 107 | 1,0751 | 0,1897 | 2,6309 | 0,4786 | | 0,0664 |
| 108 | 1,0751 | 0,1897 | 2,6533 | 0,4753 | | 0,0664 |
| 109 | 1,0751 | 0,1897 | 2,6758 | 0,4721 | | 0,0664 |
| 110 | 1,0751 | 0,1897 | 2,6982 | 0,4688 | | 0,0664 |
| 111 | 1,0751 | 0,1897 | 2,7207 | 0,4656 | | 0,0664 |
| 112 | 1,0751 | 0,1897 | 2,7432 | 0,4625 | | 0,0664 |
| 113 | 1,0751 | 0,1897 | 2,7656 | 0,4593 | | 0,0664 |
| 114 | 1,0751 | 0,1897 | 2,7881 | 0,4562 | | 0,0664 |
| 115 | 1,0751 | 0,1897 | 2,8105 | 0,4530 | | 0,0664 |
| 116 | 1,0751 | 0,1897 | 2,8330 | 0,4500 | | 0,0664 |
| 117 | 1,0751 | 0,1897 | 2,8555 | 0,4469 | | 0,0664 |
| 118 | 1,0751 | 0,1897 | 2,8779 | 0,4438 | | 0,0664 |
| 119 | 1,0751 | 0,1897 | 2,9004 | 0,4408 | | 0,0664 |
| 120 | 1,0751 | 0,1897 | 2,9229 | 0,4378 | | 0,0664 |
| 121 | 1,0751 | 0,1897 | 2,9453 | 0,4348 | | 0,0664 |
| 122 | 1,0751 | 0,1897 | 2,9678 | 0,4318 | | 0,0664 |
| 123 | 1,0751 | 0,1897 | 2,9902 | 0,4289 | | 0,0664 |
| 124 | 1,0751 | 0,1897 | 3,0127 | 0,4259 | | 0,0664 |
| 125 | 1,0751 | 0,1897 | 3,0352 | 0,4230 | | 0,0664 |
| 126 | 1,0751 | 0,1897 | 3,0576 | 0,4201 | | 0,0664 |
| 127 | 1,0751 | 0,1897 | 3,0801 | 0,4173 | | 0,0664 |
| 128 | 1,0751 | 0,1897 | 3,1025 | 0,4144 | | 0,0664 |
| 129 | 1,0751 | 0,1897 | 3,1250 | 0,4116 | | 0,0664 |
| 130 | 1,0751 | 0,1897 | 3,1475 | 0,4088 | | 0,0664 |
| 131 | 1,0751 | 0,1897 | 3,1699 | 0,4060 | | 0,0664 |
| 132 | 1,0751 | 0,1897 | 3,1924 | 0,4032 | | 0,0664 |
| 133 | 1,0751 | 0,1897 | 3,2148 | 0,4005 | | 0,0664 |
| 134 | 1,0751 | 0,1897 | 3,2373 | 0,3977 | | 0,0664 |
| 135 | 1,0751 | 0,1897 | 3,2598 | 0,3950 | | 0,0664 |
| 136 | 1,0751 | 0,1897 | 3,2822 | 0,3923 | | 0,0664 |
| 137 | 1,0751 | 0,1897 | 3,3047 | 0,3896 | | 0,0664 |
| 138 | 1,0751 | 0,1897 | 3,3271 | 0,3870 | | 0,0664 |
| 139 | 1,0751 | 0,1897 | 3,3496 | 0,3843 | | 0,0664 |
| 140 | 1,0751 | 0,1897 | 3,3721 | 0,3817 | | 0,0664 |
| 141 | 1,0751 | 0,1897 | 3,3945 | 0,3791 | | 0,0664 |
| 142 | 1,0751 | 0,1897 | 3,4170 | 0,3765 | | 0,0664 |
| 143 | 1,0751 | 0,1897 | 3,4395 | 0,3739 | | 0,0664 |
| 144 | 1,0751 | 0,1897 | 3,4619 | 0,3714 | | 0,0664 |
| 145 | 1,0751 | 0,1897 | 3,4844 | 0,3688 | | 0,0664 |
| 146 | 1,0751 | 0,1897 | 3,5068 | 0,3663 | | 0,0664 |
| 147 | 1,0751 | 0,1897 | 3,5293 | 0,3638 | | 0,0664 |

| | 25-Iterations 2 | 26-a-Final 2 | 27-b-Final 2 | 28-x column 3 | 29-y column 3 | 30 |
|---|---|---|---|---|---|---|
| 99 | 10,0000 | 1,0684 | 0,3052 | 2,4512 | 0,3916 | |
| 100 | 9,0000 | 1,0684 | 0,3052 | 2,4736 | 0,3882 | |
| 101 | 10,0000 | 1,0684 | 0,3052 | 2,4961 | 0,3848 | |
| 102 | 10,0000 | 1,0684 | 0,3052 | 2,5186 | 0,3815 | |
| 103 | 11,0000 | 1,0684 | 0,3052 | 2,5410 | 0,3782 | |
| 104 | 9,0000 | 1,0684 | 0,3052 | 2,5635 | 0,3750 | |
| 105 | 10,0000 | 1,0684 | 0,3052 | 2,5859 | 0,3718 | |
| 106 | 10,0000 | 1,0684 | 0,3052 | 2,6084 | 0,3685 | |
| 107 | 9,0000 | 1,0684 | 0,3052 | 2,6309 | 0,3654 | |
| 108 | 11,0000 | 1,0684 | 0,3052 | 2,6533 | 0,3622 | |
| 109 | 10,0000 | 1,0684 | 0,3052 | 2,6758 | 0,3591 | |
| 110 | 10,0000 | 1,0684 | 0,3052 | 2,6982 | 0,3560 | |
| 111 | 10,0000 | 1,0684 | 0,3052 | 2,7207 | 0,3529 | |
| 112 | 10,0000 | 1,0684 | 0,3052 | 2,7432 | 0,3499 | |
| 113 | 10,0000 | 1,0684 | 0,3052 | 2,7656 | 0,3469 | |
| 114 | 10,0000 | 1,0684 | 0,3052 | 2,7881 | 0,3439 | |
| 115 | 15,0000 | 1,0684 | 0,3052 | 2,8105 | 0,3409 | |
| 116 | 9,0000 | 1,0684 | 0,3052 | 2,8330 | 0,3380 | |
| 117 | 11,0000 | 1,0684 | 0,3052 | 2,8555 | 0,3351 | |
| 118 | 10,0000 | 1,0684 | 0,3052 | 2,8779 | 0,3322 | |
| 119 | 10,0000 | 1,0684 | 0,3052 | 2,9004 | 0,3293 | |
| 120 | 10,0000 | 1,0684 | 0,3052 | 2,9229 | 0,3265 | |
| 121 | 11,0000 | 1,0684 | 0,3052 | 2,9453 | 0,3237 | |
| 122 | 10,0000 | 1,0684 | 0,3052 | 2,9678 | 0,3209 | |
| 123 | 10,0000 | 1,0684 | 0,3052 | 2,9902 | 0,3181 | |
| 124 | 10,0000 | 1,0684 | 0,3052 | 3,0127 | 0,3154 | |
| 125 | 11,0000 | 1,0684 | 0,3052 | 3,0352 | 0,3127 | |
| 126 | 12,0000 | 1,0684 | 0,3052 | 3,0576 | 0,3100 | |
| 127 | 10,0000 | 1,0684 | 0,3052 | 3,0801 | 0,3073 | |
| 128 | 11,0000 | 1,0684 | 0,3052 | 3,1025 | 0,3046 | |
| 129 | 10,0000 | 1,0684 | 0,3052 | 3,1250 | 0,3020 | |
| 130 | 14,0000 | 1,0684 | 0,3052 | 3,1475 | 0,2994 | |
| 131 | 11,0000 | 1,0684 | 0,3052 | 3,1699 | 0,2968 | |
| 132 | 10,0000 | 1,0684 | 0,3052 | 3,1924 | 0,2943 | |
| 133 | 10,0000 | 1,0684 | 0,3052 | 3,2148 | 0,2917 | |
| 134 | 11,0000 | 1,0684 | 0,3052 | 3,2373 | 0,2892 | |
| 135 | 10,0000 | 1,0684 | 0,3052 | 3,2598 | 0,2867 | |
| 136 | 10,0000 | 1,0684 | 0,3052 | 3,2822 | 0,2843 | |
| 137 | 10,0000 | 1,0684 | 0,3052 | 3,3047 | 0,2818 | |
| 138 | 11,0000 | 1,0684 | 0,3052 | 3,3271 | 0,2794 | |
| 139 | 11,0000 | 1,0684 | 0,3052 | 3,3496 | 0,2770 | |
| 140 | 11,0000 | 1,0684 | 0,3052 | 3,3721 | 0,2746 | |
| 141 | 10,0000 | 1,0684 | 0,3052 | 3,3945 | 0,2722 | |
| 142 | 49,0000 | 1,0684 | 0,3052 | 3,4170 | 0,2699 | |
| 143 | 10,0000 | 1,0684 | 0,3052 | 3,4395 | 0,2676 | |
| 144 | 11,0000 | 1,0684 | 0,3052 | 3,4619 | 0,2652 | |
| 145 | 11,0000 | 1,0684 | 0,3052 | 3,4844 | 0,2630 | |
| 146 | 9,0000 | 1,0684 | 0,3052 | 3,5068 | 0,2607 | |
| 147 | 11,0000 | 1,0684 | 0,3052 | 3,5293 | 0,2584 | |

| | 31-SumSq 3 | 32-Iterations 3 | 33-a-Final 3 | 34-b-Final 3 | 35-x column 4 | 36-y column 4 |
|---|---|---|---|---|---|---|
| 99 | 0,0429 | 12,0000 | 1,0070 | 0,3854 | 2,4512 | 0,3916 |
| 100 | 0,0429 | 11,0000 | 1,0070 | 0,3854 | 2,4736 | 0,3882 |
| 101 | 0,0429 | 11,0000 | 1,0070 | 0,3854 | 2,4961 | 0,3848 |
| 102 | 0,0429 | 16,0000 | 1,0070 | 0,3854 | 2,5186 | 0,3815 |
| 103 | 0,0429 | 11,0000 | 1,0070 | 0,3854 | 2,5410 | 0,3782 |
| 104 | 0,0429 | 11,0000 | 1,0070 | 0,3854 | 2,5635 | 0,3750 |
| 105 | 0,0429 | 12,0000 | 1,0070 | 0,3854 | 2,5859 | 0,3718 |
| 106 | 0,0429 | 12,0000 | 1,0070 | 0,3854 | 2,6084 | 0,3685 |
| 107 | 0,0429 | 11,0000 | 1,0070 | 0,3854 | 2,6309 | 0,3654 |
| 108 | 0,0429 | 11,0000 | 1,0070 | 0,3854 | 2,6533 | 0,3622 |
| 109 | 0,0429 | 11,0000 | 1,0070 | 0,3854 | 2,6758 | 0,3591 |
| 110 | 0,0429 | 11,0000 | 1,0070 | 0,3854 | 2,6982 | 0,3560 |
| 111 | 0,0429 | 11,0000 | 1,0070 | 0,3854 | 2,7207 | 0,3529 |
| 112 | 0,0429 | 12,0000 | 1,0070 | 0,3854 | 2,7432 | 0,3499 |
| 113 | 0,0429 | 16,0000 | 1,0070 | 0,3854 | 2,7656 | 0,3469 |
| 114 | 0,0429 | 12,0000 | 1,0070 | 0,3854 | 2,7881 | 0,3439 |
| 115 | 0,0429 | 11,0000 | 1,0070 | 0,3854 | 2,8105 | 0,3409 |
| 116 | 0,0429 | 11,0000 | 1,0070 | 0,3854 | 2,8330 | 0,3380 |
| 117 | 0,0429 | 11,0000 | 1,0070 | 0,3854 | 2,8555 | 0,3351 |
| 118 | 0,0429 | 12,0000 | 1,0070 | 0,3854 | 2,8779 | 0,3322 |
| 119 | 0,0429 | 12,0000 | 1,0070 | 0,3854 | 2,9004 | 0,3293 |
| 120 | 0,0429 | 11,0000 | 1,0070 | 0,3854 | 2,9229 | 0,3265 |
| 121 | 0,0429 | 23,0000 | 1,0070 | 0,3854 | 2,9453 | 0,3237 |
| 122 | 0,0429 | 11,0000 | 1,0070 | 0,3854 | 2,9678 | 0,3209 |
| 123 | 0,0429 | 12,0000 | 1,0070 | 0,3854 | 2,9902 | 0,3181 |
| 124 | 0,0429 | 12,0000 | 1,0070 | 0,3854 | 3,0127 | 0,3154 |
| 125 | 0,0429 | 10,0000 | 1,0070 | 0,3854 | 3,0352 | 0,3127 |
| 126 | 0,0429 | 11,0000 | 1,0070 | 0,3854 | 3,0576 | 0,3100 |
| 127 | 0,0429 | 11,0000 | 1,0070 | 0,3854 | 3,0801 | 0,3073 |
| 128 | 0,0429 | 11,0000 | 1,0070 | 0,3854 | 3,1025 | 0,3046 |
| 129 | 0,0429 | 16,0000 | 1,0070 | 0,3854 | 3,1250 | 0,3020 |
| 130 | 0,0429 | 11,0000 | 1,0070 | 0,3854 | 3,1475 | 0,2994 |
| 131 | 0,0429 | 13,0000 | 1,0070 | 0,3854 | 3,1699 | 0,2968 |
| 132 | 0,0429 | 16,0000 | 1,0070 | 0,3854 | 3,1924 | 0,2943 |
| 133 | 0,0429 | 11,0000 | 1,0070 | 0,3854 | 3,2148 | 0,2917 |
| 134 | 0,0429 | 11,0000 | 1,0070 | 0,3854 | 3,2373 | 0,2892 |
| 135 | 0,0429 | 11,0000 | 1,0070 | 0,3854 | 3,2598 | 0,2867 |
| 136 | 0,0429 | 9,0000 | 1,0070 | 0,3854 | 3,2822 | 0,2843 |
| 137 | 0,0429 | 12,0000 | 1,0070 | 0,3854 | 3,3047 | 0,2818 |
| 138 | 0,0429 | 11,0000 | 1,0070 | 0,3854 | 3,3271 | 0,2794 |
| 139 | 0,0429 | 11,0000 | 1,0070 | 0,3854 | 3,3496 | 0,2770 |
| 140 | 0,0429 | 11,0000 | 1,0070 | 0,3854 | 3,3721 | 0,2746 |
| 141 | 0,0429 | 11,0000 | 1,0070 | 0,3854 | 3,3945 | 0,2722 |
| 142 | 0,0429 | 11,0000 | 1,0070 | 0,3854 | 3,4170 | 0,2699 |
| 143 | 0,0429 | 11,0000 | 1,0070 | 0,3854 | 3,4395 | 0,2676 |
| 144 | 0,0429 | 11,0000 | 1,0070 | 0,3854 | 3,4619 | 0,2652 |
| 145 | 0,0429 | 11,0000 | 1,0070 | 0,3854 | 3,4844 | 0,2630 |
| 146 | 0,0429 | 11,0000 | 1,0070 | 0,3854 | 3,5068 | 0,2607 |
| 147 | 0,0429 | 12,0000 | 1,0070 | 0,3854 | 3,5293 | 0,2584 |

| | 37 | 38-SumSq 4 | 39-Iterations 4 | 40-a-Final 4 | 41-b-Final 4 | 42-x column 5 |
|---|---|---|---|---|---|---|
| 99 | | 0,0429 | 10,0000 | 1,0070 | 0,3854 | 2,4512 |
| 100 | | 0,0429 | 10,0000 | 1,0070 | 0,3854 | 2,4736 |
| 101 | | 0,0429 | 10,0000 | 1,0070 | 0,3854 | 2,4961 |
| 102 | | 0,0429 | 10,0000 | 1,0070 | 0,3854 | 2,5186 |
| 103 | | 0,0429 | 10,0000 | 1,0070 | 0,3854 | 2,5410 |
| 104 | | 0,0429 | 10,0000 | 1,0070 | 0,3854 | 2,5635 |
| 105 | | 0,0429 | 10,0000 | 1,0070 | 0,3854 | 2,5859 |
| 106 | | 0,0429 | 10,0000 | 1,0070 | 0,3854 | 2,6084 |
| 107 | | 0,0429 | 10,0000 | 1,0070 | 0,3854 | 2,6309 |
| 108 | | 0,0429 | 10,0000 | 1,0070 | 0,3854 | 2,6533 |
| 109 | | 0,0429 | 10,0000 | 1,0070 | 0,3854 | 2,6758 |
| 110 | | 0,0429 | 10,0000 | 1,0070 | 0,3854 | 2,6982 |
| 111 | | 0,0429 | 10,0000 | 1,0070 | 0,3854 | 2,7207 |
| 112 | | 0,0429 | 10,0000 | 1,0070 | 0,3854 | 2,7432 |
| 113 | | 0,0429 | 9,0000 | 1,0070 | 0,3854 | 2,7656 |
| 114 | | 0,0429 | 10,0000 | 1,0070 | 0,3854 | 2,7881 |
| 115 | | 0,0429 | 10,0000 | 1,0070 | 0,3854 | 2,8105 |
| 116 | | 0,0429 | 10,0000 | 1,0070 | 0,3854 | 2,8330 |
| 117 | | 0,0429 | 10,0000 | 1,0070 | 0,3854 | 2,8555 |
| 118 | | 0,0429 | 10,0000 | 1,0070 | 0,3854 | 2,8779 |
| 119 | | 0,0429 | 11,0000 | 1,0070 | 0,3854 | 2,9004 |
| 120 | | 0,0429 | 10,0000 | 1,0070 | 0,3854 | 2,9229 |
| 121 | | 0,0429 | 10,0000 | 1,0070 | 0,3854 | 2,9453 |
| 122 | | 0,0429 | 10,0000 | 1,0070 | 0,3854 | 2,9678 |
| 123 | | 0,0429 | 10,0000 | 1,0070 | 0,3854 | 2,9902 |
| 124 | | 0,0429 | 9,0000 | 1,0070 | 0,3854 | 3,0127 |
| 125 | | 0,0429 | 10,0000 | 1,0070 | 0,3854 | 3,0352 |
| 126 | | 0,0429 | 10,0000 | 1,0070 | 0,3854 | 3,0576 |
| 127 | | 0,0429 | 10,0000 | 1,0070 | 0,3854 | 3,0801 |
| 128 | | 0,0429 | 10,0000 | 1,0070 | 0,3854 | 3,1025 |
| 129 | | 0,0429 | 10,0000 | 1,0070 | 0,3854 | 3,1250 |
| 130 | | 0,0429 | 10,0000 | 1,0070 | 0,3854 | 3,1475 |
| 131 | | 0,0429 | 10,0000 | 1,0070 | 0,3854 | 3,1699 |
| 132 | | 0,0429 | 10,0000 | 1,0070 | 0,3854 | 3,1924 |
| 133 | | 0,0429 | 10,0000 | 1,0070 | 0,3854 | 3,2148 |
| 134 | | 0,0429 | 10,0000 | 1,0070 | 0,3854 | 3,2373 |
| 135 | | 0,0429 | 10,0000 | 1,0070 | 0,3854 | 3,2598 |
| 136 | | 0,0429 | 10,0000 | 1,0070 | 0,3854 | 3,2822 |
| 137 | | 0,0429 | 10,0000 | 1,0070 | 0,3854 | 3,3047 |
| 138 | | 0,0429 | 10,0000 | 1,0070 | 0,3854 | 3,3271 |
| 139 | | 0,0429 | 10,0000 | 1,0070 | 0,3854 | 3,3496 |
| 140 | | 0,0429 | 10,0000 | 1,0070 | 0,3854 | 3,3721 |
| 141 | | 0,0429 | 10,0000 | 1,0070 | 0,3854 | 3,3945 |
| 142 | | 0,0429 | 11,0000 | 1,0070 | 0,3854 | 3,4170 |
| 143 | | 0,0429 | 10,0000 | 1,0070 | 0,3854 | 3,4395 |
| 144 | | 0,0429 | 9,0000 | 1,0070 | 0,3854 | 3,4619 |
| 145 | | 0,0429 | 10,0000 | 1,0070 | 0,3854 | 3,4844 |
| 146 | | 0,0429 | 10,0000 | 1,0070 | 0,3854 | 3,5068 |
| 147 | | 0,0429 | 10,0000 | 1,0070 | 0,3854 | 3,5293 |

| | 43-y column 5 | 44 | 45-SumSq 5 | 46-Iterations 5 | 47-a-Final 5 | 48-b-Final 5 |
|---|---|---|---|---|---|---|
| 99 | 0,5588 | | 0,1194 | 9,0000 | 1,0036 | 0,2389 |
| 100 | 0,5558 | | 0,1194 | 10,0000 | 1,0036 | 0,2389 |
| 101 | 0,5528 | | 0,1194 | 9,0000 | 1,0036 | 0,2389 |
| 102 | 0,5499 | | 0,1194 | 9,0000 | 1,0036 | 0,2389 |
| 103 | 0,5469 | | 0,1194 | 9,0000 | 1,0036 | 0,2389 |
| 104 | 0,5440 | | 0,1194 | 9,0000 | 1,0036 | 0,2389 |
| 105 | 0,5411 | | 0,1194 | 10,0000 | 1,0036 | 0,2389 |
| 106 | 0,5382 | | 0,1194 | 8,0000 | 1,0036 | 0,2389 |
| 107 | 0,5353 | | 0,1194 | 10,0000 | 1,0036 | 0,2389 |
| 108 | 0,5325 | | 0,1194 | 9,0000 | 1,0036 | 0,2389 |
| 109 | 0,5296 | | 0,1194 | 8,0000 | 1,0036 | 0,2389 |
| 110 | 0,5268 | | 0,1194 | 9,0000 | 1,0036 | 0,2389 |
| 111 | 0,5240 | | 0,1194 | 9,0000 | 1,0036 | 0,2389 |
| 112 | 0,5212 | | 0,1194 | 9,0000 | 1,0036 | 0,2389 |
| 113 | 0,5184 | | 0,1194 | 8,0000 | 1,0036 | 0,2389 |
| 114 | 0,5156 | | 0,1194 | 9,0000 | 1,0036 | 0,2389 |
| 115 | 0,5128 | | 0,1194 | 10,0000 | 1,0036 | 0,2389 |
| 116 | 0,5101 | | 0,1194 | 10,0000 | 1,0036 | 0,2389 |
| 117 | 0,5074 | | 0,1194 | 9,0000 | 1,0036 | 0,2389 |
| 118 | 0,5047 | | 0,1194 | 9,0000 | 1,0036 | 0,2389 |
| 119 | 0,5020 | | 0,1194 | 9,0000 | 1,0036 | 0,2389 |
| 120 | 0,4993 | | 0,1194 | 7,0000 | 1,0036 | 0,2389 |
| 121 | 0,4966 | | 0,1194 | 9,0000 | 1,0036 | 0,2389 |
| 122 | 0,4939 | | 0,1194 | 10,0000 | 1,0036 | 0,2389 |
| 123 | 0,4913 | | 0,1194 | 9,0000 | 1,0036 | 0,2389 |
| 124 | 0,4887 | | 0,1194 | 10,0000 | 1,0036 | 0,2389 |
| 125 | 0,4861 | | 0,1194 | 10,0000 | 1,0036 | 0,2389 |
| 126 | 0,4834 | | 0,1194 | 9,0000 | 1,0036 | 0,2389 |
| 127 | 0,4809 | | 0,1194 | 9,0000 | 1,0036 | 0,2389 |
| 128 | 0,4783 | | 0,1194 | 8,0000 | 1,0036 | 0,2389 |
| 129 | 0,4757 | | 0,1194 | 9,0000 | 1,0036 | 0,2389 |
| 130 | 0,4732 | | 0,1194 | 10,0000 | 1,0036 | 0,2389 |
| 131 | 0,4707 | | 0,1194 | 9,0000 | 1,0036 | 0,2389 |
| 132 | 0,4681 | | 0,1194 | 10,0000 | 1,0036 | 0,2389 |
| 133 | 0,4656 | | 0,1194 | 9,0000 | 1,0036 | 0,2389 |
| 134 | 0,4631 | | 0,1194 | 7,0000 | 1,0036 | 0,2389 |
| 135 | 0,4607 | | 0,1194 | 9,0000 | 1,0036 | 0,2389 |
| 136 | 0,4582 | | 0,1194 | 9,0000 | 1,0036 | 0,2389 |
| 137 | 0,4557 | | 0,1194 | 9,0000 | 1,0036 | 0,2389 |
| 138 | 0,4533 | | 0,1194 | 9,0000 | 1,0036 | 0,2389 |
| 139 | 0,4509 | | 0,1194 | 10,0000 | 1,0036 | 0,2389 |
| 140 | 0,4485 | | 0,1194 | 8,0000 | 1,0036 | 0,2389 |
| 141 | 0,4461 | | 0,1194 | 9,0000 | 1,0036 | 0,2389 |
| 142 | 0,4437 | | 0,1194 | 9,0000 | 1,0036 | 0,2389 |
| 143 | 0,4413 | | 0,1194 | 9,0000 | 1,0036 | 0,2389 |
| 144 | 0,4389 | | 0,1194 | 9,0000 | 1,0036 | 0,2389 |
| 145 | 0,4366 | | 0,1194 | 9,0000 | 1,0036 | 0,2389 |
| 146 | 0,4343 | | 0,1194 | 9,0000 | 1,0036 | 0,2389 |
| 147 | 0,4319 | | 0,1194 | 9,0000 | 1,0036 | 0,2389 |

| | 49-x column 6 | 50-y column 6 | 51 | 52-SumSq 6 | 53-Iterations 6 | 54-a-Final 6 |
|---|---|---|---|---|---|---|
| 99 | 2,4512 | 0,4444 | | 0,0865 | 10,0000 | 0,9419 |
| 100 | 2,4736 | 0,4414 | | 0,0865 | 10,0000 | 0,9419 |
| 101 | 2,4961 | 0,4384 | | 0,0865 | 10,0000 | 0,9419 |
| 102 | 2,5186 | 0,4354 | | 0,0865 | 10,0000 | 0,9419 |
| 103 | 2,5410 | 0,4324 | | 0,0865 | 11,0000 | 0,9419 |
| 104 | 2,5635 | 0,4294 | | 0,0865 | 10,0000 | 0,9419 |
| 105 | 2,5859 | 0,4265 | | 0,0865 | 10,0000 | 0,9419 |
| 106 | 2,6084 | 0,4235 | | 0,0865 | 10,0000 | 0,9419 |
| 107 | 2,6309 | 0,4206 | | 0,0865 | 10,0000 | 0,9419 |
| 108 | 2,6533 | 0,4177 | | 0,0865 | 10,0000 | 0,9419 |
| 109 | 2,6758 | 0,4149 | | 0,0865 | 9,0000 | 0,9419 |
| 110 | 2,6982 | 0,4120 | | 0,0865 | 10,0000 | 0,9419 |
| 111 | 2,7207 | 0,4092 | | 0,0865 | 11,0000 | 0,9419 |
| 112 | 2,7432 | 0,4064 | | 0,0865 | 9,0000 | 0,9419 |
| 113 | 2,7656 | 0,4036 | | 0,0865 | 10,0000 | 0,9419 |
| 114 | 2,7881 | 0,4008 | | 0,0865 | 10,0000 | 0,9419 |
| 115 | 2,8105 | 0,3981 | | 0,0865 | 10,0000 | 0,9419 |
| 116 | 2,8330 | 0,3954 | | 0,0865 | 11,0000 | 0,9419 |
| 117 | 2,8555 | 0,3926 | | 0,0865 | 10,0000 | 0,9419 |
| 118 | 2,8779 | 0,3900 | | 0,0865 | 10,0000 | 0,9419 |
| 119 | 2,9004 | 0,3873 | | 0,0865 | 10,0000 | 0,9419 |
| 120 | 2,9229 | 0,3846 | | 0,0865 | 10,0000 | 0,9419 |
| 121 | 2,9453 | 0,3820 | | 0,0865 | 10,0000 | 0,9419 |
| 122 | 2,9678 | 0,3794 | | 0,0865 | 11,0000 | 0,9419 |
| 123 | 2,9902 | 0,3768 | | 0,0865 | 10,0000 | 0,9419 |
| 124 | 3,0127 | 0,3742 | | 0,0865 | 10,0000 | 0,9419 |
| 125 | 3,0352 | 0,3716 | | 0,0865 | 10,0000 | 0,9419 |
| 126 | 3,0576 | 0,3691 | | 0,0865 | 10,0000 | 0,9419 |
| 127 | 3,0801 | 0,3665 | | 0,0865 | 10,0000 | 0,9419 |
| 128 | 3,1025 | 0,3640 | | 0,0865 | 11,0000 | 0,9419 |
| 129 | 3,1250 | 0,3615 | | 0,0865 | 10,0000 | 0,9419 |
| 130 | 3,1475 | 0,3590 | | 0,0865 | 10,0000 | 0,9419 |
| 131 | 3,1699 | 0,3566 | | 0,0865 | 10,0000 | 0,9419 |
| 132 | 3,1924 | 0,3541 | | 0,0865 | 10,0000 | 0,9419 |
| 133 | 3,2148 | 0,3517 | | 0,0865 | 10,0000 | 0,9419 |
| 134 | 3,2373 | 0,3493 | | 0,0865 | 10,0000 | 0,9419 |
| 135 | 3,2598 | 0,3469 | | 0,0865 | 10,0000 | 0,9419 |
| 136 | 3,2822 | 0,3445 | | 0,0865 | 10,0000 | 0,9419 |
| 137 | 3,3047 | 0,3422 | | 0,0865 | 10,0000 | 0,9419 |
| 138 | 3,3271 | 0,3398 | | 0,0865 | 10,0000 | 0,9419 |
| 139 | 3,3496 | 0,3375 | | 0,0865 | 10,0000 | 0,9419 |
| 140 | 3,3721 | 0,3352 | | 0,0865 | 10,0000 | 0,9419 |
| 141 | 3,3945 | 0,3329 | | 0,0865 | 10,0000 | 0,9419 |
| 142 | 3,4170 | 0,3306 | | 0,0865 | 10,0000 | 0,9419 |
| 143 | 3,4395 | 0,3283 | | 0,0865 | 10,0000 | 0,9419 |
| 144 | 3,4619 | 0,3261 | | 0,0865 | 10,0000 | 0,9419 |
| 145 | 3,4844 | 0,3238 | | 0,0865 | 10,0000 | 0,9419 |
| 146 | 3,5068 | 0,3216 | | 0,0865 | 10,0000 | 0,9419 |
| 147 | 3,5293 | 0,3194 | | 0,0865 | 10,0000 | 0,9419 |

| | 55-b-Final 6 |
|---|---|
| 99 | 0,3064 |
| 100 | 0,3064 |
| 101 | 0,3064 |
| 102 | 0,3064 |
| 103 | 0,3064 |
| 104 | 0,3064 |
| 105 | 0,3064 |
| 106 | 0,3064 |
| 107 | 0,3064 |
| 108 | 0,3064 |
| 109 | 0,3064 |
| 110 | 0,3064 |
| 111 | 0,3064 |
| 112 | 0,3064 |
| 113 | 0,3064 |
| 114 | 0,3064 |
| 115 | 0,3064 |
| 116 | 0,3064 |
| 117 | 0,3064 |
| 118 | 0,3064 |
| 119 | 0,3064 |
| 120 | 0,3064 |
| 121 | 0,3064 |
| 122 | 0,3064 |
| 123 | 0,3064 |
| 124 | 0,3064 |
| 125 | 0,3064 |
| 126 | 0,3064 |
| 127 | 0,3064 |
| 128 | 0,3064 |
| 129 | 0,3064 |
| 130 | 0,3064 |
| 131 | 0,3064 |
| 132 | 0,3064 |
| 133 | 0,3064 |
| 134 | 0,3064 |
| 135 | 0,3064 |
| 136 | 0,3064 |
| 137 | 0,3064 |
| 138 | 0,3064 |
| 139 | 0,3064 |
| 140 | 0,3064 |
| 141 | 0,3064 |
| 142 | 0,3064 |
| 143 | 0,3064 |
| 144 | 0,3064 |
| 145 | 0,3064 |
| 146 | 0,3064 |
| 147 | 0,3064 |

| | 1 | 2 | 3 | 4 | 5 | 6 |
|---|---|---|---|---|---|---|
| 148 | | | | | | |
| 149 | | | | | | |
| 150 | | | | | | |
| 151 | | | | | | |
| 152 | | | | | | |
| 153 | | | | | | |
| 154 | | | | | | |
| 155 | | | | | | |
| 156 | | | | | | |
| 157 | | | | | | |
| 158 | | | | | | |
| 159 | | | | | | |
| 160 | | | | | | |
| 161 | | | | | | |
| 162 | | | | | | |
| 163 | | | | | | |
| 164 | | | | | | |
| 165 | | | | | | |
| 166 | | | | | | |
| 167 | | | | | | |
| 168 | | | | | | |
| 169 | | | | | | |
| 170 | | | | | | |
| 171 | | | | | | |
| 172 | | | | | | |
| 173 | | | | | | |
| 174 | | | | | | |
| 175 | | | | | | |
| 176 | | | | | | |
| 177 | | | | | | |
| 178 | | | | | | |
| 179 | | | | | | |
| 180 | | | | | | |
| 181 | | | | | | |
| 182 | | | | | | |
| 183 | | | | | | |
| 184 | | | | | | |
| 185 | | | | | | |
| 186 | | | | | | |
| 187 | | | | | | |
| 188 | | | | | | |
| 189 | | | | | | |
| 190 | | | | | | |
| 191 | | | | | | |
| 192 | | | | | | |
| 193 | | | | | | |
| 194 | | | | | | |
| 195 | | | | | | |
| 196 | | | | | | |

| | 7-x column | 8-y column | 9 | 10-SumSq | 11-Iterations | 12-a-Final |
|---|---|---|---|---|---|---|
| 148 | 3,5518 | 0,5375 | | 0,0411 | 8,0000 | 1,0972 |
| 149 | 3,5742 | 0,5351 | | 0,0411 | 8,0000 | 1,0972 |
| 150 | 3,5967 | 0,5327 | | 0,0411 | 7,0000 | 1,0972 |
| 151 | 3,6191 | 0,5303 | | 0,0411 | 8,0000 | 1,0972 |
| 152 | 3,6416 | 0,5279 | | 0,0411 | 7,0000 | 1,0972 |
| 153 | 3,6641 | 0,5255 | | 0,0411 | 7,0000 | 1,0972 |
| 154 | 3,6865 | 0,5232 | | 0,0411 | 10,0000 | 1,0972 |
| 155 | 3,7090 | 0,5208 | | 0,0411 | 7,0000 | 1,0972 |
| 156 | 3,7314 | 0,5185 | | 0,0411 | 7,0000 | 1,0972 |
| 157 | 3,7539 | 0,5161 | | 0,0411 | 7,0000 | 1,0972 |
| 158 | 3,7764 | 0,5138 | | 0,0411 | 7,0000 | 1,0972 |
| 159 | 3,7988 | 0,5115 | | 0,0411 | 7,0000 | 1,0972 |
| 160 | 3,8213 | 0,5092 | | 0,0411 | 7,0000 | 1,0972 |
| 161 | 3,8438 | 0,5069 | | 0,0411 | 7,0000 | 1,0972 |
| 162 | 3,8662 | 0,5046 | | 0,0411 | 7,0000 | 1,0972 |
| 163 | 3,8887 | 0,5023 | | 0,0411 | 7,0000 | 1,0972 |
| 164 | 3,9111 | 0,5001 | | 0,0411 | 7,0000 | 1,0972 |
| 165 | 3,9336 | 0,4978 | | 0,0411 | 7,0000 | 1,0972 |
| 166 | 3,9561 | 0,4956 | | 0,0411 | 7,0000 | 1,0972 |
| 167 | 3,9785 | 0,4934 | | 0,0411 | 7,0000 | 1,0972 |
| 168 | 4,0010 | 0,4911 | | 0,0411 | 7,0000 | 1,0972 |
| 169 | 4,0234 | 0,4889 | | 0,0411 | 7,0000 | 1,0972 |
| 170 | 4,0459 | 0,4867 | | 0,0411 | 7,0000 | 1,0972 |
| 171 | 4,0684 | 0,4845 | | 0,0411 | 7,0000 | 1,0972 |
| 172 | 4,0908 | 0,4824 | | 0,0411 | 7,0000 | 1,0972 |
| 173 | 4,1133 | 0,4802 | | 0,0411 | 7,0000 | 1,0972 |
| 174 | 4,1357 | 0,4780 | | 0,0411 | 7,0000 | 1,0972 |
| 175 | 4,1582 | 0,4759 | | 0,0411 | 7,0000 | 1,0972 |
| 176 | 4,1807 | 0,4737 | | 0,0411 | 7,0000 | 1,0972 |
| 177 | 4,2031 | 0,4716 | | 0,0411 | 7,0000 | 1,0972 |
| 178 | 4,2256 | 0,4695 | | 0,0411 | 7,0000 | 1,0972 |
| 179 | 4,2480 | 0,4674 | | 0,0411 | 7,0000 | 1,0972 |
| 180 | 4,2705 | 0,4653 | | 0,0411 | 7,0000 | 1,0972 |
| 181 | 4,2930 | 0,4632 | | 0,0411 | 7,0000 | 1,0972 |
| 182 | 4,3154 | 0,4611 | | 0,0411 | 7,0000 | 1,0972 |
| 183 | 4,3379 | 0,4590 | | 0,0411 | 7,0000 | 1,0972 |
| 184 | 4,3604 | 0,4569 | | 0,0411 | 7,0000 | 1,0972 |
| 185 | 4,3828 | 0,4549 | | 0,0411 | 7,0000 | 1,0972 |
| 186 | 4,4053 | 0,4528 | | 0,0411 | 7,0000 | 1,0972 |
| 187 | 4,4277 | 0,4508 | | 0,0411 | 7,0000 | 1,0972 |
| 188 | 4,4502 | 0,4488 | | 0,0411 | 7,0000 | 1,0972 |
| 189 | 4,4727 | 0,4467 | | 0,0411 | 7,0000 | 1,0972 |
| 190 | 4,4951 | 0,4447 | | 0,0411 | 7,0000 | 1,0972 |
| 191 | 4,5176 | 0,4427 | | 0,0411 | 7,0000 | 1,0972 |
| 192 | 4,5400 | 0,4407 | | 0,0411 | 7,0000 | 1,0972 |
| 193 | 4,5625 | 0,4387 | | 0,0411 | 7,0000 | 1,0972 |
| 194 | 4,5850 | 0,4368 | | 0,0411 | 7,0000 | 1,0972 |
| 195 | 4,6074 | 0,4348 | | 0,0411 | 7,0000 | 1,0972 |
| 196 | 4,6299 | 0,4328 | | 0,0411 | 7,0000 | 1,0972 |

| | 13-b-Final | 14-x column 1 | 15-y column 1 | 16 | 17-SumSq 1 | 18-Iterations 1 |
|---|---|---|---|---|---|---|
| 148 | 0,2009 | 3,5518 | 0,5480 | | 0,0352 | 7,0000 |
| 149 | 0,2009 | 3,5742 | 0,5457 | | 0,0352 | 12,0000 |
| 150 | 0,2009 | 3,5967 | 0,5434 | | 0,0352 | 33,0000 |
| 151 | 0,2009 | 3,6191 | 0,5410 | | 0,0352 | 7,0000 |
| 152 | 0,2009 | 3,6416 | 0,5387 | | 0,0352 | 7,0000 |
| 153 | 0,2009 | 3,6641 | 0,5364 | | 0,0352 | 7,0000 |
| 154 | 0,2009 | 3,6865 | 0,5342 | | 0,0352 | 10,0000 |
| 155 | 0,2009 | 3,7090 | 0,5319 | | 0,0352 | 7,0000 |
| 156 | 0,2009 | 3,7314 | 0,5296 | | 0,0352 | 7,0000 |
| 157 | 0,2009 | 3,7539 | 0,5274 | | 0,0352 | 7,0000 |
| 158 | 0,2009 | 3,7764 | 0,5251 | | 0,0352 | 7,0000 |
| 159 | 0,2009 | 3,7988 | 0,5229 | | 0,0352 | 7,0000 |
| 160 | 0,2009 | 3,8213 | 0,5207 | | 0,0352 | 7,0000 |
| 161 | 0,2009 | 3,8438 | 0,5185 | | 0,0352 | 7,0000 |
| 162 | 0,2009 | 3,8662 | 0,5163 | | 0,0352 | 10,0000 |
| 163 | 0,2009 | 3,8887 | 0,5141 | | 0,0352 | 7,0000 |
| 164 | 0,2009 | 3,9111 | 0,5119 | | 0,0352 | 7,0000 |
| 165 | 0,2009 | 3,9336 | 0,5097 | | 0,0352 | 7,0000 |
| 166 | 0,2009 | 3,9561 | 0,5075 | | 0,0352 | 7,0000 |
| 167 | 0,2009 | 3,9785 | 0,5054 | | 0,0352 | 7,0000 |
| 168 | 0,2009 | 4,0010 | 0,5032 | | 0,0352 | 7,0000 |
| 169 | 0,2009 | 4,0234 | 0,5011 | | 0,0352 | 7,0000 |
| 170 | 0,2009 | 4,0459 | 0,4990 | | 0,0352 | 7,0000 |
| 171 | 0,2009 | 4,0684 | 0,4968 | | 0,0352 | 7,0000 |
| 172 | 0,2009 | 4,0908 | 0,4947 | | 0,0352 | 7,0000 |
| 173 | 0,2009 | 4,1133 | 0,4926 | | 0,0352 | 7,0000 |
| 174 | 0,2009 | 4,1357 | 0,4905 | | 0,0352 | 10,0000 |
| 175 | 0,2009 | 4,1582 | 0,4884 | | 0,0352 | 7,0000 |
| 176 | 0,2009 | 4,1807 | 0,4864 | | 0,0352 | 7,0000 |
| 177 | 0,2009 | 4,2031 | 0,4843 | | 0,0352 | 10,0000 |
| 178 | 0,2009 | 4,2256 | 0,4822 | | 0,0352 | 7,0000 |
| 179 | 0,2009 | 4,2480 | 0,4802 | | 0,0352 | 7,0000 |
| 180 | 0,2009 | 4,2705 | 0,4781 | | 0,0352 | 7,0000 |
| 181 | 0,2009 | 4,2930 | 0,4761 | | 0,0352 | 7,0000 |
| 182 | 0,2009 | 4,3154 | 0,4741 | | 0,0352 | 10,0000 |
| 183 | 0,2009 | 4,3379 | 0,4721 | | 0,0352 | 7,0000 |
| 184 | 0,2009 | 4,3604 | 0,4701 | | 0,0352 | 7,0000 |
| 185 | 0,2009 | 4,3828 | 0,4681 | | 0,0352 | 7,0000 |
| 186 | 0,2009 | 4,4053 | 0,4661 | | 0,0352 | 7,0000 |
| 187 | 0,2009 | 4,4277 | 0,4641 | | 0,0352 | 7,0000 |
| 188 | 0,2009 | 4,4502 | 0,4621 | | 0,0352 | 7,0000 |
| 189 | 0,2009 | 4,4727 | 0,4601 | | 0,0352 | 7,0000 |
| 190 | 0,2009 | 4,4951 | 0,4582 | | 0,0352 | 7,0000 |
| 191 | 0,2009 | 4,5176 | 0,4562 | | 0,0352 | 7,0000 |
| 192 | 0,2009 | 4,5400 | 0,4543 | | 0,0352 | 9,0000 |
| 193 | 0,2009 | 4,5625 | 0,4524 | | 0,0352 | 7,0000 |
| 194 | 0,2009 | 4,5850 | 0,4504 | | 0,0352 | 7,0000 |
| 195 | 0,2009 | 4,6074 | 0,4485 | | 0,0352 | 7,0000 |
| 196 | 0,2009 | 4,6299 | 0,4466 | | 0,0352 | 7,0000 |

| | 19-a-Final 1 | 20-b-Final 1 | 21-x column 2 | 22-y column 2 | 23 | 24-SumSq 2 |
|---|---|---|---|---|---|---|
| 148 | 1,0751 | 0,1897 | 3,5518 | 0,3613 | | 0,0664 |
| 149 | 1,0751 | 0,1897 | 3,5742 | 0,3588 | | 0,0664 |
| 150 | 1,0751 | 0,1897 | 3,5967 | 0,3564 | | 0,0664 |
| 151 | 1,0751 | 0,1897 | 3,6191 | 0,3540 | | 0,0664 |
| 152 | 1,0751 | 0,1897 | 3,6416 | 0,3515 | | 0,0664 |
| 153 | 1,0751 | 0,1897 | 3,6641 | 0,3491 | | 0,0664 |
| 154 | 1,0751 | 0,1897 | 3,6865 | 0,3468 | | 0,0664 |
| 155 | 1,0751 | 0,1897 | 3,7090 | 0,3444 | | 0,0664 |
| 156 | 1,0751 | 0,1897 | 3,7314 | 0,3420 | | 0,0664 |
| 157 | 1,0751 | 0,1897 | 3,7539 | 0,3397 | | 0,0664 |
| 158 | 1,0751 | 0,1897 | 3,7764 | 0,3374 | | 0,0664 |
| 159 | 1,0751 | 0,1897 | 3,7988 | 0,3351 | | 0,0664 |
| 160 | 1,0751 | 0,1897 | 3,8213 | 0,3328 | | 0,0664 |
| 161 | 1,0751 | 0,1897 | 3,8438 | 0,3305 | | 0,0664 |
| 162 | 1,0751 | 0,1897 | 3,8662 | 0,3282 | | 0,0664 |
| 163 | 1,0751 | 0,1897 | 3,8887 | 0,3260 | | 0,0664 |
| 164 | 1,0751 | 0,1897 | 3,9111 | 0,3238 | | 0,0664 |
| 165 | 1,0751 | 0,1897 | 3,9336 | 0,3216 | | 0,0664 |
| 166 | 1,0751 | 0,1897 | 3,9561 | 0,3194 | | 0,0664 |
| 167 | 1,0751 | 0,1897 | 3,9785 | 0,3172 | | 0,0664 |
| 168 | 1,0751 | 0,1897 | 4,0010 | 0,3150 | | 0,0664 |
| 169 | 1,0751 | 0,1897 | 4,0234 | 0,3129 | | 0,0664 |
| 170 | 1,0751 | 0,1897 | 4,0459 | 0,3107 | | 0,0664 |
| 171 | 1,0751 | 0,1897 | 4,0684 | 0,3086 | | 0,0664 |
| 172 | 1,0751 | 0,1897 | 4,0908 | 0,3065 | | 0,0664 |
| 173 | 1,0751 | 0,1897 | 4,1133 | 0,3044 | | 0,0664 |
| 174 | 1,0751 | 0,1897 | 4,1357 | 0,3023 | | 0,0664 |
| 175 | 1,0751 | 0,1897 | 4,1582 | 0,3003 | | 0,0664 |
| 176 | 1,0751 | 0,1897 | 4,1807 | 0,2982 | | 0,0664 |
| 177 | 1,0751 | 0,1897 | 4,2031 | 0,2962 | | 0,0664 |
| 178 | 1,0751 | 0,1897 | 4,2256 | 0,2941 | | 0,0664 |
| 179 | 1,0751 | 0,1897 | 4,2480 | 0,2921 | | 0,0664 |
| 180 | 1,0751 | 0,1897 | 4,2705 | 0,2901 | | 0,0664 |
| 181 | 1,0751 | 0,1897 | 4,2930 | 0,2882 | | 0,0664 |
| 182 | 1,0751 | 0,1897 | 4,3154 | 0,2862 | | 0,0664 |
| 183 | 1,0751 | 0,1897 | 4,3379 | 0,2842 | | 0,0664 |
| 184 | 1,0751 | 0,1897 | 4,3604 | 0,2823 | | 0,0664 |
| 185 | 1,0751 | 0,1897 | 4,3828 | 0,2804 | | 0,0664 |
| 186 | 1,0751 | 0,1897 | 4,4053 | 0,2784 | | 0,0664 |
| 187 | 1,0751 | 0,1897 | 4,4277 | 0,2765 | | 0,0664 |
| 188 | 1,0751 | 0,1897 | 4,4502 | 0,2747 | | 0,0664 |
| 189 | 1,0751 | 0,1897 | 4,4727 | 0,2728 | | 0,0664 |
| 190 | 1,0751 | 0,1897 | 4,4951 | 0,2709 | | 0,0664 |
| 191 | 1,0751 | 0,1897 | 4,5176 | 0,2691 | | 0,0664 |
| 192 | 1,0751 | 0,1897 | 4,5400 | 0,2672 | | 0,0664 |
| 193 | 1,0751 | 0,1897 | 4,5625 | 0,2654 | | 0,0664 |
| 194 | 1,0751 | 0,1897 | 4,5850 | 0,2636 | | 0,0664 |
| 195 | 1,0751 | 0,1897 | 4,6074 | 0,2618 | | 0,0664 |
| 196 | 1,0751 | 0,1897 | 4,6299 | 0,2600 | | 0,0664 |

| | 25-Iterations 2 | 26-a-Final 2 | 27-b-Final 2 | 28-x column 3 | 29-y column 3 | 30 |
|---|---|---|---|---|---|---|
| 148 | 11,0000 | 1,0684 | 0,3052 | 3,5518 | 0,2562 | |
| 149 | 11,0000 | 1,0684 | 0,3052 | 3,5742 | 0,2540 | |
| 150 | 10,0000 | 1,0684 | 0,3052 | 3,5967 | 0,2518 | |
| 151 | 11,0000 | 1,0684 | 0,3052 | 3,6191 | 0,2497 | |
| 152 | 11,0000 | 1,0684 | 0,3052 | 3,6416 | 0,2475 | |
| 153 | 11,0000 | 1,0684 | 0,3052 | 3,6641 | 0,2454 | |
| 154 | 14,0000 | 1,0684 | 0,3052 | 3,6865 | 0,2433 | |
| 155 | 11,0000 | 1,0684 | 0,3052 | 3,7090 | 0,2412 | |
| 156 | 11,0000 | 1,0684 | 0,3052 | 3,7314 | 0,2391 | |
| 157 | 11,0000 | 1,0684 | 0,3052 | 3,7539 | 0,2370 | |
| 158 | 11,0000 | 1,0684 | 0,3052 | 3,7764 | 0,2350 | |
| 159 | 11,0000 | 1,0684 | 0,3052 | 3,7988 | 0,2329 | |
| 160 | 11,0000 | 1,0684 | 0,3052 | 3,8213 | 0,2309 | |
| 161 | 11,0000 | 1,0684 | 0,3052 | 3,8438 | 0,2290 | |
| 162 | 11,0000 | 1,0684 | 0,3052 | 3,8662 | 0,2270 | |
| 163 | 11,0000 | 1,0684 | 0,3052 | 3,8887 | 0,2250 | |
| 164 | 11,0000 | 1,0684 | 0,3052 | 3,9111 | 0,2231 | |
| 165 | 11,0000 | 1,0684 | 0,3052 | 3,9336 | 0,2212 | |
| 166 | 11,0000 | 1,0684 | 0,3052 | 3,9561 | 0,2193 | |
| 167 | 11,0000 | 1,0684 | 0,3052 | 3,9785 | 0,2174 | |
| 168 | 11,0000 | 1,0684 | 0,3052 | 4,0010 | 0,2155 | |
| 169 | 9,0000 | 1,0684 | 0,3052 | 4,0234 | 0,2136 | |
| 170 | 11,0000 | 1,0684 | 0,3052 | 4,0459 | 0,2118 | |
| 171 | 9,0000 | 1,0684 | 0,3052 | 4,0684 | 0,2100 | |
| 172 | 10,0000 | 1,0684 | 0,3052 | 4,0908 | 0,2082 | |
| 173 | 10,0000 | 1,0684 | 0,3052 | 4,1133 | 0,2064 | |
| 174 | 16,0000 | 1,0684 | 0,3052 | 4,1357 | 0,2046 | |
| 175 | 8,0000 | 1,0684 | 0,3052 | 4,1582 | 0,2028 | |
| 176 | 9,0000 | 1,0684 | 0,3052 | 4,1807 | 0,2011 | |
| 177 | 9,0000 | 1,0684 | 0,3052 | 4,2031 | 0,1993 | |
| 178 | 8,0000 | 1,0684 | 0,3052 | 4,2256 | 0,1976 | |
| 179 | 10,0000 | 1,0684 | 0,3052 | 4,2480 | 0,1959 | |
| 180 | 10,0000 | 1,0684 | 0,3052 | 4,2705 | 0,1942 | |
| 181 | 22,0000 | 1,0684 | 0,3052 | 4,2930 | 0,1926 | |
| 182 | 10,0000 | 1,0684 | 0,3052 | 4,3154 | 0,1909 | |
| 183 | 9,0000 | 1,0684 | 0,3052 | 4,3379 | 0,1893 | |
| 184 | 8,0000 | 1,0684 | 0,3052 | 4,3604 | 0,1876 | |
| 185 | 9,0000 | 1,0684 | 0,3052 | 4,3828 | 0,1860 | |
| 186 | 10,0000 | 1,0684 | 0,3052 | 4,4053 | 0,1844 | |
| 187 | 14,0000 | 1,0684 | 0,3052 | 4,4277 | 0,1828 | |
| 188 | 8,0000 | 1,0684 | 0,3052 | 4,4502 | 0,1812 | |
| 189 | 10,0000 | 1,0684 | 0,3052 | 4,4727 | 0,1797 | |
| 190 | 14,0000 | 1,0684 | 0,3052 | 4,4951 | 0,1781 | |
| 191 | 14,0000 | 1,0684 | 0,3052 | 4,5176 | 0,1766 | |
| 192 | 9,0000 | 1,0684 | 0,3052 | 4,5400 | 0,1751 | |
| 193 | 10,0000 | 1,0684 | 0,3052 | 4,5625 | 0,1736 | |
| 194 | 14,0000 | 1,0684 | 0,3052 | 4,5850 | 0,1721 | |
| 195 | 10,0000 | 1,0684 | 0,3052 | 4,6074 | 0,1706 | |
| 196 | 11,0000 | 1,0684 | 0,3052 | 4,6299 | 0,1691 | |

| | 31-SumSq 3 | 32-Iterations 3 | 33-a-Final 3 | 34-b-Final 3 | 35-x column 4 | 36-y column 4 |
|---|---|---|---|---|---|---|
| 148 | 0,0429 | 11,0000 | 1,0070 | 0,3854 | 3,5518 | 0,2562 |
| 149 | 0,0429 | 11,0000 | 1,0070 | 0,3854 | 3,5742 | 0,2540 |
| 150 | 0,0429 | 11,0000 | 1,0070 | 0,3854 | 3,5967 | 0,2518 |
| 151 | 0,0429 | 12,0000 | 1,0070 | 0,3854 | 3,6191 | 0,2497 |
| 152 | 0,0429 | 11,0000 | 1,0070 | 0,3854 | 3,6416 | 0,2475 |
| 153 | 0,0429 | 16,0000 | 1,0070 | 0,3854 | 3,6641 | 0,2454 |
| 154 | 0,0429 | 12,0000 | 1,0070 | 0,3854 | 3,6865 | 0,2433 |
| 155 | 0,0429 | 16,0000 | 1,0070 | 0,3854 | 3,7090 | 0,2412 |
| 156 | 0,0429 | 16,0000 | 1,0070 | 0,3854 | 3,7314 | 0,2391 |
| 157 | 0,0429 | 10,0000 | 1,0070 | 0,3854 | 3,7539 | 0,2370 |
| 158 | 0,0429 | 11,0000 | 1,0070 | 0,3854 | 3,7764 | 0,2350 |
| 159 | 0,0429 | 12,0000 | 1,0070 | 0,3854 | 3,7988 | 0,2329 |
| 160 | 0,0429 | 12,0000 | 1,0070 | 0,3854 | 3,8213 | 0,2309 |
| 161 | 0,0429 | 12,0000 | 1,0070 | 0,3854 | 3,8438 | 0,2290 |
| 162 | 0,0429 | 12,0000 | 1,0070 | 0,3854 | 3,8662 | 0,2270 |
| 163 | 0,0429 | 10,0000 | 1,0070 | 0,3854 | 3,8887 | 0,2250 |
| 164 | 0,0429 | 12,0000 | 1,0070 | 0,3854 | 3,9111 | 0,2231 |
| 165 | 0,0429 | 12,0000 | 1,0070 | 0,3854 | 3,9336 | 0,2212 |
| 166 | 0,0429 | 15,0000 | 1,0070 | 0,3854 | 3,9561 | 0,2193 |
| 167 | 0,0429 | 9,0000 | 1,0070 | 0,3854 | 3,9785 | 0,2174 |
| 168 | 0,0429 | 12,0000 | 1,0070 | 0,3854 | 4,0010 | 0,2155 |
| 169 | 0,0429 | 12,0000 | 1,0070 | 0,3854 | 4,0234 | 0,2136 |
| 170 | 0,0429 | 12,0000 | 1,0070 | 0,3854 | 4,0459 | 0,2118 |
| 171 | 0,0429 | 41,0000 | 1,0070 | 0,3854 | 4,0684 | 0,2100 |
| 172 | 0,0429 | 12,0000 | 1,0070 | 0,3854 | 4,0908 | 0,2082 |
| 173 | 0,0429 | 12,0000 | 1,0070 | 0,3854 | 4,1133 | 0,2064 |
| 174 | 0,0429 | 12,0000 | 1,0070 | 0,3854 | 4,1357 | 0,2046 |
| 175 | 0,0429 | 12,0000 | 1,0070 | 0,3854 | 4,1582 | 0,2028 |
| 176 | 0,0429 | 12,0000 | 1,0070 | 0,3854 | 4,1807 | 0,2011 |
| 177 | 0,0429 | 12,0000 | 1,0070 | 0,3854 | 4,2031 | 0,1993 |
| 178 | 0,0429 | 13,0000 | 1,0070 | 0,3854 | 4,2256 | 0,1976 |
| 179 | 0,0429 | 12,0000 | 1,0070 | 0,3854 | 4,2480 | 0,1959 |
| 180 | 0,0429 | 12,0000 | 1,0070 | 0,3854 | 4,2705 | 0,1942 |
| 181 | 0,0429 | 11,0000 | 1,0070 | 0,3854 | 4,2930 | 0,1926 |
| 182 | 0,0429 | 12,0000 | 1,0070 | 0,3854 | 4,3154 | 0,1909 |
| 183 | 0,0429 | 12,0000 | 1,0070 | 0,3854 | 4,3379 | 0,1893 |
| 184 | 0,0429 | 10,0000 | 1,0070 | 0,3854 | 4,3604 | 0,1876 |
| 185 | 0,0429 | 12,0000 | 1,0070 | 0,3854 | 4,3828 | 0,1860 |
| 186 | 0,0429 | 15,0000 | 1,0070 | 0,3854 | 4,4053 | 0,1844 |
| 187 | 0,0429 | 12,0000 | 1,0070 | 0,3854 | 4,4277 | 0,1828 |
| 188 | 0,0429 | 10,0000 | 1,0070 | 0,3854 | 4,4502 | 0,1812 |
| 189 | 0,0429 | 10,0000 | 1,0070 | 0,3854 | 4,4727 | 0,1797 |
| 190 | 0,0429 | 12,0000 | 1,0070 | 0,3854 | 4,4951 | 0,1781 |
| 191 | 0,0429 | 12,0000 | 1,0070 | 0,3854 | 4,5176 | 0,1766 |
| 192 | 0,0429 | 9,0000 | 1,0070 | 0,3854 | 4,5400 | 0,1751 |
| 193 | 0,0429 | 10,0000 | 1,0070 | 0,3854 | 4,5625 | 0,1736 |
| 194 | 0,0429 | 10,0000 | 1,0070 | 0,3854 | 4,5850 | 0,1721 |
| 195 | 0,0429 | 10,0000 | 1,0070 | 0,3854 | 4,6074 | 0,1706 |
| 196 | 0,0429 | 10,0000 | 1,0070 | 0,3854 | 4,6299 | 0,1691 |

| | 37 | 38-SumSq 4 | 39-Iterations 4 | 40-a-Final 4 | 41-b-Final 4 | 42-x column 5 |
|---|---|---|---|---|---|---|
| 148 | | 0,0429 | 10,0000 | 1,0070 | 0,3854 | 3,5518 |
| 149 | | 0,0429 | 10,0000 | 1,0070 | 0,3854 | 3,5742 |
| 150 | | 0,0429 | 10,0000 | 1,0070 | 0,3854 | 3,5967 |
| 151 | | 0,0429 | 10,0000 | 1,0070 | 0,3854 | 3,6191 |
| 152 | | 0,0429 | 10,0000 | 1,0070 | 0,3854 | 3,6416 |
| 153 | | 0,0429 | 10,0000 | 1,0070 | 0,3854 | 3,6641 |
| 154 | | 0,0429 | 8,0000 | 1,0070 | 0,3854 | 3,6865 |
| 155 | | 0,0429 | 10,0000 | 1,0070 | 0,3854 | 3,7090 |
| 156 | | 0,0429 | 10,0000 | 1,0070 | 0,3854 | 3,7314 |
| 157 | | 0,0429 | 10,0000 | 1,0070 | 0,3854 | 3,7539 |
| 158 | | 0,0429 | 10,0000 | 1,0070 | 0,3854 | 3,7764 |
| 159 | | 0,0429 | 10,0000 | 1,0070 | 0,3854 | 3,7988 |
| 160 | | 0,0429 | 10,0000 | 1,0070 | 0,3854 | 3,8213 |
| 161 | | 0,0429 | 10,0000 | 1,0070 | 0,3854 | 3,8438 |
| 162 | | 0,0429 | 10,0000 | 1,0070 | 0,3854 | 3,8662 |
| 163 | | 0,0429 | 10,0000 | 1,0070 | 0,3854 | 3,8887 |
| 164 | | 0,0429 | 10,0000 | 1,0070 | 0,3854 | 3,9111 |
| 165 | | 0,0429 | 11,0000 | 1,0070 | 0,3854 | 3,9336 |
| 166 | | 0,0429 | 10,0000 | 1,0070 | 0,3854 | 3,9561 |
| 167 | | 0,0429 | 11,0000 | 1,0070 | 0,3854 | 3,9785 |
| 168 | | 0,0429 | 10,0000 | 1,0070 | 0,3854 | 4,0010 |
| 169 | | 0,0429 | 11,0000 | 1,0070 | 0,3854 | 4,0234 |
| 170 | | 0,0429 | 9,0000 | 1,0070 | 0,3854 | 4,0459 |
| 171 | | 0,0429 | 11,0000 | 1,0070 | 0,3854 | 4,0684 |
| 172 | | 0,0429 | 10,0000 | 1,0070 | 0,3854 | 4,0908 |
| 173 | | 0,0429 | 8,0000 | 1,0070 | 0,3854 | 4,1133 |
| 174 | | 0,0429 | 11,0000 | 1,0070 | 0,3854 | 4,1357 |
| 175 | | 0,0429 | 9,0000 | 1,0070 | 0,3854 | 4,1582 |
| 176 | | 0,0429 | 9,0000 | 1,0070 | 0,3854 | 4,1807 |
| 177 | | 0,0429 | 10,0000 | 1,0070 | 0,3854 | 4,2031 |
| 178 | | 0,0429 | 11,0000 | 1,0070 | 0,3854 | 4,2256 |
| 179 | | 0,0429 | 11,0000 | 1,0070 | 0,3854 | 4,2480 |
| 180 | | 0,0429 | 9,0000 | 1,0070 | 0,3854 | 4,2705 |
| 181 | | 0,0429 | 11,0000 | 1,0070 | 0,3854 | 4,2930 |
| 182 | | 0,0429 | 10,0000 | 1,0070 | 0,3854 | 4,3154 |
| 183 | | 0,0429 | 7,0000 | 1,0070 | 0,3854 | 4,3379 |
| 184 | | 0,0429 | 9,0000 | 1,0070 | 0,3854 | 4,3604 |
| 185 | | 0,0429 | 11,0000 | 1,0070 | 0,3854 | 4,3828 |
| 186 | | 0,0429 | 10,0000 | 1,0070 | 0,3854 | 4,4053 |
| 187 | | 0,0429 | 7,0000 | 1,0070 | 0,3854 | 4,4277 |
| 188 | | 0,0429 | 11,0000 | 1,0070 | 0,3854 | 4,4502 |
| 189 | | 0,0429 | 10,0000 | 1,0070 | 0,3854 | 4,4727 |
| 190 | | 0,0429 | 11,0000 | 1,0070 | 0,3854 | 4,4951 |
| 191 | | 0,0429 | 11,0000 | 1,0070 | 0,3854 | 4,5176 |
| 192 | | 0,0429 | 10,0000 | 1,0070 | 0,3854 | 4,5400 |
| 193 | | 0,0429 | 9,0000 | 1,0070 | 0,3854 | 4,5625 |
| 194 | | 0,0429 | 11,0000 | 1,0070 | 0,3854 | 4,5850 |
| 195 | | 0,0429 | 9,0000 | 1,0070 | 0,3854 | 4,6074 |
| 196 | | 0,0429 | 10,0000 | 1,0070 | 0,3854 | 4,6299 |

| | 43-y column 5 | 44 | 45-SumSq 5 | 46-Iterations 5 | 47-a-Final 5 | 48-b-Final 5 |
|---|---|---|---|---|---|---|
| 148 | 0,4296 | | 0,1194 | 10,0000 | 1,0036 | 0,2389 |
| 149 | 0,4273 | | 0,1194 | 9,0000 | 1,0036 | 0,2389 |
| 150 | 0,4250 | | 0,1194 | 9,0000 | 1,0036 | 0,2389 |
| 151 | 0,4228 | | 0,1194 | 9,0000 | 1,0036 | 0,2389 |
| 152 | 0,4205 | | 0,1194 | 9,0000 | 1,0036 | 0,2389 |
| 153 | 0,4182 | | 0,1194 | 9,0000 | 1,0036 | 0,2389 |
| 154 | 0,4160 | | 0,1194 | 9,0000 | 1,0036 | 0,2389 |
| 155 | 0,4138 | | 0,1194 | 10,0000 | 1,0036 | 0,2389 |
| 156 | 0,4116 | | 0,1194 | 10,0000 | 1,0036 | 0,2389 |
| 157 | 0,4094 | | 0,1194 | 9,0000 | 1,0036 | 0,2389 |
| 158 | 0,4072 | | 0,1194 | 9,0000 | 1,0036 | 0,2389 |
| 159 | 0,4050 | | 0,1194 | 9,0000 | 1,0036 | 0,2389 |
| 160 | 0,4028 | | 0,1194 | 9,0000 | 1,0036 | 0,2389 |
| 161 | 0,4007 | | 0,1194 | 10,0000 | 1,0036 | 0,2389 |
| 162 | 0,3985 | | 0,1194 | 9,0000 | 1,0036 | 0,2389 |
| 163 | 0,3964 | | 0,1194 | 9,0000 | 1,0036 | 0,2389 |
| 164 | 0,3943 | | 0,1194 | 9,0000 | 1,0036 | 0,2389 |
| 165 | 0,3922 | | 0,1194 | 9,0000 | 1,0036 | 0,2389 |
| 166 | 0,3901 | | 0,1194 | 9,0000 | 1,0036 | 0,2389 |
| 167 | 0,3880 | | 0,1194 | 10,0000 | 1,0036 | 0,2389 |
| 168 | 0,3859 | | 0,1194 | 9,0000 | 1,0036 | 0,2389 |
| 169 | 0,3838 | | 0,1194 | 9,0000 | 1,0036 | 0,2389 |
| 170 | 0,3818 | | 0,1194 | 9,0000 | 1,0036 | 0,2389 |
| 171 | 0,3797 | | 0,1194 | 9,0000 | 1,0036 | 0,2389 |
| 172 | 0,3777 | | 0,1194 | 9,0000 | 1,0036 | 0,2389 |
| 173 | 0,3757 | | 0,1194 | 9,0000 | 1,0036 | 0,2389 |
| 174 | 0,3737 | | 0,1194 | 10,0000 | 1,0036 | 0,2389 |
| 175 | 0,3717 | | 0,1194 | 9,0000 | 1,0036 | 0,2389 |
| 176 | 0,3697 | | 0,1194 | 9,0000 | 1,0036 | 0,2389 |
| 177 | 0,3677 | | 0,1194 | 9,0000 | 1,0036 | 0,2389 |
| 178 | 0,3657 | | 0,1194 | 9,0000 | 1,0036 | 0,2389 |
| 179 | 0,3638 | | 0,1194 | 9,0000 | 1,0036 | 0,2389 |
| 180 | 0,3618 | | 0,1194 | 9,0000 | 1,0036 | 0,2389 |
| 181 | 0,3599 | | 0,1194 | 9,0000 | 1,0036 | 0,2389 |
| 182 | 0,3580 | | 0,1194 | 9,0000 | 1,0036 | 0,2389 |
| 183 | 0,3561 | | 0,1194 | 8,0000 | 1,0036 | 0,2389 |
| 184 | 0,3542 | | 0,1194 | 10,0000 | 1,0036 | 0,2389 |
| 185 | 0,3523 | | 0,1194 | 8,0000 | 1,0036 | 0,2389 |
| 186 | 0,3504 | | 0,1194 | 8,0000 | 1,0036 | 0,2389 |
| 187 | 0,3485 | | 0,1194 | 8,0000 | 1,0036 | 0,2389 |
| 188 | 0,3466 | | 0,1194 | 10,0000 | 1,0036 | 0,2389 |
| 189 | 0,3448 | | 0,1194 | 8,0000 | 1,0036 | 0,2389 |
| 190 | 0,3429 | | 0,1194 | 10,0000 | 1,0036 | 0,2389 |
| 191 | 0,3411 | | 0,1194 | 9,0000 | 1,0036 | 0,2389 |
| 192 | 0,3393 | | 0,1194 | 10,0000 | 1,0036 | 0,2389 |
| 193 | 0,3375 | | 0,1194 | 7,0000 | 1,0036 | 0,2389 |
| 194 | 0,3357 | | 0,1194 | 9,0000 | 1,0036 | 0,2389 |
| 195 | 0,3339 | | 0,1194 | 10,0000 | 1,0036 | 0,2389 |
| 196 | 0,3321 | | 0,1194 | 10,0000 | 1,0036 | 0,2389 |

| | 49-x column 6 | 50-y column 6 | 51 | 52-SumSq 6 | 53-Iterations 6 | 54-a-Final 6 |
|---|---|---|---|---|---|---|
| 148 | 3,5518 | 0,3172 | | 0,0865 | 10,0000 | 0,9419 |
| 149 | 3,5742 | 0,3150 | | 0,0865 | 10,0000 | 0,9419 |
| 150 | 3,5967 | 0,3129 | | 0,0865 | 10,0000 | 0,9419 |
| 151 | 3,6191 | 0,3107 | | 0,0865 | 10,0000 | 0,9419 |
| 152 | 3,6416 | 0,3086 | | 0,0865 | 10,0000 | 0,9419 |
| 153 | 3,6641 | 0,3065 | | 0,0865 | 10,0000 | 0,9419 |
| 154 | 3,6865 | 0,3044 | | 0,0865 | 10,0000 | 0,9419 |
| 155 | 3,7090 | 0,3023 | | 0,0865 | 10,0000 | 0,9419 |
| 156 | 3,7314 | 0,3002 | | 0,0865 | 10,0000 | 0,9419 |
| 157 | 3,7539 | 0,2982 | | 0,0865 | 10,0000 | 0,9419 |
| 158 | 3,7764 | 0,2961 | | 0,0865 | 10,0000 | 0,9419 |
| 159 | 3,7988 | 0,2941 | | 0,0865 | 10,0000 | 0,9419 |
| 160 | 3,8213 | 0,2921 | | 0,0865 | 10,0000 | 0,9419 |
| 161 | 3,8438 | 0,2901 | | 0,0865 | 10,0000 | 0,9419 |
| 162 | 3,8662 | 0,2881 | | 0,0865 | 10,0000 | 0,9419 |
| 163 | 3,8887 | 0,2861 | | 0,0865 | 10,0000 | 0,9419 |
| 164 | 3,9111 | 0,2841 | | 0,0865 | 10,0000 | 0,9419 |
| 165 | 3,9336 | 0,2822 | | 0,0865 | 9,0000 | 0,9419 |
| 166 | 3,9561 | 0,2802 | | 0,0865 | 10,0000 | 0,9419 |
| 167 | 3,9785 | 0,2783 | | 0,0865 | 10,0000 | 0,9419 |
| 168 | 4,0010 | 0,2764 | | 0,0865 | 10,0000 | 0,9419 |
| 169 | 4,0234 | 0,2745 | | 0,0865 | 10,0000 | 0,9419 |
| 170 | 4,0459 | 0,2726 | | 0,0865 | 10,0000 | 0,9419 |
| 171 | 4,0684 | 0,2708 | | 0,0865 | 10,0000 | 0,9419 |
| 172 | 4,0908 | 0,2689 | | 0,0865 | 10,0000 | 0,9419 |
| 173 | 4,1133 | 0,2671 | | 0,0865 | 10,0000 | 0,9419 |
| 174 | 4,1357 | 0,2652 | | 0,0865 | 10,0000 | 0,9419 |
| 175 | 4,1582 | 0,2634 | | 0,0865 | 10,0000 | 0,9419 |
| 176 | 4,1807 | 0,2616 | | 0,0865 | 10,0000 | 0,9419 |
| 177 | 4,2031 | 0,2598 | | 0,0865 | 10,0000 | 0,9419 |
| 178 | 4,2256 | 0,2580 | | 0,0865 | 10,0000 | 0,9419 |
| 179 | 4,2480 | 0,2563 | | 0,0865 | 10,0000 | 0,9419 |
| 180 | 4,2705 | 0,2545 | | 0,0865 | 10,0000 | 0,9419 |
| 181 | 4,2930 | 0,2528 | | 0,0865 | 10,0000 | 0,9419 |
| 182 | 4,3154 | 0,2510 | | 0,0865 | 10,0000 | 0,9419 |
| 183 | 4,3379 | 0,2493 | | 0,0865 | 10,0000 | 0,9419 |
| 184 | 4,3604 | 0,2476 | | 0,0865 | 10,0000 | 0,9419 |
| 185 | 4,3828 | 0,2459 | | 0,0865 | 10,0000 | 0,9419 |
| 186 | 4,4053 | 0,2442 | | 0,0865 | 10,0000 | 0,9419 |
| 187 | 4,4277 | 0,2425 | | 0,0865 | 10,0000 | 0,9419 |
| 188 | 4,4502 | 0,2409 | | 0,0865 | 10,0000 | 0,9419 |
| 189 | 4,4727 | 0,2392 | | 0,0865 | 10,0000 | 0,9419 |
| 190 | 4,4951 | 0,2376 | | 0,0865 | 10,0000 | 0,9419 |
| 191 | 4,5176 | 0,2359 | | 0,0865 | 10,0000 | 0,9419 |
| 192 | 4,5400 | 0,2343 | | 0,0865 | 10,0000 | 0,9419 |
| 193 | 4,5625 | 0,2327 | | 0,0865 | 10,0000 | 0,9419 |
| 194 | 4,5850 | 0,2311 | | 0,0865 | 10,0000 | 0,9419 |
| 195 | 4,6074 | 0,2295 | | 0,0865 | 10,0000 | 0,9419 |
| 196 | 4,6299 | 0,2280 | | 0,0865 | 10,0000 | 0,9419 |

| | 55-b-Final 6 |
|---|---|
| 148 | 0,3064 |
| 149 | 0,3064 |
| 150 | 0,3064 |
| 151 | 0,3064 |
| 152 | 0,3064 |
| 153 | 0,3064 |
| 154 | 0,3064 |
| 155 | 0,3064 |
| 156 | 0,3064 |
| 157 | 0,3064 |
| 158 | 0,3064 |
| 159 | 0,3064 |
| 160 | 0,3064 |
| 161 | 0,3064 |
| 162 | 0,3064 |
| 163 | 0,3064 |
| 164 | 0,3064 |
| 165 | 0,3064 |
| 166 | 0,3064 |
| 167 | 0,3064 |
| 168 | 0,3064 |
| 169 | 0,3064 |
| 170 | 0,3064 |
| 171 | 0,3064 |
| 172 | 0,3064 |
| 173 | 0,3064 |
| 174 | 0,3064 |
| 175 | 0,3064 |
| 176 | 0,3064 |
| 177 | 0,3064 |
| 178 | 0,3064 |
| 179 | 0,3064 |
| 180 | 0,3064 |
| 181 | 0,3064 |
| 182 | 0,3064 |
| 183 | 0,3064 |
| 184 | 0,3064 |
| 185 | 0,3064 |
| 186 | 0,3064 |
| 187 | 0,3064 |
| 188 | 0,3064 |
| 189 | 0,3064 |
| 190 | 0,3064 |
| 191 | 0,3064 |
| 192 | 0,3064 |
| 193 | 0,3064 |
| 194 | 0,3064 |
| 195 | 0,3064 |
| 196 | 0,3064 |

| | 1 | 2 | 3 | 4 | 5 | 6 |
|---|---|---|---|---|---|---|
| 197 | | | | | | |
| 198 | | | | | | |
| 199 | | | | | | |
| 200 | | | | | | |
| 201 | | | | | | |
| 202 | | | | | | |
| 203 | | | | | | |
| 204 | | | | | | |
| 205 | | | | | | |
| 206 | | | | | | |
| 207 | | | | | | |
| 208 | | | | | | |
| 209 | | | | | | |
| 210 | | | | | | |
| 211 | | | | | | |
| 212 | | | | | | |
| 213 | | | | | | |
| 214 | | | | | | |
| 215 | | | | | | |
| 216 | | | | | | |
| 217 | | | | | | |
| 218 | | | | | | |
| 219 | | | | | | |
| 220 | | | | | | |
| 221 | | | | | | |
| 222 | | | | | | |
| 223 | | | | | | |
| 224 | | | | | | |
| 225 | | | | | | |
| 226 | | | | | | |
| 227 | | | | | | |
| 228 | | | | | | |
| 229 | | | | | | |
| 230 | | | | | | |
| 231 | | | | | | |
| 232 | | | | | | |
| 233 | | | | | | |
| 234 | | | | | | |
| 235 | | | | | | |
| 236 | | | | | | |
| 237 | | | | | | |
| 238 | | | | | | |
| 239 | | | | | | |
| 240 | | | | | | |
| 241 | | | | | | |
| 242 | | | | | | |
| 243 | | | | | | |
| 244 | | | | | | |
| 245 | | | | | | |

| | 7-x column | 8-y column | 9 | 10-SumSq | 11-Iterations | 12-a-Final |
|---|---|---|---|---|---|---|
| 197 | 4,6523 | 0,4309 | | 0,0411 | 7,0000 | 1,0972 |
| 198 | 4,6748 | 0,4290 | | 0,0411 | 7,0000 | 1,0972 |
| 199 | 4,6973 | 0,4270 | | 0,0411 | 7,0000 | 1,0972 |
| 200 | 4,7197 | 0,4251 | | 0,0411 | 6,0000 | 1,0972 |
| 201 | 4,7422 | 0,4232 | | | | |
| 202 | 4,7646 | 0,4213 | | | | |
| 203 | 4,7871 | 0,4194 | | | | |
| 204 | 4,8096 | 0,4175 | | | | |
| 205 | 4,8320 | 0,4156 | | | | |
| 206 | 4,8545 | 0,4138 | | | | |
| 207 | 4,8770 | 0,4119 | | | | |
| 208 | 4,8994 | 0,4100 | | | | |
| 209 | 4,9219 | 0,4082 | | | | |
| 210 | 4,9443 | 0,4064 | | | | |
| 211 | 4,9668 | 0,4045 | | | | |
| 212 | 4,9893 | 0,4027 | | | | |
| 213 | 5,0117 | 0,4009 | | | | |
| 214 | 5,0342 | 0,3991 | | | | |
| 215 | 5,0566 | 0,3973 | | | | |
| 216 | 5,0791 | 0,3955 | | | | |
| 217 | 5,1016 | 0,3937 | | | | |
| 218 | 5,1240 | 0,3919 | | | | |
| 219 | 5,1465 | 0,3902 | | | | |
| 220 | 5,1689 | 0,3884 | | | | |
| 221 | 5,1914 | 0,3867 | | | | |
| 222 | 5,2139 | 0,3849 | | | | |
| 223 | 5,2363 | 0,3832 | | | | |
| 224 | 5,2588 | 0,3815 | | | | |
| 225 | 5,2813 | 0,3798 | | | | |
| 226 | 5,3037 | 0,3780 | | | | |
| 227 | 5,3262 | 0,3763 | | | | |
| 228 | 5,3486 | 0,3747 | | | | |
| 229 | 5,3711 | 0,3730 | | | | |
| 230 | 5,3936 | 0,3713 | | | | |
| 231 | 5,4160 | 0,3696 | | | | |
| 232 | 5,4385 | 0,3680 | | | | |
| 233 | 5,4609 | 0,3663 | | | | |
| 234 | 5,4834 | 0,3646 | | | | |
| 235 | 5,5059 | 0,3630 | | | | |
| 236 | 5,5283 | 0,3614 | | | | |
| 237 | 5,5508 | 0,3597 | | | | |
| 238 | 5,5732 | 0,3581 | | | | |
| 239 | 5,5957 | 0,3565 | | | | |
| 240 | 5,6182 | 0,3549 | | | | |
| 241 | 5,6406 | 0,3533 | | | | |
| 242 | 5,6631 | 0,3517 | | | | |
| 243 | 5,6855 | 0,3501 | | | | |
| 244 | 5,7080 | 0,3486 | | | | |
| 245 | 5,7305 | 0,3470 | | | | |

| | 13-b-Final | 14-x column 1 | 15-y column 1 | 16 | 17-SumSq 1 | 18-Iterations 1 |
|---|---|---|---|---|---|---|
| 197 | 0,2009 | 4,6523 | 0,4447 | | 0,0352 | 7,0000 |
| 198 | 0,2009 | 4,6748 | 0,4428 | | 0,0352 | 7,0000 |
| 199 | 0,2009 | 4,6973 | 0,4409 | | 0,0352 | 7,0000 |
| 200 | 0,2009 | 4,7197 | 0,4391 | | 0,0352 | 6,0000 |
| 201 | | 4,7422 | 0,4372 | | | |
| 202 | | 4,7646 | 0,4353 | | | |
| 203 | | 4,7871 | 0,4335 | | | |
| 204 | | 4,8096 | 0,4317 | | | |
| 205 | | 4,8320 | 0,4298 | | | |
| 206 | | 4,8545 | 0,4280 | | | |
| 207 | | 4,8770 | 0,4262 | | | |
| 208 | | 4,8994 | 0,4244 | | | |
| 209 | | 4,9219 | 0,4226 | | | |
| 210 | | 4,9443 | 0,4208 | | | |
| 211 | | 4,9668 | 0,4190 | | | |
| 212 | | 4,9893 | 0,4172 | | | |
| 213 | | 5,0117 | 0,4154 | | | |
| 214 | | 5,0342 | 0,4136 | | | |
| 215 | | 5,0566 | 0,4119 | | | |
| 216 | | 5,0791 | 0,4101 | | | |
| 217 | | 5,1016 | 0,4084 | | | |
| 218 | | 5,1240 | 0,4067 | | | |
| 219 | | 5,1465 | 0,4049 | | | |
| 220 | | 5,1689 | 0,4032 | | | |
| 221 | | 5,1914 | 0,4015 | | | |
| 222 | | 5,2139 | 0,3998 | | | |
| 223 | | 5,2363 | 0,3981 | | | |
| 224 | | 5,2588 | 0,3964 | | | |
| 225 | | 5,2813 | 0,3947 | | | |
| 226 | | 5,3037 | 0,3930 | | | |
| 227 | | 5,3262 | 0,3914 | | | |
| 228 | | 5,3486 | 0,3897 | | | |
| 229 | | 5,3711 | 0,3880 | | | |
| 230 | | 5,3936 | 0,3864 | | | |
| 231 | | 5,4160 | 0,3847 | | | |
| 232 | | 5,4385 | 0,3831 | | | |
| 233 | | 5,4609 | 0,3815 | | | |
| 234 | | 5,4834 | 0,3798 | | | |
| 235 | | 5,5059 | 0,3782 | | | |
| 236 | | 5,5283 | 0,3766 | | | |
| 237 | | 5,5508 | 0,3750 | | | |
| 238 | | 5,5732 | 0,3734 | | | |
| 239 | | 5,5957 | 0,3718 | | | |
| 240 | | 5,6182 | 0,3703 | | | |
| 241 | | 5,6406 | 0,3687 | | | |
| 242 | | 5,6631 | 0,3671 | | | |
| 243 | | 5,6855 | 0,3656 | | | |
| 244 | | 5,7080 | 0,3640 | | | |
| 245 | | 5,7305 | 0,3625 | | | |

| | 19-a-Final 1 | 20-b-Final 1 | 21-x column 2 | 22-y column 2 | 23 | 24-SumSq 2 |
|---|---|---|---|---|---|---|
| 197 | 1,0751 | 0,1897 | 4,6523 | 0,2582 | | 0,0664 |
| 198 | 1,0751 | 0,1897 | 4,6748 | 0,2565 | | 0,0664 |
| 199 | 1,0751 | 0,1897 | 4,6973 | 0,2547 | | 0,0664 |
| 200 | 1,0751 | 0,1897 | 4,7197 | 0,2530 | | 1,3142 |
| 201 | | | 4,7422 | 0,2512 | | |
| 202 | | | 4,7646 | 0,2495 | | |
| 203 | | | 4,7871 | 0,2478 | | |
| 204 | | | 4,8096 | 0,2461 | | |
| 205 | | | 4,8320 | 0,2444 | | |
| 206 | | | 4,8545 | 0,2428 | | |
| 207 | | | 4,8770 | 0,2411 | | |
| 208 | | | 4,8994 | 0,2395 | | |
| 209 | | | 4,9219 | 0,2378 | | |
| 210 | | | 4,9443 | 0,2362 | | |
| 211 | | | 4,9668 | 0,2346 | | |
| 212 | | | 4,9893 | 0,2330 | | |
| 213 | | | 5,0117 | 0,2314 | | |
| 214 | | | 5,0342 | 0,2298 | | |
| 215 | | | 5,0566 | 0,2282 | | |
| 216 | | | 5,0791 | 0,2267 | | |
| 217 | | | 5,1016 | 0,2251 | | |
| 218 | | | 5,1240 | 0,2236 | | |
| 219 | | | 5,1465 | 0,2221 | | |
| 220 | | | 5,1689 | 0,2205 | | |
| 221 | | | 5,1914 | 0,2190 | | |
| 222 | | | 5,2139 | 0,2175 | | |
| 223 | | | 5,2363 | 0,2161 | | |
| 224 | | | 5,2588 | 0,2146 | | |
| 225 | | | 5,2813 | 0,2131 | | |
| 226 | | | 5,3037 | 0,2117 | | |
| 227 | | | 5,3262 | 0,2102 | | |
| 228 | | | 5,3486 | 0,2088 | | |
| 229 | | | 5,3711 | 0,2074 | | |
| 230 | | | 5,3936 | 0,2059 | | |
| 231 | | | 5,4160 | 0,2045 | | |
| 232 | | | 5,4385 | 0,2031 | | |
| 233 | | | 5,4609 | 0,2017 | | |
| 234 | | | 5,4834 | 0,2004 | | |
| 235 | | | 5,5059 | 0,1990 | | |
| 236 | | | 5,5283 | 0,1976 | | |
| 237 | | | 5,5508 | 0,1963 | | |
| 238 | | | 5,5732 | 0,1949 | | |
| 239 | | | 5,5957 | 0,1936 | | |
| 240 | | | 5,6182 | 0,1923 | | |
| 241 | | | 5,6406 | 0,1910 | | |
| 242 | | | 5,6631 | 0,1897 | | |
| 243 | | | 5,6855 | 0,1884 | | |
| 244 | | | 5,7080 | 0,1871 | | |
| 245 | | | 5,7305 | 0,1858 | | |

| | 25-Iterations 2 | 26-a-Final 2 | 27-b-Final 2 | 28-x column 3 | 29-y column 3 | 30 |
|---|---|---|---|---|---|---|
| 197 | 10,0000 | 1,0684 | 0,3052 | 4,6523 | 0,1677 | |
| 198 | 10,0000 | 1,0684 | 0,3052 | 4,6748 | 0,1662 | |
| 199 | 13,0000 | 1,0684 | 0,3052 | 4,6973 | 0,1648 | |
| 200 | 22,0000 | 892,1815 | 27,1747 | 4,7197 | 0,1634 | |
| 201 | | | | 4,7422 | 0,1619 | |
| 202 | | | | 4,7646 | 0,1606 | |
| 203 | | | | 4,7871 | 0,1592 | |
| 204 | | | | 4,8096 | 0,1578 | |
| 205 | | | | 4,8320 | 0,1564 | |
| 206 | | | | 4,8545 | 0,1551 | |
| 207 | | | | 4,8770 | 0,1538 | |
| 208 | | | | 4,8994 | 0,1524 | |
| 209 | | | | 4,9219 | 0,1511 | |
| 210 | | | | 4,9443 | 0,1498 | |
| 211 | | | | 4,9668 | 0,1485 | |
| 212 | | | | 4,9893 | 0,1472 | |
| 213 | | | | 5,0117 | 0,1460 | |
| 214 | | | | 5,0342 | 0,1447 | |
| 215 | | | | 5,0566 | 0,1435 | |
| 216 | | | | 5,0791 | 0,1422 | |
| 217 | | | | 5,1016 | 0,1410 | |
| 218 | | | | 5,1240 | 0,1398 | |
| 219 | | | | 5,1465 | 0,1386 | |
| 220 | | | | 5,1689 | 0,1374 | |
| 221 | | | | 5,1914 | 0,1362 | |
| 222 | | | | 5,2139 | 0,1350 | |
| 223 | | | | 5,2363 | 0,1339 | |
| 224 | | | | 5,2588 | 0,1327 | |
| 225 | | | | 5,2813 | 0,1316 | |
| 226 | | | | 5,3037 | 0,1304 | |
| 227 | | | | 5,3262 | 0,1293 | |
| 228 | | | | 5,3486 | 0,1282 | |
| 229 | | | | 5,3711 | 0,1271 | |
| 230 | | | | 5,3936 | 0,1260 | |
| 231 | | | | 5,4160 | 0,1249 | |
| 232 | | | | 5,4385 | 0,1238 | |
| 233 | | | | 5,4609 | 0,1228 | |
| 234 | | | | 5,4834 | 0,1217 | |
| 235 | | | | 5,5059 | 0,1207 | |
| 236 | | | | 5,5283 | 0,1196 | |
| 237 | | | | 5,5508 | 0,1186 | |
| 238 | | | | 5,5732 | 0,1176 | |
| 239 | | | | 5,5957 | 0,1166 | |
| 240 | | | | 5,6182 | 0,1156 | |
| 241 | | | | 5,6406 | 0,1146 | |
| 242 | | | | 5,6631 | 0,1136 | |
| 243 | | | | 5,6855 | 0,1126 | |
| 244 | | | | 5,7080 | 0,1116 | |
| 245 | | | | 5,7305 | 0,1107 | |

| | 31-SumSq 3 | 32-Iterations 3 | 33-a-Final 3 | 34-b-Final 3 | 35-x column 4 | 36-y column 4 |
|---|---|---|---|---|---|---|
| 197 | 0,0429 | 10,0000 | 1,0070 | 0,3854 | 4,6523 | 0,1677 |
| 198 | 0,0429 | 11,0000 | 1,0070 | 0,3854 | 4,6748 | 0,1662 |
| 199 | 0,7225 | 22,0000 | 6807,1438 | 35,3029 | 4,6973 | 0,1648 |
| 200 | 0,7225 | 22,0000 | 19219,7409 | 39,4548 | 4,7197 | 0,1634 |
| 201 | | | | | 4,7422 | 0,1619 |
| 202 | | | | | 4,7646 | 0,1606 |
| 203 | | | | | 4,7871 | 0,1592 |
| 204 | | | | | 4,8096 | 0,1578 |
| 205 | | | | | 4,8320 | 0,1564 |
| 206 | | | | | 4,8545 | 0,1551 |
| 207 | | | | | 4,8770 | 0,1538 |
| 208 | | | | | 4,8994 | 0,1524 |
| 209 | | | | | 4,9219 | 0,1511 |
| 210 | | | | | 4,9443 | 0,1498 |
| 211 | | | | | 4,9668 | 0,1485 |
| 212 | | | | | 4,9893 | 0,1472 |
| 213 | | | | | 5,0117 | 0,1460 |
| 214 | | | | | 5,0342 | 0,1447 |
| 215 | | | | | 5,0566 | 0,1435 |
| 216 | | | | | 5,0791 | 0,1422 |
| 217 | | | | | 5,1016 | 0,1410 |
| 218 | | | | | 5,1240 | 0,1398 |
| 219 | | | | | 5,1465 | 0,1386 |
| 220 | | | | | 5,1689 | 0,1374 |
| 221 | | | | | 5,1914 | 0,1362 |
| 222 | | | | | 5,2139 | 0,1350 |
| 223 | | | | | 5,2363 | 0,1339 |
| 224 | | | | | 5,2588 | 0,1327 |
| 225 | | | | | 5,2813 | 0,1316 |
| 226 | | | | | 5,3037 | 0,1304 |
| 227 | | | | | 5,3262 | 0,1293 |
| 228 | | | | | 5,3486 | 0,1282 |
| 229 | | | | | 5,3711 | 0,1271 |
| 230 | | | | | 5,3936 | 0,1260 |
| 231 | | | | | 5,4160 | 0,1249 |
| 232 | | | | | 5,4385 | 0,1238 |
| 233 | | | | | 5,4609 | 0,1228 |
| 234 | | | | | 5,4834 | 0,1217 |
| 235 | | | | | 5,5059 | 0,1207 |
| 236 | | | | | 5,5283 | 0,1196 |
| 237 | | | | | 5,5508 | 0,1186 |
| 238 | | | | | 5,5732 | 0,1176 |
| 239 | | | | | 5,5957 | 0,1166 |
| 240 | | | | | 5,6182 | 0,1156 |
| 241 | | | | | 5,6406 | 0,1146 |
| 242 | | | | | 5,6631 | 0,1136 |
| 243 | | | | | 5,6855 | 0,1126 |
| 244 | | | | | 5,7080 | 0,1116 |
| 245 | | | | | 5,7305 | 0,1107 |

| | 37 | 38-SumSq 4 | 39-Iterations 4 | 40-a-Final 4 | 41-b-Final 4 | 42-x column 5 |
|---|---|---|---|---|---|---|
| 197 | | 0,0429 | 9,0000 | 1,0070 | 0,3854 | 4,6523 |
| 198 | | 0,0429 | 11,0000 | 1,0070 | 0,3854 | 4,6748 |
| 199 | | 0,0429 | 9,0000 | 1,0070 | 0,3854 | 4,6973 |
| 200 | | 0,0429 | 10,0000 | 1,0070 | 0,3854 | 4,7197 |
| 201 | | | | | | 4,7422 |
| 202 | | | | | | 4,7646 |
| 203 | | | | | | 4,7871 |
| 204 | | | | | | 4,8096 |
| 205 | | | | | | 4,8320 |
| 206 | | | | | | 4,8545 |
| 207 | | | | | | 4,8770 |
| 208 | | | | | | 4,8994 |
| 209 | | | | | | 4,9219 |
| 210 | | | | | | 4,9443 |
| 211 | | | | | | 4,9668 |
| 212 | | | | | | 4,9893 |
| 213 | | | | | | 5,0117 |
| 214 | | | | | | 5,0342 |
| 215 | | | | | | 5,0566 |
| 216 | | | | | | 5,0791 |
| 217 | | | | | | 5,1016 |
| 218 | | | | | | 5,1240 |
| 219 | | | | | | 5,1465 |
| 220 | | | | | | 5,1689 |
| 221 | | | | | | 5,1914 |
| 222 | | | | | | 5,2139 |
| 223 | | | | | | 5,2363 |
| 224 | | | | | | 5,2588 |
| 225 | | | | | | 5,2813 |
| 226 | | | | | | 5,3037 |
| 227 | | | | | | 5,3262 |
| 228 | | | | | | 5,3486 |
| 229 | | | | | | 5,3711 |
| 230 | | | | | | 5,3936 |
| 231 | | | | | | 5,4160 |
| 232 | | | | | | 5,4385 |
| 233 | | | | | | 5,4609 |
| 234 | | | | | | 5,4834 |
| 235 | | | | | | 5,5059 |
| 236 | | | | | | 5,5283 |
| 237 | | | | | | 5,5508 |
| 238 | | | | | | 5,5732 |
| 239 | | | | | | 5,5957 |
| 240 | | | | | | 5,6182 |
| 241 | | | | | | 5,6406 |
| 242 | | | | | | 5,6631 |
| 243 | | | | | | 5,6855 |
| 244 | | | | | | 5,7080 |
| 245 | | | | | | 5,7305 |

| | 43-y column 5 | 44 | 45-SumSq 5 | 46-Iterations 5 | 47-a-Final 5 | 48-b-Final 5 |
|---|---|---|---|---|---|---|
| 197 | 0,3303 | | 0,1194 | 9,0000 | 1,0036 | 0,2389 |
| 198 | 0,3285 | | 0,1194 | 10,0000 | 1,0036 | 0,2389 |
| 199 | 0,3268 | | 0,1194 | 10,0000 | 1,0036 | 0,2389 |
| 200 | 0,3250 | | 0,1194 | 9,0000 | 1,0036 | 0,2389 |
| 201 | 0,3233 | | | | | |
| 202 | 0,3216 | | | | | |
| 203 | 0,3198 | | | | | |
| 204 | 0,3181 | | | | | |
| 205 | 0,3164 | | | | | |
| 206 | 0,3147 | | | | | |
| 207 | 0,3130 | | | | | |
| 208 | 0,3114 | | | | | |
| 209 | 0,3097 | | | | | |
| 210 | 0,3080 | | | | | |
| 211 | 0,3064 | | | | | |
| 212 | 0,3048 | | | | | |
| 213 | 0,3031 | | | | | |
| 214 | 0,3015 | | | | | |
| 215 | 0,2999 | | | | | |
| 216 | 0,2983 | | | | | |
| 217 | 0,2967 | | | | | |
| 218 | 0,2951 | | | | | |
| 219 | 0,2935 | | | | | |
| 220 | 0,2920 | | | | | |
| 221 | 0,2904 | | | | | |
| 222 | 0,2888 | | | | | |
| 223 | 0,2873 | | | | | |
| 224 | 0,2858 | | | | | |
| 225 | 0,2842 | | | | | |
| 226 | 0,2827 | | | | | |
| 227 | 0,2812 | | | | | |
| 228 | 0,2797 | | | | | |
| 229 | 0,2782 | | | | | |
| 230 | 0,2767 | | | | | |
| 231 | 0,2752 | | | | | |
| 232 | 0,2737 | | | | | |
| 233 | 0,2723 | | | | | |
| 234 | 0,2708 | | | | | |
| 235 | 0,2694 | | | | | |
| 236 | 0,2679 | | | | | |
| 237 | 0,2665 | | | | | |
| 238 | 0,2651 | | | | | |
| 239 | 0,2637 | | | | | |
| 240 | 0,2622 | | | | | |
| 241 | 0,2608 | | | | | |
| 242 | 0,2594 | | | | | |
| 243 | 0,2581 | | | | | |
| 244 | 0,2567 | | | | | |
| 245 | 0,2553 | | | | | |

| | 49-x column 6 | 50-y column 6 | 51 | 52-SumSq 6 | 53-Iterations 6 | 54-a-Final 6 |
|---|---|---|---|---|---|---|
| 197 | 4,6523 | 0,2264 | | 0,0865 | 10,0000 | 0,9419 |
| 198 | 4,6748 | 0,2248 | | 0,0865 | 10,0000 | 0,9419 |
| 199 | 4,6973 | 0,2233 | | 0,0865 | 9,0000 | 0,9419 |
| 200 | 4,7197 | 0,2218 | | 0,0865 | 10,0000 | 0,9419 |
| 201 | 4,7422 | 0,2203 | | | | |
| 202 | 4,7646 | 0,2187 | | | | |
| 203 | 4,7871 | 0,2172 | | | | |
| 204 | 4,8096 | 0,2157 | | | | |
| 205 | 4,8320 | 0,2143 | | | | |
| 206 | 4,8545 | 0,2128 | | | | |
| 207 | 4,8770 | 0,2113 | | | | |
| 208 | 4,8994 | 0,2099 | | | | |
| 209 | 4,9219 | 0,2085 | | | | |
| 210 | 4,9443 | 0,2070 | | | | |
| 211 | 4,9668 | 0,2056 | | | | |
| 212 | 4,9893 | 0,2042 | | | | |
| 213 | 5,0117 | 0,2028 | | | | |
| 214 | 5,0342 | 0,2014 | | | | |
| 215 | 5,0566 | 0,2000 | | | | |
| 216 | 5,0791 | 0,1986 | | | | |
| 217 | 5,1016 | 0,1973 | | | | |
| 218 | 5,1240 | 0,1959 | | | | |
| 219 | 5,1465 | 0,1946 | | | | |
| 220 | 5,1689 | 0,1933 | | | | |
| 221 | 5,1914 | 0,1919 | | | | |
| 222 | 5,2139 | 0,1906 | | | | |
| 223 | 5,2363 | 0,1893 | | | | |
| 224 | 5,2588 | 0,1880 | | | | |
| 225 | 5,2813 | 0,1867 | | | | |
| 226 | 5,3037 | 0,1854 | | | | |
| 227 | 5,3262 | 0,1842 | | | | |
| 228 | 5,3486 | 0,1829 | | | | |
| 229 | 5,3711 | 0,1816 | | | | |
| 230 | 5,3936 | 0,1804 | | | | |
| 231 | 5,4160 | 0,1792 | | | | |
| 232 | 5,4385 | 0,1779 | | | | |
| 233 | 5,4609 | 0,1767 | | | | |
| 234 | 5,4834 | 0,1755 | | | | |
| 235 | 5,5059 | 0,1743 | | | | |
| 236 | 5,5283 | 0,1731 | | | | |
| 237 | 5,5508 | 0,1719 | | | | |
| 238 | 5,5732 | 0,1707 | | | | |
| 239 | 5,5957 | 0,1696 | | | | |
| 240 | 5,6182 | 0,1684 | | | | |
| 241 | 5,6406 | 0,1672 | | | | |
| 242 | 5,6631 | 0,1661 | | | | |
| 243 | 5,6855 | 0,1650 | | | | |
| 244 | 5,7080 | 0,1638 | | | | |
| 245 | 5,7305 | 0,1627 | | | | |

| | 55-b-Final 6 |
|---|---|
| 197 | 0,3064 |
| 198 | 0,3064 |
| 199 | 0,3064 |
| 200 | 0,3064 |
| 201 | |
| 202 | |
| 203 | |
| 204 | |
| 205 | |
| 206 | |
| 207 | |
| 208 | |
| 209 | |
| 210 | |
| 211 | |
| 212 | |
| 213 | |
| 214 | |
| 215 | |
| 216 | |
| 217 | |
| 218 | |
| 219 | |
| 220 | |
| 221 | |
| 222 | |
| 223 | |
| 224 | |
| 225 | |
| 226 | |
| 227 | |
| 228 | |
| 229 | |
| 230 | |
| 231 | |
| 232 | |
| 233 | |
| 234 | |
| 235 | |
| 236 | |
| 237 | |
| 238 | |
| 239 | |
| 240 | |
| 241 | |
| 242 | |
| 243 | |
| 244 | |
| 245 | |

| | 1 | 2 | 3 | 4 | 5 | 6 |
|---|---|---|---|---|---|---|
| 246 | | | | | | |
| 247 | | | | | | |
| 248 | | | | | | |
| 249 | | | | | | |
| 250 | | | | | | |
| 251 | | | | | | |
| 252 | | | | | | |
| 253 | | | | | | |
| 254 | | | | | | |
| 255 | | | | | | |
| 256 | | | | | | |
| 257 | | | | | | |

| | 7-x column | 8-y column | 9 | 10-SumSq | 11-Iterations | 12-a-Final |
|---|---|---|---|---|---|---|
| 246 | 5,7529 | 0,3454 | | | | |
| 247 | 5,7754 | 0,3439 | | | | |
| 248 | 5,7979 | 0,3423 | | | | |
| 249 | 5,8203 | 0,3408 | | | | |
| 250 | 5,8428 | 0,3392 | | | | |
| 251 | 5,8652 | 0,3377 | | | | |
| 252 | 5,8877 | 0,3362 | | | | |
| 253 | 5,9102 | 0,3347 | | | | |
| 254 | 5,9326 | 0,3332 | | | | |
| 255 | 5,9551 | 0,3317 | | | | |
| 256 | 5,9775 | 0,3302 | | | | |
| 257 | 6,0000 | 0,3287 | | | | |

| | 13-b-Final | 14-x column 1 | 15-y column 1 | 16 | 17-SumSq 1 | 18-Iterations 1 |
|---|---|---|---|---|---|---|
| 246 | | 5,7529 | 0,3609 | | | |
| 247 | | 5,7754 | 0,3594 | | | |
| 248 | | 5,7979 | 0,3578 | | | |
| 249 | | 5,8203 | 0,3563 | | | |
| 250 | | 5,8428 | 0,3548 | | | |
| 251 | | 5,8652 | 0,3533 | | | |
| 252 | | 5,8877 | 0,3518 | | | |
| 253 | | 5,9102 | 0,3503 | | | |
| 254 | | 5,9326 | 0,3488 | | | |
| 255 | | 5,9551 | 0,3473 | | | |
| 256 | | 5,9775 | 0,3459 | | | |
| 257 | | 6,0000 | 0,3444 | | | |

| | 19-a-Final 1 | 20-b-Final 1 | 21-x column 2 | 22-y column 2 | 23 | 24-SumSq 2 |
|---|---|---|---|---|---|---|
| 246 | | | 5,7529 | 0,1845 | | |
| 247 | | | 5,7754 | 0,1833 | | |
| 248 | | | 5,7979 | 0,1820 | | |
| 249 | | | 5,8203 | 0,1808 | | |
| 250 | | | 5,8428 | 0,1795 | | |
| 251 | | | 5,8652 | 0,1783 | | |
| 252 | | | 5,8877 | 0,1771 | | |
| 253 | | | 5,9102 | 0,1759 | | |
| 254 | | | 5,9326 | 0,1747 | | |
| 255 | | | 5,9551 | 0,1735 | | |
| 256 | | | 5,9775 | 0,1723 | | |
| 257 | | | 6,0000 | 0,1711 | | |

| | 25-Iterations 2 | 26-a-Final 2 | 27-b-Final 2 | 28-x column 3 | 29-y column 3 | 30 |
|---|---|---|---|---|---|---|
| 246 | | | | 5,7529 | 0,1097 | |
| 247 | | | | 5,7754 | 0,1088 | |
| 248 | | | | 5,7979 | 0,1078 | |
| 249 | | | | 5,8203 | 0,1069 | |
| 250 | | | | 5,8428 | 0,1060 | |
| 251 | | | | 5,8652 | 0,1051 | |
| 252 | | | | 5,8877 | 0,1042 | |
| 253 | | | | 5,9102 | 0,1033 | |
| 254 | | | | 5,9326 | 0,1024 | |
| 255 | | | | 5,9551 | 0,1015 | |
| 256 | | | | 5,9775 | 0,1006 | |
| 257 | | | | 6,0000 | 0,0997 | |

| | 31-SumSq 3 | 32-Iterations 3 | 33-a-Final 3 | 34-b-Final 3 | 35-x column 4 | 36-y column 4 |
|---|---|---|---|---|---|---|
| 246 | | | | | 5,7529 | 0,1097 |
| 247 | | | | | 5,7754 | 0,1088 |
| 248 | | | | | 5,7979 | 0,1078 |
| 249 | | | | | 5,8203 | 0,1069 |
| 250 | | | | | 5,8428 | 0,1060 |
| 251 | | | | | 5,8652 | 0,1051 |
| 252 | | | | | 5,8877 | 0,1042 |
| 253 | | | | | 5,9102 | 0,1033 |
| 254 | | | | | 5,9326 | 0,1024 |
| 255 | | | | | 5,9551 | 0,1015 |
| 256 | | | | | 5,9775 | 0,1006 |
| 257 | | | | | 6,0000 | 0,0997 |

| | 37 | 38-SumSq 4 | 39-Iterations 4 | 40-a-Final 4 | 41-b-Final 4 | 42-x column 5 |
|---|---|---|---|---|---|---|
| 246 | | | | | | 5,7529 |
| 247 | | | | | | 5,7754 |
| 248 | | | | | | 5,7979 |
| 249 | | | | | | 5,8203 |
| 250 | | | | | | 5,8428 |
| 251 | | | | | | 5,8652 |
| 252 | | | | | | 5,8877 |
| 253 | | | | | | 5,9102 |
| 254 | | | | | | 5,9326 |
| 255 | | | | | | 5,9551 |
| 256 | | | | | | 5,9775 |
| 257 | | | | | | 6,0000 |

| | 43-y column 5 | 44 | 45-SumSq 5 | 46-Iterations 5 | 47-a-Final 5 | 48-b-Final 5 |
|---|---|---|---|---|---|---|
| 246 | 0,2539 | | | | | |
| 247 | 0,2526 | | | | | |
| 248 | 0,2512 | | | | | |
| 249 | 0,2499 | | | | | |
| 250 | 0,2485 | | | | | |
| 251 | 0,2472 | | | | | |
| 252 | 0,2459 | | | | | |
| 253 | 0,2446 | | | | | |
| 254 | 0,2433 | | | | | |
| 255 | 0,2420 | | | | | |
| 256 | 0,2407 | | | | | |
| 257 | 0,2394 | | | | | |

| | 49-x column 6 | 50-y column 6 | 51 | 52-SumSq 6 | 53-Iterations 6 | 54-a-Final 6 |
|---|---|---|---|---|---|---|
| 246 | 5,7529 | 0,1616 | | | | |
| 247 | 5,7754 | 0,1605 | | | | |
| 248 | 5,7979 | 0,1594 | | | | |
| 249 | 5,8203 | 0,1583 | | | | |
| 250 | 5,8428 | 0,1572 | | | | |
| 251 | 5,8652 | 0,1561 | | | | |
| 252 | 5,8877 | 0,1550 | | | | |
| 253 | 5,9102 | 0,1540 | | | | |
| 254 | 5,9326 | 0,1529 | | | | |
| 255 | 5,9551 | 0,1519 | | | | |
| 256 | 5,9775 | 0,1508 | | | | |
| 257 | 6,0000 | 0,1498 | | | | |

| | 55-b-Final 6 |
|---|---|
| 246 | |
| 247 | |
| 248 | |
| 249 | |
| 250 | |
| 251 | |
| 252 | |
| 253 | |
| 254 | |
| 255 | |
| 256 | |
| 257 | |

**Nonlinear Regression - Dynamic Fitting** **czwartek, maj 07, 2026, 15:43:01**

**Data Source: Copy of Data 1 in Notebook1 half life calculations.JNB**
**Equation: Exponential Decay; Single, 2 Parameter**
f = a*exp(-b*x)

**Dynamic Fit Options:**

| | |
|---|---|
| Total Number of Fits | 200 |
| Maximum Number of Iterations | 200 |

**Parameter Ranges for Initial Estimates:**

| | Minimum | Maximum |
|---|---|---|
| a | -0,8343 | 2,5030 |
| b | 0,0000 | 2,6039 |

**Summary of Fit Results:**

| | |
|---|---|
| Converged | 100,0% |
| Singular Solutions | 1,0% |

**Results for the Overall Best-Fit Solution:**

| R | Rsqr | Adj Rsqr | Standard Error of Estimate |
|---|---|---|---|
| 0,9504 | 0,9033 | 0,8791 | 0,1036 |

| | Coefficient | Std. Error | t | P |
|---|---|---|---|---|
| a | 1,0070 | 0,1120 | 8,9945 | 0,0008 |
| b | 0,3854 | 0,0782 | 4,9284 | 0,0079 |

**Analysis of Variance:**

Analysis of Variance:

| | DF | SS | MS |
|---|---|---|---|
| Regression | 2 | 1,6795 | 0,8398 |
| Residual | 4 | 0,0429 | 0,0107 |
| Total | 6 | 1,7225 | 0,2871 |

Corrected for the mean of the observations:

| | DF | SS | MS | F | P |
|---|---|---|---|---|---|
| Regression | 1 | 0,4010 | 0,4010 | 37,3499 | 0,0036 |
| Residual | 4 | 0,0429 | 0,0107 | | |
| Total | 5 | 0,4440 | 0,0888 | | |

**Statistical Tests:**

**Normality Test (Shapiro-Wilk)** Passed (P = 0,2984)

W Statistic= 0,8861 Significance Level = <0,0001

**Constant Variance Test** Passed (P = 0,0600)

**Fit Equation Description:**

```
[Variables]
x = col(6)
y = col(4)
reciprocal_y = 1/abs(y)
reciprocal_ysquare = 1/y^2
'Automatic Initial Parameter Estimate Functions
F(q)=if(size(x)>1; if(total(abs(y))>0; ape(x;log(abs(y));1;0;1); -306); 0)
asign(q)=if(mean(q)>=0;1;-1)
[Parameters]
a = if(F(0)[1]< 307; if(F(0)[1]>-307; asign(y)*10^F(0)[1]; asign(y)*10^(-307)); asign(y)*10^307) "Auto {{previous: 1,00703}} {{MinRange: 0,99}} {{MaxRange: 1}}
b = if(x50(x;y)-min(x)=0; 1;  -ln(,5)/(x50(x;y)-min(x)))  "Auto {{previous: 0,385366}} {{MinRange: 0}} {{MaxRange: 1}}
[Equation]
f = a*exp(-b*x)
fit f to y
"fit f to y with weight reciprocal_y
"fit f to y with weight reciprocal_ysquare
[Constraints]
b>0
[Options]
tolerance=1e-10
stepsize=1
iterations=200
```

Number of Iterations Performed = 11
